# Neurogenomic landscape associated with status-dependent cooperative behavior

**DOI:** 10.1101/2022.10.05.510848

**Authors:** Peri E. Bolton, T. Brandt Ryder, Roslyn Dakin, Jennifer L. Houtz, Ignacio T. Moore, Christopher N. Balakrishnan, Brent M. Horton

## Abstract

The neurogenomic mechanisms mediating male-male reproductive cooperative behaviors are unknown. We leveraged extensive transcriptomic and behavioral data on a neotropical bird (*Pipra filicauda*) that perform cooperative courtship displays to understand these mechanisms. Cooperative display is modulated by testosterone, it promotes cooperation in non-territorial birds, but suppresses cooperation in territory holders. We sought to understand the neurogenomic underpinnings of these three related traits: social status, cooperative behavior, and testosterone phenotype. To do this, we profiled gene expression in 10 brain nuclei spanning the social decision-making network (SDMN), and two key endocrine tissues that regulate social behavior. We associated gene expression with each bird’s behavioral and endocrine profile derived from three years of repeated measures taken from free-living birds in the Ecuadorian Amazon. We found distinct landscapes of constitutive gene expression were associated with social status, testosterone phenotype, and cooperation, reflecting the modular organization and engagement of neuroendocrine tissues. Sex-steroid and neuropeptide signaling appeared to be important in mediating status-specific relationships between testosterone and cooperation, suggesting shared regulatory mechanisms with male aggressive and sexual behaviors. We also identified differentially regulated genes involved in cellular activity and synaptic potentiation, suggesting multiple mechanisms underpin these genomic states. Finally, we identified SDMN-wide gene expression differences between territorial and floater males that could form the basis of “status-specific” neurophysiological phenotypes, potentially mediated by testosterone and growth hormone. Overall, our findings provide new, systems-level insights into the mechanisms of cooperative behavior and suggest that differences in neurogenomic state are the basis for individual differences in social behavior.

## Introduction

Cooperation is the synchronized behavior(s) of individuals to produce a net fitness benefit for one or more individuals (Taborsky & Taborsky, 2015). An individual’s decisions about cooperation are based on immediate social stimuli, past experiences, developmental effects, and genetics. These effects contribute to repeatable, among-individual variation in behaviors – a behavioral phenotype – where some individuals are more cooperative than others. To understand the mechanisms of cooperation, it is important to understand not only the elements of neurophysiological signal processing, but also the genetic and neurophysiological components that underpin behavioral phenotypes. Understanding these mechanisms enables us to link how genetic and neurophysiological architecture can constrain and facilitate the evolution of cooperation (Hofmann et al., 2014; Soares et al., 2010; Taborsky & Taborsky, 2015). However, the mechanisms underlying the diversity of behaviors related to cooperation remain unclear (Díaz-Muñoz et al., 2014; Kasper et al., 2017). In particular, our understanding of the neuroendocrine mechanisms underlying seemingly paradoxical male-male reproductive cooperative behaviors is only emerging (Díaz-Muñoz et al., 2014; DuVal & Goymann, 2011; Jones & DuVal, 2021; Loveland et al., 2021; Ryder et al., 2020; Vernasco et al., 2020).

Differences in cooperative behavioral phenotype are partly achieved through pleiotropic yet modular effects of hormone signaling systems (Cox, 2020; Ketterson et al., 2009). Hormonal pleiotropy mediates phenotypic integration, whereby multiple traits covary with a hormonal signal (Cox, 2020; Ketterson et al., 2009). Trait independence is facilitated by variation in hormone receptor expression, where tissues across the body are modular in their response to hormone signals, enabling diverse and flexible responses to hormonal and social signals (Ketterson et al., 2009; Lipshutz et al., 2019). In particular, sex steroid concentration and receptor distribution in neural tissues are important mediators of social behaviors, including cooperation (Kasper et al., 2017; Soares et al., 2010). The neural substrates modulating social behaviors are conserved across vertebrates, in the social decision making network (SDMN) (O’Connell & Hofmann, 2011). The SDMN comprises reciprocally interconnected brain nuclei rich in steroid receptors, steroidogenic enzymes, and steroid-modulated neuropeptide systems (Goodson, 2005; Newman, 1999; O’Connell & Hofmann, 2011). This modular and dynamic network activates different combinations of brain regions in response to different social contexts (Goodson, 2005; Newman, 1999). Therefore, the neurological response in a single nucleus is not reflective of all systems engaged in modulating a given behavior. Consistent variation among individuals in neuroendocrine gene expression across the SDMN likely underlies repeatable individual differences in social behavior and behavioral phenotype. For example, variation in the expression of genes involved in sex-steroid signaling pathways across SDMN nuclei can modify the male brain’s sensitivity to testosterone, resulting in distinct behavioral responses to a specific hormone signal. Indeed, recent research suggests that constitutive gene expression or neurogenomic state, characterized by co-expressed gene suites in behaviorally relevant brain regions, is associated with behavioral phenotypes (Antunes et al., 2021; Kabelik et al., 2021; Lattin et al., 2021).

For the first time, we characterize constitutive gene expression across the neuroendocrine system in relation to variation in a male-male cooperative behavior. The wire-tailed manakin (*Pipra filicauda*) is a neotropical lek breeding bird in which unrelated males perform cooperative displays to attract females (Figure 1A). Territorial and subordinate non-territorial (floater) males form long-term display partnerships and form complex social networks (Figure 1A, B) (Ryder et al., 2008; Ryder, Blake, et al., 2011). Status and cooperation are directly linked to fitness, whereby territory holders with more display partners sire more offspring, and floaters with more partners are more likely to ascend to territorial status (Ryder et al., 2008, 2009). Individuals have a repeatable ‘testosterone phenotype’ where territorial males tend to have higher testosterone than floaters (Figure 1D), and circulating testosterone has a status-specific effect on the cooperative display behavior (Ryder et al., 2020; Ryder, Horton, et al., 2011). Experimental and observational evidence show that higher testosterone levels are antagonistic to cooperation in territorial males but promote cooperation in floater males (Ryder et al., 2020; Vernasco et al., 2020) (Figure 1C).

**Fig. 1.**
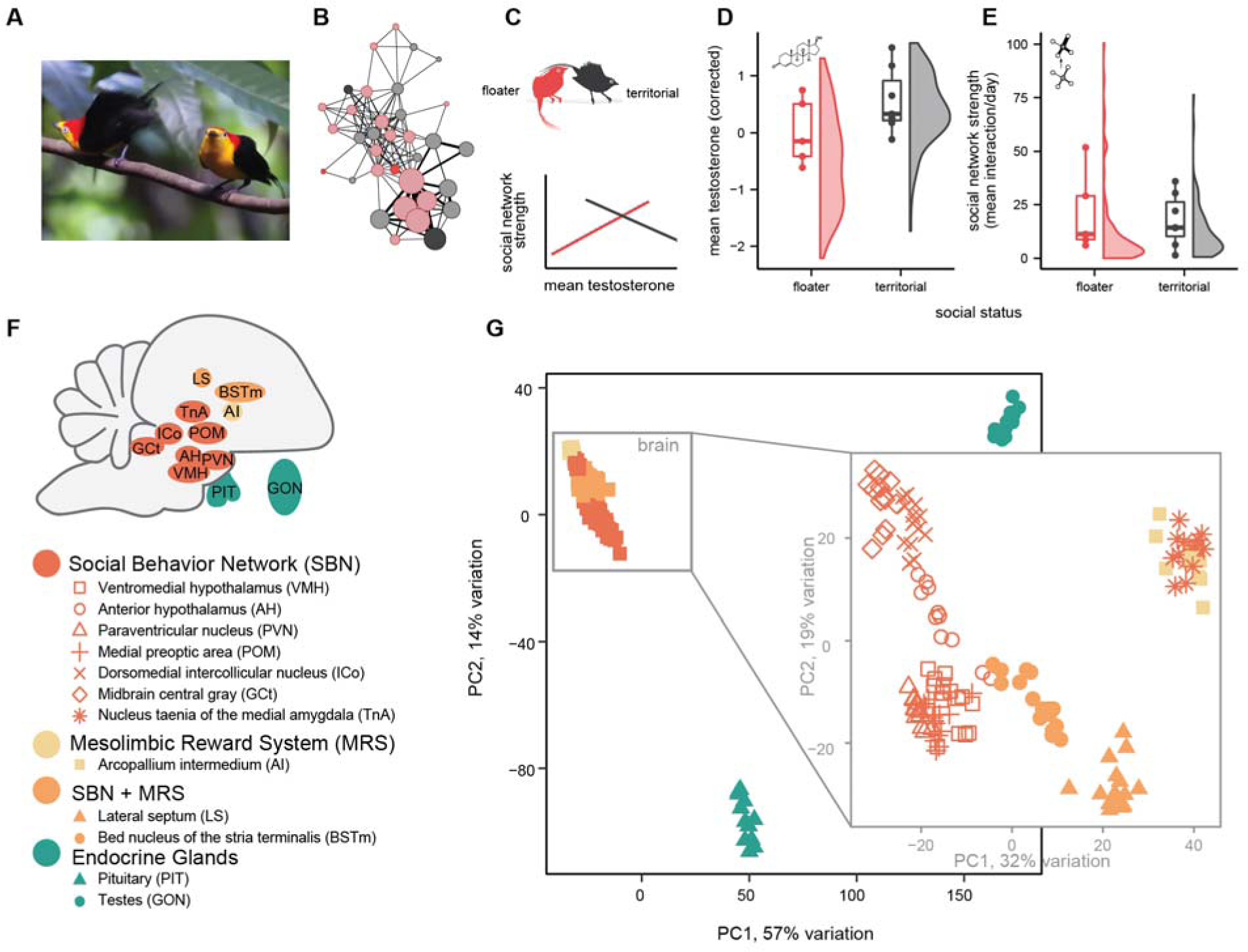
Testosterone and neuroendocrine tissues are involved in the cooperative behavior of wire-tailed manakins. A) Males engage in cooperative displays forming complex social networks. B) Example network shows single lek over one week, where node size indicates a male’s average social network strength (measure of overall time spent interacting), and edge width represents interaction frequency. Node color denotes floater (red) and territorial (gray)status, and darker colors were included in the study. C) Testosterone phenotype (mean testosterone (corrected)) influences cooperative tendency in territorial and floater males differently. D) Repeated measures of testosterone for 12 individuals (points) from a larger population (density). E) as D but proximity-data logging measures of social network strength.. F) Brain and endocrine tissues sampled for RNAseq. Structures not to scale. Colors represent functional groups (O’Connell & Hofmann, 2011). G & inset) Principal component analysis reveals distinct gene expression profiles among brain regions and endocrine tissues (colors and shapes as F). (Photo: Alice Boyle). Note: sample size N=16 for social status, N=12 for testosterone and network strength.

To explore the neurogenomic basis of the status-specific androgenic modulation of cooperative behavior in these birds, we quantify transcriptome-level gene expression in the SDMN, pituitary and testes of free-living male wire-tailed manakins in the Ecuadorian Amazon. Captive studies can obscure ecologically and evolutionary relevant variation in environmental and behavioral responses (Griffith et al., 2017; Kasper et al., 2017), and we demonstrate the exciting applicability of neuro-transcriptional methods in challenging rainforest conditions. We aim to characterize the genes and neuroendocrine pathways that modulate individual variation in cooperative behavior, social status, and testosterone phenotype as a modifier of both social status and cooperative behavior. Given the androgenic regulation of cooperative display (Ryder et al., 2020; Vernasco et al., 2020), we hypothesize that differences in gene regulation will manifest in SDNM nuclei with high density of sex-steroid receptors and endocrine tissues (Goodson, 2005; Newman, 1999), and will involve expression of candidate genes relating to sex-steroid signaling, metabolism, and neuropeptide systems (Goodson & Thompson, 2010; Kingsbury & Wilson, 2016). We anticipate brain region-specific gene expression patterns within the modular SDMN (Figure 1G), as well as systemic effects of testosterone on gene expression across the brain (Cox, 2020; Ketterson et al., 2009). We discuss our results in the context of modes of phenotypic evolution and similarities with mechanisms of other cooperative and social behaviors.

## Materials and Methods

All methods described here were approved by Smithsonian Animal Care and Use Committee (protocols 12-12,14-25,17-11), as well as the Ecuadorian Ministry of the Environment (MAE-DNB-CM-2015-0008), and samples were transported with export permit (006-016-EXP-1C-FAU-DNB/MA) and import permit (USDA APHIS 126133). Detailed methods can be found in the supplementary material, and full code from analyses in R v 4.0.2 are available on GitHub (https://periperipatus.github.io/PIFI_brain_transcriptome/) and FigShare (10.25573/data.22186516).

### Field Methods

We sampled 16 wild territorial and floater wire-tailed manakin males for RNA sequencing from Tiputini Biodiversity Station in the lowland Amazon rainforest of Ecuador (Table S1.1). Males were categorized by social status: where the territorial birds (n=9) were males in definitive plumage that held a consistent display territory over the study period, while floaters (n=7) had either definitive or pre-definitive plumage and did not occupy consistent territories within leks (Ryder et al., 2008). We used repeated testosterone sampling on 12 of these males to characterize their testosterone phenotype (Figure 1D, Supplement 1). A male’s testosterone phenotype is represented by mean of the residuals of a linear regression from repeated sampling of testosterone concentration, and this measure was previously shown to be repeatable within individuals (Supplement 1 for more detail, see also (Ryder et al., 2020)). Previous research has shown that territorial males have, on average, higher circulating testosterone than floaters (Ryder et al., 2020; Ryder, Horton, et al., 2011), and is reflected in our sampled males (Figure 1D).

Social network structure and individual variation in cooperative display were characterized using autonomous radio-telemetry and tag proximity detections within territories on a total population of 180 male manakins over multiple years (Dakin & Ryder, 2018; Ryder et al., 2012, 2020). We used this tagging data to calculate social network strength for the same 12 males with repeated testosterone measures. Social network strength is a proxy for cooperative tendency and is the time spent interacting with other males averaged over the study period (Figure 1E), and was the most repeatable within individuals (Ryder et al., 2020). This particular proxy for cooperation showed the strongest status-specific relationship with an individual’s testosterone phenotype (Ryder et al., 2020) (Figure 1C), but the overall frequency of cooperative interactions did not differ between territorial and floater males (Figure 1E).

### Microdissection and RNAseq

Methods for brain extraction, preservation, cryosectioning, microdissection, and RNA extraction followed those published previously (Horton, Ryder, et al., 2020). Pituitaries were preserved in RNAlater™ (Invitrogen) at ambient temperatures until imported into the United States. We excised tissue from 10 different nuclei involved in social behavior – nine of which are from the SDMN. The SDMN reflects the interconnectivity of two networks involved in regulating social behavior (O’Connell & Hofmann, 2011): Social Behavior Network (SBN) (Goodson, 2005; Newman, 1999) and the Mesolimbic Reward System (MRS) (O’Connell & Hofmann, 2011; Olds & Milner, 1954). Figure 1F indicates the nomenclature used and the relationships of these nuclei to the SBN and MRS. To clarify bird specific nomenclature: the arcopallium intermedium (AI) is a region that may be partly homologous the basolateral amygdala of mammals (Figure S2.5), which is hypothesized to play a role in androgen-dependent manakin display behavior (Fusani et al., 2014); and the nucleus taenia (TnA) is homologous to the mammalian medial amygdala (O’Connell & Hofmann, 2011). The paraventricular nucleus (PVN) is not yet formally recognized as part of the SDMN, despite its neuropeptide projections to nodes of the SDMN and established role as a major regulator of vertebrate social behavior (Goodson & Kingsbury, 2013; Goodson & Thompson, 2010). We additionally sampled the pituitary gland and testes immediately after the brains. Testes were handled the same as brains (Horton, Ryder, et al., 2020) and pituitaries were preserved in RNAlater™ (Invitrogen) at ambient temperatures until imported into the United States.

Microdissected tissues from both hemispheres were combined for sequencing. RNA was extracted and sequenced according to previously described methods (Horton, Ryder, et al., 2020). RNA libraries for the various tissues and field seasons were randomized amongst sequencing batches (Table S1.1-S1.2). All samples analyzed are associated with BioProject PRJNA437157, and SRR12660169-198, SRR19521260-271, SRR19521432-575.

### Gene expression analyses

Reads were aligned to the annotated *Pipra filicauda* genome (GCA_003945595.1) using splice-aware aligner STAR v2.7.5 (Dobin et al., 2013), and gene-level counts were obtained from featureCounts v2.0.1 (Liao et al., 2014). A total of 170 libraries were retained after quality filtering in each tissue (Figures S1.2-S1.4 & Figures S3.1-3.2), but the number of individuals used in each tissue varies (Table S1.1-S1.2). To further characterize tissue and brain nucleus function in this species (Horton, Ryder, et al., 2020), we characterized differences in gene expression among tissues and brain nuclei using principal components analysis in PCAtools v2.8.0 and Weighted Gene Co-Expression Analysis (WGCNA v1.71) (Blighe & Lun, 2020; Langfelder & Horvath, 2008) (Supplement 2). All analyses were conducted in R v 4.0.2 (R Core Team, 2020).

To describe the role of tissues and brain nuclei in relation to hormonal and behavioral phenotypes we analyzed differential expression using DESeq2 v1.36 (Love et al., 2014) and co-expression using WGCNA (Langfelder & Horvath, 2008) in each tissue and nucleus separately. For a subset of tissues, we had an extra four individuals characterized for social status from a previous study (Horton, Ryder, et al., 2020), for which behavioral and hormone sampling data were unavailable. Owing to this, and to not overparameterize the models that account for batch effects, we identified differentially expressed genes (DEGs) associated with each of our traits of interest (social status, testosterone phenotype and social network strength) separately. These were run as separate negative binomial generalized linear models for each trait that accounted for batch effects where necessary using DESeq2 (Love et al., 2014) (Supplement 3, Figures S3.1 - S3.2). Testosterone and social network strength were treated as continuous variables. Significant DEGs were considered to be correlated with the predictor variable, and Log_2_ Fold Change (LFC) value represents the LFC of gene expression per unit of the predictor variable. In all cases, significant DEGs were identified using the Wald Test and the default FDR corrected p-value (q<0.1). The identity of many hub genes identified by WGCNA were similar to DEGs identified byDESeq2, so for brevity and simplicity we focus mostly on the differential expression results, but the full results are presented in Supplement 5 (WGCNA Figures S5.1-S5.14) and Supplement 3 (differential expression Figures S3.3-3.43).

Due to the key role testosterone plays in modulating behavior in this species (Ryder et al., 2020), we further analyzed nine candidate genes involved in sex-hormone metabolism and signaling (Figure S4.1), as well as nine steroid sensitive neuropeptides and their receptors (Figure S4.2; Table S1.3). Candidate genes showing differential gene expression with an uncorrected p < 0.05 was considered “significant” and examined further. In addition to analyzing models of status, mean testosterone and social network strength separately, we also explored whether candidate genes were involved in mediating status-specific relationships between mean testosterone and cooperative behavior (Ryder et al., 2020) by extracting “significant” candidate genes from a final DESeq2 interaction analysis (model: expression ∼ status x mean testosterone). We primarily used this model to identify candidate genes with status-specific expression rather than discovering new DEGs due to overfitting concerns. The resulting putative gene-trait relationships were plotted (Figures S4.3-S4.23), and those with highly influential observations or R2 ≤ 0.2 were excluded. For candidates derived from the interaction effect models or genes that were associated with multiple traits, the most convincing relationship was determined using a stepdown AIC procedure (Supplement 4). Considering the smaller set of candidate genes, uncorrected p-values were deemed significant (p<0.05), but we also report the FDR corrected values.

To assess neuroanatomical constraints on gene expression patterns, we examined overlap among brain nuclei by comparing ranked gene-lists (Supplement 6). The gene-lists were ranked using -log10 transformed p-values multiplied by the expression direction. Similarity between tissue pairs was measured using Pearson correlation and rank-rank hypergeometric overlap (RRHO) test (Cahill et al., 2018).

Testosterone has pleiotropic effects, influencing gene expression in similar ways in multiple tissues (Cox, 2020; Ketterson et al., 2009). We reasoned that these effects would be reflected in shared (systemic) signatures of differential expression across brain nuclei. To explore systemic gene expression patterns, we calculated median p-values from the previous analysis and identified genes consistently differentially expressed across the entire brain (p-value < 0.05, transformed value > |1.3|). Further, we correlated eigenvalues of each principal component (PC) from the whole-brain PCA analysis (Figure 1G inset) with our variables of interest using pcatools::eigencorplot() (Blighe & Lun, 2020) (Supplement 1). Correlations with social status were validated by a linear mixed model in the nlme package v3.1 (Pinheiro et al., 2016), which included individual as a random effect and accounted for correlated batch variables (Supplement 1).

We used clusterProfiler v3.16.1(Yu et al., 2012) to determine patterns of functionality in DEGs and WGCNA modules with Gene Ontology enrichment analyses. If the total number of FDR corrected DEGs was less than 40, we used gene set enrichment analysis instead which leverages the rank order of p-values and direction of expression (Supplement 3).

## Results

As expected, gene expression analyses revealed substantial differences among tissues and brain regions (Figure 1G; Figure S2.4). These expression patterns were more similar among adjacent nuclei, forming major brain-region groupings of the amygdala, hypothalamus, and midbrain (Figure 1G). Additionally, our 18 candidate neuroendocrine genes (Table S1.3) displayed tissue and brain region-specific expression (Table 2.1, Figures S4.1-4.2).

### Social status and testosterone phenotype are associated with tissue and brain region-specific patterns of gene expression

We observed a landscape of gene expression in association with testosterone phenotype (Figure 2A) and social status (Figure 2B), with a varying number of differentially expressed genes (DEGs) or co-expression modules identified across different tissues. Similarly, across brain regions, we identified relationships between many of our 18 candidate genes and status, testosterone phenotype, or their interaction (Figure 2C). Mean testosterone was a continuous predictor in DESeq2, as such DEGs indicate up-or downregulation with a higher testosterone phenotype (Figure 2A), while DEGs identified in social status indicated up or down regulation in territorial males (Figure 2B). Here, we primarily focus on the tissues with large numbers of DEGs and significant candidate genes for the main manuscript, but the co-expression results were similar to our differential expression results, and all results are shown in Supplements 3 and 5. The number of DEGs in each tissue showed a similar pattern between testosterone phenotype and status compared to other trait comparisons (R = 0.5, p = 0.08, Figure S3.7), which was consistent across co-expression modules (Supplement 5) and candidate gene analysis (Figure 2C & Figure 3B). This similarity was expected due to the correlation between testosterone and status (Figure S1.1) (Ryder et al., 2020; Ryder, Horton, et al., 2011).

**Fig. 2.**
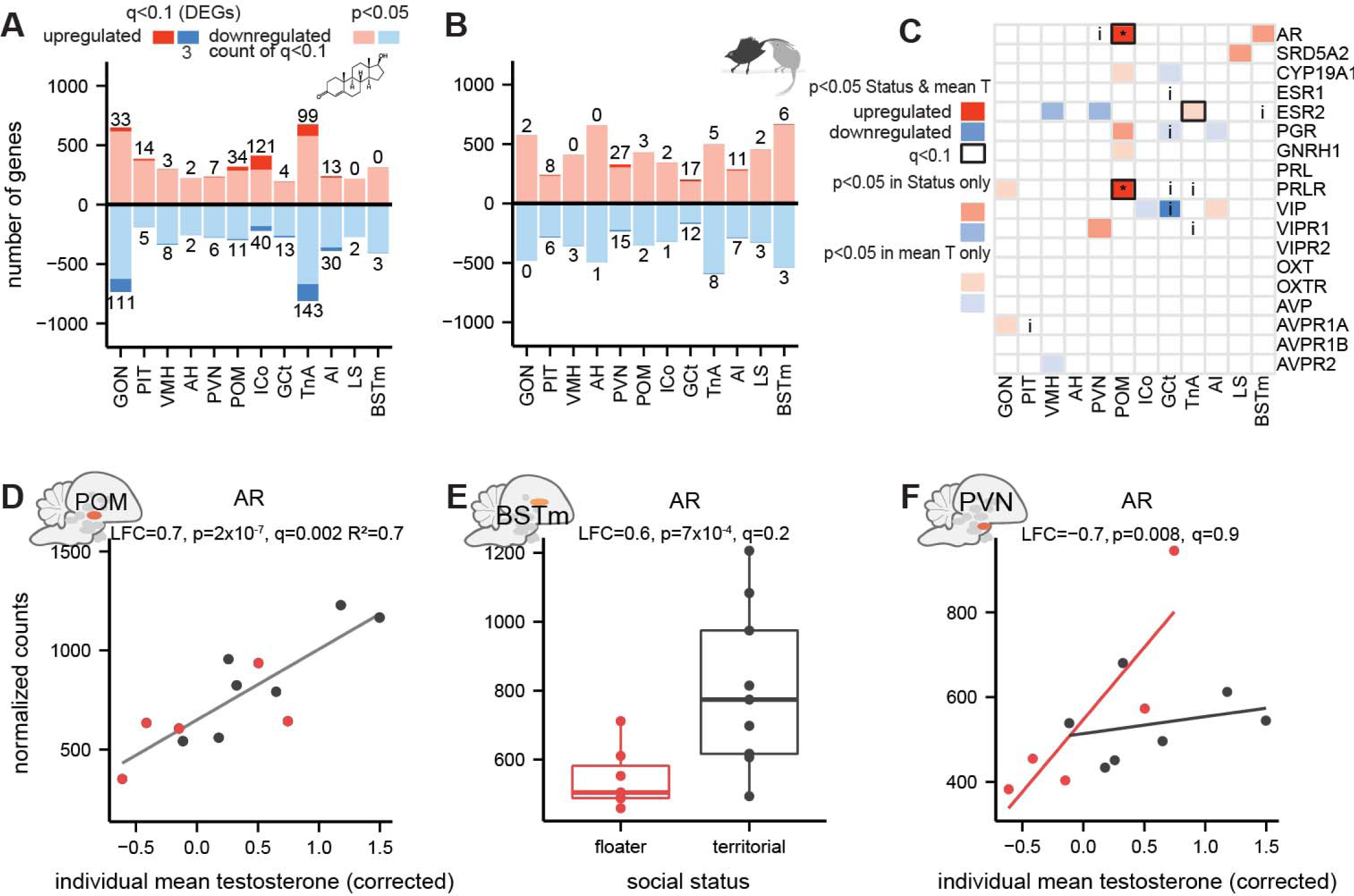
Neurogenomic landscape of social status and hormone phenotype. A) Landscape of gene expression in male wire-tailed manakins with high testosterone phenotype (mean testosterone (corrected)) relative to lower testosterone phenotype. Red bars indicate number genes upregulated in males with higher testosterone phenotype, blue indicates genes downregulated in males with higher testosterone. Darker colors and text indicate number of differentially expressed genes after FDR correction. B) Landscape of gene expression in relation to social status, where red indicates genes upregulated in territorial males. C) Summary of candidate gene analysis and filtering, where colors indicate direction of expression, intensity indicates which trait is associated, and “i” indicates there was an interaction effect, and black boxes indicate result was significant after FDR correction. D) Upregulation of AR in POM of males with higher testosterone, where point colors indicate social status (red = floater; black = territorial. Fitted line from linear regression. Subheader includes results from DESeq2: Log_2_ Fold Change (LFC), uncorrected p-value (p), and correction for multiple testing (q) and R^2^ calculated separately. E) Upregulation of AR in BSTm of territorial males. F) Status-specific expression of AR in PVN in relation to testosterone phenotype. Note: Sample sizes vary in each tissue due to quality filtering. Maximum sample size for analysis of social status was N=16, while only 12 individuals were sampled for testosterone and behavior.

**Fig. 3.**
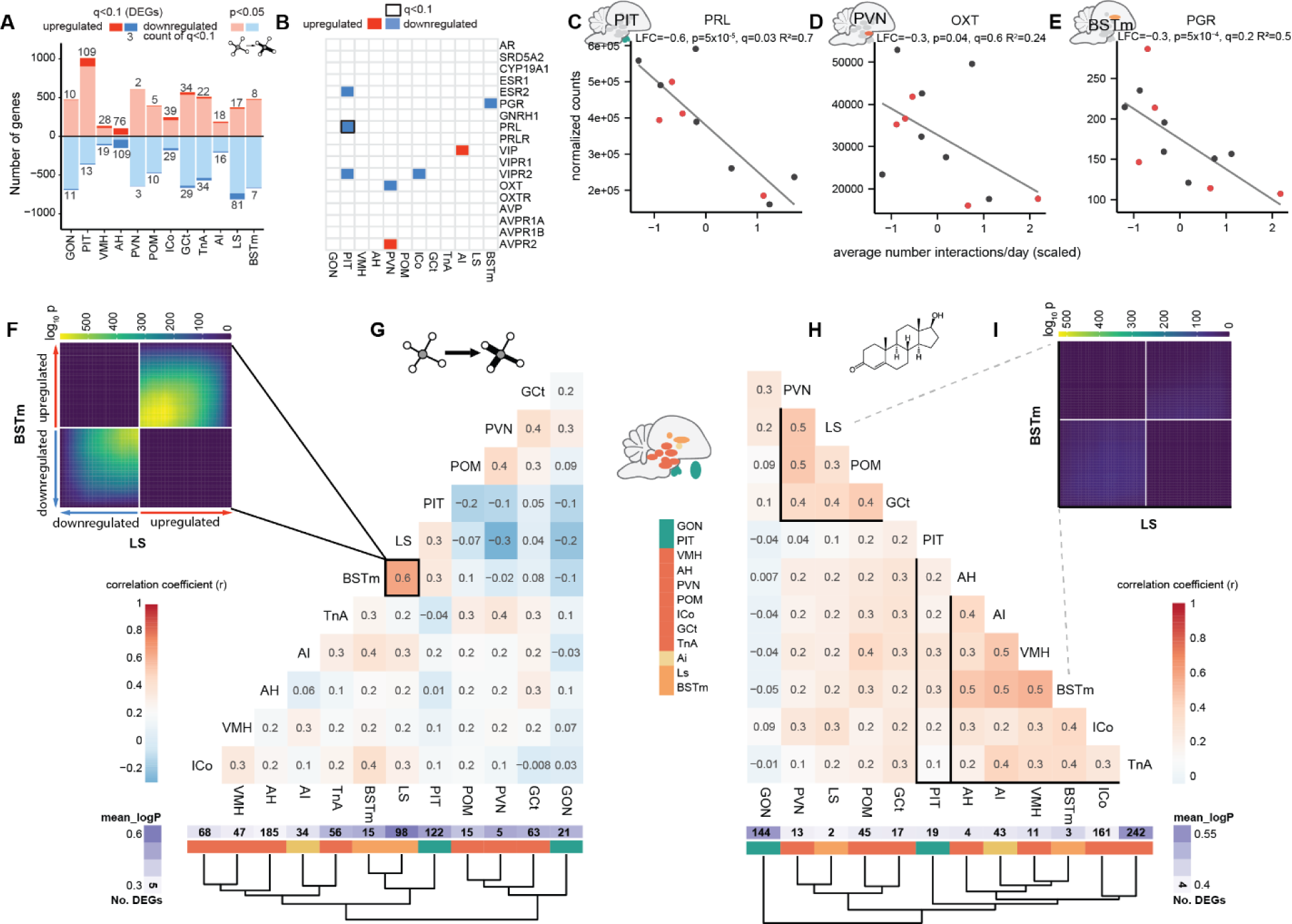
A distinct landscape of gene expression in relation to cooperative behavior. A) Gene expression landscape in male wire-tailed manakins with higher social network strength. Legend as fig. 2A. B) Summary of candidate gene analysis and filtering, where colors indicate direction of expression in in individuals with higher social network strength. Black boxes indicates significant results after FDR correction. C) Downregulation of prolactin (PRL) in the pituitary of males with higher social network strengths D) Downregulation of oxytocin (OXT) in PVN of males with higher social network strength E) Downregulation of progesterone receptor (PGR) in BSTm of males with higher social network strength. F) RRHO2 analysis shows similar gene expression profiles associated with social network strength in the LS and BSTm, G) while network strength related gene expression similarity is low among other tissues. H) Gene expression similarity among tissues in relation to testosterone phenotype follows the ontogeny of brain regions, I) No correlation between testosterone-related gene expression profiles in LS and BSTm. G,H) Pearson correlation coefficients of gene expression between tissues, represented by cell color and number. Gene expression p-values were log_10_ transformed and multiplied by direction of expression. Black boxes outline significant tissue correlation clusters based on 1000 bootstrap resamples.

The testes and TnA each had over 1,000 genes differentially expressed (at uncorrected p-value < 0.05) in association with both testosterone phenotype and status (144 and 252 genes after FDR correction q<0.01 in association with mean testosterone; 2 and 13 after FDR correction in association with social status). DEGs identified in association with testosterone phenotype in the testes were enriched for genes involved with response to corticosterone (GO:0051412) and synaptic transmission and organization (e.g., GO:0007268, GO:0050808) among others (Figure S3.20). Notable DEGs included steroidogenic acute regulatory protein (STAR, LFC = −1.1, p = 2×10^−7^, q = 0.0003, R^2^ = 0.6) and candidate gene AVPR1A (LFC = 0.8, p = 5×10^−5^, q = 0.01, R^2^ = 0.7; Figure S4.4). Note that Log_2_ Fold Change (LFC) represents a change in gene expression per unit of the predictor variable. These transcriptional differences do not correspond to significant differences in testis size in relation to testosterone phenotype (Figure S3.44, Table S3.1), but territorial males have larger testes (p=0.04, Table S3.1)

Among the 252 DEGs associated with testosterone phenotype in TnA, there was non-significant enrichment for genes involved in neural crest migration (GO:0001755), learning and memory (GO:0007613, GO:0007612), and long-term synaptic potentiation (e.g., GO:0060291) (Figure S3.28). Notably, the candidate gene estrogen receptor 2 (ESR2; aka Erβ) was upregulated in TnA of males with higher testosterone (Figure 2C; LFC = 0.8, p = 0.001, q = 0.08, R^2^ = 0.4). Other notable DEGs in TnA included upregulation of immediate early gene JUN, also a ‘coral1’ hub gene (module membership (MM) = 0.93; LFC = 0.61, p = 0.0004, q = 0.05) and corticotropin releasing hormone receptor 2 (CRHR2, LFC = 1, q = 0.006, R^2^ = 0.5).

Not all nuclei showed similar patterns between status and testosterone phenotype. For example, ICo of the midbrain showed 161 DEGs associated with testosterone phenotype after FDR correction but did not have substantial association with social status (3 DEGs). Enriched GO terms in ICo were primarily related to motor proteins and cilia (e.g., GO:0003341, Figure S3.26). Intriguingly, although the POM did not reveal a high number of DEGs in association with testosterone phenotype (45), our analysis revealed differential expression for compelling gene ontology categories and candidate neuroendocrine genes (Figure S3.25, Figure S4.10, Figure S5.7). Notably, males with higher testosterone exhibited upregulation of candidate gene androgen receptor (AR) in POM (Figure 2D). The ‘floralwhite’ module, which comprised many upregulated DEGs, was significantly associated with both testosterone phenotype and, to a lesser extent, social status (Figure S5.6-5.7). This module showed enrichment for biological processes related to G protein-coupled receptor signaling pathways (GO:0007186) and neuropeptide signaling pathway (GO:0007218), with statistically non-significant enrichment for steroid receptor signaling (GO:0030518), learning and memory (GO:0007611), and nervous system development (GO:0007399) (Figure S3.7). Indeed, many of the shared hub-genes and DEGs interact with candidate sex hormone pathways, such as prolactin releasing hormone (PRLH, MM = 0.91, LFC = 2.4, q = 5×10^−5^, R^2^ = 0.6, Figure S3.25A), growth regulating estrogen receptor binding 1 (GREB1, MM = 0.86, LFC = 0.6, q = 3×10^−5^, R^2^ = 0.7, Figure S3.25A), and gonadotropin releasing hormone 1 (GNRH1, MM = 0.76, LFC = 0.4, p = 0.006, q = 0.2, R^2^ = 0.4). Other notable ‘floralwhite’ hub genes include the candidate gene progesterone receptor (PGR, MM = 0.81) which was also associated with social status (Figure 2C; LFC = 0.4 p = 0.006, q = 0.7) and serotonin receptor 5A (HTR5A, MM = 0.87, LFC = 0.4, p = 0.04, q = 0.6). Candidate gene aromatase (LOC113993669/CYP19A1) was also upregulated in males with higher testosterone phenotype (Figure 2C, LFC = 0.6, p = 0.02, q = 0.4, R^2^ = 0.5), but was found in the ‘lightcoral’ co-expression module (MM= 0.86).

Despite overall similarity in gene expression associations with testosterone phenotype and status, certain brain regions exhibited stronger associations with either status or the interaction between status and testosterone phenotype (Figure 2B). Likewise, the 18 candidate neuroendocrine genes also showed varying associations with status or the interaction across different nuclei (Figure 2C). PVN of the hypothalamus showed the most DEGs associated with social status after FDR correction (42). These DEGs were enriched for cytoskeletal movement genes involved in intracellular transport and cilia (e.g., GO:0003341, GO:0061640), and non-significant enrichment for cholesterol biosynthesis (GO:0006695, GO:0045540) (Figure S3.12B). Candidate gene expression also revealed status-specific associations with testosterone phenotype in the PVN, that is a statistical interaction between status and testosterone phenotype (Figure 2C). In particular, AR expression was positively correlated with testosterone phenotype in floater males, but not in territorial males (Figure 2F; LFC = −0.7, p = 0.008, q = 0.9). The PVN of territorial males also showed higher VIPR1 (Figure S4.8; LFC = 0.3, p = 0.02, q = 0.4) and lower ESR2 expression (LFC = −0.6, p = 0.006, q = 0.2). In BSTm, a region that bridges the SDMN and MRS, there were more than 1000 DEGs, but only nine after FDR correction. Among these, territorial males showed higher AR expression compared to floater males (Figure 2E; LFC = 0.6, p = 0.0007, q =0.2). Further, testosterone metabolism gene 5-alpha reductase (SRD5A2) was upregulated in the LS of territorial males (Figure S4.20; LFC = 0.7, p = 0.0005, q = 0.4). In the TnA, the VIP system was implicated in status-specific responses to testosterone, with VIPR1 upregulated in association with testosterone in territorial males (Figure S4.17; LFC = 1, p = 0.04 q = 0.7).

GCt was a hotbed for gene expression associated with social status and status × testosterone phenotype interaction (Figure 1B, Figure S3.6). In relation to social status, gene set enrichment analysis revealed a significant downregulation of genes involved in synapse assembly (GO:0051965), response to estradiol (GO:0032355), as well as upregulation of genes related to RNA processing (e.g., GO:0000184, GO:0006396), and translation (e.g., GO:0006413, GO:0006412) (Figure S3.15). Candidate neuroendocrine genes showed status specific expression in association with testosterone phenotype in this region including progesterone receptor (PGR, Figure S4.15; LFC = 1.9, p= 0.003, q = 0.1), estrogen receptor 1 (ESR1; aka Erα; LFC = 1.1 p = 0.04, q = 0.4), prolactin receptor (PRLR; LFC = 0.7, p = 0.01, q = 0.2), and vasoactive intestinal peptide (VIP; Figure S4.15; LFC = 1.6, q = 0.005, q = 0.1). Further, aromatase was downregulated with higher testosterone phenotype in the GCt (LFC = −1.3, p = 0.002, q = 0.3, R^2^ = 0.4).

### Cooperative behavior is associated with a distinct gene expression landscape

Social network strength (the number of cooperative interactions between individuals) varies widely among territorial and floater males (Figure 1E), representing a key dimension of social behavior in this species. We predicted that the associations between gene expression and cooperative behavior would be different from those observed for social status and testosterone phenotype. Indeed, across tissues we found a distinct landscape of constitutive gene expression in relation to cooperative behavior (Figure 3A, Figure S3.5), as well as among candidate genes (Figure 3B). AH had the highest number of DEGs associated with a male’s network strength, with non-significant enrichment for genes involved in RNA metabolism (e.g., GO:0043928) and regulation of translation (e.g., GO:0017148), development (e.g., GO:0061061, GO:0042472) and synaptic potentiation (e.g., GO:0007274) (Figure S3.35). The pituitary also showed an abundance of differentially expressed genes associated with male’s network strength (109 genes after FDR correction q < 0.1), with non-significant enrichment for actin filament organization (GO:0007015), neural differentiation (e.g., GO:0031103, GO:0045664) and the cell cycle (e.g., GO:0043065, GO:0071158.) (Figure S3.33B). Within the pituitary, candidate genes prolactin (PRL) (Figure 3C; LFC = −0.6, p=5×10^−5^ q = 0.03, R^2^ = 0.7), ESR2 (Figure S4.5; LFC = −0.7, p = 0.04, q = 0.5, R^2^ = 0.5) and VIPR2 (Figure S4.5; LFC = −0.4, p = 0.008, q = 0.3, R^2^ = 0.4) were downregulated in highly cooperative males. Intriguingly, in the PVN there was a weak negative correlation between the expression of the candidate gene oxytocin (OXT) and network strength (Figure 3D; LFC = −0.3, p = 0.04, q = 0.6, R^2^ = 0.2). Furthermore, although there was less representation of sex-steroid related pathways in the candidate gene analysis (Figure 2B), PGR was downregulated in males with higher social network strength in BSTm (Figure 2E; LFC = - 0.3, p = 5×10^−4^, q = 0.2, R^2^ = 0.5).

Many genes in the LS were differentially expressed in association with an individual’s social network strength (98 genes after FDR correction q < 0.1). Among these, the top non-significant terms were broadly involved in development (e.g., GO:0048538, GO:0048593), and neural organization (e.g., GO:0099175, GO:0050771) (Figure S3.42). No candidate genes were linked to social network strength in LS. Although few genes were differentially expressed in BSTm related to social network strength, RRHO analysis showed that LS and BSTm shared similar expression associations with strength (Cahill et al., 2018) (Figure 3G, F). In contrast, LS and BSTm showed limited similarity in gene expression related to testosterone phenotype (Figure 4H, I) or social status (Figure S6.1 & S6.4), suggesting their unique role in modulating cooperative behavior. Pairwise correlations of gene expression associated with testosterone phenotype aligned with neuroanatomical similarity (Figure 3H) possibly reflecting similarities in pathways related to sex-steroid signaling (Figure 2C), but LS/BSTm similarity was the only neuroanatomical signal in pairwise gene expression associated with strength.

**Fig. 4:**
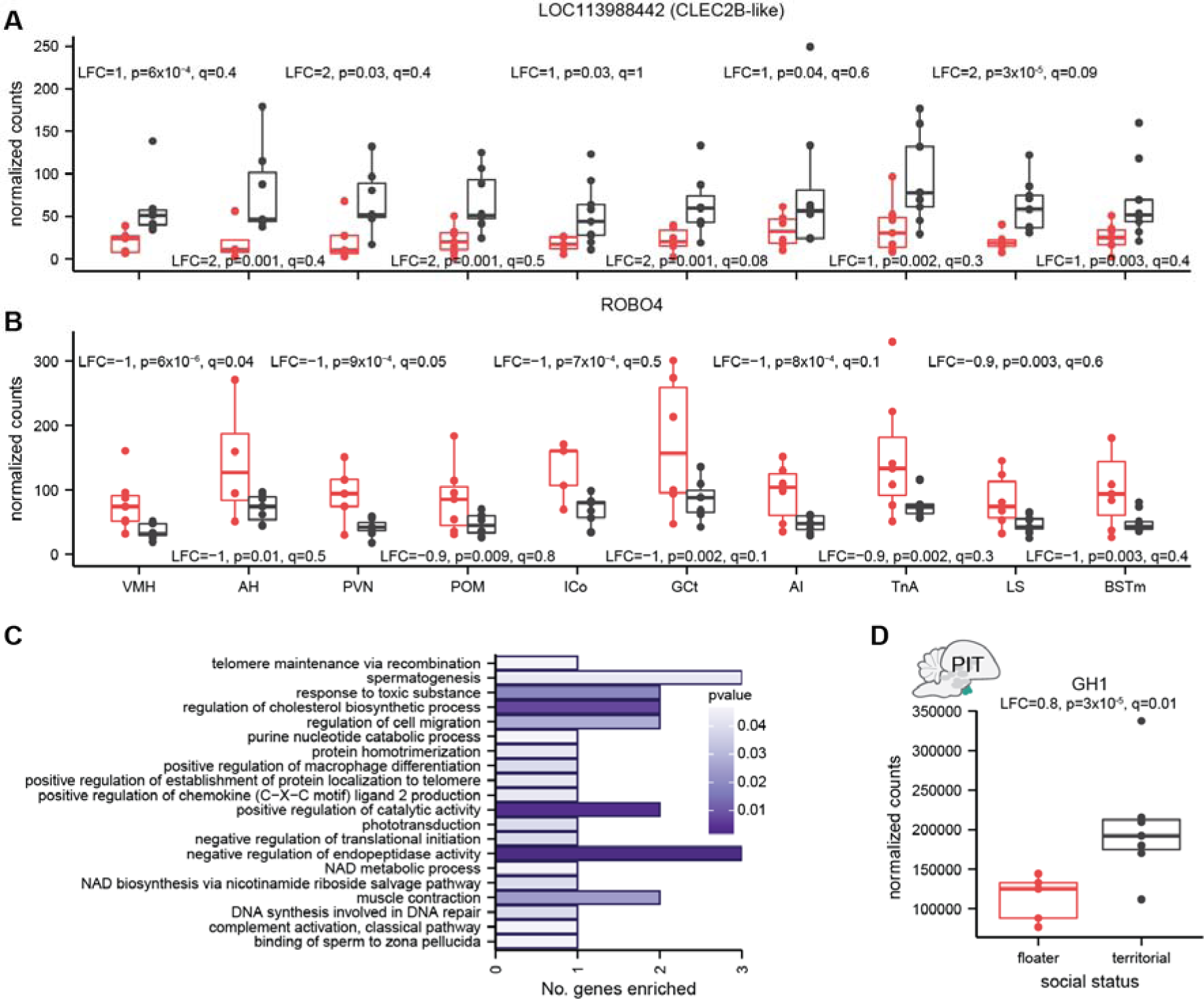
Global neural gene expression differences between social classes. A) Immune gene CLEC2B-like was consistently upregulated in the brains of territorial individuals. B) Developmental gene ROBO4 was consistently downregulated in the brains of territorial males. C) Enriched GO categories among genes that were consistently up- or downregulated between territorial and floater males, where bar color indicates uncorrected p-value. D) Upregulation of growth hormone transcript in the pituitary of territorial males could explain differences between status classes.

### Testosterone phenotype and social status are associated with systemic gene expression

We reasoned that testosterone, with its pleiotropic effects and role in social status in manakins (Ryder et al., 2020) would have systemic impacts on gene expression, resulting in similar expression patterns across multiple regions. Social status and testosterone phenotype showed the highest number of consistently differentially expressed genes across tissues (60 and 66 genes with median uncorrected p < 0.05 respectively, Figure S6.2), while social network strength had only 10 consistently differentially expressed genes (Figure S6.3 & Table S6.3). Further, PC5 of the whole brain analysis was significantly associated with social status in a linear mixed model, accounting for batch confounds and repeated sampling of individuals (Figure S2.1-S2.3). There was overlap between genes and GO categories among the consistently differentially regulated genes with social status and testosterone phenotype (Tables S6.1-S6.2), as well as with genes associated with PC5. But hereon we focus on genes related to social status for simplicity.

The top two consistently differentially regulated genes were upregulation of the immune system gene CLEC2B in the brains of territorial males (Figure 4A) and downregulation of the development gene ROBO4 (Figure 4B). Enrichment analysis reflected the functional categories of these genes, although no significant terms were found after FDR correction (Figure 4C). There was also evidence for global differential regulation of telomere maintenance genes (Figure 4C), with downregulation of POLD-1-like (LOC114000574) and upregulation of RAD51 in territorial males. Further, there was evidence for global regulation of cholesterol biosynthesis (Figure 4C), and glutathione transferases (Figure S2.3, Table S6.2), both of which are involved in steroid hormone biosynthesis. These findings of global expression differences among status classes could be partially explained by the upregulation of growth hormone (GH1) in the pituitary of territorial males (Figure 4D; LFC = 0.8, p = 3×10^−5^, q = 0.01), which remained consistent in a model accounting for the interaction between social status and mean T (LFC = 0.9, p = 6×10^−5^, q = 0.05).

## Discussion

Using the most comprehensive transcriptome-level sampling of the SDMN to-date (Antunes et al., 2021; Bentz, George, et al., 2021; Bentz, Niederhuth, et al., 2021; Kabelik et al., 2021; Lopes & König, 2020), our findings highlight both modular and integrative neurogenomic organization underlying individual variation in cooperative behavior. The relative roles of each are central to the development and expression of any complex trait (Cox, 2020; Ketterson et al., 2009; Lipshutz et al., 2019).

### Different landscapes of gene expression with each trait reflect modular organization of the neuroendocrine system

Modularity is reflected in how different brain nuclei, endocrine tissues, and genetic pathways are associated with the expression of different traits. These neurogenomic landscapes, characterized by the upregulation of neuroendocrine genes in some brain regions and downregulation in others, enabling differential sensitivity to testosterone and other signaling molecules among SDMN nuclei. These neurogenomic states can be likened to the activation of immediate early genes (IEGs) across the SDMN after a social stimulus (Goodson, 2005; Newman, 1999). In that ‘landscape’ model, it is the pattern of IEG expression across multiple brain nuclei that correlates with behavioral responses, rather than the up or downregulation of particular genes within a single nucleus. This original model considered this landscape of IEG expression (and later neuropeptides) in the context of proximate responses to social stimuli. Our findings extend this model in two ways: by considering transcriptomic variation and how long-term neurogenomic states underlie individual variation in social behavior.

In the landscape of constitutive gene expression in male wire-tailed manakins, most brain regions exhibit differences in gene expression but the brain regions with higher numbers of DEGs or candidate gene expression may be more important for mediating certain traits. Some brain regions showed associations between gene expression and both social status and testosterone phenotype (e.g., POM) or only testosterone phenotype (e.g., TnA, testes). This may be because these regions are involved in testosterone mediated traits irrespective of social status (O’Connell & Hofmann, 2012). Meanwhile, others appeared to be more related to social status and status specific relationships with testosterone phenotype (e.g., PVN, GCt), or an individual’s strength of cooperative behavior (e.g., PIT, LS). Across brain regions, there were variations in the identity and function of genes, but certain functions consistently emerged, including neural development, synaptic potentiation or organization, and genes related to cellular metabolism or transcription/translation regulation. These results suggest that individual variation is mediated by neurogenomic states that encode synaptic modifications and connectivity (mechanism of memory formation), larger scale shifts in cellular activity, and neural structure (Clayton et al., 2019; George et al., 2020). These mechanisms may interact with variation in elements of sex-steroid signaling that we investigated in detail (*sensu “*neuroendocrine action potential” (Clayton et al., 2019)).

Our findings provide strong evidence of modularity in the expression of steroid-related genes, particularly in relation to social status, testosterone phenotype, or their interaction. We found evidence that status-specific regulation of behavior could be modulated directly by testosterone through AR in multiple nuclei (e.g., POM, PVN, BSTm), which is consistent with previous findings of AR expression linked to display performance in golden-collared manakins (B. A. Schlinger et al., 2013). Further, the conversion of testosterone to estradiol by the enzyme aromatase and subsequent binding to estrogen receptors is a key pathway in the regulation of testosterone-dependent male behaviors (B. Schlinger & Balthazart, 2013), and we found evidence for the role of aromatase and estrogen signaling role in multiple nuclei.

The co-upregulation of steroid related genes in the POM of males with high testosterone phenotype (and territorial males) point to this region’s role as a potent steroid regulator. The POM is involved in regulating male aggression, sexual behavior and parental care (O’Connell & Hofmann, 2011), and stimulates gonadal testosterone production through gonadotropin releasing hormone (*GnRH*). Interestingly, we observed increase in GNRH1 transcription, which suggest enhanced steroid production in the testes. Our finding of co-upregulation of AR and GNRH1 may seem contradictory due to the negative regulatory effects of androgens and AR on *GnRH* (Brayman et al., 2012). However, these may not interact at the cell population level or may show different temporal patterns of translation. The activation of POM and its steroid receptor concentrations have been associated with social status differences and social status ascension cichlid fish (Maruska et al., 2013), and mice (Lee et al., 2022). In the same cichlid species, different preoptic cell types, including homologs for PVN, showed higher activation in one of the two partners in cooperative territorial defense (Weitekamp & Hofmann, 2017). Testosterone can link the POM to the Hypothalamic Pituitary Adrenal (HPA) axis via the PVN (Williamson et al., 2010) – another important endocrine regulator of social status (DuVal & Goymann, 2011; Goymann & Wingfield, 2004; Jones & DuVal, 2021). In our study, the PVN showed status-specific gene expression many genes, including AR, and could be an important mediator of status-specific regulation of cooperative display. Thus, close linkages between the PVN and POM (Williamson et al., 2010), suggest these nuclei could be involved in multiple elements of manakin behavior depending on cell types, social contexts and timescales investigated.

Our results also highlight the importance of neural VIP expression in modulating multiple aspects of social behavior (Horton, Michael, et al., 2020; Kingsbury & Wilson, 2016), as this neuropeptide system exhibited widespread differential expression patterns across all traits examined. An intriguing finding in our study is the potential involvement of VIP and other steroid receptors in mediating status-specific responses to testosterone in GCt. Although tyrosine hydroxylase (TH) neurons in GCt have been associated with courtship behaviors (Ben-Tov et al., 2021; Goodson et al., 2009), our novel findings point to the association between VIP and courtship behaviors in the GCt (Kingsbury & Wilson, 2016). GCt coordinates social stimuli and downstream motor processes that control behavior. For example, in songbirds, GCt links POM to the song control system and thereby mediates androgen-dependent singing behavior (Haakenson et al., 2020). Given that cooperative display in wire-tailed manakins requires coordinated physical (and vocal displays in cooperative *Chiroxiphia* manakins (Trainer et al., 2002)), the status-specific differential expression of genes for steroid receptors and steroid-sensitive neuropeptides in GCt suggest that this SDMN region may play a unique role in mediating status-specific relationships between testosterone and cooperative behavior.

The differential gene expression observed in TnA suggests its role as a hub for regulating testosterone-mediated traits. TnA, the avian homolog of the medial amygdala (O’Connell & Hofmann, 2011), integrates sensory information and regulates diverse social behaviors (Raam & Hong, 2021). Our results may reflect persistent activation of this region, based on the upregulation of IEGs such as JUN and SYT7 in males with high testosterone phenotype, as well as differential regulation of genes involved in long-term synaptic potentiation (Clayton, 2000; Clayton et al., 2019; Marrone et al., 2008; O’Connell et al., 2012). Further, our results implicate potential estrogenic regulation in this region via upregulation of ESR2 in males with higher testosterone phenotype. Estrogenic pathways through ESR1 have been shown to regulate status-specific aggression in birds (Horton et al., 2014; Merritt et al., 2020) and prosocial behavior in prairie voles (Cushing et al., 2008). Although ESR2 is not well studied in birds, it may function in the medial amygdala of mammals to regulate social recognition (Lymer et al., 2018), and sexual regulation (Nakata et al., 2016), and is associated with higher social rank in the forebrains of cichlid fish (Burmeister et al., 2007). Finally, the fish homolog of TnA appears to mediate aggression and cooperation in cooperative territory defense in an actor-specific manner (Weitekamp et al., 2017; Weitekamp & Hofmann, 2017), supporting a role for modulating cooperation in wire-tailed manakins.

There were distinct patterns of gene expression across sampled tissues in association with social network strength (time spent interacting with other males), and some of these signals suggest derivation of cooperative display from aggressive behaviors (Díaz-Muñoz et al., 2014). Many genes were differentially expressed in AH in association with strength, a region primarily linked to regulation of aggression through AVP and VIP (Horton, Michael, et al., 2020; O’Connell & Hofmann, 2011). Although these were not differentially expressed in AH, we found a weak negative relationship between OXT expression and social network strength in PVN. This negative relationship is contradictory to oxytocin’s role in promoting sociality (as in humans (Feldman, 2012). The relationship observed in the wire-tailed manakin suggests that oxytocin inhibits cooperation, further supporting the hypothesis that cooperation is derived from aggressive or dominance displays in this species. However, nonapeptides (like OXT and AVP) have complex and sometimes species-specific relationships with affiliative behavior and social status (Goodson & Thompson, 2010; Lee et al., 2022; Maruska et al., 2022). Thus, further research is needed to clarify the role of nonapeptides in this species.

LS showed many differentially expressed genes associated with cooperation, and highly cooperative individuals may show differences in neural structure based on the enriched GO terms. Patterns of expression in BSTm were similar, but weaker based on fewer statistically significant DEGs. BSTm has neural projections into LS, and vasopressin (AVP) and VIP neurons in the BSTm and receptors for these neuropeptides in the LS are associated with affiliative behavior and gregariousness in birds (Kelly et al., 2011; Kelly & Goodson, 2013; Kingsbury & Wilson, 2016). This suggests cooperative display is regulated by some of the same conserved brain regions as other types of affiliative behavior. Those studies focused on the acute processing of social information in these nuclei. In contrast, our study suggests that constitutive differences in gene expression and neural structure in these regions may underlie individual variation in cooperative tendencies, potentially affecting their excitability in response to social information.

Our results reveal a variety of mechanisms that could contribute to individual variation in cooperative behavior, and its status-specific modulation by testosterone. We identified multiple forms of neuroplasticity are involved, through steroid receptor variation, synaptic potentiation, neural development, as well as cellular activity (Clayton et al., 2019). Indeed, testosterone manipulations are known to induce changes in neuroanatomy in behaviorally relevant brain nuclei (Kabelik et al., 2008), suggesting a direct role for testosterone-mediated neural growth and flexibility in modulating male-male cooperation (Soares et al., 2010). Our results provide promising avenues for further study on the mechanisms underlying cooperative behavior, such as examination of neuroanatomical features and the use of hormonal or other pharmaceutical manipulations and RNAi technologies to examine causal relationships between neural gene expression and behavior (e.g., (Merritt et al., 2020)).

### Brain-wide expression patterns reflect the role of testosterone as a phenotypic integrator

The pleiotropic effects of testosterone are predicted to lead to similarities in gene expression among tissues, supporting its role as a mediator of phenotypic integration. The extent of gene expression similarities and differences among tissues may reflect the degree of integration and independence (or modularity), that can either constrain or facilitate adaptive evolution (Ketterson et al., 2009; Lipshutz et al., 2019). We identified similarities in testosterone associated gene expression between brain regions that broadly recapitulated tissue ontology. Additionally, we identified consistent (systemic) gene expression differences throughout the brain, predominantly associated with social status and testosterone phenotype. However, systemic results did not explain as much variation in overall neural gene expression compared with certain personality traits (Kabelik et al., 2021; Lattin et al., 2021), suggesting that the brain region-specific patterns may be more important in modulating male wire-tailed manakin behavior. The association between social status and metabolic, immune, and telomere gene ontology categories reflect differences in allostatic load among status classes in male-male cooperative systems (Goymann & Wingfield, 2004; Jones & DuVal, 2021; Vernasco et al., 2021). The differential global regulation of telomeres may explain the relative stability of telomere length in territorial wire-tailed manakin males (Vernasco et al., 2021), suggesting metabolic profiles associated with territoriality. Cooperation itself may also have metabolic costs, as more cooperative male manakins have been shown to have shorter telomeres (Vernasco et al., 2021), and social bond strength has been associated with signatures of energy metabolism and stress (Anderson et al., 2022; Simons et al., 2022).

Our results highlight two additional hormones, prolactin and growth hormone, that may have a system-wide role in the status-specific modulation of cooperative display behavior. Transcripts for these hormones were differentially expressed in the pituitary, and may reflect organism wide metabolic profiles. The anterior pituitary is the production site for both hormones, and thus gene expression patterns are likely good predictors for circulating levels of those hormones and thus signal strength. PRL transcript was downregulated in more cooperative males. Prolactin is best known for its role in promoting parental care behaviors, including in cooperative breeders (Schoech et al., 1996), but emerging studies suggest other functions such as inhibiting aggression in cooperative breeders (Gilbert et al., 2022; Medger et al., 2019) and promoting nest defense behavior (Mohamed et al., 2016). The negative relationship observed appears contradictory to the role of prolactin in promoting prosocial behavior, and again, may reflect the origin of these cooperative displays from aggressive interactions (Díaz-Muñoz et al., 2014). In addition, growth hormone transcript (GH1) showed higher expression in territorial males. This was surprising because ascension in social rank in the cooperatively displaying manakins is associated with age (DuVal, 2007; McDonald, 1989; Ryder et al., 2008) and GH1 expression is higher in older territorial males. However, in *Anolis* lizards, stimulation of the growth hormone axis is associated with increased testosterone that enables phenotypic divergence between males and females (Cox et al., 2017). In manakins growth-hormone could thus be involved in testosterone production and have subtle effects on multiple systems enabling the status-specific regulation of behaviors. Status differences in GH1 expression may provide an explanation for the consistent patterns of neural development and protein synthesis observed in our results (Sonksen, 2006). These expression differences could reflect the heightened metabolic demands placed on territorial males during cooperative displays, involving enhanced muscle performance and the regulation of oxidative stress (Hu et al., 2019). The neuroethological role of growth hormone is poorly understood, but it may be linked to stress (Greenwood & Landon, 1966; McCormick et al., 1998), and has been shown to increase aggression (Johansson et al., 2004; Jönsson et al., 1998; Matte, 1981). While the growth hormone inhibition system (via somatostatin) is an androgen-independent modulator of aggression and social status in cichlid fish (Hofmann & Fernald, 2000; Trainor & Hofmann, 2006), and natal androgen treatment increased somatostatin receptor expression in zebra finches (Bentz, Niederhuth, et al., 2021). Thus, our results suggest that the growth hormone axis may play a role in the status-specific regulation of wire-tailed manakin cooperative displays, where increased GH and androgens in territorial males potentially inhibit cooperation by promoting aggression. Future work on social behavior should consider the interplay between growth hormone, somatostatin, sex steroids, and glucocorticoids to gain new insights into their roles in behavioral regulation.

## Conclusion

Our study reveals correlations between gene regulation, social status, cooperation, and testosterone phenotype across the male wire-tailed manakin’s SDMN and HPG axis, involving a plurality of gene functions, including sex-steroid and neuropeptide signaling. This landscape of gene expression speaks to the modular nature of behavioral mechanisms (Goodson, 2005; Lipshutz et al., 2019; Newman, 1999) and highlights the continuum between phenotypic integration and modularity (Ketterson et al., 2009). Unlike previous studies focusing on limited candidate genes or specific brain regions, our work is the most comprehensive examination of patterns across multiple nuclei in the SDMN. While our findings are correlative, they provide valuable insights into the nuclei, genomic pathways and endocrine mechanisms underlying understudied male-male cooperative behaviors, and its status-specific regulation in this species. Although questions remain on the acute mediators of cooperative display behavior in the wire-tailed manakin, we demonstrate the power and flexibility of a system-wide approach, and our findings generate hypotheses for further experiments to implicate causality. We also contribute to the emerging concept that constitutive gene expression landscapes across the brain are associated with consistent behavioral phenotypes (Antunes et al., 2021; Kabelik et al., 2021; Lattin et al., 2021). We propose that landscapes of constitutive gene expression, comprising multiple genes across many tissues and brain regions, form the foundation of consistent inter-individual differences in behavior, upon which flexible and acute responses to social stimuli are built.

## Acknowledgements

We want to acknowledge long hours of fieldwork in Ecuador by Ben Vernasco and Camilo Alfonso-Cuta that made the behavioral foundation of this work. Thank you to them, and the staff at Tiputini Biodiversity Station. Thank you to Eric Shuppe and to Matthew Fuxjager for use of laboratory resources. Thank you to David Clayton, Julie George, Sarah London, and Barney Schlinger for helpful discussions and manuscript reviewing, and Vanessa González for advice on data analysis.

## Data Accessibility

Raw read data are available associated with BioProject PRJNA437157, and all code used to conduct analyses is available at (doi:10.25573/data.22186516) or https://github.com/periperipatus/PIFI_brain_transcriptome

## Author contributions

TBR, BMH, ITM, and CNB designed and funded the research, TBR, RD, BMH, ITM, JLH collected data and performed field data analysis, PEB led the analysis and writing, CNB and BMH provided supervision. All authors contributed to review and editing of the manuscript.

