## Supplementary Material for "Neurogenomic landscape associated with status-dependent cooperative behavior"

### 1. Sampling & RNAseq

#### Methods & Results

##### Sample Collection & Behavioral Observations

Male wire-tailed manakins (*Pipra filicauda*) were studied for behavior on their leks at the Tiputini Biodiversity Station in the Orellana province of eastern Ecuador as part of a larger study on the role of testosterone on cooperative lekking behavior over three field seasons: 2015-2016, 2016-2017, and 2017-2018 (Horton *et al.* 2020; Ryder *et al.* 2020). Social and lek attendance behaviors were recorded using an automated telemetry system and verified by human observation (Ryder *et al.* 2012; Ryder *et al.* 2020; Vernasco *et al.* 2020). Up to three times per field season, males were captured in mist nets and blood sampled for circulating testosterone (T). Capture and blood sampling was conducted on leks between 6:00 - 11:00 am, outside the behavioral telemetry observational period (Ryder *et al.* 2020), and without reference to last observed behaviors. Time spent in the net can have a subtle but significant effect on plasma testosterone (Vernasco *et al.* 2019). Thus, nets were monitored with GoPro cameras to record capture and blood sampling times, and these times were used to statistically correct testosterone measures (see below) (Vernasco *et al.* 2019; Ryder *et al.* 2020).

A subset of males (n=16) of known social status were subsequently euthanized for RNAseq analysis of neural and endocrine tissues. These males were characterized as a territorial male (n=9) or a floater male (n=7); floaters included birds with definitive or pre-definitive plumage. For the 12 individuals with behavioral observations from telemetry, we extracted multiple testosterone (T) and behavioral variables from the full dataset (Ryder *et al.* 2012; Dakin and Ryder 2018; Ryder *et al.* 2020). Previous work on this species has shown that a male’s mean testosterone is associated with his social behavior, so we included the mean of each individual’s suite of corrected T measures in our analyses, as per (Ryder *et al.* 2020). The corrected T was the residuals from a linear regression of log-transformed T that accounts for season, date, time of day, and time spent in net, we then took the arithmetic mean of these residuals for each measurement to calculate a male’s mean testosterone (mean_T) (Vernasco *et al.* 2019; Ryder *et al.* 2020). Finally, we also included the average telemetry observations for strength of cooperative behavior (Strength.all_study), which represented the number of cooperative interactions an individual engaged in per day averaged over the entire study period (Ryder *et al.* 2020). We included strength as the metric of sociality because it was the most repeatable measurement of cooperation and best associated with circulating T phenotype (Ryder *et al.* 2020). Status, individual T phenotype (mean_T), and cooperative behavior (Strength.all_study) all covary (Ryder *et al.* 2020), but the only significant correlation in our subsample of the population was a positive relationship between cooperative behavior and mean T phenotype in floater status males (Figure S1.1).

Four males (two territorial and two floater) collected during the 2015-16 field season were not characterized for social behavior or testosterone, but they served to establish and validate now published protocols for brain extraction and tissue preservation in the field, and microdissection and RNA extraction in the lab (Horton *et al.* 2020) (Table S1.1).

Methods for brain extraction, tissue preservation, cryosectioning, microdissection, and RNA extraction followed those published in (Horton *et al.* 2020). Briefly, birds were euthanized by decapitation near the site of capture immediately after blood sampling for testosterone. Whole brains were extracted within 4-6 minutes of sacrifice and immediately frozen in powdered dry ice. Brains were kept frozen on dry ice in the field (< 4 hours) until transferred into a 21-day liquid nitrogen-charged Arctic Express™ dry shipper (Thermo Scientific) where they were stored at cryogenic temperatures until imported into the United States. Once in the United States, brains were stored at -80°C until cryosectioning, microdissection and RNA extraction. The pituitary gland and testes were extracted immediately after the brain. Testes were frozen, transported, and stored in the same manner as brains. Pituitaries were preserved in RNAlater™ (Invitrogen) at ambient temperatures until imported into the United States, after which excess RNAlater solution was removed before storing pituitaries at -80°C until RNA extraction. To minimize batch effects, cryosectioning, microdissection, and RNA extraction from tissues collected in the latter two field seasons (2016-2017 and 2017-2018) were conducted at the same time, where the order in which an individual’s tissues were processed was randomized with respect to male social status and year.

For each bird we excised tissue from 10 different brain nuclei. Eight of these nuclei are nodes of the highly interconnected Social Behavior Network (SBN) of the vertebrate brain (Newman 1999; Goodson *et al.* 2005). The SBN nuclei included the ventromedial hypothalamus (VMH), anterior hypothalamus (AH), medial preoptic area (POM), nucleus taenia (TnA), two midbrain regions, medial bed nucleus of the stria terminalis (BSTm), and lateral septum (LS). The TnA is considered by numerous authors to be the avian homologue of the mammalian medial amygdala (O’Connell and Hofmann 2011), although efforts to further define putative subdivisions of the avian arcopallium and homologies to mammalian amygdalar regions are ongoing (Mello *et al.* 2019). Here, we sampled the region of the brain most commonly recognized as TnA in avian studies, and which corresponds to the region referred to as the medial ventral arcopallium (AMV) by Mello *et al.* 2019. For the midbrain, we sampled both the midbrain central gray (GCT) and dorsomedial intercollicular nucleus (ICo), which collectively appear to be homologous to the mammalian periaqueductal gray (PAG) (Kingsbury *et al.* 2011). Two of the SBN nuclei sampled, BSTm and LS, are also nodes of the Mesolimbic Reward System (MRS), reflecting the interconnectivity of the two networks, both of which play key roles in the regulation of vertebrate social behavior (O’Connell and Hofmann 2011). We also sampled the arcopallium intermedium (AI), a region that may be, in part, homologous the basolateral amygdala of mammals, a node of the MRS (O’Connell and Hofmann 2011), which is hypothesized to play a role in androgen-dependent manakin display behavior(Fusani *et al.* 2014). Finally, we sampled the paraventricular nucleus (PVN), a region that is not yet formally recognized as part of the SBN despite its neuropeptide projections to nodes of the SBN and established role as a major regulator of vertebrate social behavior (Goodson and Thompson 2010; Goodson and Kingsbury 2013).

Methods for the microdissection of VMH, POM, TnA, LS, BSTm, ICo, and AI are described in Horton et al. (2020); methods for the additional regions sampling in this study are as follows. We sampled GCt, as delineated by (Kingsbury *et al.* 2011), by taking 0.5 mm diameter tissue punches from the left and right hemispheres in three or four consecutive sections. We sampled AH, as delineated by (Goodson *et al.* 2012), by taking 0.35mm punches from the left and right hemispheres of two to three consecutive sections. Because the PVN, as delineated by (Goodson *et al.* 2005) is located at the midline of the brain, it was sampled by taking 0.5 mm punches centered on the midline to simultaneously capture the left and right portions of the region in two to three consecutive sections. In addition to the 10 brain nuclei, the pituitary gland (PIT) and testes (GON) were extracted and sequenced to represent the full hypothalamic-pituitary-gonadal (HPG) axis. For a graphical representation of the tissues sampled see Figure 1G, with a full list of individuals collected in Table S1.1 and tissues in Table S1.2.

##### RNA sequencing and filtering

RNA was extracted and sequenced according to previously described methods (Horton *et al.* 2020). RNA from the gonads, pituitary, and 10 brain regions were prepared as individual TruSeq Stranded mRNA Sample Prep Kit libraries, making a total of 186 mRNA libraries(Table S1.1 & S1.2, and [GitHub](https://github.com/periperipatus/PIFI_brain_transcriptome/blob/main/data_unfiltered/Samplekey_SRA_accessions.csv) data). RNA libraries for the various tissues and field seasons were randomized amongst sequencing batches (Table S1.1 & S1.2, and [GitHub](https://github.com/periperipatus/PIFI_brain_transcriptome/blob/main/data_unfiltered/Samplekey_SRA_accessions.csv) data). A selection of samples from the original study on brain gene expression in male wire-tailed manakins (Horton *et al.* 2020) were included in this library preparation process (see [GitHub](https://github.com/periperipatus/PIFI_brain_transcriptome/blob/main/data_unfiltered/Samplekey_SRA_accessions.csv) data), to investigate library and sequencing batch effects. All library preparations and sequencing were performed at the University of Illinois Roy J Carver Biotechnology Center, and libraries were sequenced on an Illumina HiSeq 40000, across three flow-cells. Reads were mapped to the annotated *Pipra filicauda* genome (GCA_003945595.1) using splice-aware mapper STAR v.2.7.5 (Dobin *et al.* 2013), and transcripts were counted against exons using featureCounts v.2.0.1 (Liao *et al.* 2014).

Three samples were removed from further analysis due to low library size and poor mapping rates (Figure S1.2). We used hierarchical clustering of samples run in 2016 and 2018 and principal component analysis (PCA) ordination to characterize batch differences between replicate samples using *PCAtools* in R v. 4.0.2 (Blighe and Lun 2020; R Core Team 2020). All replicates were each other’s closest match, and differed in a relatively small fraction of multidimensional PCA space. Thus, we concluded that samples ran in 2016 and 2018 were not sufficiently different, and thus RNAseq data from 2016 were included in the analyses for the current study (Figure S1.3). Across all tissues, genes with an average count of less than five were removed. Then we used hierarchical clustering of samples and principal component analysis (PCA) ordination to identify potential outlier samples. Potential outliers were defined by large distances between other tissues in PCA and clustering outside the tissue in hierarchical clustering.

For most downstream analyses, individual tissues were analyzed separately because a) we wanted to capture differential expression of genes in one tissue that might be lowly expressed in another tissue (causing high dispersion), and b) because some tissues were analyzed on a single sequencing run, and because sample sizes were not equal for each tissue, a model accounting for batch and tissue effects could not be used. For each tissue, therefore, a second round of filtering was applied, where 1) genes with an average count of less than five were removed, and 2) genes were removed if > 50% of the samples had zero reads for that gene. Finally, for samples identified as potential outliers during the filtering above, they were removed if they had an overall network scaled connectivity of < –2.5 in the sample adjacency matrix within an individual tissue. After this filtering process for individual tissues, only a single sample (PFT2) from POM was removed, as it was consistently an outlier (Figure S1.4). After excluding samples during the filtering process described above, a total of 170 unique samples were included in analyses.

All samples analyzed are associated with Bioproject PRJNA437157, and SRR12660169-198, SRR19521260-271, SRR19521432-575. The file that matches SRA numbers with our code and metadata available on the [GitHub](https://github.com/periperipatus/PIFI_brain_transcriptome/blob/main/data_unfiltered/Samplekey_SRA_accessions.csv).

##### Candidate Genes

We predicted that our variables of interest would show evidence of differential expression in neuroendocrine genes known to be associated with testosterone signaling and vertebrate social behavior (Soares *et al.* 2010; Díaz-Muñoz *et al.* 2014; Kingsbury and Wilson 2016; Potticary and Duckworth 2021). These candidate genes included those for steroidogenic enzymes and sex steroid receptors, as well as key neuropeptides and their receptors. A total of 18 candidate genes to selected for closer investigation for evidence of differential expression associated with mean T, social status, or cooperative behavior. These genes and rationale for their inclusion are described in Table S1.3. Note that we follow the nomenclatural recommendations of Kelly and Goodson (2014) and use oxytocin and vasopressin rather than mesotocin and vasotocin as they are commonly used in the bird literature. This is because these genes are orthologous across animals and differential nomenclature in within each group is confusing (Kelly and Goodson 2014; Theofanopoulou *et al.* 2021). Further details of this analysis is in Supplement 4.

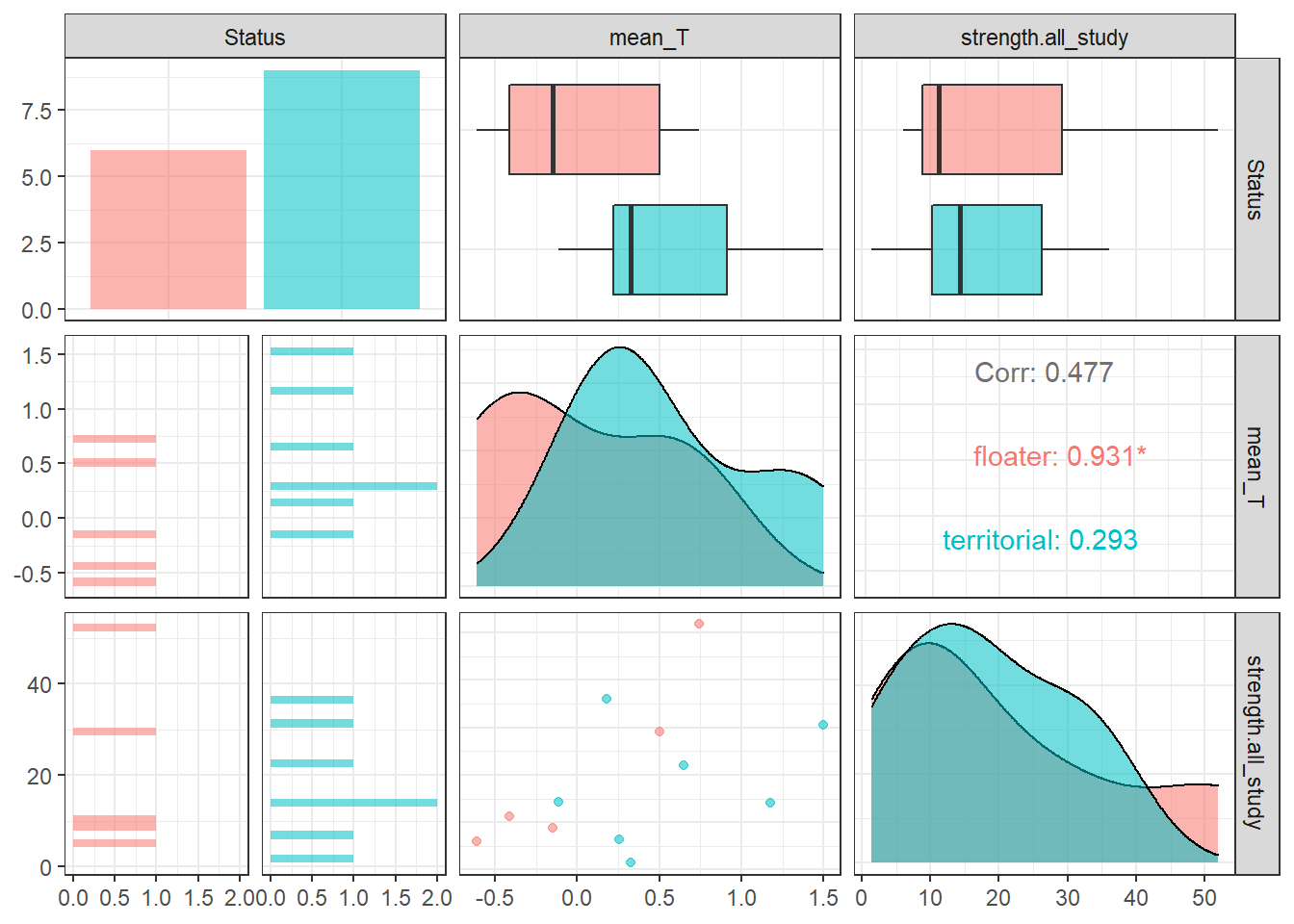

###### **Figure S1.1 Relationships between predictor variables**.

For individuals with repeated samples of mean testosterone phenotype (mean_T) and strength of cooperative behavior (strength.all_study, a social network measure). Linear regression found there was no statistically significant relationship between mean T and strength (p=0.12, correlation coefficient R=0.48).

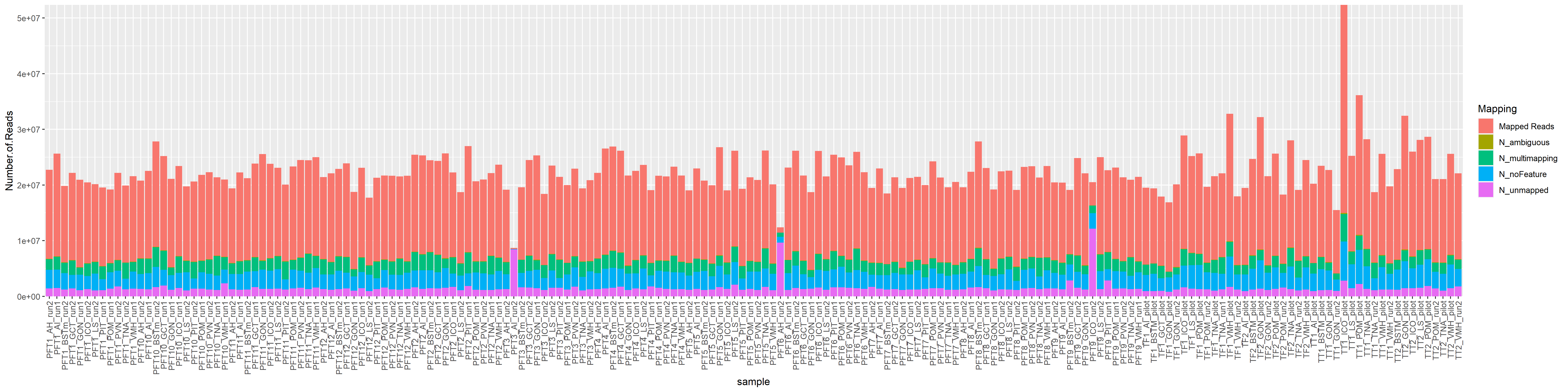

###### **Figure S1.2 Mapping summary for each library.**

Derived from STAR alignment output. Red denotes the number of mapped reads.

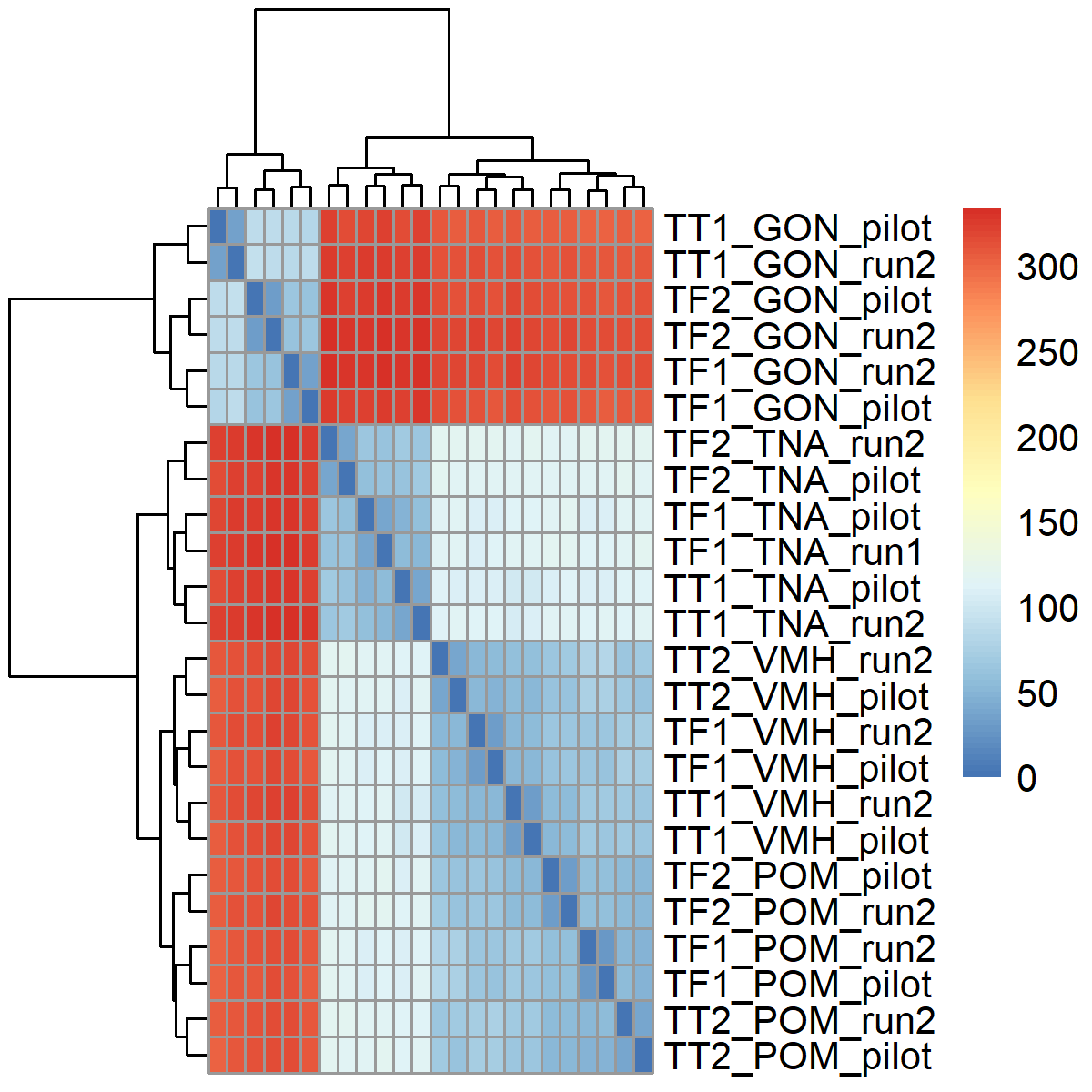

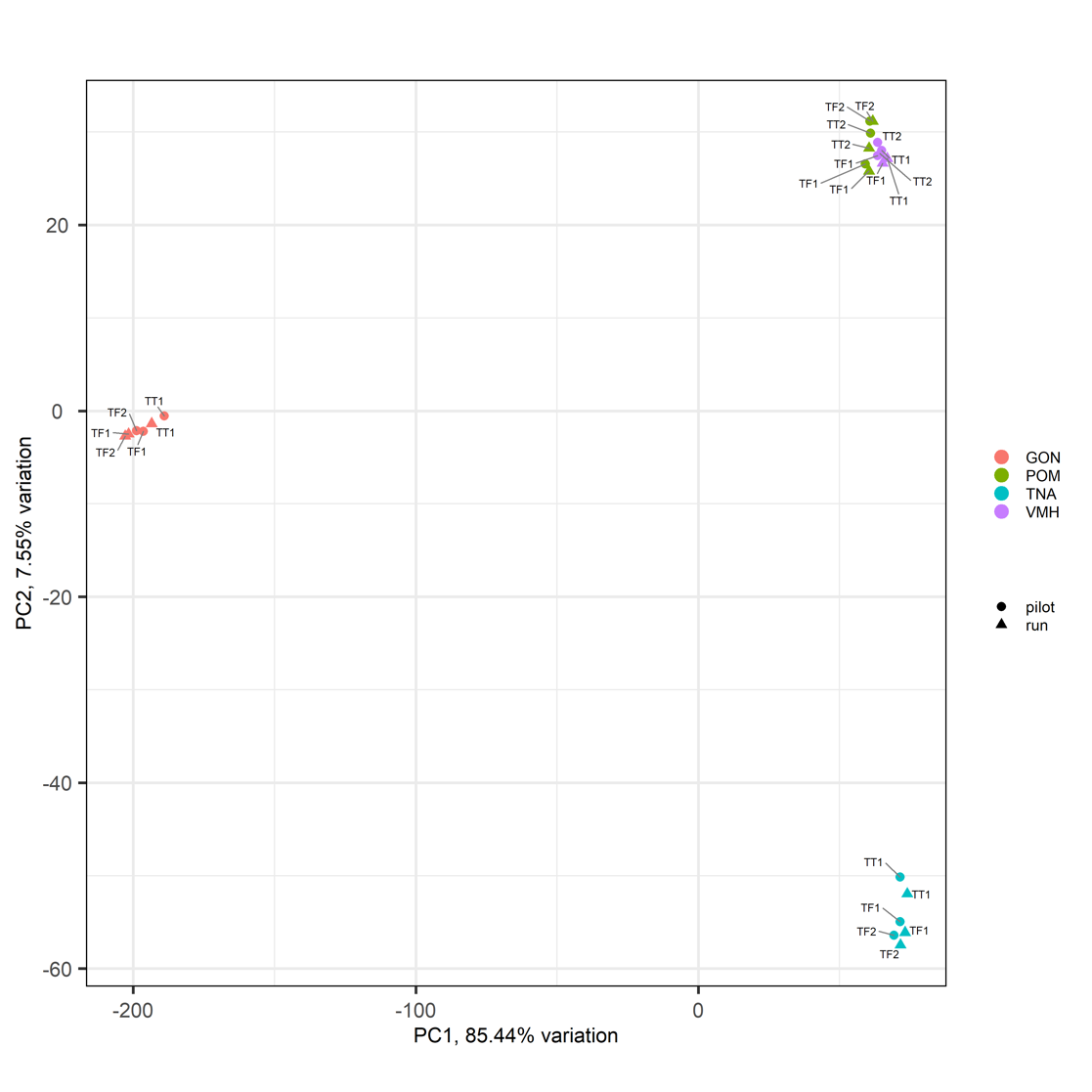

**A	B**

###### **Figure S1.3 Replicate samples visualization**.

A) PCA plot showing replicate samples for GON, POM, TNA, and VMH with individual’s sample ID annotated. B) Hierarchical clustering of between sample distances shows each replicate is each other’s most similar gene expression profile.

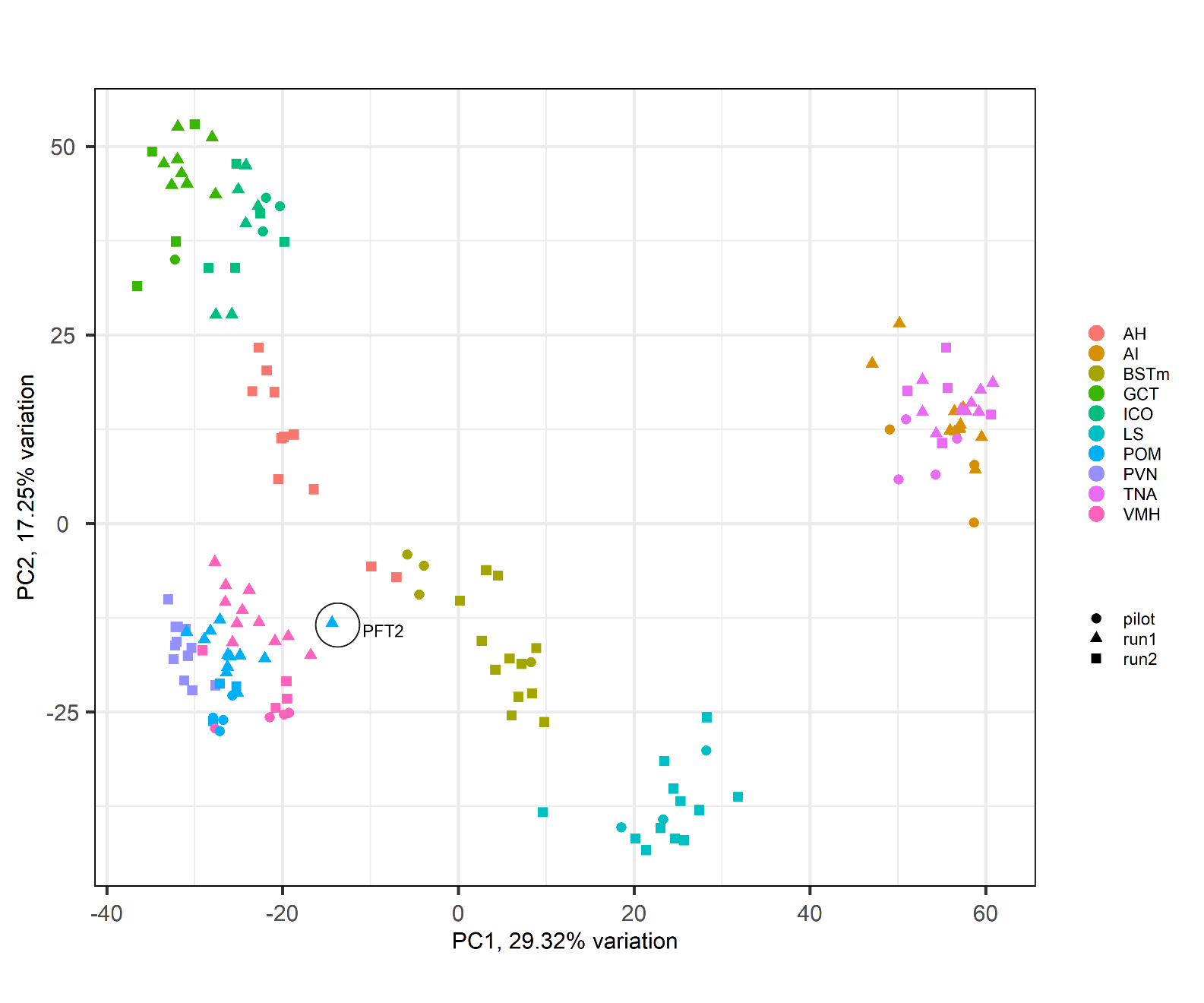

###### **Figure S1.4 Principal Component Analysis of brain gene expression profiles**

Color coded according to tissue, with point shapes representing sequencing batches. After individual tissue analysis for outliers, only PFT2 from the POM was consistently an outlier, and was subsequently removed from the analysis.

###### **Table S1.1 Individual birds sacrificed for this study**.

Including the number of tissues and brain regions used in the final study. Full sample details included in supplementary data.

| **Harvest ID** | **Tissues** | **Batches** | **Plumage and Status** | **Telemetry data?** | **Date of Sacrifice** | **Range of Raw Read Count** | **Comments** |
| --- | --- | --- | --- | --- | --- | --- | --- |
| PFT1 | 12 | 1 & 2 | Definitive territorial | Yes | 7-Jan-17 | 19207955 - 25637079 |  |
| PFT10 | 12 | 1 & 2 | Definitive territorial | Yes | 9-Jan-18 | 19754429 - 27805752 |  |
| PFT11 | 12 | 1 & 2 | Pre-definitive floater | Yes | 13-Jan-18 | 19396967 - 25574346 |  |
| PFT12 | 12 | 1 & 2 | Definitive territorial | Yes | 17-Jan-18 | 17731541 - 23884288 |  |
| PFT2 | 11 | 1 & 2 | Definitive territorial | Yes | 8-Jan-17 | 18701708 - 26975959 | Excluded from POM as consistently an outlier |
| PFT3 | 11* | 1 & 2 | Definitive floater | Yes | 8-Jan-17 | 8689632 - 25299167 | Excluded from PIT Strength DGE analysis owing to super high leverage; AI excluded due to low read count low mapping rate |
| PFT4 | 12 | 1 & 2 | Definitive territorial | Yes | 12-Jan-17 | 19085287 - 26926927 |  |
| PFT5 | 12 | 1 & 2 | Definitive floater | Yes | 12-Jan-17 | 19030162-26788439 |  |
| PFT6 | 11 | 1 & 2 | Definitive floater | Yes | 14-Jan-17 | 12401000 - 26148177 | Excluded AH due to low read count and poor mapping |
| PFT7 | 12 | 1 & 2 | Definitive territorial | Yes | 16-Jan-17 | 18480724 - 24247559 |  |
| PFT8 | 12 | 1 & 2 | Definitive territorial | Yes | 6-Jan-18 | 19106958 - 27808709 |  |
| PFT9 | 11 | 1 & 2 | Pre-definitive floater | Yes | 9-Jan-18 | 19097739 - 25000658 | Excluded ICO due to poor mapping |
| TF1 | 9 | pilot | Pre-definitive floater | No | 20-Dec-15 | 16905547 - 32806460 | Data published in Horton et al 2020 GBB https://doi.org/10.1111/gbb.12560; technical replicates in new batches |
| TF2 | 9 | pilot | Pre-definitive floater | No | 22-Dec-15 | 18289603 - 32213367 | Data published in Horton et al 2020 GBB https://doi.org/10.1111/gbb.12561 |
| TT1 | 8 | pilot | Definitive territorial | No | 20-Dec-15 | 15485248 - 52344603 | Data published in Horton et al 2020 GBB https://doi.org/10.1111/gbb.12560; technical replicates in batch 2 |
| TT2 | 9 | pilot | Definitive territorial | No | 22-Dec-15 | 21055879 - 32455212 | Data published in Horton et al 2020 GBB https://doi.org/10.1111/gbb.12561 |

###### **Table S1.2 List of tissues included in the study**

Including tissue codes used throughout this paper. Samples were sequenced over three batches, including the original pilot published in Horton *et al.* (2020). Numbers in brackets indicate number of samples used in the final analysis.

| **Name** | **Code** | **Pilot** | **Batch 1** | **Batch 2** | **Territorial** | **Floater** | **Telemetry** | **Comment** |
| --- | --- | --- | --- | --- | --- | --- | --- | --- |
| Anterior Hypothalamus | AH | - | - | 12(11) | 7 | 5(4) | 7 | Excluded PFT6 due to low counts and mapping |
| Arcopallium Intermedium | AI | 3 | 11 | 1(0) | 8 | 7(6) | 12(11) | Excluded PFT3 from run 2 due to low reads and mapping |
| Bed Nucleus of the Stria Terminalis | BSTm | 4 |  | 12 | 9 | 7 | 12 |  |
| Midbrain Central Grey | GCt | 1 | 8 | 4 | 7 | 6 | 12 |  |
| Testes | GON | 4 | 12 | 3(0) | 9 | 7 | 12 | TT1 and TF1&2 were repeated in run 2, but only pilot were included in final analysis |
| Intercollicular Nucleus | ICo | 3 | 6 | 6(5) | 9 | 6(5) | 12(11) | PFT9 was excluded due to poor mapping |
| Lateral Septum | LS | 3 |  | 12 | 9 | 6 | 12 |  |
| Pituitary | PIT | - | 8 | 4 | 7 | 5 | 12 | PFT3 was excluded from cooperation analysis due to high leverage. |
| Medial Preoptic Area | POM | 4 | 11(10) | 3(0) | 9(8) | 7 | 12(11) | TT1 and TF1&2 were repeated in run 2, but only pilot were included in final analysis; & PFT2 was excluded from all analysis due to an outlier gene expression profile |
| Paraventricular Nucleus | PVN | - | - | 12 | 7 | 5 | 12 |  |
| Nucleus Taenia | TnA | 4 | 10(9) | 5(3) | 9 | 7 | 12 |  |
| Ventromedial Hypothalamus | VMH | 4 | 11 | 4(1) | 9 | 7 | 12 | TT1&2 and TF1 were repeated in batch 2 |

###### **Table S1.3 List of candidate sex steroid related genes and neuropeptides**

These were included in this study for special focus. We include the NCBI gene accession numbers for the *Pipra* V1 genome used here.

| **Gene** | **V1 NCBI ID** | **HGNC Gene Name** | **Rationale** | **Reference** |
| --- | --- | --- | --- | --- |
| Androgen Receptor | 113993078 | AR | Status-specific regulation of behavior by testosterone. AR binds to testosterone and its derivative DHT. | (Schuppe *et al.* 2020; Ryder *et al.* 2020) |
| 5-alpha-reductase | 113992458 | SRD5A2 | Converts testosterone to dihydro-testosterone. |  |
| Aromatase | 113993669 | CYP19A1 | Converts testosterone to estradiol, also an important regulator of male social behavior. | (Schlinger and Balthazart 2013; Ubuka and Tsutsui 2014) |
| Estrogen receptor α | 113994625 | ESR1 | Receptor for estrogens, and regulator of social behavior. | (Horton *et al.* 2014; Maney *et al.* 2015) |
| Estrogen receptor β | 113985669 | ESR2 | Receptor for estrogens, not well characterized in birds, but ESR β has been linked to behavior in mammals | (Ubuka and Tsutsui 2014; Zuloaga *et al.* 2020) |
| Progesterone Receptor | 113997323 | PGR | Social behavior regulator | (Soares *et al.* 2010) |
| Gonadotropin releasing hormone | 113986838 | GNRH1 | Master regulator of gonadal steroid production (via direct modulation of gonadotropes). |  |
| Prolactin | 114002390 | PRL | Involved in regulation of care-giving behavior, including cooperative breeders and males. | (Potticary and Duckworth 2021) |
| Prolactin Receptor | 114000922 | PRLR | Receptor, as above |  |
| Vasoactive intestinal peptide | 113994612 | VIP | Prolactin-release and its expression across the brain is associated with social behaviors | (Kingsbury and Wilson 2016) |
| VIP receptor 1 | 113996142 | VIPR1 | Receptor, see above. |  |
| VIP receptor 2 | 113987732 | VIPR2 | Receptor, see above |  |
| Oxytocin | 113983511 | OXT | Oxytocin is a positive regulator of affiliative behavior | (Soares *et al.* 2010) |
| Oxytocin receptor | 114001344 | OXTR | Receptor, see above |  |
| Vasopressin | 113983498 | AVP | Arginine-vasopressin (Vasotocin in birds) is a negative regulator of affiliative behavior | (Soares *et al.* 2010) |
| Vasopressin receptor 1A | 113987995 | AVPR1A | Receptor, see above |  |
| Vasopressin receptor 1B | 113999504 | AVPR1B | Receptor, see above |  |
| Vasopressin receptor 2 | 113982601 | AVPR2 | Receptor, see above |  |

### 2. System-Wide Analyses: WGCNA and PCA

#### Introduction

This study is the most detailed assessment of the transcriptome of multiple nuclei in a wild animal brain. Previously, we assessed seven brain regions using RNAseq of the male wire-tailed manakins using two territorial and two floater individuals (Horton *et al.* 2020). We were interested in how the addition of three brain regions in this study influenced the tissue-specific patterns of co-expression seen previously (Horton *et al.* 2020).

#### Methods

This work follows a previous study of gene expression across the SBN in manakins (Horton *et al.* 2020), except that three additional brain nuclei were included in this study, including the midbrain central gray (GCt), paraventricular nucleus (PVN), and anterior hypothalamus (AH). To explore how overall patterns of gene expression were distributed among these larger suite of brain nuclei, we ran a principal component analysis (PCA) and weighted gene co-expression network analysis (WGCNA) (Langfelder and Horvath 2008).

##### Principal Component Analysis (PCA)

We ran two PCA analyses using *PCAtools* package in R v. 4.0.2 (Blighe and Lun 2020; R Core Team 2020). The first included the brain and HPG tissues of the pituitary (PIT) and testes (GON) (171 libraries), and the second included only the brain (143 libraries). All data were filtered to remove genes with an average count of less than 20 were removed, and genes with >50% of samples had zero reads. In the brain only subset we removed a sample from the POM (PFT2) because it was identified as an outlier in single tissue analyses (See section 1 & 3), but removing this individual had no appreciable effect on our results (not shown). All data were transformed for use in PCA using *DESeq2*’s getVarianceStabilizedData() function (Love *et al.* 2014). We then correlated batch, tissue type, and interest variables with our PC axes, to discern global patterns. When PC axes were associated with an interest variable and other variables, we used linear mixed models in the *nlme* R package to test for the effect of our interest variable on gene expression on that PC. The general model form was PC axis ~ (Batch) + (Year) + Tissue*Trait + (1|Individual), where Individual is the random effect.

##### Weighted Gene Co-expression Network Analysis (WGCNA)

To explore how replicable the co-expression patterns from the first *Pipra* brain study were, we ran WGCNA (Langfelder and Horvath 2008) excluding samples from Horton *et al.* (2020). We excluded a sample from the POM (PFT2) as the single tissue analyses identified them as an outlier. Then, for all 10 brain nuclei we filtered the data to exclude genes that had a mean count of <20 across 116 libraries. Data were transformed for use in WGCNA using *DESeq2*’s getVarianceStabilizedData() function (Love *et al.* 2014).

Using a soft-thresholding power (β) of 20, we built a signed adjacency matrix using Pearson correlations between genes. We used a minimum module size of 30 genes and used dynamic tree cutting and a module similarity threshold of 0.3 to discern the modules of gene co-expression. We then Pearson correlated the module eigengene with our 10 brain nuclei, bird ID, sequencing flow cell (Batch), and our interest variables (Status, Mean T, and Strength) to discern any whole-brain co-expression patterns associated with our traits. Hub genes were determined by module membership scores, which is the correlation between the individual gene’s expression pattern and the module eigengene (Langfelder and Horvath 2008). Gene expression patterns of interest were visualized using the normalized counts from *DESeq2* (Love *et al.* 2014).

#### Results & Discussion

##### All Tissues

PC1 and PC2 explained 57% and 14% of the variation in gene expression, respectively, for all tissues sampled. There were three distinct clusters representing all the brain nuclei, the pituitary, and the gonads (main text: Figure 1G). PC3 further explained 8% of the variation in gene expression and separated the brain nuclei from each other. PC axes 1,3,5 and 10 were significantly correlated with tissue of origin. Only in the main two axes, PC1 was positively associated with gene expression in the testes, and genes with positive loading were associated with glucose metabolism (HK3), germ cells (PKDREJ, BOLL), and flagellar motility (DRC7). Conversely, low values of PC1 were associated with brain tissues, and showed associations with genes involved in neural function (ENO2, SLC1A2). Low values of PC2 are associated with differentiating the Pituitary, and key pituitary transcripts are associated with this axis (e.g. prolactin (PRL), growth hormone (GH1), TSH subunit B (TSHB), propriomelanocortin (POMC)).

##### Brain: Social Decision Making Network (SDMN)

Even within the brain, the majority of gene expression variation was explained by differences between brain regions (Figure 1G inset, Figure S2.4) and was reflected in tissue-specific gene co-expression networks (Figure S2.7). For the brain only analyses, PC1 and PC2 explained 32% and 19% of the variation in gene expression (Figure S2.4A), while axis 3 explained 11% of the variation (Figure S2.4B). The correlation analysis suggested only 1,3,7,8 and 9 were significantly correlated with brain nucleus of origin.

PC5 was found to be significantly associated with social status (Figure S2.4C), even after using a mixed model to correct for the year confound, as well as individual and tissue effects (Figure S2.5). Interestingly, a candidate gene AVP was strongly associated with PC5 loadings (Figure S2.6), but did not yield an effect in any single tissue (Supplement 4). PC5 loadings also suggested global differential regulation of GRINA and GSTA3 and suggests the status classes differ in cholesterol processing and neural excitability across the brain.

The genes associated with axis loadings (Figure S2.6) were plausible in terms of our understanding of the heterogeneous distribution of these genes across the SBN. Many of the top genes in PC axes 1 and 2 were previously identified as associated with brain differentiation in this species, such as GAL, MC3R, SYT2, DDC, CBLN2, C1QL3, and PDYN (Horton *et al.* 2020). Unlike the previous work, however, we found OXT and AVP as associated with brain region differentiation.

We identified 22 modules of gene co-expression in the ten brain nuclei dataset (Figure S2.7). The expression patterns are more similar between adjacent nuclei, forming major groupings of the amygdala, hypothalamus, and midbrain. Module hub genes, whose expression is highly correlated with module eigengenes (kME) (Langfelder and Horvath 2008), broadly recapitulated previously documented patterns in the manakin brain with additional resolution provided by the enhanced sampling in the present study (Table S2.1) (Horton *et al.* 2020). All modules but ‘honeydew1’, ‘darkorange2’ and ‘grey’ (the module with “leftover” genes that don’t fit other co-expression modules) had at least one highly significant (p<0.009) correlation with a tissue (Figure S2.7). We visualized the top hub genes for modules associated with Status and Strength (‘darkturquoise’, ‘honeydew1’, ‘paleturquoise’ and ‘darkorange2’), but these did not provide strong relationships (in terms of slope and variance) when separated across tissues (shown on [GitHub](https://periperipatus.github.io/PIFI_brain_transcriptome/03_WGCNA_wholebrain.html)). As such, these results are not discussed further.

At both the broad scale and candidate gene level, we found similar patterns of gene expression as previous research using a subset of these same individuals (Horton *et al.* 2020). We found a similar signal of different co-expression signatures between hypothalamic (VMH, AH, PVN, POM), and the midbrain (GCt, ICo), and the forebrain.

Like previous findings, we found modules that were strongly positively associated with thalamic nuclei (‘navajowhite2’. ‘yellow’, and ‘tan’) were strongly negatively correlated with amygdala nuclei (AI and TnA). For each module we also observed similar module assignment of the key neurohormones, neurotransmitter receptors, hormone receptors and enzymes as identified in Horton *et al.* (2020) (Table S2.1). The ‘navajowhite2’ and the ‘tan’ modules carried most of the steroid-related genes, as well as neurodevelopmental gene OTP, and were associated with increased expression in most hypothalamic nuclei. These modules share most genes with the ‘black’ and ‘blue’ modules associated with the POM and VMH in previous work. The ‘navajowhite2’ module contained 20 genes important in the SBN, plus three genes related to the growth hormone pathway. We even recover similarities in the strong association of some modules with the lateral septum (LS), and similar but non-significant trends in the nearby medial bed nucleus of the stria terminalis (BSTm). Finally, expression of the androgen receptor (AR) and estrogen receptor α (ESR1) are broadly similar to that previously described in the related golden-collared manakin (*Manacus vitellinus*) (Fusani *et al.* 2014), but quantitative variation of these and other candidate genes across species could provide mechanistic basis for differences in social structure across manakins (Pipridae) (Prum 1994).

There were co-expression modules that distinguished the tissues unique to this study from other related tissues.

The anterior hypothalamus (AH) differed from other hypothalamic nuclei in the direction and magnitude co-expression of ‘navajowhite2’, ‘yellow’ and ‘darkolivegreen’ modules. In particular, the ‘navajowhite2’ is full of steroid receptors but showed significant downregulation in the AH. The ‘yellow’ module is positively associated with all the hypothalamic nuclei, but is inversely associated with the AH, suggesting much lower expression in this brain region. The top three hub genes for the yellow module (CFAP54, CFAP221, DNAH9) are associated with cilia and flagella. This may indicate that the microdissections for the hypothalamic nuclei have captured distinct differences in primary cilium organelles, important in hypothalamic regulation of homeostasis (Yang *et al.* 2021). The yellow module also describes some differences in neurotransmitter sensitivity in the AH, as serotonin receptor 2B (HTR2B) is a hub gene that is lowly expressed in the AH, and very highly in the PVN, relative to the other hypothalamic nuclei. Genes in the ’darkolivegreen’ module appear to be uniquely upregulated in the AH. This module included GO categories for protein synthesis and modification, as well as brain and eye development, though none of these were significant after FDR correction. Conversely, the PVN was more like other hypothalamic nuclei, but differed in its much stronger signal of upregulation of genes in the ‘yellow’ module. This module contains statistically significant enrichment for cilium-related GO categories and contains serotonin receptor 2B (HTR2B). There is subtle variation in module expression across the remaining hypothalamic nuclei but varies prominently among candidate genes. For example, POM has high expression of GNRH1 (Figure S4.1) and is a known site of *gnrh* synthesis.

The midbrain central grey (GCt) tissues differed from midbrain tissue intercollicular nucleus (ICo) primarily by strength of association with co-expression modules ‘black’, ‘bisque4’, ‘darkseagreen4’, ‘purple’, and ‘violet’. Compared with the nearby ICo, genes in the ‘black’ module mores strongly associated with increased expression in the GCt. This module includes candidate gene 5-alpha-reductase (SRD5A2) (Table S2.1). Genes in ‘bisque4’ were most strongly downregulated in the GCt, and included GO processes for RNA splicing, cilium assembly, synapse assembly, and neural cell-adhesion. This module also included notable downregulation for important neural receptors such as NPY1R and VIPR2 in the GCt (Table S2.1). Finally, the ‘tan’ module showed stronger upregulation in the GCt compared with the ICo. This module contains tyrosine hydroxylase and vasoactive polypeptide, and the GCt in birds shows a concentration of VIP-ir and TH-ir neurons (Kingsbury *et al.* 2011).

In a recent work, the TnA was revised to be included within the zebra finch arcopallium (renaming it to AMV) (Mello *et al.* 2019), and our data gives us an opportunity to explore genes differentiating the TnA and the AI in a suboscine bird. Both the previous study and ours found differences in the co-expression modules associated with each nucleus (Horton *et al.* 2020). In our study, these regions are united by similar expression of ‘tan’, ‘yellowgreen’, and ‘sienna3’ modules. The ‘sienna3’ module is similar in composition to the previous study, and both these regions show strong upregulation of key neuroendocrine genes including dopamine receptor D5 (DRD5) and cannabinoid receptor 1 (CNR1) (Horton *et al.* 2020). This module also contained C1QL3 (module membership=0.85) and ETV1 (module membership=0.85) (Figure S2.8A,B), which *in situ* studies previously demonstrated as important markers unique to the zebra finch arcopallium (Mello *et al.* 2019). The two regions are differentiated by upregulation of 488 genes in the ‘plum1’ module in the TnA, and upregulation of 333 genes in the ‘mediumpurple3’ module in the AI. Some known markers supported the TnA/AMV designation: KCTD12 was in the unique ‘plum1’ module (mm=0.81), along with MGP (mm=0.66), and HTR1D was a ‘hub’ gene (mm=0.93) despite no record of its association with this region in the zebra finch (Mello *et al.* 2019). We found mixed results for markers for the AI: HTR2A showed poor module membership in the ‘yellowgreen’ module (mm=0.42), but visualization showed strong expression in the AI (and LS) (Figure S2.8C), PCP4 was associated with the uniquely AI module medium purple (mm=0.79) (Figure S2.8D), while other Ai markers SCUBE1 and PVALB showed higher expression in the TnA (Figure S2.8E,F).

The PCA analyses and brain-wide co-expression analysis revealed biologically realistic differentiation in gene expression across the sampled tissues of the wire-tailed manakin. Compared with *in situ* methods, we uncovered a trove of potential differences in molecular architecture in comparison with oscine passerines (Saldanha *et al.* 2000; Fusani *et al.* 2014; Eaton *et al.* 2018; Mello *et al.* 2019). It remains to be seen what elements of these differences in molecular architecture are owing to their unique courtship displays (Lindsay *et al.* 2015), or are shared across suboscines. We also find suggestions of system-wide differences in gene expression between territorial and floater birds which is discussed further in the main text, and supplement section 6.

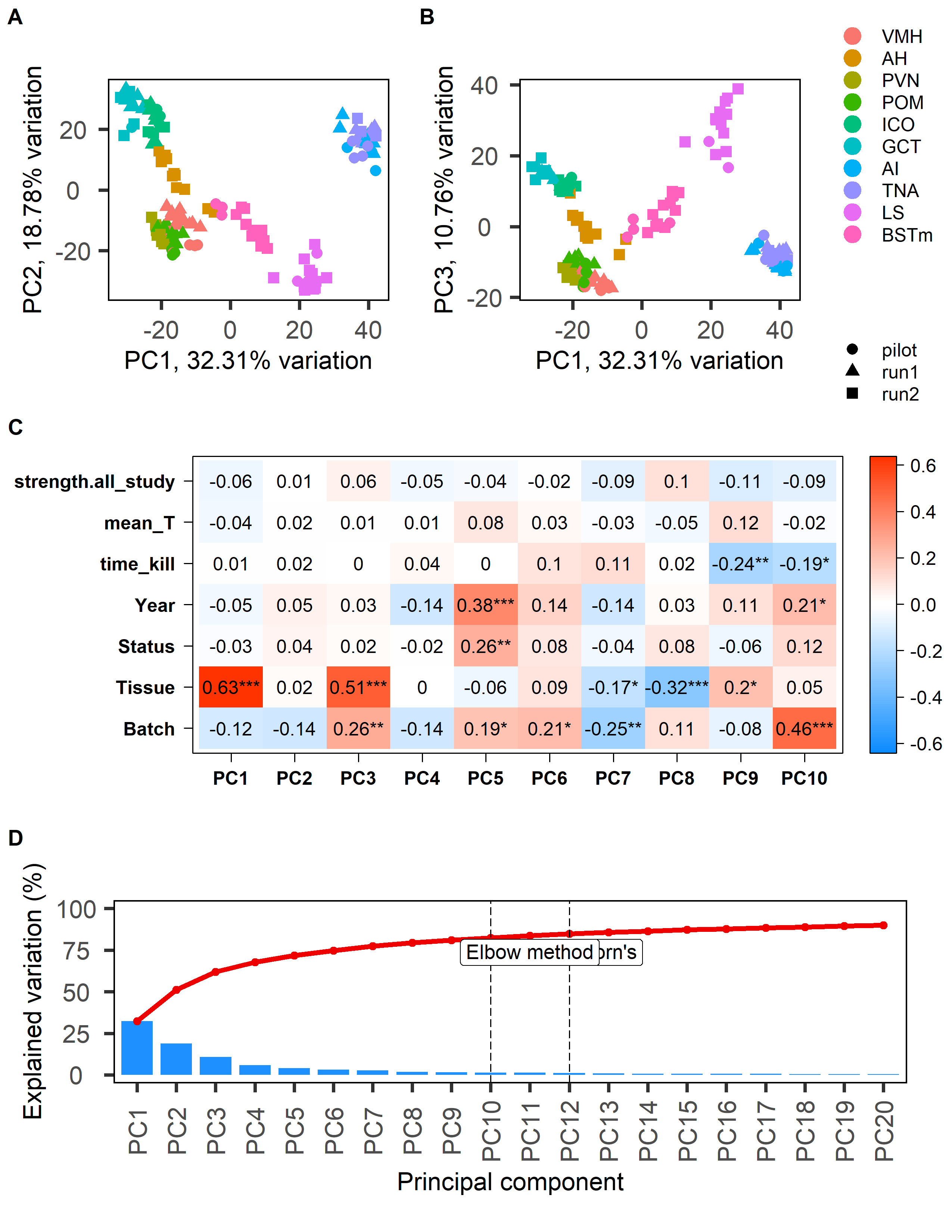

###### **Figure S2.1: PCA results for brain tissues only and relationship to predictor variables.**

Ordination plots for PC1 and PC2 (A) and PC1 and 3 (B), and the association of the top 10 eigenvectors with our interest and nuisance variables (C).

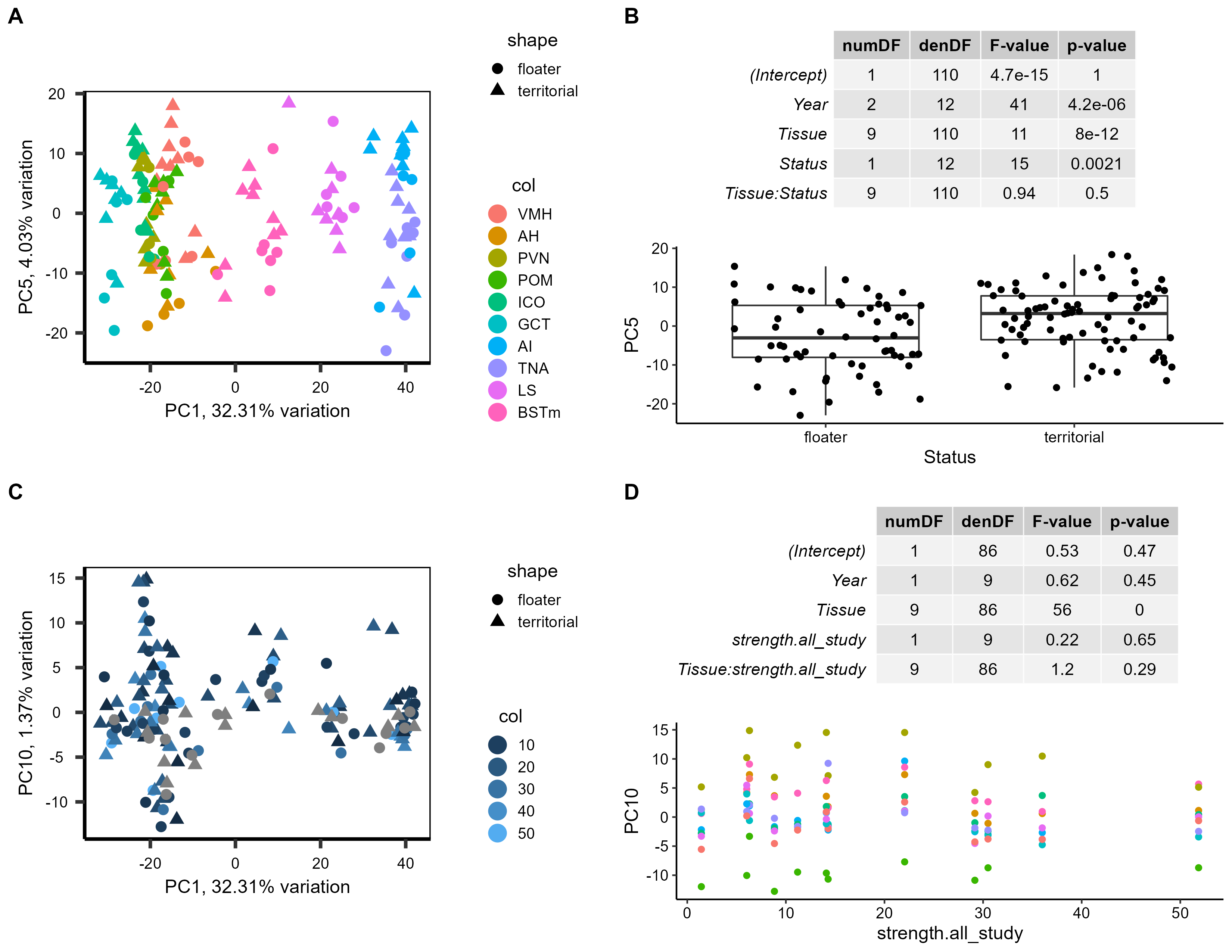

###### **Figure S2.2: Results from PCA and mixed model analysis on PC axes correlated with social status.**

A) Ordination plot of PC1 and PC5, the latter of which was significantly associated with social Status, B) Results from the ANOVA on the linear mixed model PC5~ Year + Tissue*Status + (1|Harvest_ID), and plot of PC5 against social Status.

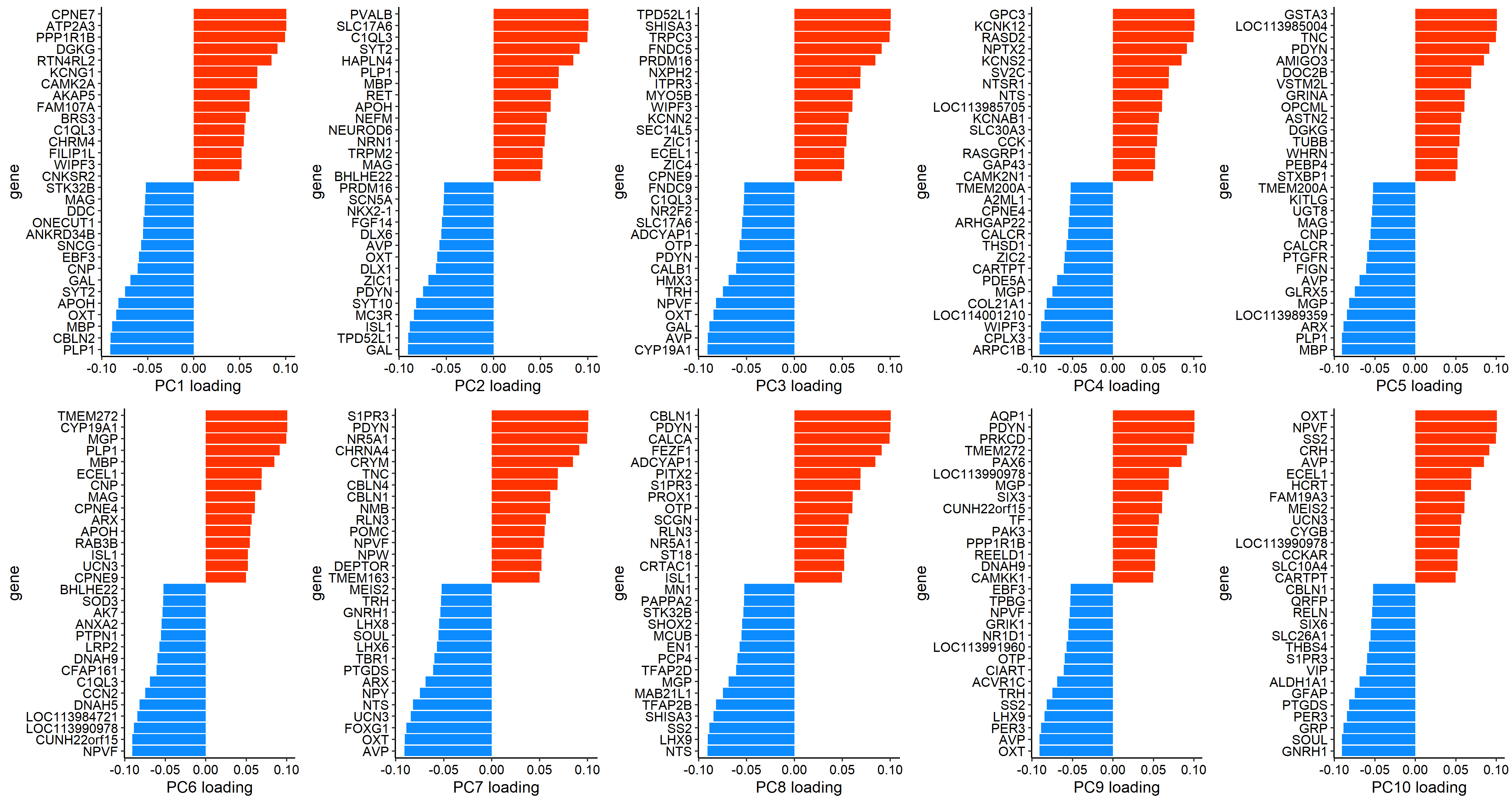

###### **Figure S2.3: Loadings for 10 PC axes for the analysis including only the brain nuclei**.

Where red bars indicate genes with positive associations with the PC axis, and blue indicates negative associations. Where possible genes with generic identifiers (LOC-) were replaced with the human gene name based on a BLAST search protocol (See section 3).

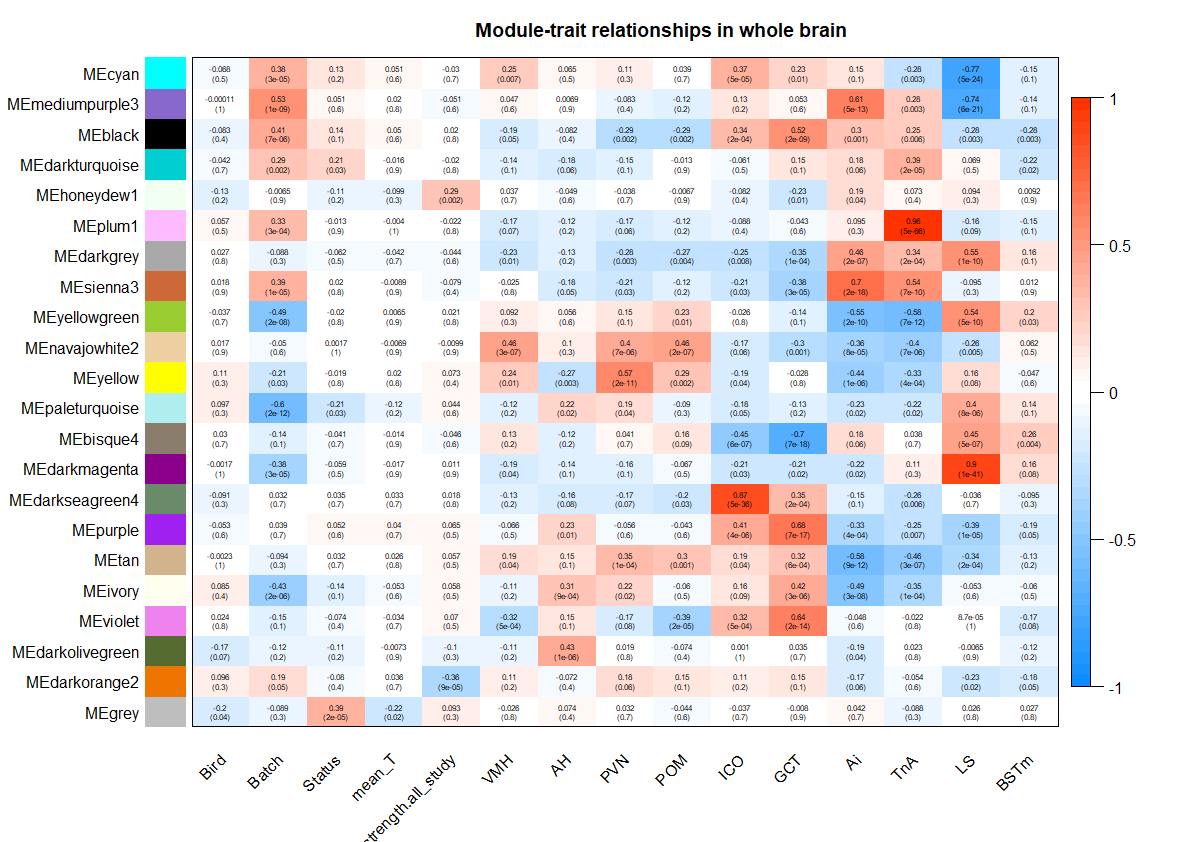

###### **Figure S2.4: Co-expression modules in the brain associated with each brain region.**

Correlation plot from WGCNA showing correlation coefficients and raw p-values between gene co-expression module eigengene and the individual bird ID, sequencing flow cell (Batch), our traits of interest (Status, mean T, and Strength), as well as the 10 brain nuclei of the SBN and MRS. Red squares indicate positive correlations, and blue is negative correlations. The Pearson correlation coefficient is indicated in the text of each square, and its p-value indicated in parentheses.

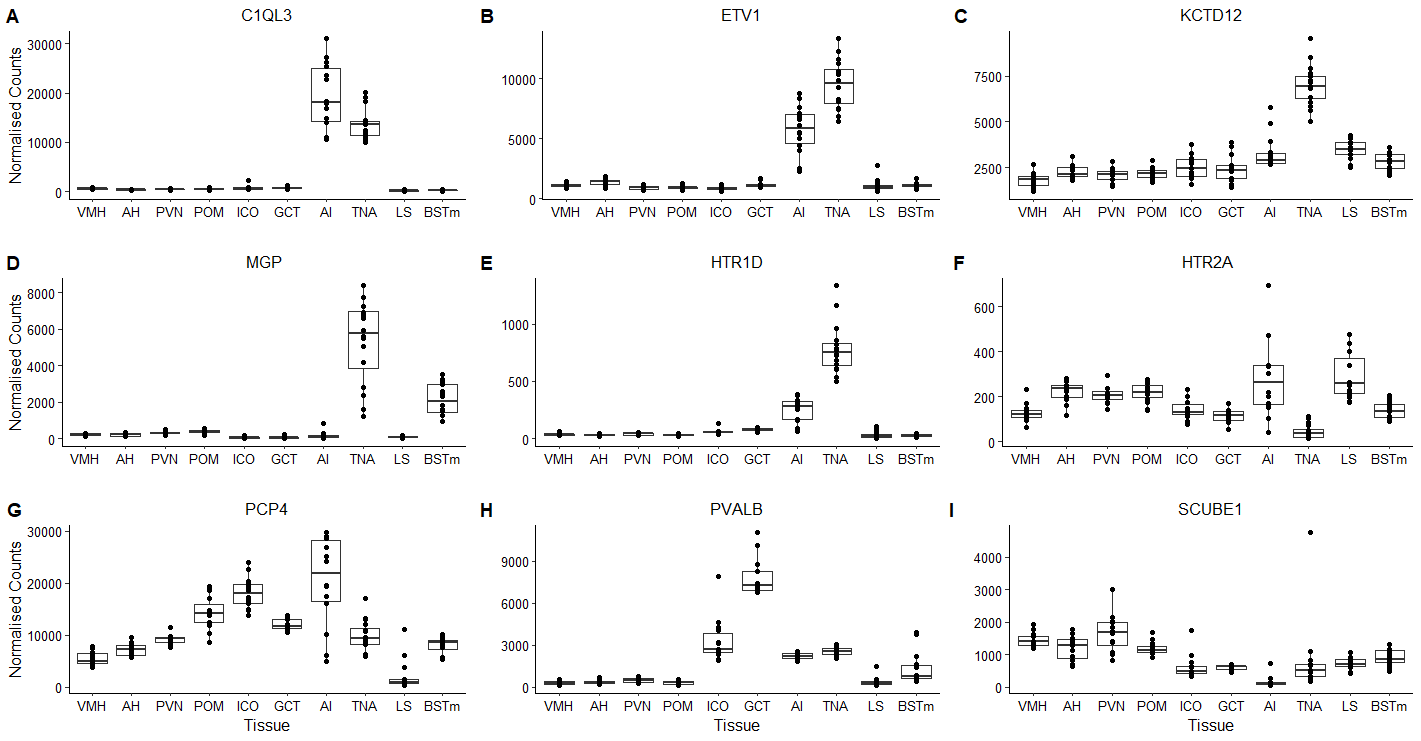

###### **Figure S2.5: Gene expression in Arcopallial markers.**

mRNA abundance across the SBN for genes previously identified to differentiate Arcopallial regions (AI, TnA) (Mello *et al.* 2019). Normalization method is the Median of Ratios from DESeq2 (Love *et al.* 2014).

###### **Table S2.1: Module assignment for key neurohormones, receptors, and enzymes**

These genes identified in the previous study of *Pipra filicauda* brains as important for SBN differentiation (Horton *et al.* 2020). Additional genes related to the growth hormone secretory pathway were included. Bolded genes are those in our candidate gene list we expected to be associated with our traits of interest.

| **Module** | **Gene** | **Name** | **Module Membership** |
| --- | --- | --- | --- |
| Navajowhite2 | MC3R | Melanocortin Receptor 3R | 0.96 |
|  | GAL | Galanin | 0.94 |
|  | **PGR** | Progesterone Receptor | 0.94 |
|  | GHSR | Growth Hormone Secretagogue Receptor | 0.93 |
|  | GHR | Growth Hormone Receptor | 0.87 |
|  | **ESR2** | Estrogen Receptor 2 | 0.86 |
|  | PDYN | Prodynorphin | 0.84 |
|  | PENK | Proenkephalin | 0.84 |
|  | **ESR1** | Estrogen Receptor 1 | 0.84 |
|  | TRHR | Thyrotropin-Releasing Hormone Receptor | 0.83 |
|  | TRH | Thyrotropin-Releasing Hormone | 0.83 |
|  | GHRH | Growth Hormone Releasing Hormone | 0.80 |
|  | **PRLH** | Prolactin releasing hormone | 0.79 |
|  | **LOC113983511 (OXT)** | Neurophysin-1-like (Oxytocin precursor) | 0.73 |
|  | **LOC113983498 (AVP)** | Vasotocin-Neurophysin-1 | 0.72 |
|  | **LOC113993669 (CYP19A1)** | Aromatase | 0.67 |
|  | **GNRH1** | Gonadotropin Releasing Hormone 1 | 0.62 |
|  | GALR3 | Galanin Receptor 3 | 0.62 |
|  | NPVF | Neuropeptide VF precursor | 0.60 |
|  | POMC | Pro-opiomelanocortin | 0.58 |
|  | **AR** | Androgen Receptor | 0.51 |
|  | NTSR1 | Neurotensin Receptor 1 | 0.50 |
|  | CALCB | Calcitonin-related Polypeptide Beta | 0.49 |
| Tan | DDC | DOPA decarboxylase | 0.85 |
|  | DRD2 | Dopamine Receptor 2 | 0.84 |
|  | TH | Tyrosine Hydroxylase | 0.83 |
|  | GNRH2 | Gonadotropin Releasing Hormone 2 | 0.68 |
|  | HTR1F | Serotonin Receptor 1F | 0.67 |
|  | **VIP** | Vasoactive Intestinal Polypeptide | 0.53 |
|  | HTR1A | Serotonin Receptor 1A | 0.45 |
|  | **OXTR** | Mesotocin Receptor | 0.40 |
| Yellowgreen | GALR1 | Galanin Receptor 1 | 0.78 |
|  | **VIPR1** | VIP Receptor 1 | 0.77 |
|  | **PRLR** | Prolactin Receptor | 0.76 |
|  | HTR2A | Serotonin Receptor 2A | 0.41 |
| cyan | GABRB2 | GABA Receptor Type A Receptor Subunit Beta2 | 0.58 |
|  | TAC1 | Tachykinin | 0.57 |
|  | SST | Somatostatin | 0.50 |
| Sienna3 | CNR1 | Cannabinoid Receptor 1 | 0.85 |
|  | DRD5 | Dopamine Receptor 5 | 0.80 |
|  | GABBR1 | GABA Receptor Type B Receptor Subunit 1 | 0.79 |
|  | GABBR2 | GABA Receptor Type B Subunit 2 | 0.79 |
|  | INSR | Insulin Receptor | 0.79 |
|  | SSTR4 | Somatostatin Receptor 4 | 0.75 |
|  | HTR2C | Serotonin Receptor 2C | 0.73 |
|  | GABRR3 | GABA Receptor Type A Receptor Subunit Rho3 | 0.62 |
|  | SSTR2 | Somatostatin Receptor 2 | 0.44 |
|  | SSTR1 | Somatostatin Receptor 1 | 0.32 |
| Bisque4 | GABRB1 | GABA Receptor Type A Receptor Subunit Beta1 | 0.92 |
|  | NPY1R | Neuropeptide Y Receptor 1 | 0.81 |
|  | NPY5R | Neuropeptide Y Receptor 5 | 0.76 |
|  | **VIPR2** | VIP Receptor 2 | 0.69 |
|  | NTS | Neurotensin | 0.56 |
|  | NPY | Neuropeptide Y | 0.52 |
| Darkgrey | DRD1 | Dopamine Receptor 1 | 0.93 |
|  | HTR6 | Serotonin Receptor 6 | 0.86 |
|  | HTR1B | Serotonin Receptor 1B | 0.65 |
|  | HTR7 | Serotonin Receptor 7 | 0.42 |
|  | DRD3 | Dopamine Receptor 3 | 0.38 |
| Black | HTRA2 | Serotonin Receptor A2 | 0.67 |
|  | **SRD5A2** | 5-Alpha-Reductase 2 | 0.53 |

### 3. Per Tissue Differential Expression Analysis

#### Methods

We examined each tissue individually for genes differentially expressed according to the traits of interest (status, mean testosterone, strength) using *DESeq2* (Love *et al.* 2014). After filtering (Supplement Section 1), each tissue was examined for statistically significant variation in expression according to confound variables of sequencing batch and capture year using correlation with principal components of gene expression using R package *PCAtools* (Blighe and Lun 2020). There were four additional samples for social status (pilot) samples, and some tissues were run over multiple batches, others on the same plate (see Table S1.1, S1.2). Owing to this complicated sampling, we used a decision tree framework when analyzing each tissue and variable of interest which is outlined in Figure S3.1.

*DESeq2* would give a convergence warning when raw social network strength data was used, and we scaled (centered around zero) those data based on the *DESeq2* recommendations. Following Klaus (2014), we corrected hill-shaped (skewed towards 1) p-value distributions using R package *fdrtool* v1.2.15 (Klaus and Strimmer 2020). This distribution often occurs when there is higher variance in the count data than is accounted for in the original *DESeq2* model (Klaus 2014), and *fdrtool* is a way of re-estimating the null variance and correcting the p-values (Klaus and Strimmer 2020).

Differentially expressed genes were identified using *DESeq2*’s negative binomial generalized linear models that included Batch and/or Year variables where necessary (model form: Gene Exp ~ (Batch) + (Year) + Trait[1:3]), with the full logic of variable inclusion described in Figure S3.1. Genes significantly differentially expressed were identified using the Wald Test and an FDR corrected p-value of 0.1 (q<0.1). The top six most significantly differentially expressed genes were plotted to validate relationship with predictor variables using *ggplot2* v3.3.2 (Wickham *et al.* 2020).

Finally, the individual with the highest strength value (PFT3) was associated with many genes with a highly influential outlier in the pituitary (Figure S3.2A). Although we used a customized function that leveraged Cook’s Distance to identify and discard genes with outlying samples for most tissues (see [GitHub](https://github.com/periperipatus/PIFI_brain_transcriptome/blob/analysis_code/02_DESeq2.Rmd) for code), this script was not sensitive enough to remove these genes. Because PFT3 was associated with a higher cook’s distance than other samples in this analysis (Figure S3.2B), it was removed, and the analysis repeated without it.

The results from these analyses were also used in the candidate gene analyses (Section 3). For the candidate gene analysis, we also explored whether these genes had status-specific gene expression by running an additional DESeq2 model: Gene Exp ~ (Batch) + (Year) + Status*mean_T.

##### Gene Ontology

There are 5817 genes in the *Pipra filicauda* V1 genome annotated as unknown (e.g. LOC113997056). To maximize the number of genes with Gene Ontology (GO) annotations we extracted the CDS from the *Pipra filicauda* genome, and conducted multiple reciprocal BLAST searches against human, manakins, and a selected set of well annotated birds and reptiles. The consensus gene symbol from these multiple BLAST results was assigned to the *Pipra* genes. These best symbols were then used to extract the human GO terms from geneontology.org GAF (version 2021-02-01). Using these annotations, we used R package *ClusterProfiler* v 3.16.1 (Yu *et al.* 2012) to conduct GO enrichment on genes identified as differentially expressed, or assigned to WGCNA modules (Section 5), and these are mostly discussed in the main manuscript. In cases where there were few significant differentially expressed genes (<40), we ran gene set enrichment analysis (GSEA) on the full gene list. This approach leverages the order of genes in the gene list (ranked on signed p-value) to describe the types of genes that show some relationship between expression and the explanatory variable. To help parse the information for each tissue, all GO terms were clustered and summarized using R package *rrvgo* v 1.0.2 (Sayols 2020), using a semantic similarity of ≥ 0.7.

##### Testis Size

To accompany data on gene expression differences in the testes, we compared the measured size of the testes from individuals included in this study. Both left and right testis were measured on extraction for both length and width using calipers (mm). We then calculated testis volume for each testis using the previously published formula to calculate testis size (Moore *et al.* 2002):

$$\frac{4}{3}\pi(0.5\times length)(0.5\times{width}^{2})$$

Then, we calculated the average testis size per individual. We then tested using ANOVA whether there was a difference between territorial and floater males in the average testis size, as well as left and right separately (n=16). Then, to explore the effects of testosterone on testis size we ran a linear model for testis size with interaction effect for social status and mean testosterone (n=12).

#### Results & Discussion

Here we present an abridged version of the differential expression analysis because it is long and repetitive. The full set of results are deposited as an Rmarkdown file [here](https://github.com/periperipatus/PIFI_brain_transcriptome/blob/analysis_code/02_DESeq2.Rmd).

Across the sampled tissues, we found a landscape of differential gene expression in association with social status (Figure S3.3), mean testosterone phenotype (Figure S3.4), and strength of cooperative behavior (social network strength) (Figure S3.5). We also explored whether our candidate gene set had status-specific gene expression by conducting a differential expression analysis with an interaction effect (Status x mean_T), which similarly showed variation in expression across tissues (Figure S3.6). Although we report the volcano plots here for completeness, we only used this method for extracting candidate genes with status-specific expression (Section 4), and do not interpret them further. There was some evidence of a correlation between the number of DEGs recovered by social status, and those recovered according to meant T (Figure S3.7)

While there were only two significant DEGs after FDR correction in the testes (Figure S3.8A) associated with social status, there were interesting GO categories with near-significant downregulation (Figure S3.8B). These terms were clustered by semantic similarity into sperm development (parent term: penetration of zona pellucida) and responses to steroid hormone (Figure S3.8C). A t-test showed there was no significant difference between size of left and right testes in the 16 sampled males (t=1.41, df=28.42, p=0.17). There was a tendency for territorial males to have larger testes than floaters (Figure 4.44 A,B). The linear model describing whether testis size is related to the interaction between status and mean testosterone, provided some evidence that (after correcting for testosterone) territorial males had larger testes (Table 3.2). This, paired with enrichment for genes relating to sperm performance and spermatogenesis by social status (Figure S3.8), and a large number of differentially expressed genes in males with higher testosterone phenotype (Figure S3.20) may suggest enhanced fertility for territorial males.

The remainder of the results for social status in each tissue are presented in figures S3.9-3.19. The results for each tissue in association with mean testosterone phenotype (mean T) are presented in figures S3.20-S3.31, and the results in association with social network strength are presented in figures S.32-44. Note that for the testes we do not expect a biologically meaningful relationship with social network strength, as this tissue is unlikely to be involved in the production of this behavior. However, we include it in the manuscript as all tissues were analyzed for all comparisons.

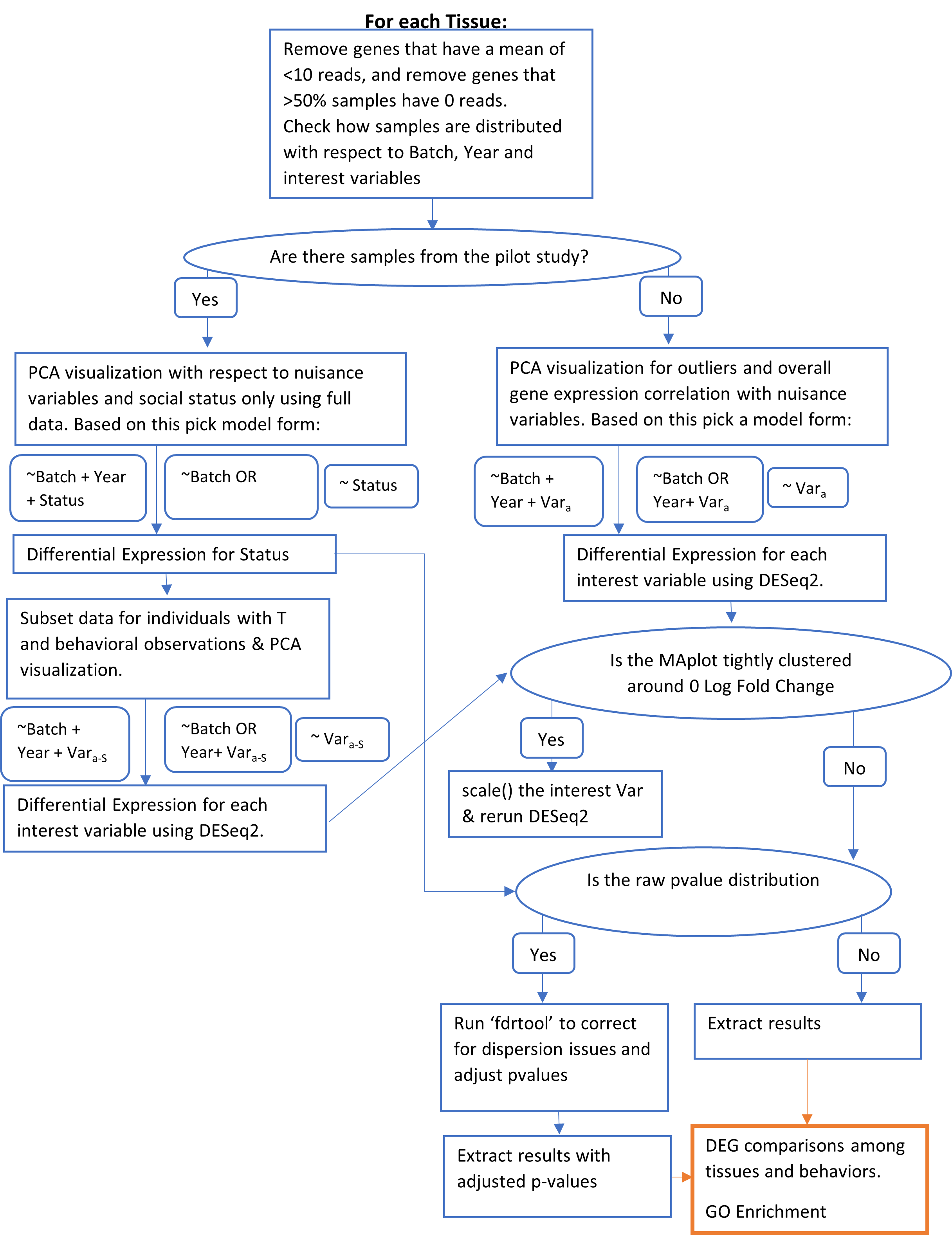

###### **Figure S3.1: Workflow for differential expression analysis in DESeq2.**

Explains decisions around whether batch variables were included in the DESeq2 model, and any other corrections applied to the data. The hill-shaped p-value distribution correction came was applied according to tutorial recommendations (Klaus 2014).

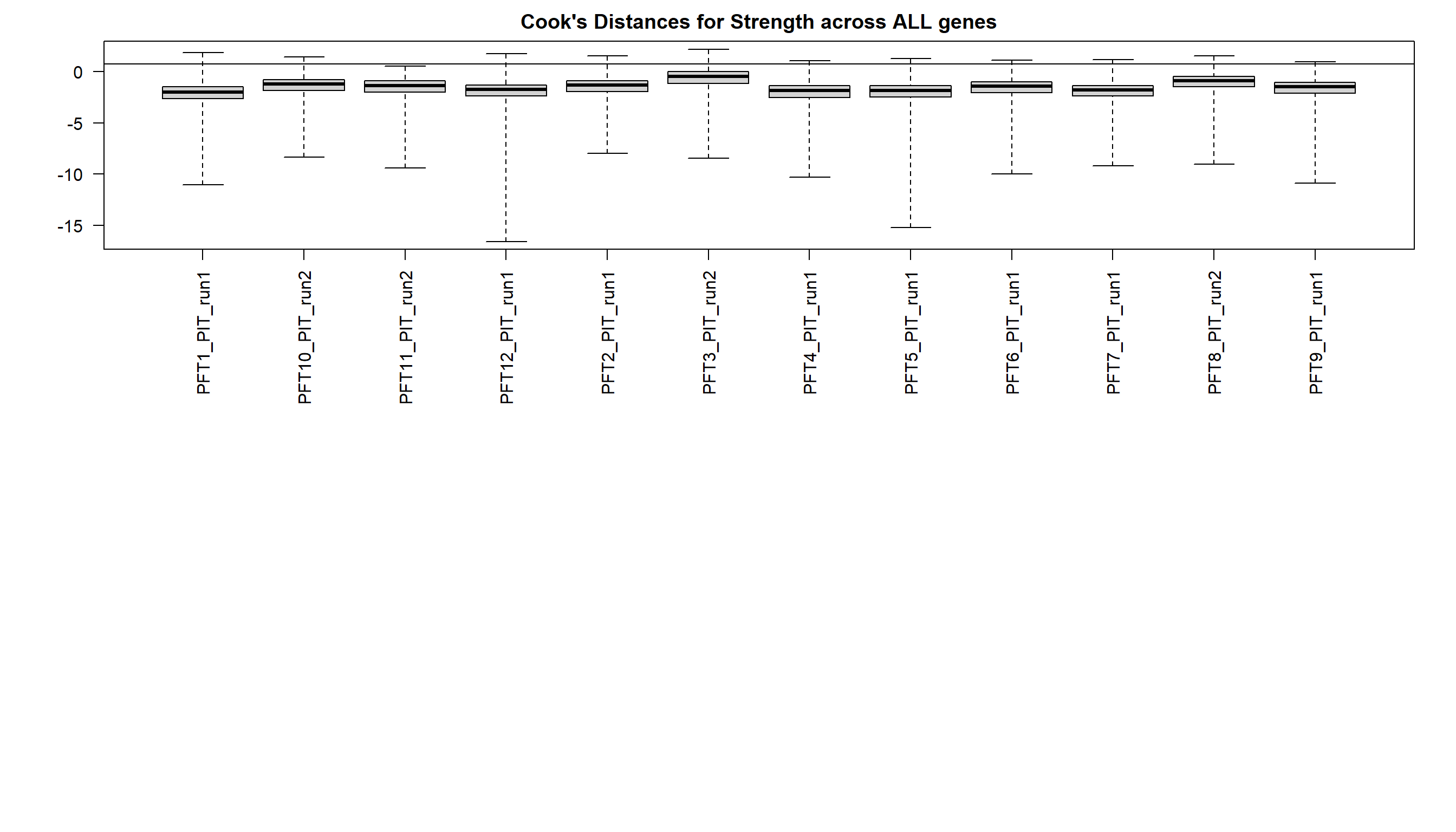

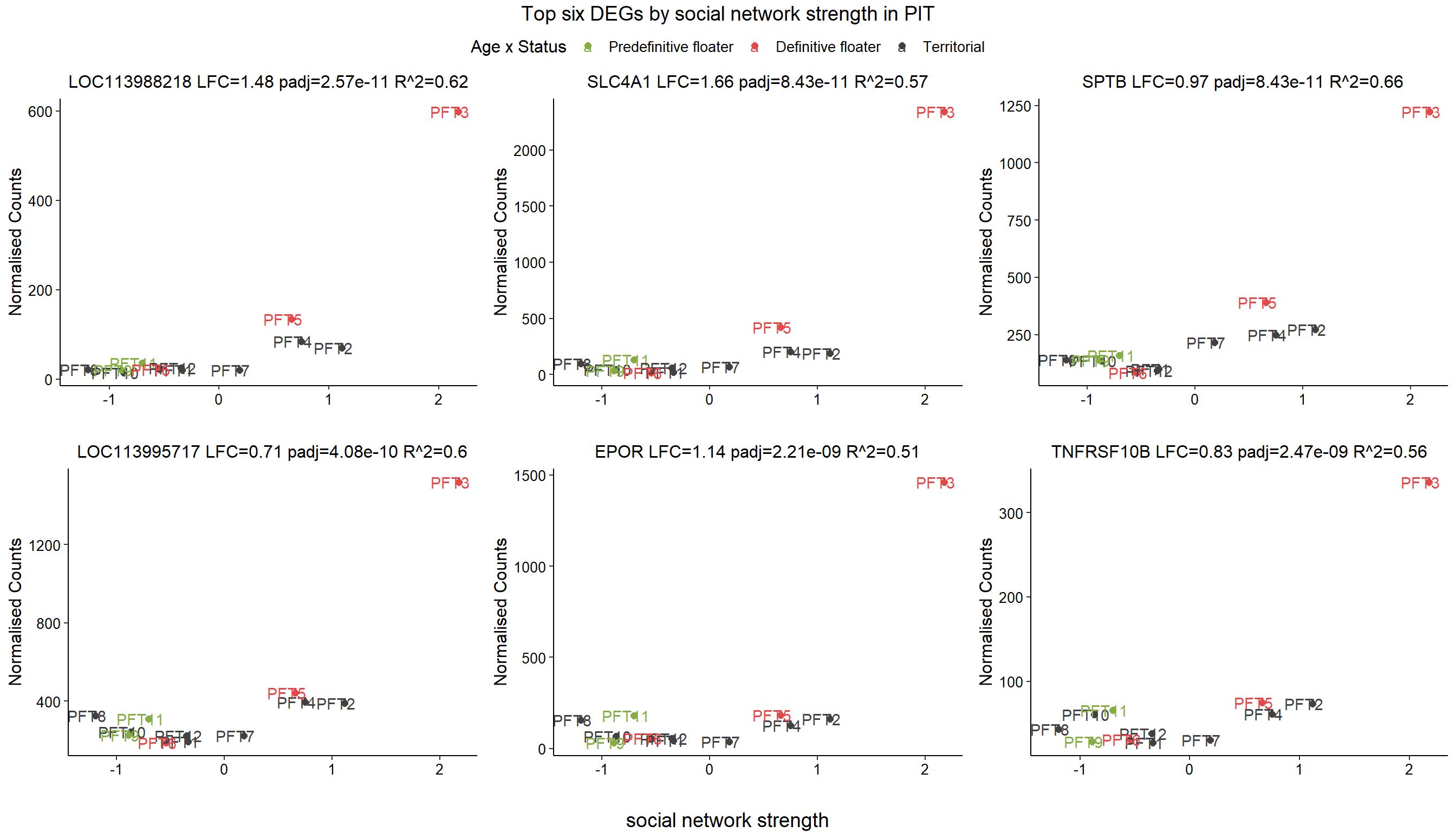

###### Figure S3.2: Outlier sample in DEG analysis of cooperation in pituitary.

A

B

Initial results from differential gene expression in the pituitary according to social network strength showing that PFT3 is an extreme outlier (A) in the top genes, as well as its overall leverage measured by Cook’s Distance (B). Thus, this sample was removed from the comparison with social network strength

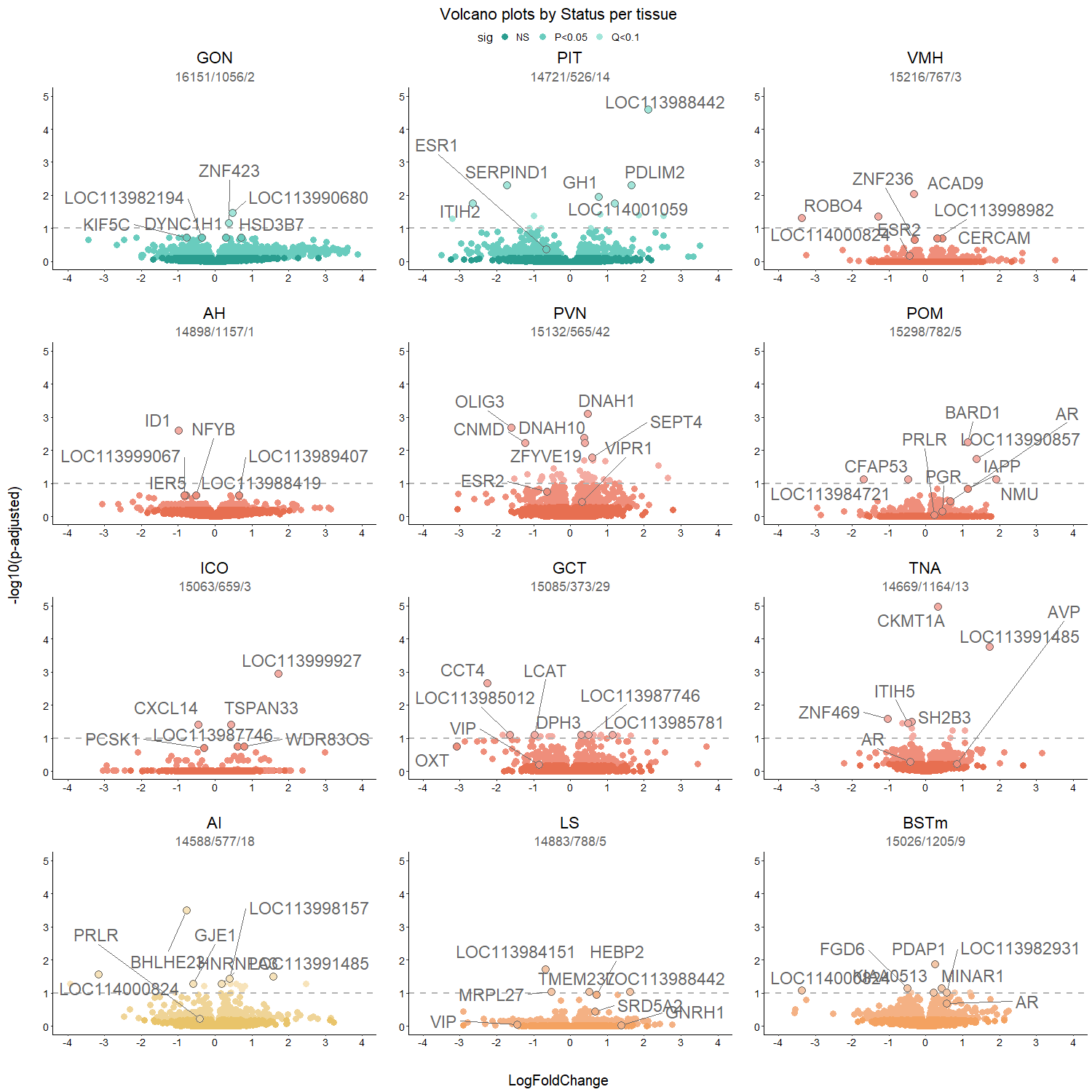

###### **Figure S3.3: Social status volcano plots**

From differential expression analysis in DESeq2 by social status, where the x-axis is log_2_ Fold-change, and positive values indicate higher expression in territorial individuals. The y-axis is log_10_ transformed p-values. Each tissue type is color coded according to functional networks described in Figure 1C. Results within each tissue are color coded by significance level, where genes that met the FDR q<0.1 threshold in the lightest color. The gene symbols for the top six most differentially expressed genes (based on p-value), and our candidate genes are highlighted if they had a raw p<0.05.

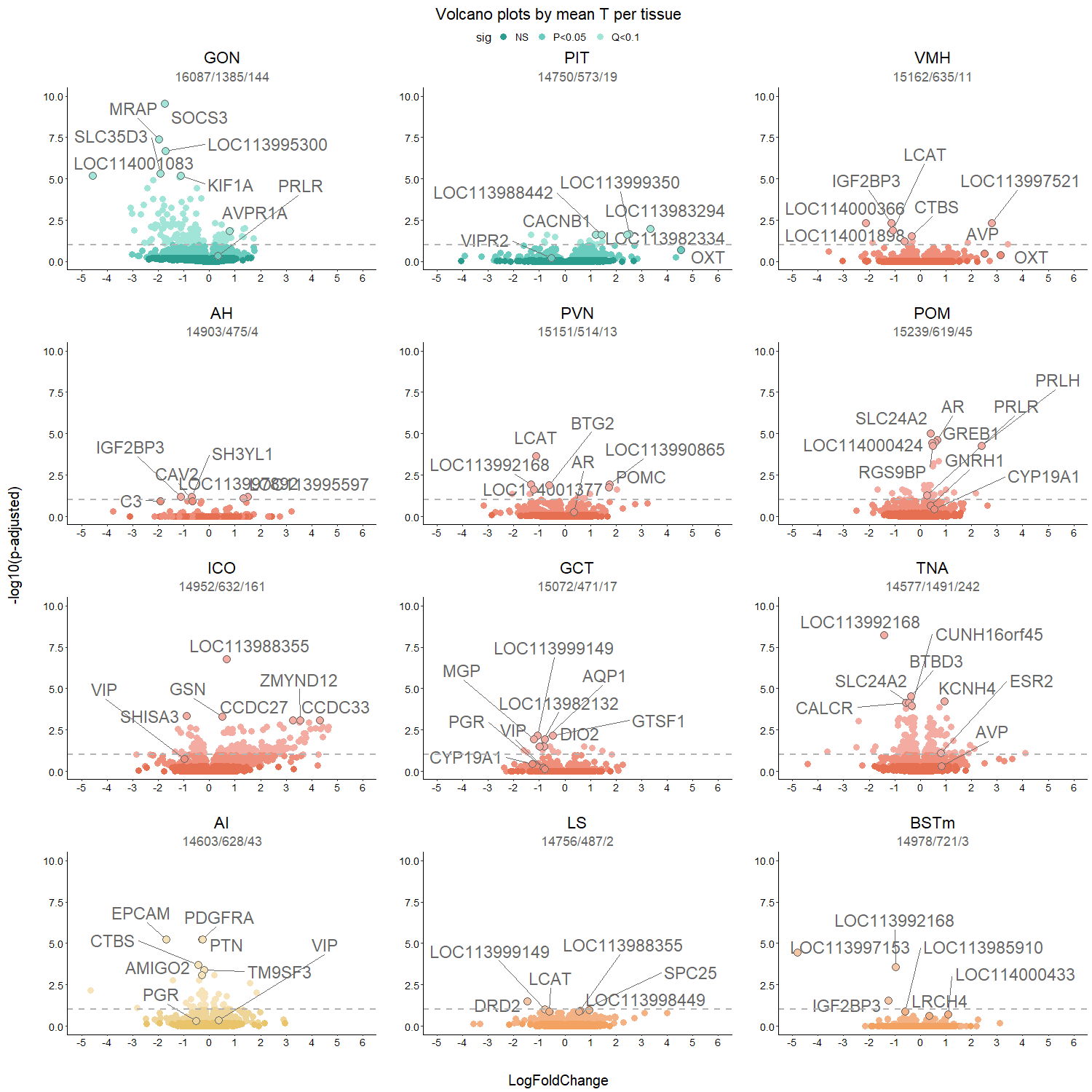

###### **Figure S3.4: Mean testosterone phenotype volcano plots.**

From differential expression analysis in DESeq2 by mean T phenotype, where the x-axis is log_2_ Fold-change, and positive values indicate higher expression in individuals with higher mean T. The y-axis is log_10_ transformed p-values. Each tissue type is color coded according to functional networks described in Figure 1C. Results within each tissue are color coded by significance level, where genes that met the FDR q<0.1 threshold in the lightest color. The gene symbols for the top six most differentially expressed genes (based on p-value), and our candidate genes are highlighted if they had a raw p<0.05.

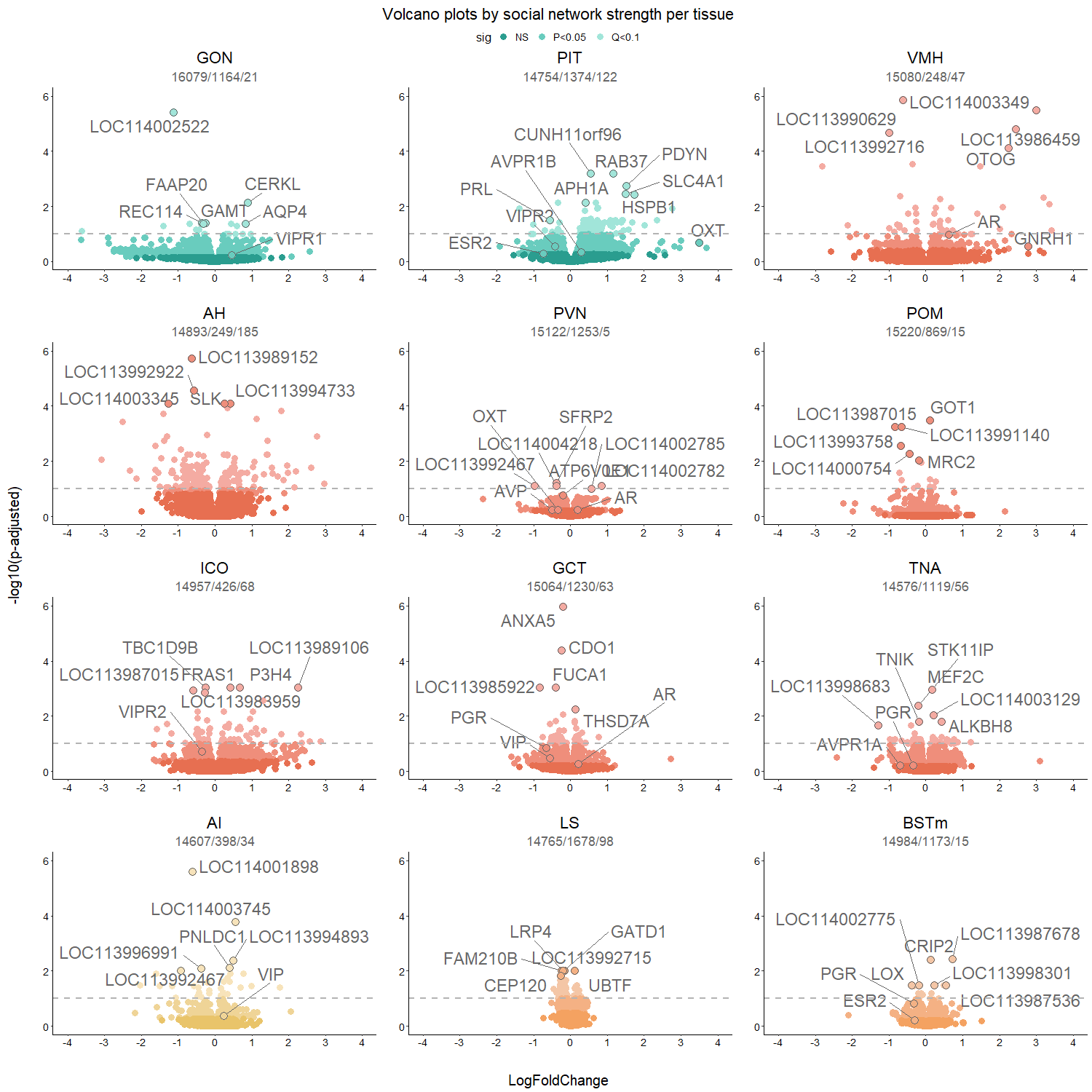

###### **Figure S3.5: Social network strength volcano plots.**

From differential expression analysis in DESeq2 by average social network strength of an individual over the study period, where the x-axis is log_2_ Fold-change, and positive values indicate higher expression in individuals with higher strengths of association. The y-axis is log_10_ transformed p-values. Each tissue type is color coded according to functional networks described in Figure 1C. Results within each tissue are color coded by significance level, where genes that met the FDR q<0.1 threshold in the lightest color. The gene symbols for the top six most differentially expressed genes (based on p-value), and our candidate genes are highlighted if they had a raw p<0.05.

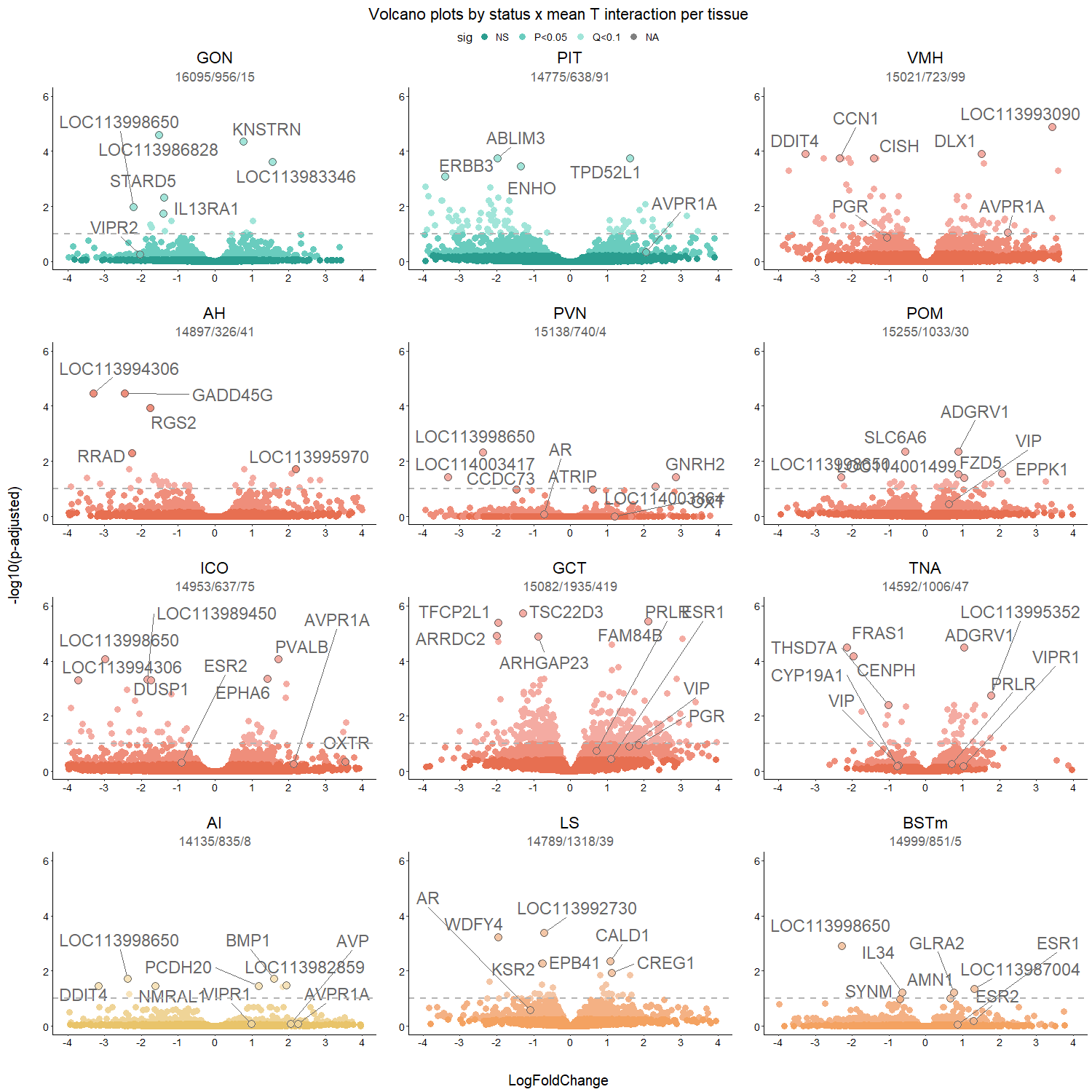

###### **Figure S3.6: Status x mean T interaction volcano plots.**

Differential expression analysis in DESeq2 of status-specific gene expression (Status x mean_T). The explanations are as in previous figures, except that the log fold change represents the difference between the specific effect of mean T on territorial gene expression vs the effect in floater individuals (reference level).

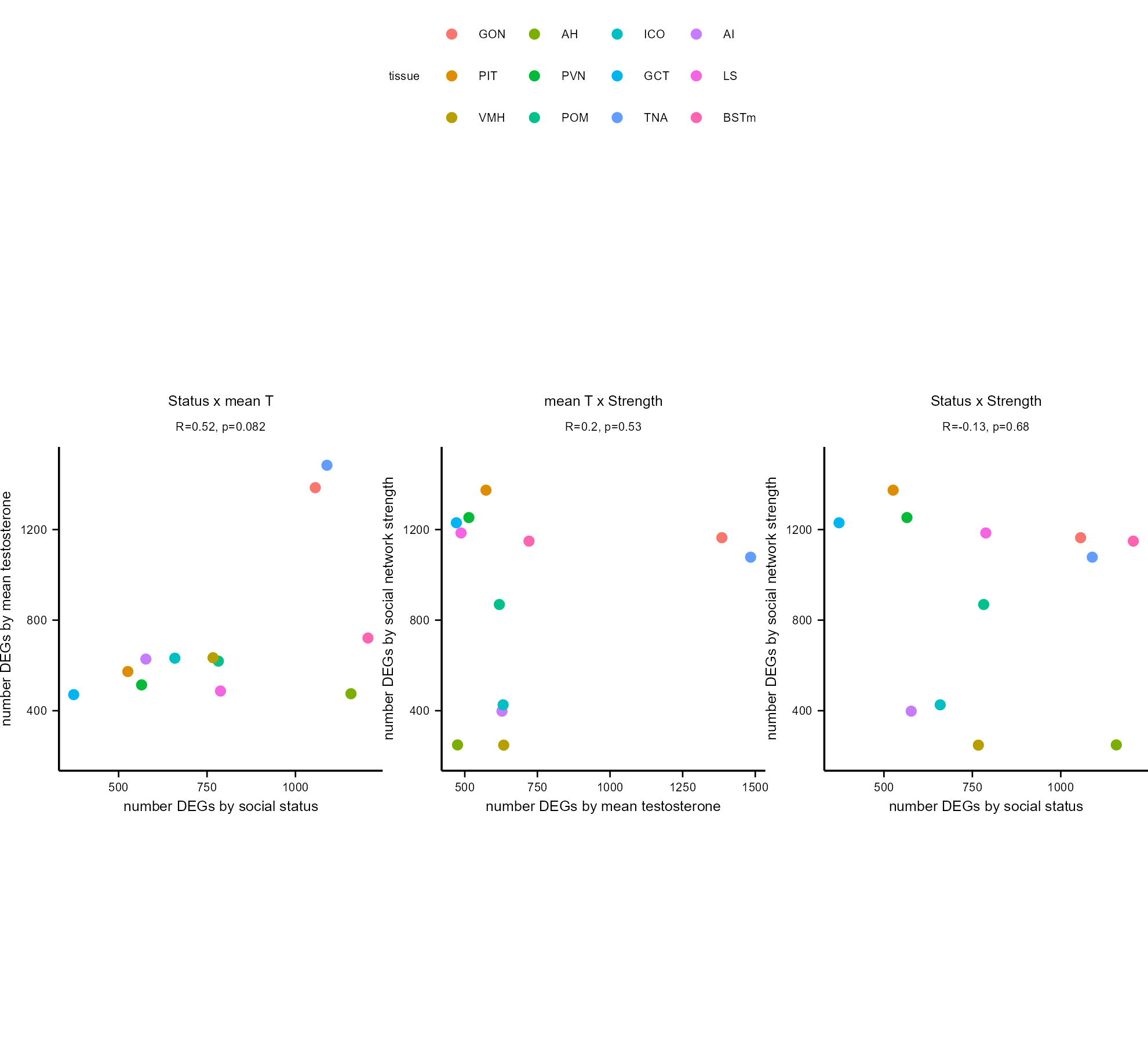

###### Figure S3.7: Correlation between number of differentially expressed genes in each tissue and comparison.

Colors indicate the tissue type, and the subheading indicates the correlation coefficient and its p-value.

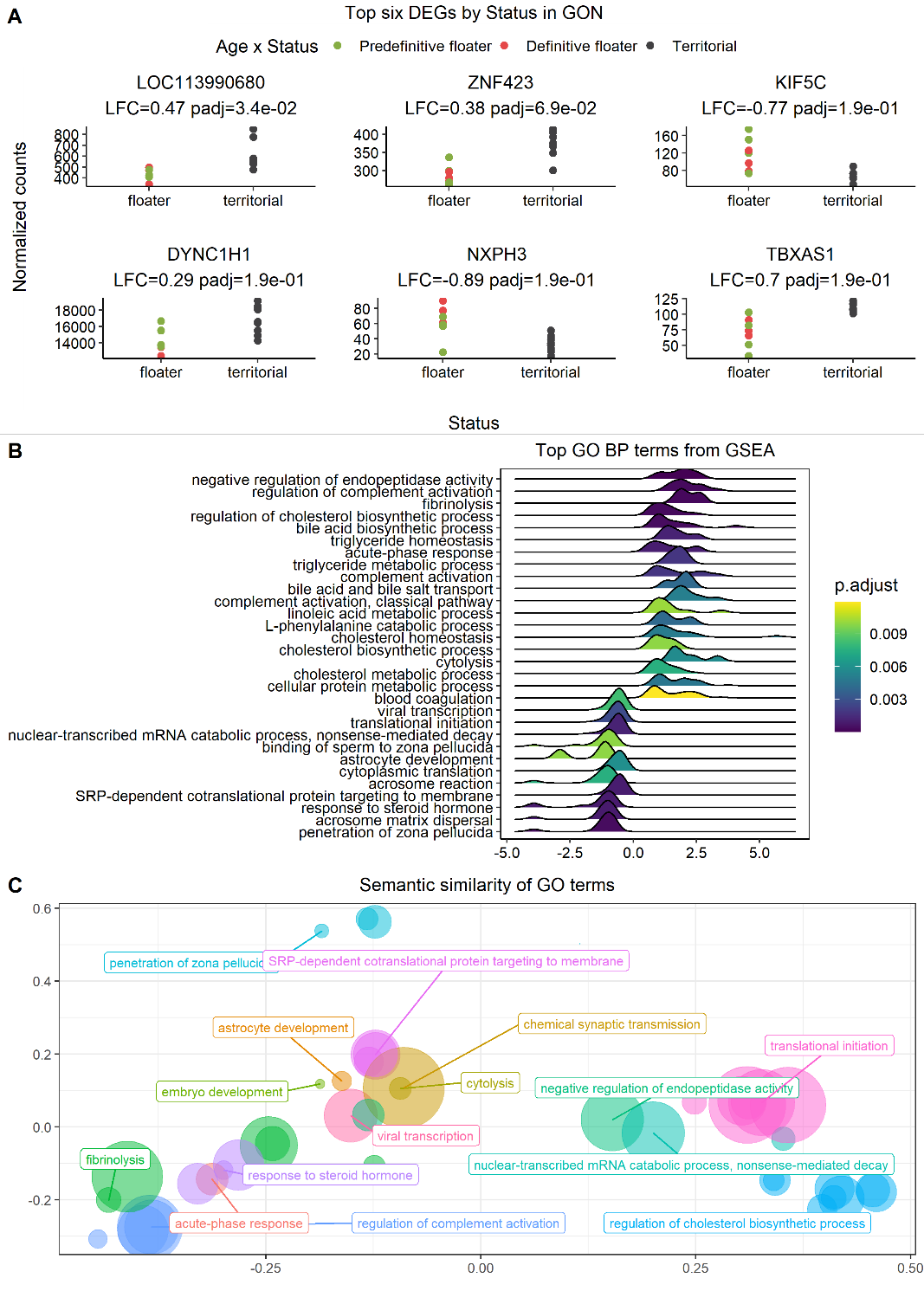

###### **Figure S3.8: Status associated DEGs in the testes (GON).**

A) The six genes with the lowest p-values for difference in gene expression between floater and territorial birds. B) Ridge plot showing the density of GO terms along a directional, log-transformed of gene p-values, therefore genes with low p-values are on the left and right edges of the plot. C) Bubble plot of semantically clustered GO terms where colors indicate the clustered GO terms, with the parent term labelled. The three-dimensional distance between points represents the semantic distance between terms, and the bubble size indicates enrichment score.

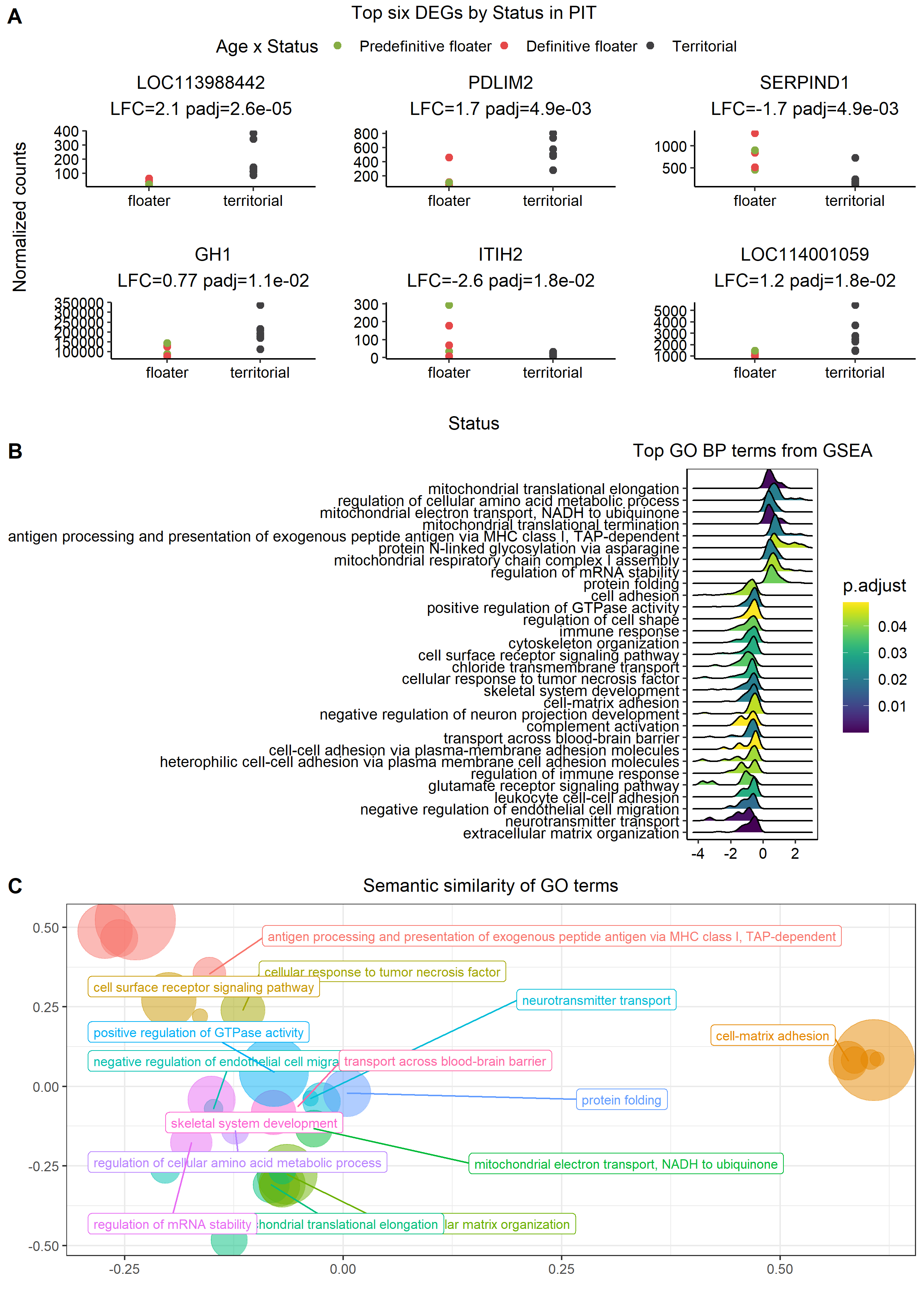

###### **Figure S3.9: Status associated DEGs in the pituitary (PIT).**

Explanations as Figure S3.8.

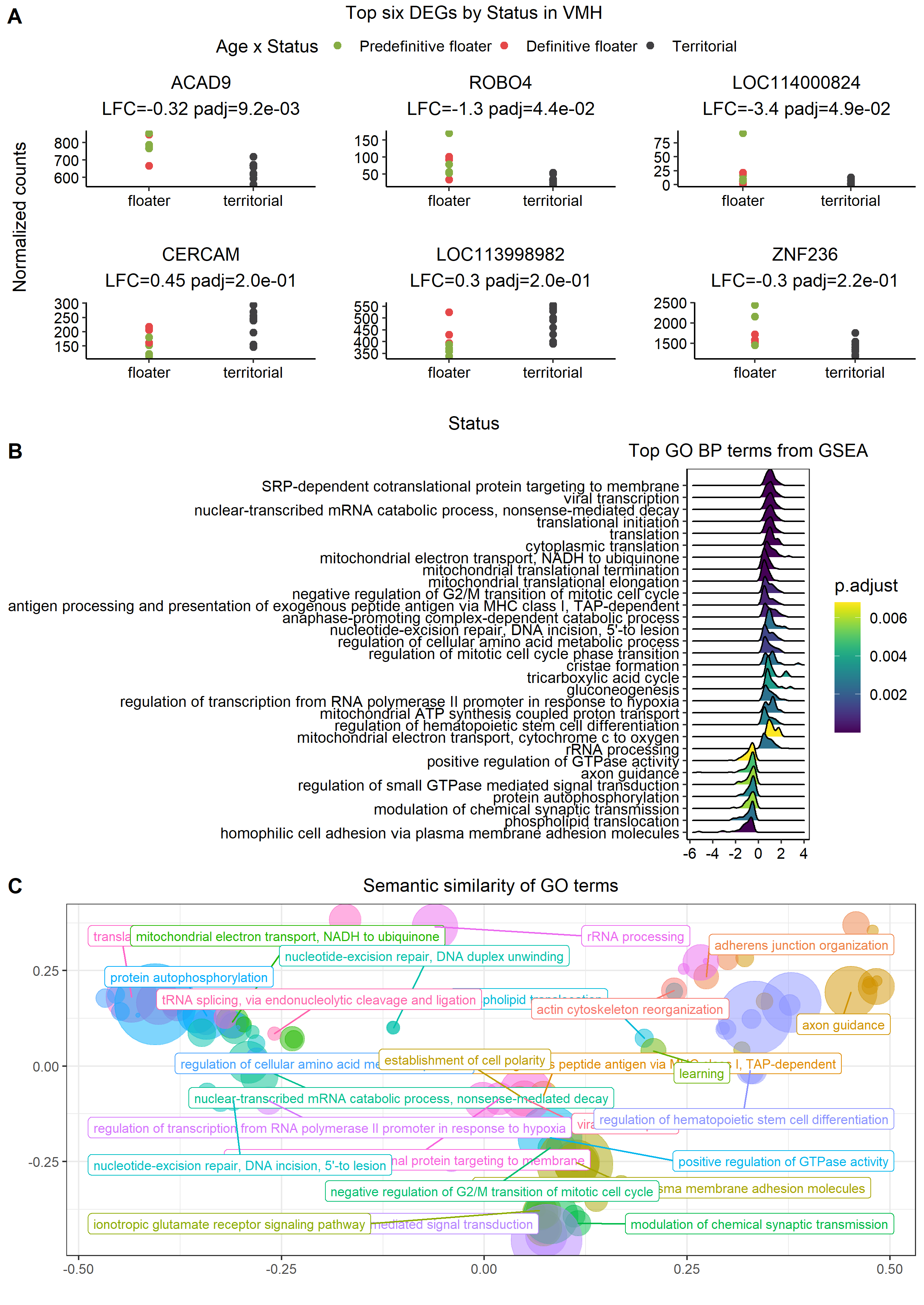

###### **Figure S3.10: Status associated DEGs in the ventromedial hypothalamus (VMH).**

Explanations as Figure S3.8.

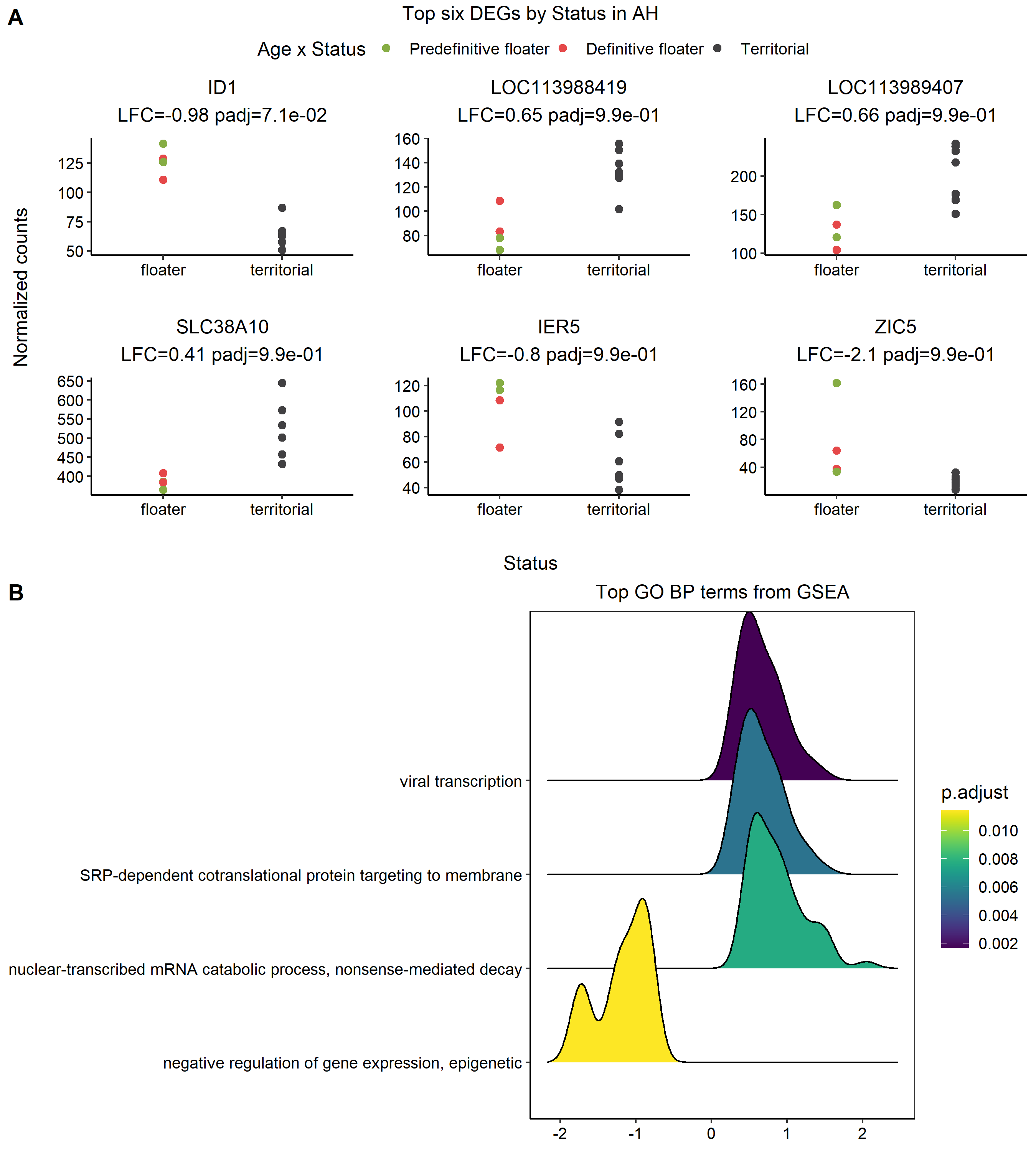

###### **Figure S3.11: Status associated DEGs in the anterior hypothalamus (AH).**

Explanations as Figure S3.8. There is no semantic similarity analysis here as only four GO terms were clustered in the gene list in this analysis.

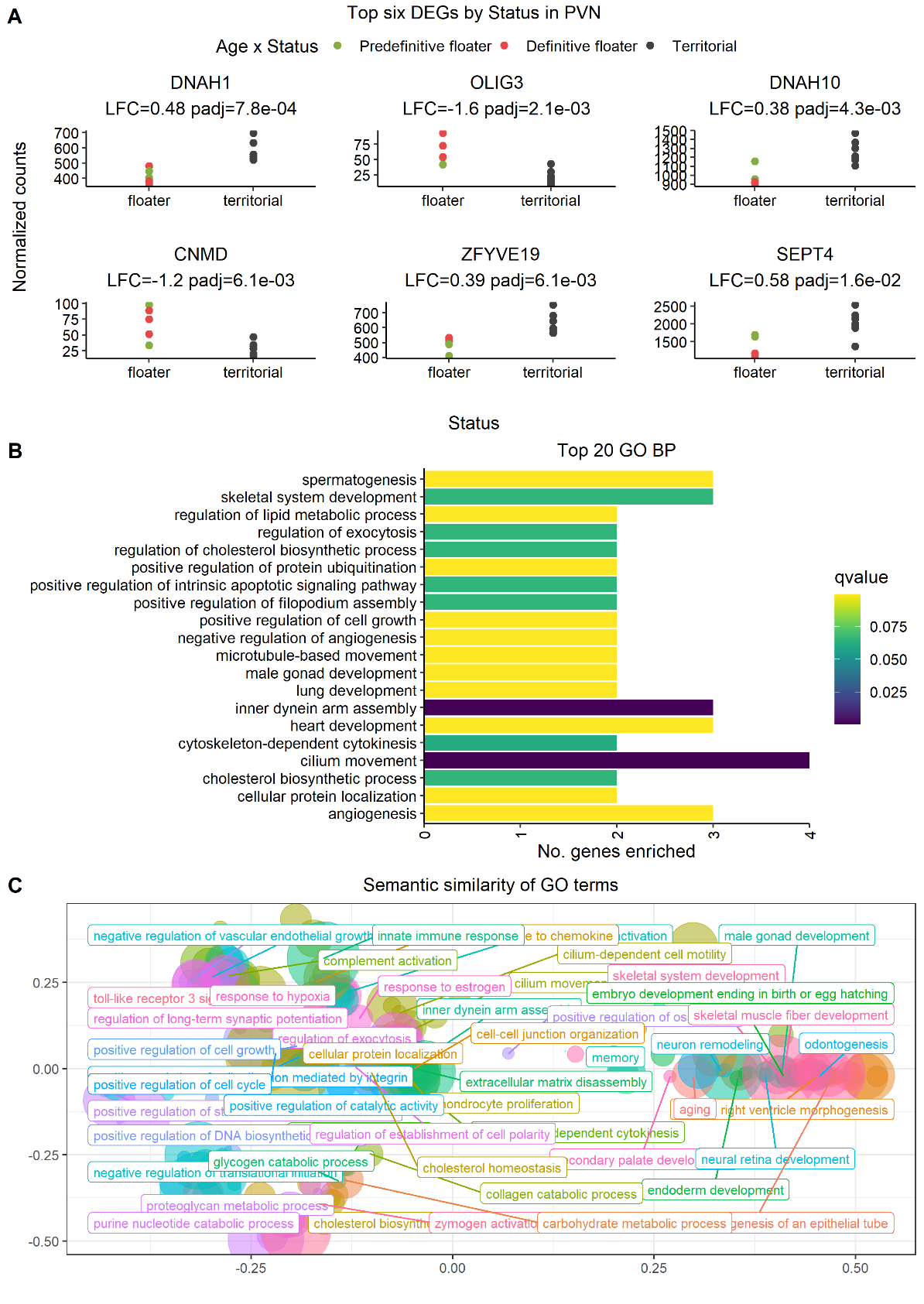

###### **Figure S3.12: Status associated DEGs in the paraventricular nucleus (PVN).**

A) The six genes with the lowest p-values for difference in gene expression between floater and territorial birds. B) Results from simple enrichment analysis of the 42 statistically significant differentially expressed genes, with bar color indicating the FDR corrected p-value. C) as Figure S3.8.

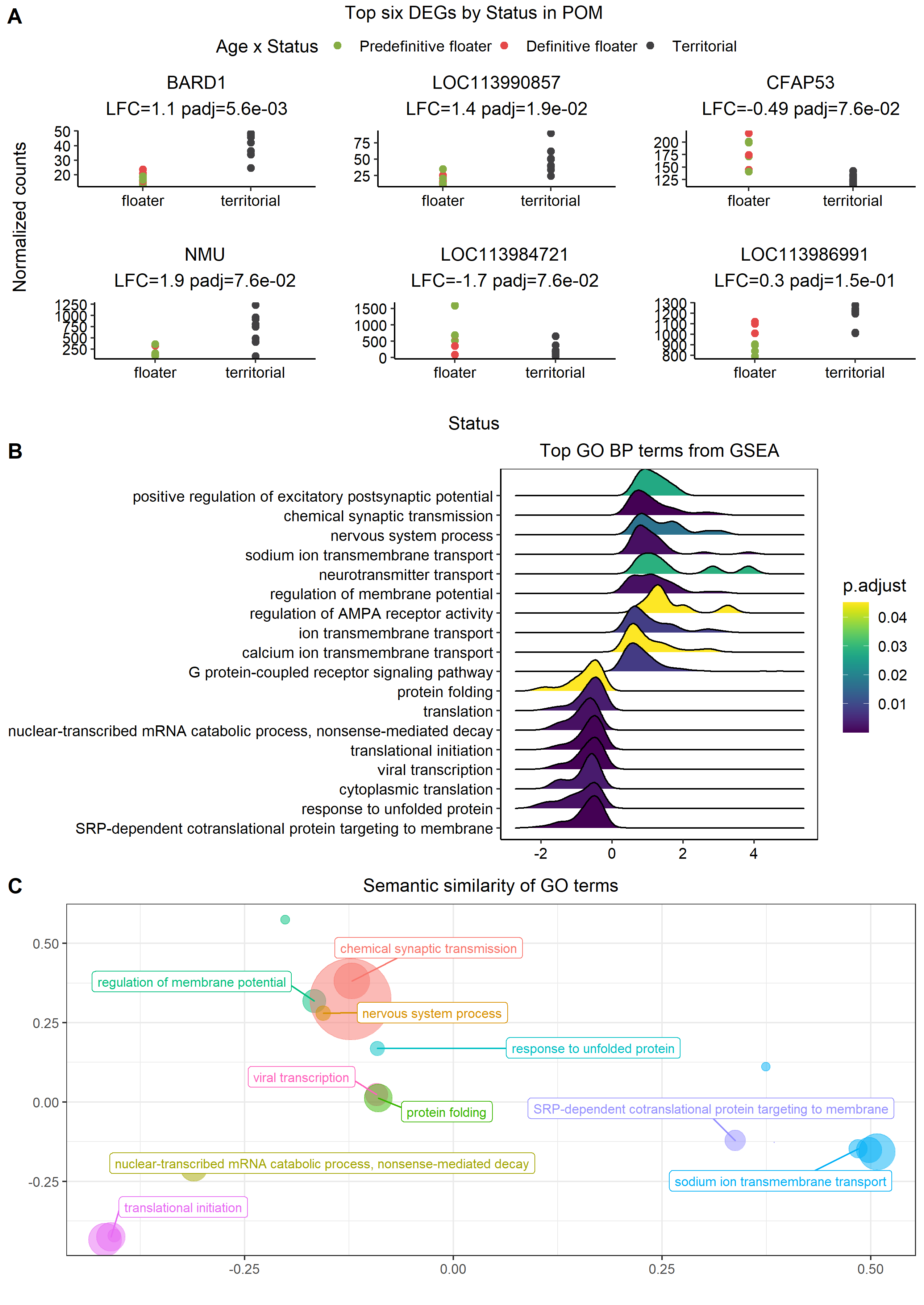

###### **Figure S3.13: Status associated DEGs in the medial preoptic nucleus (PVN).**

Explanations as Figure S3.8

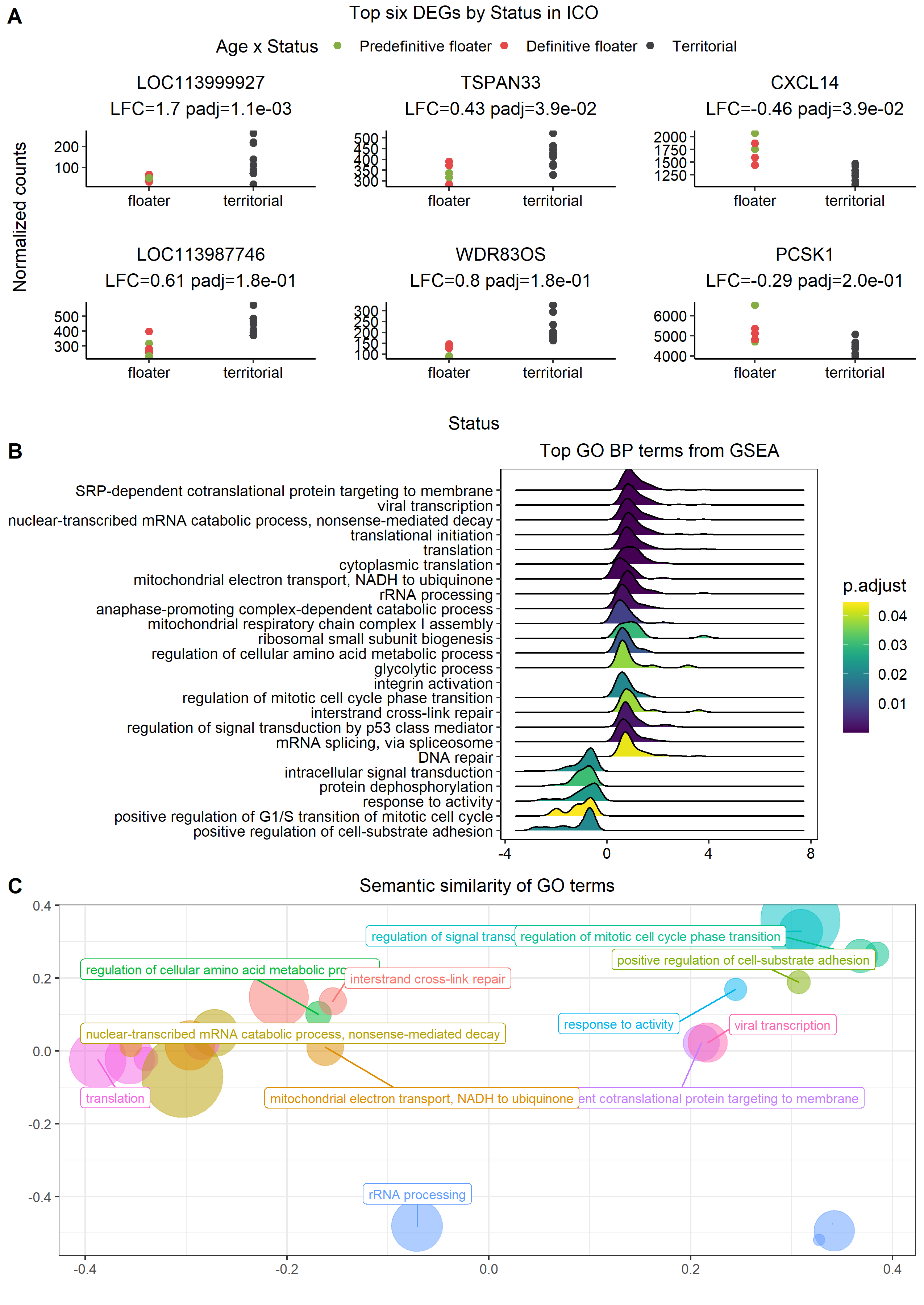

###### **Figure S3.14: Status associated DEGs in the intercollicular nucleus (ICo).**

Explanations as Figure S3.8

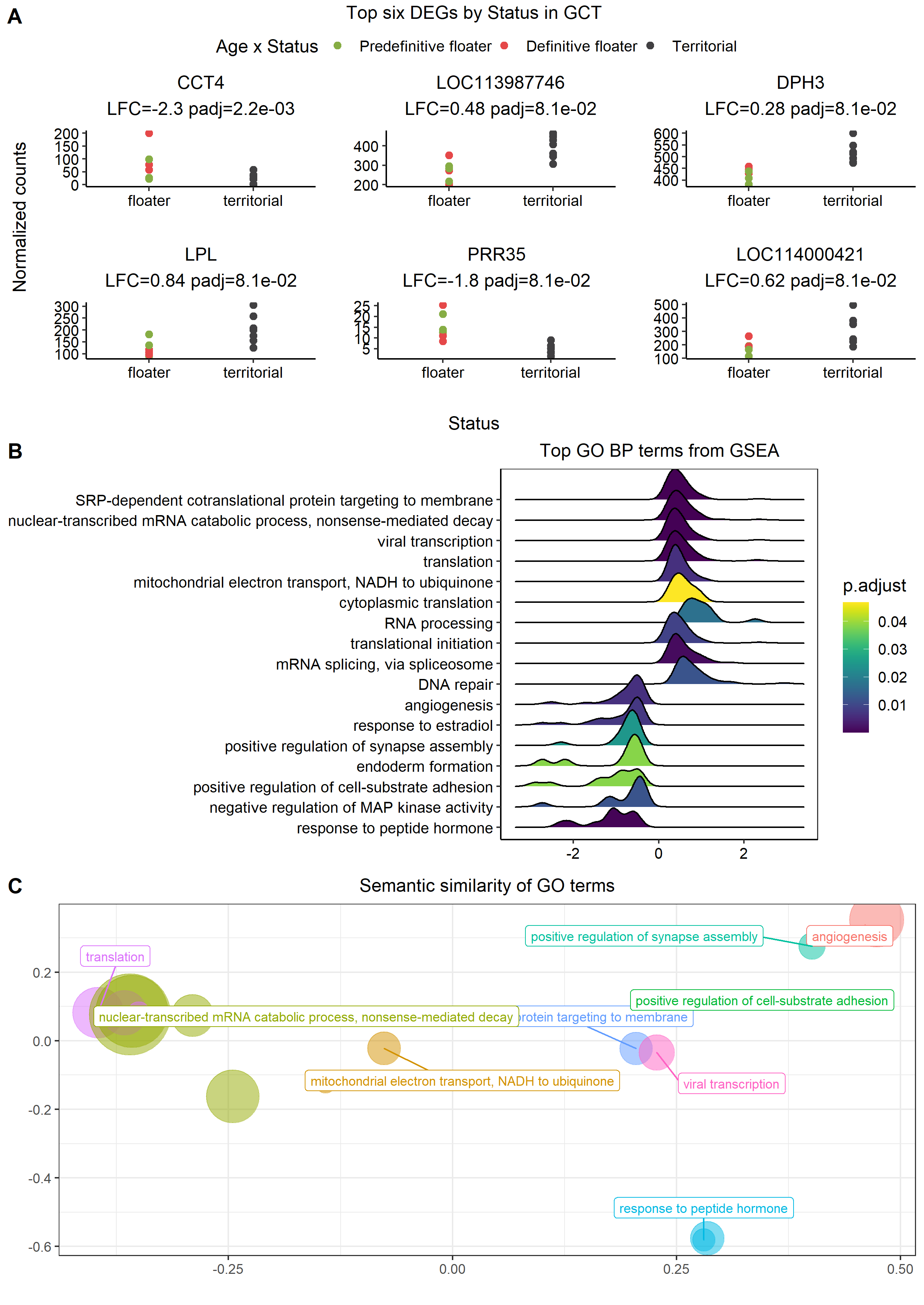

###### **Figure S3.15: Status associated DEGs in the midbrain central grey (GCt).**

Explanations as Figure S3.8

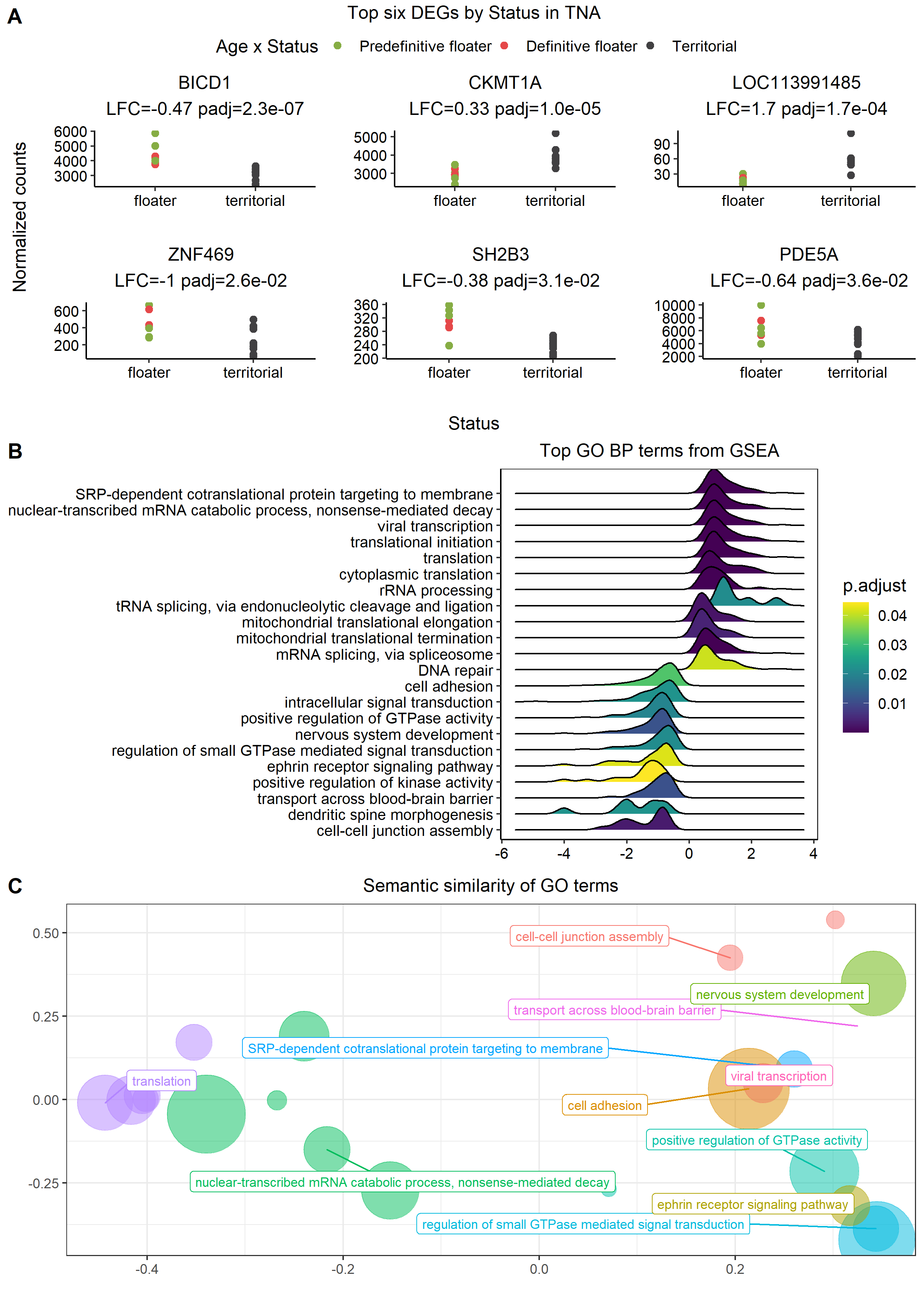

###### **Figure S3.16: Status associated DEGs in the nucleus taenia (TnA).**

Explanations as Figure S3.8

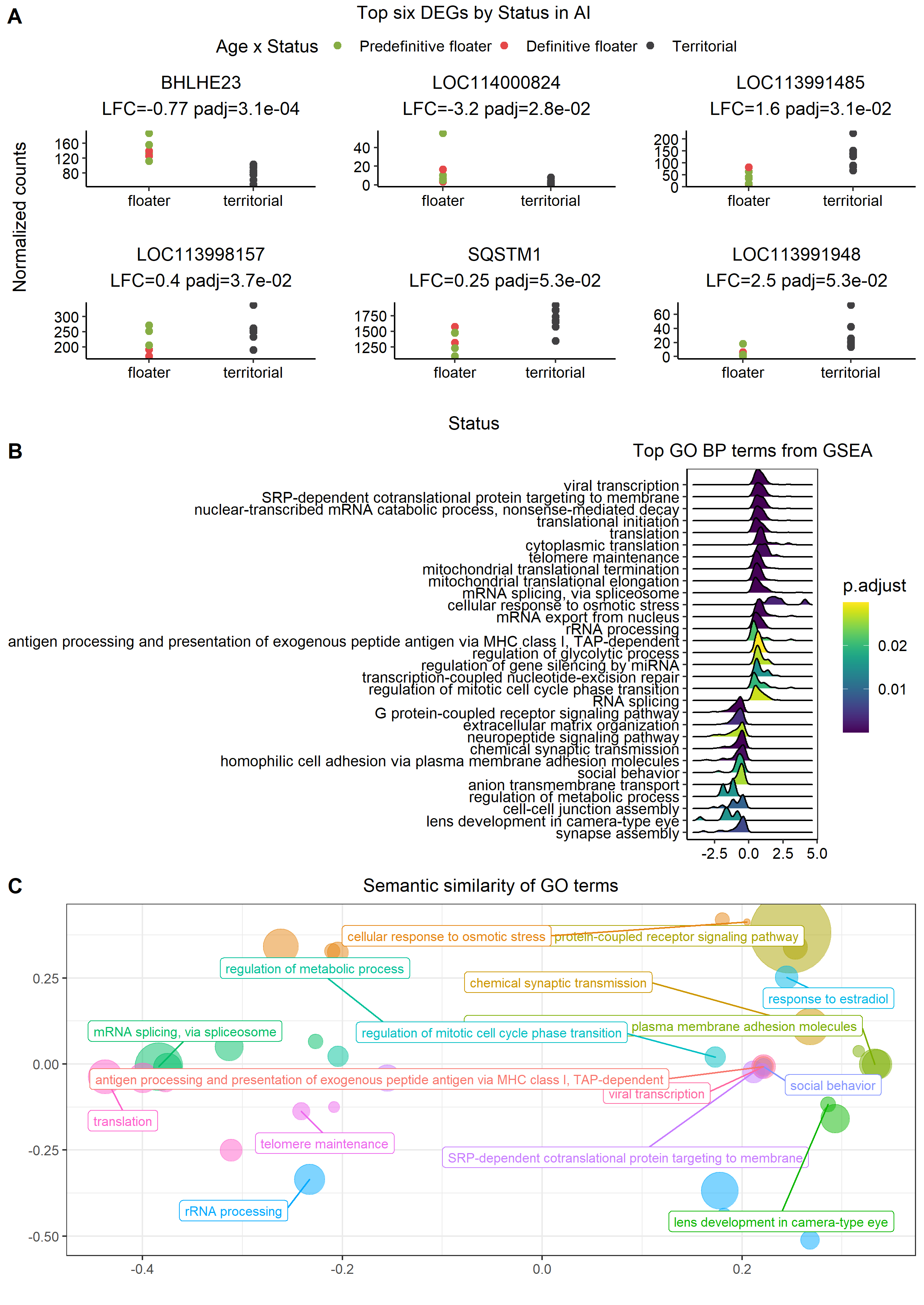

###### Figure S3.17: Status associated DEGs in the arcopallium intermedium (AI).

Explanation as in Figure S3.8.

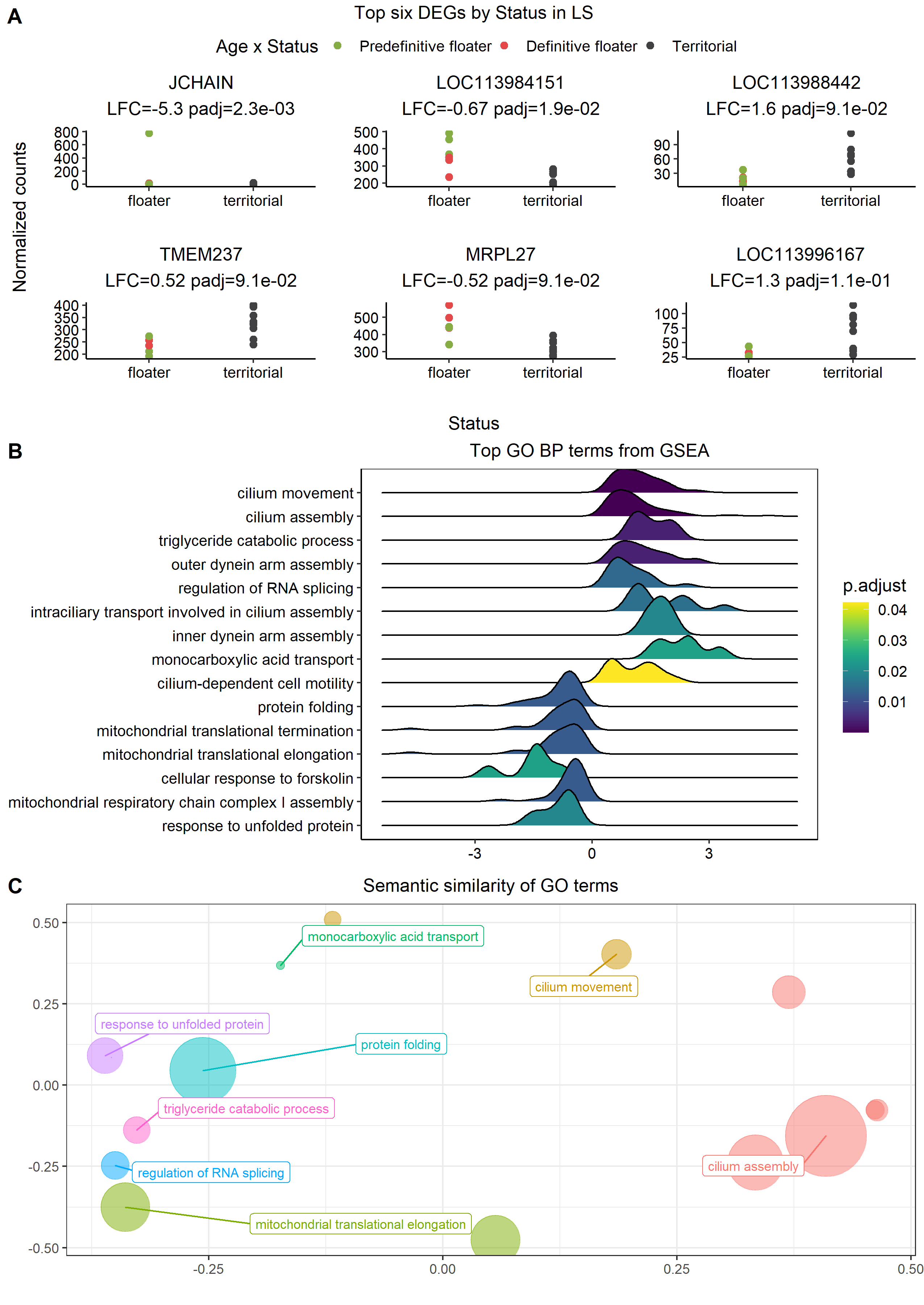

###### **Figure S3.18: Status associated DEGs in the lateral septum (LS).**

Explanations as Figure S3.8.

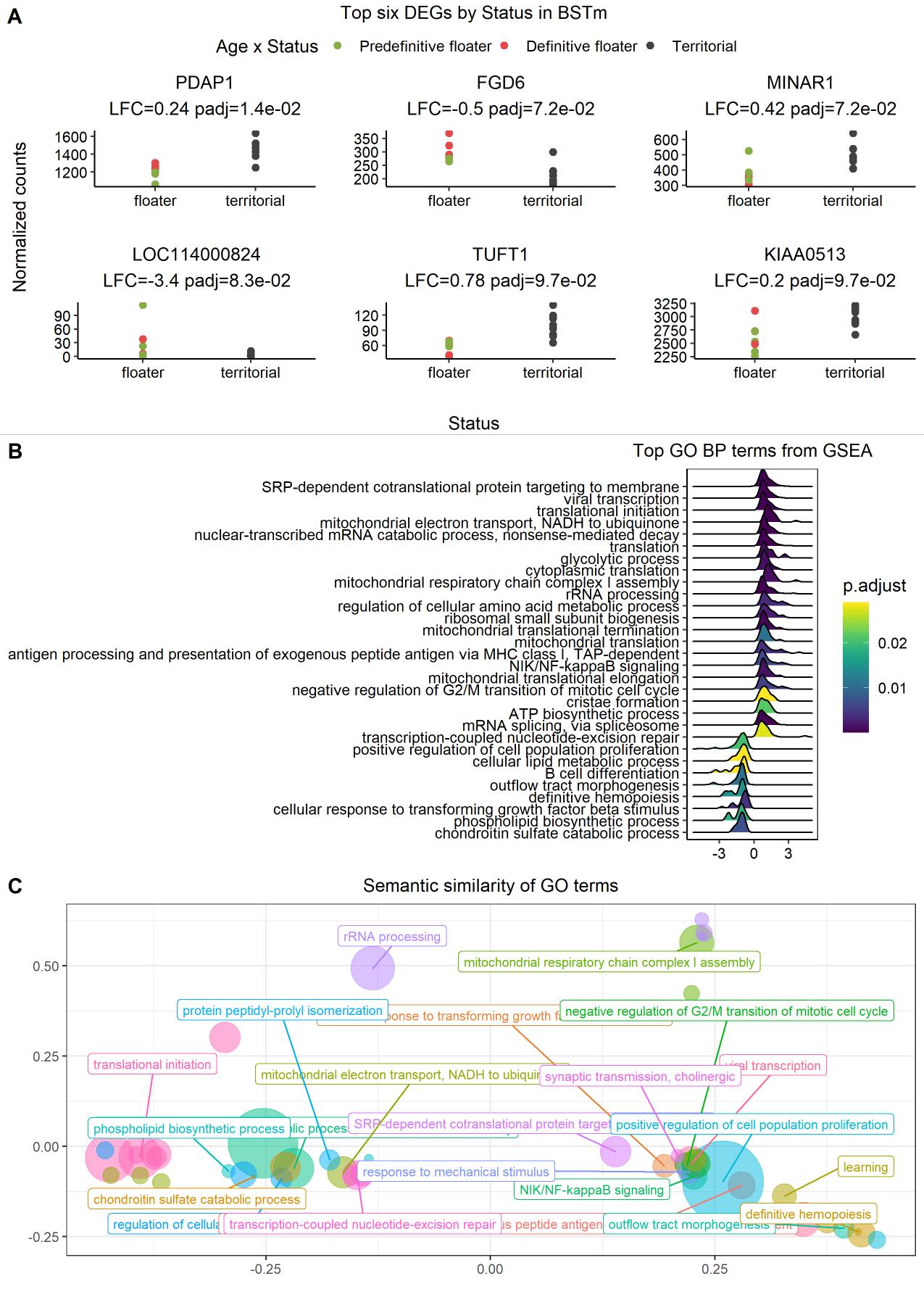

###### **Figure S3.19: Status associated DEGs in the bed nucleus of the stria terminalis (BSTm).**

Explanations as Figure S3.8

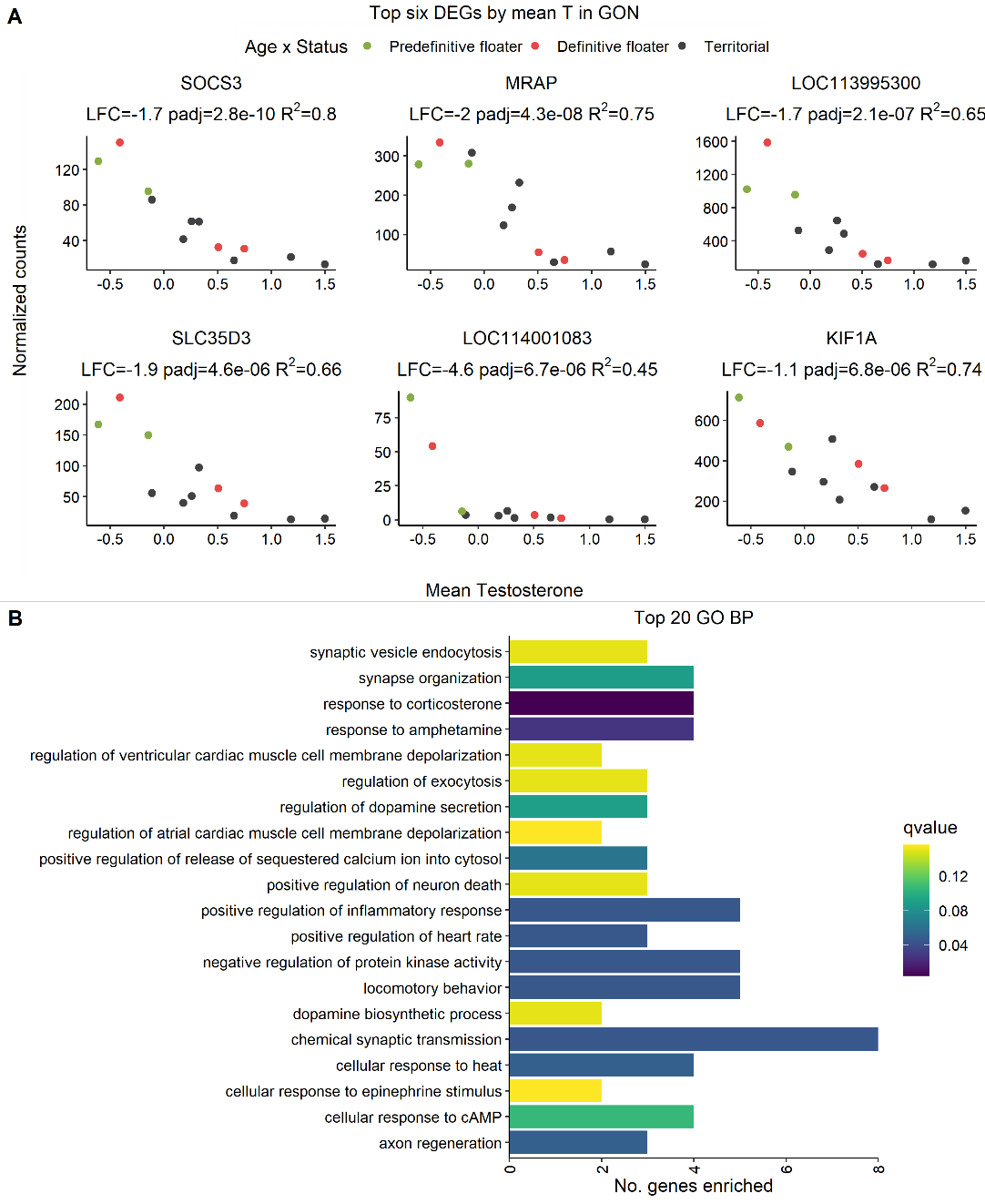

###### **Figure S3.20: Mean testosterone phenotype associated DEGs in the testes (GON).**

A) The six genes with the lowest p-values for difference in gene expression in individuals with high or low mean testosterone phenotype (mean T). B) Ridge plot showing the density of GO terms along a directional, log-transformed of gene p-values, therefore genes with low p-values are on the left and right edges of the plot. The bubble plot contained too many terms to be useful, but are presented as in Figure S3.7.

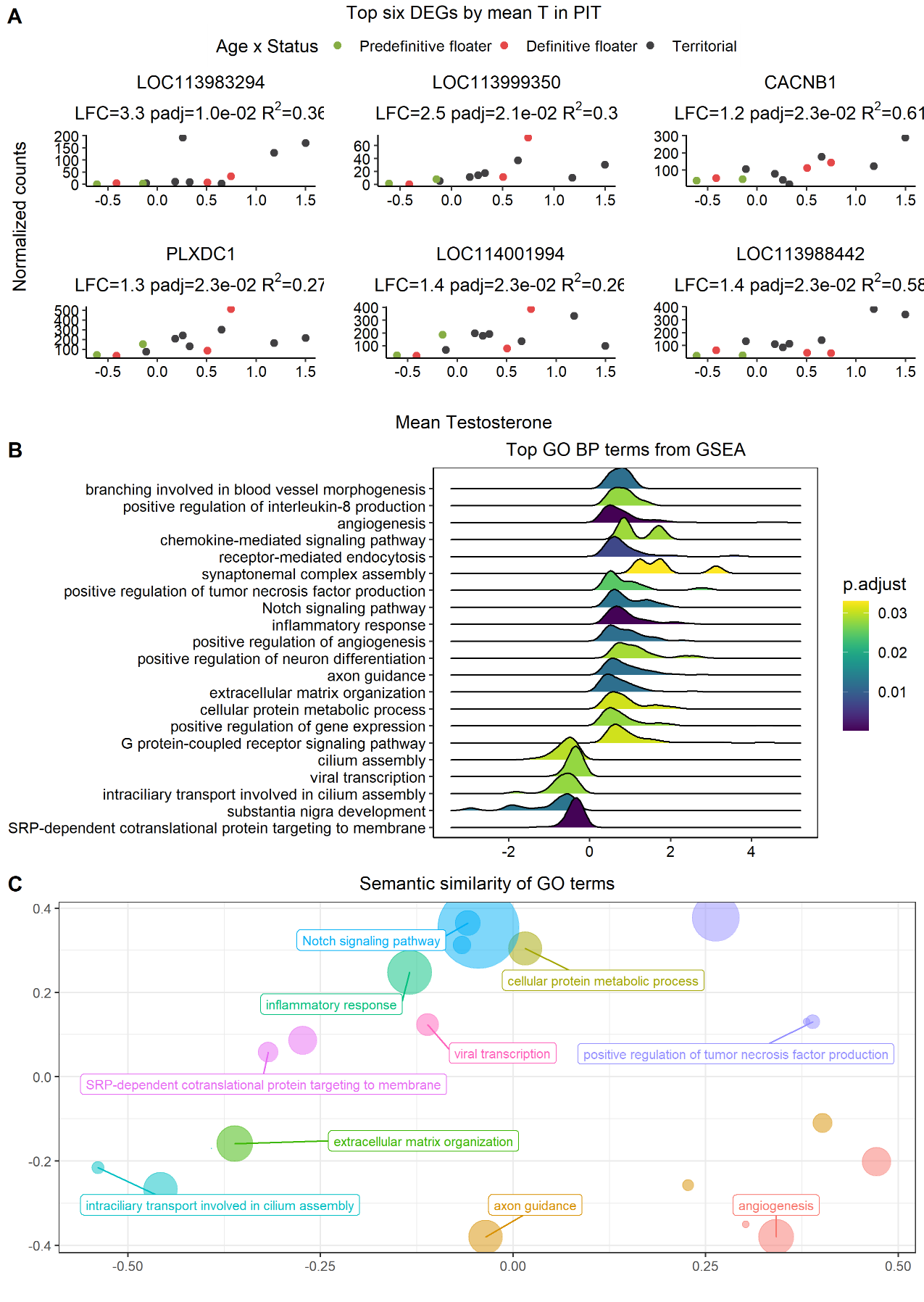

###### **Figure S3.21: Mean testosterone phenotype associated DEGs in the pituitary (PIT).**

Explanations as Figure S3.20.

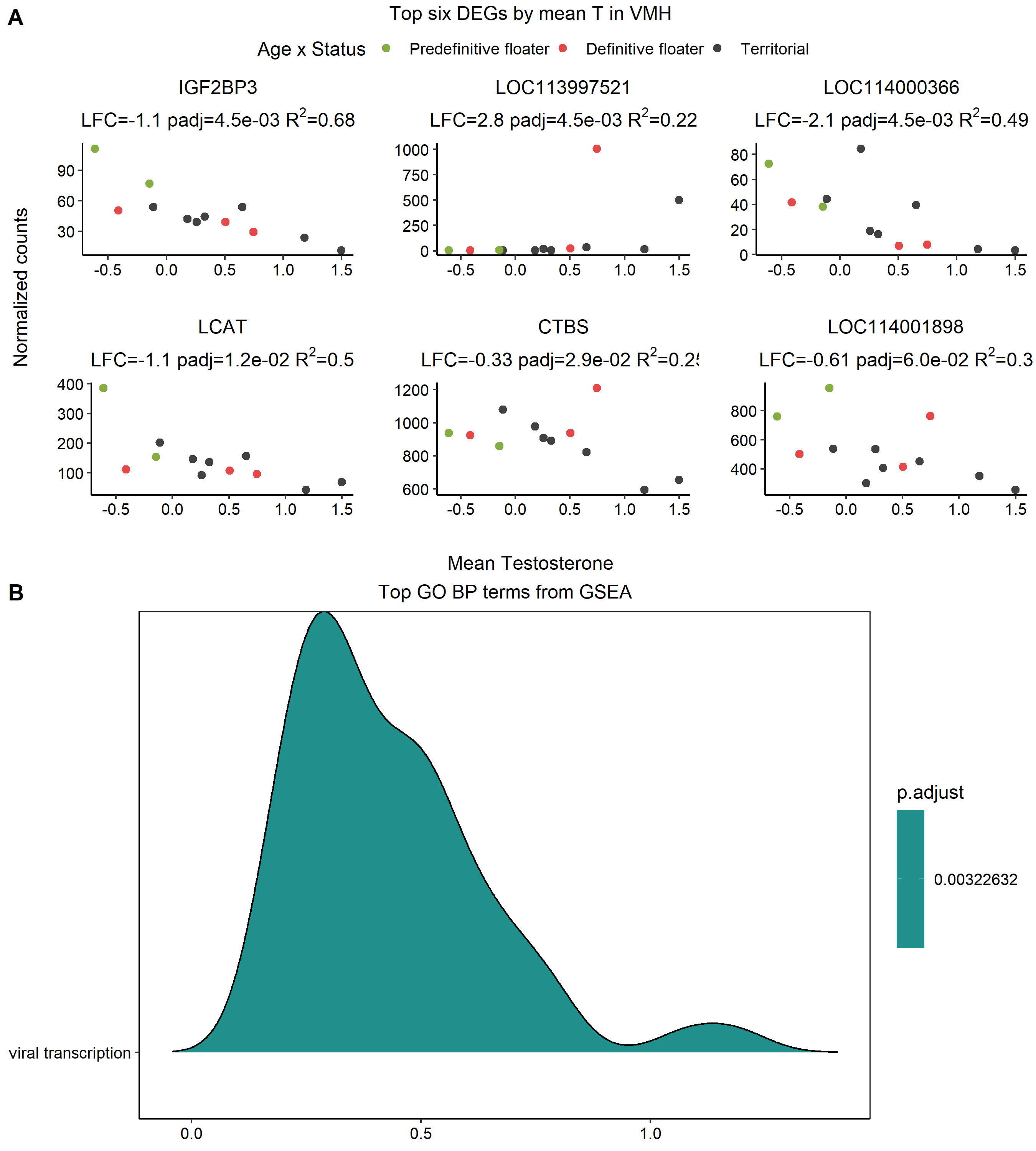

###### Figure S3.22: Mean testosterone phenotype associated DEGs in the ventromedial hypothalamus (VMH).

Explanations as in Figure S3.20. Viral transcription was the only GO term with a coherent signal along the gene list.

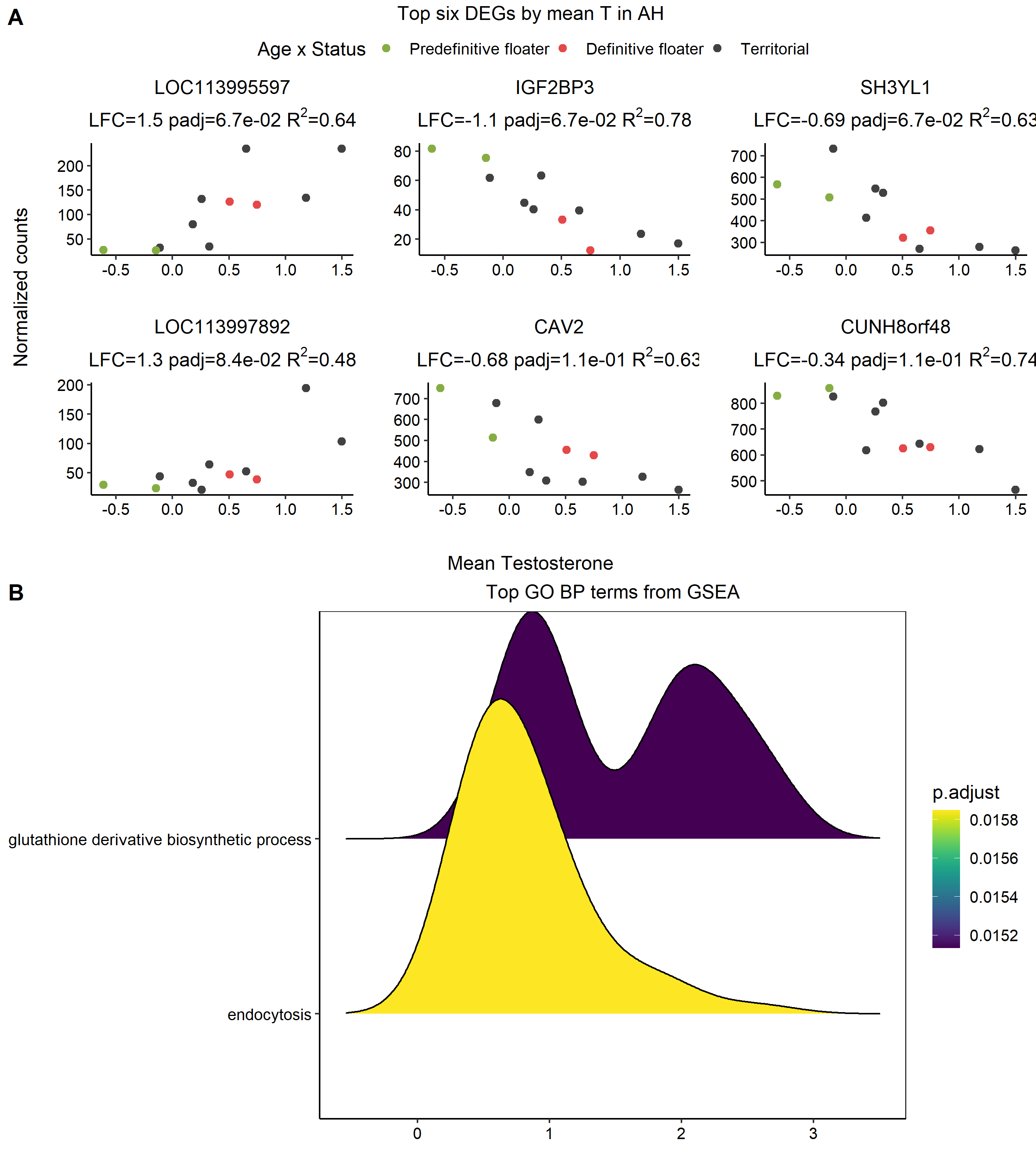

###### Figure S3.23: Mean testosterone phenotype associated DEGs in the anterior hypothalamus (AH).

Explanation as in Figure S3.20.

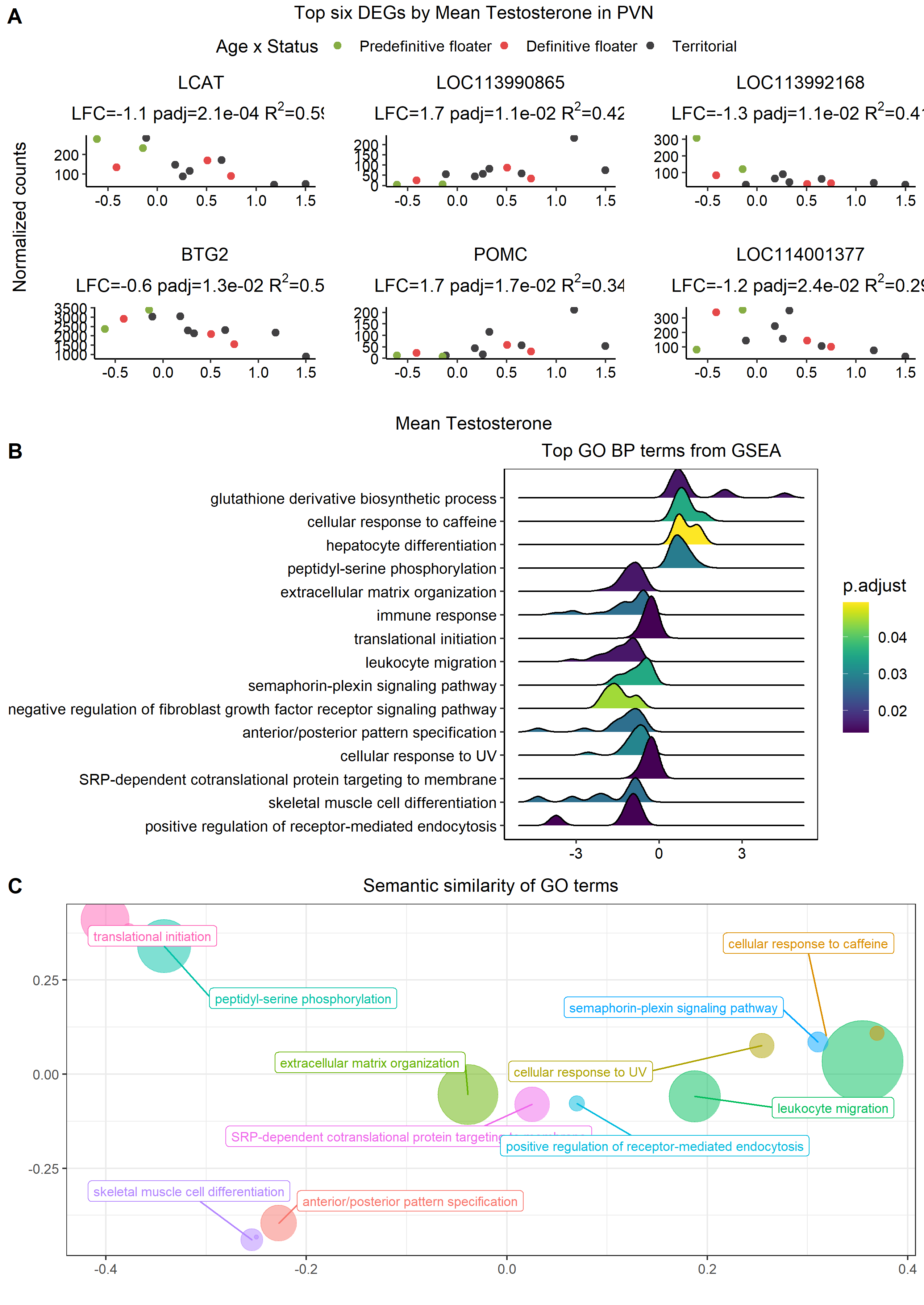

###### Figure S3.24: Mean testosterone phenotype associated DEGs in the paraventricular nucleus (PVN).

Explanation as in Figure S3.20.

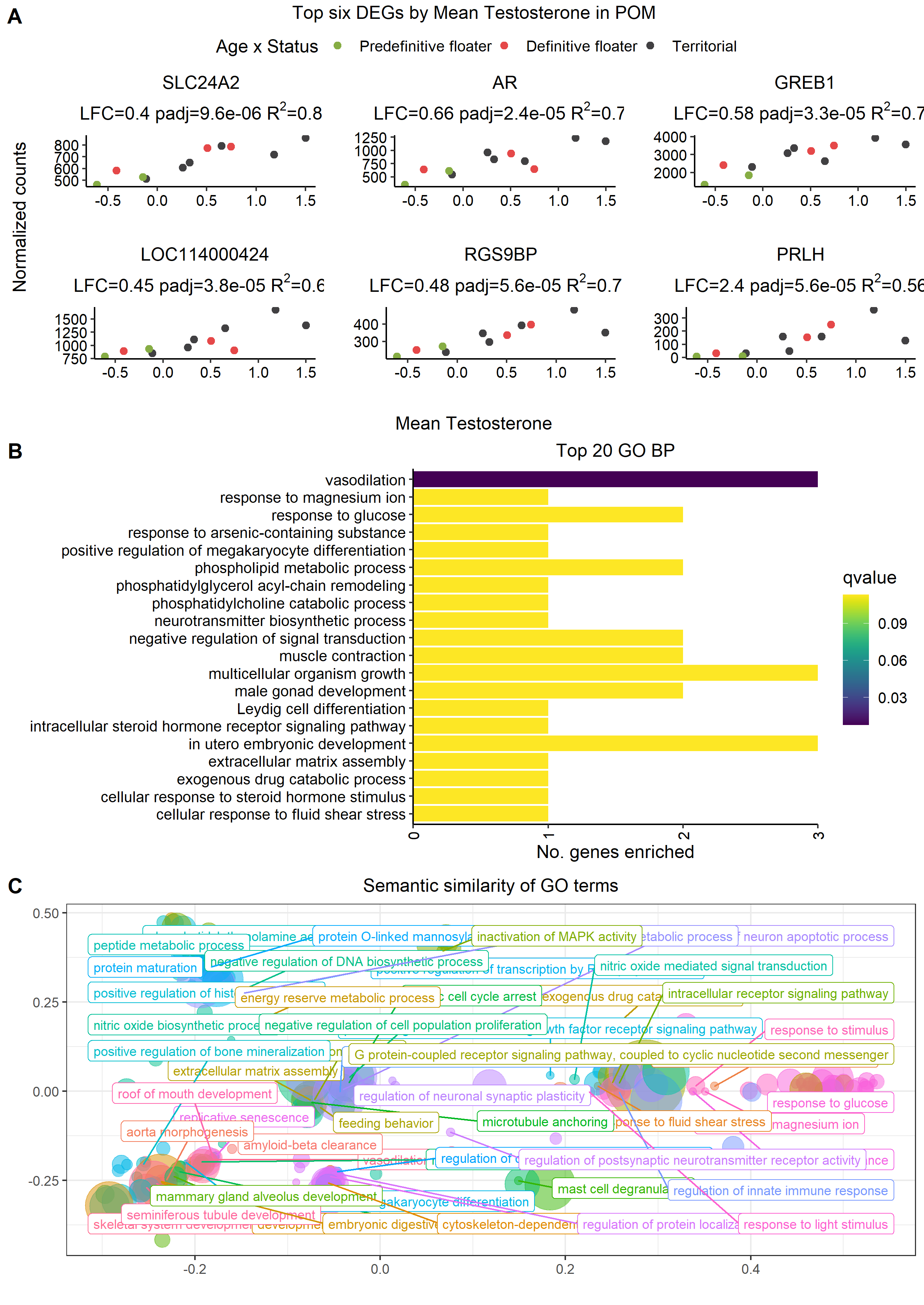

###### Figure S3.25: Mean testosterone phenotype associated DEGs in the medial preoptic area (POM).

Explanations as in Figure S3.20.

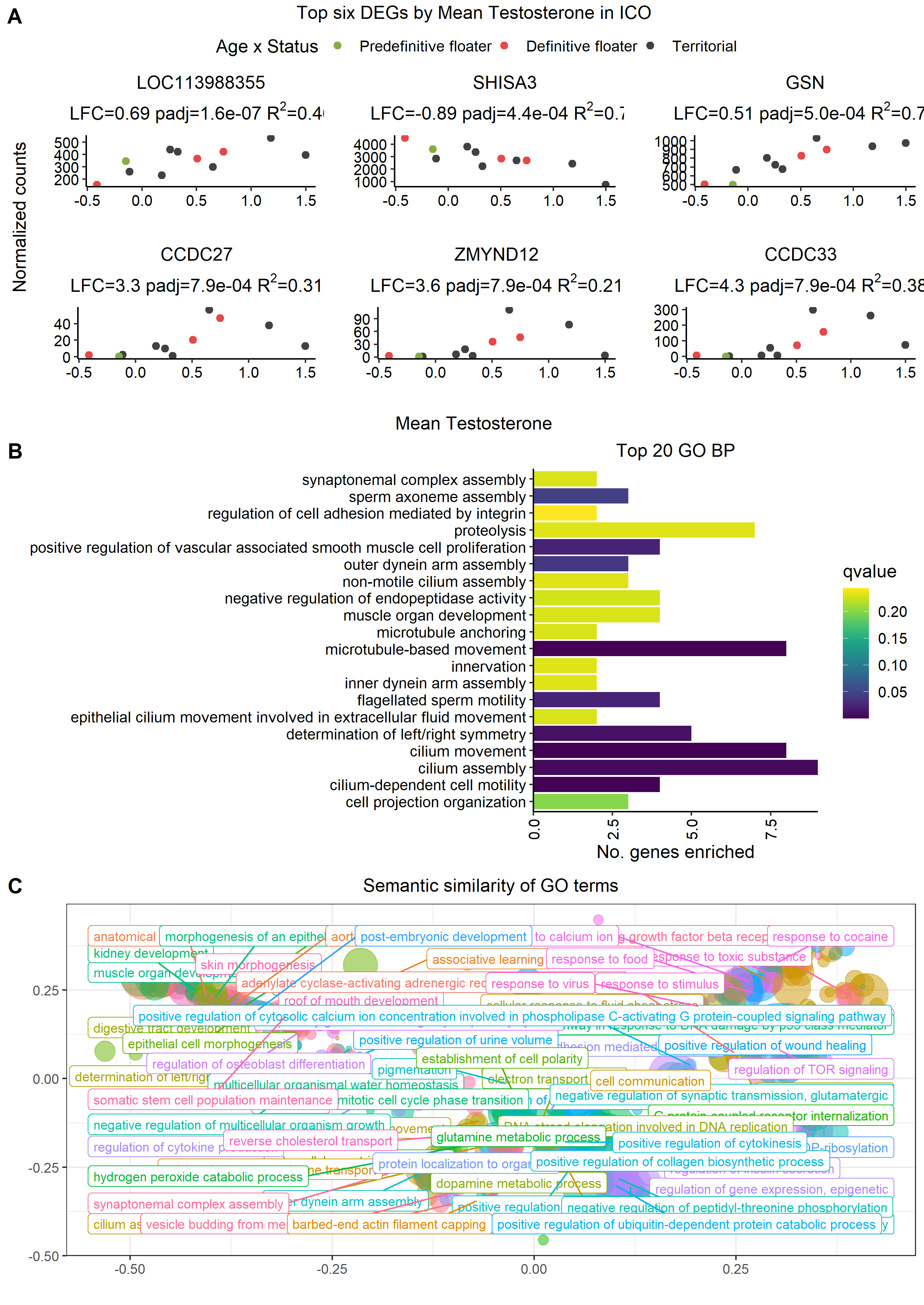

###### Figure S3.26: Mean testosterone phenotype associated DEGs in the intercollicular nucleus (ICo).

Explanations as in Figure S3.20.

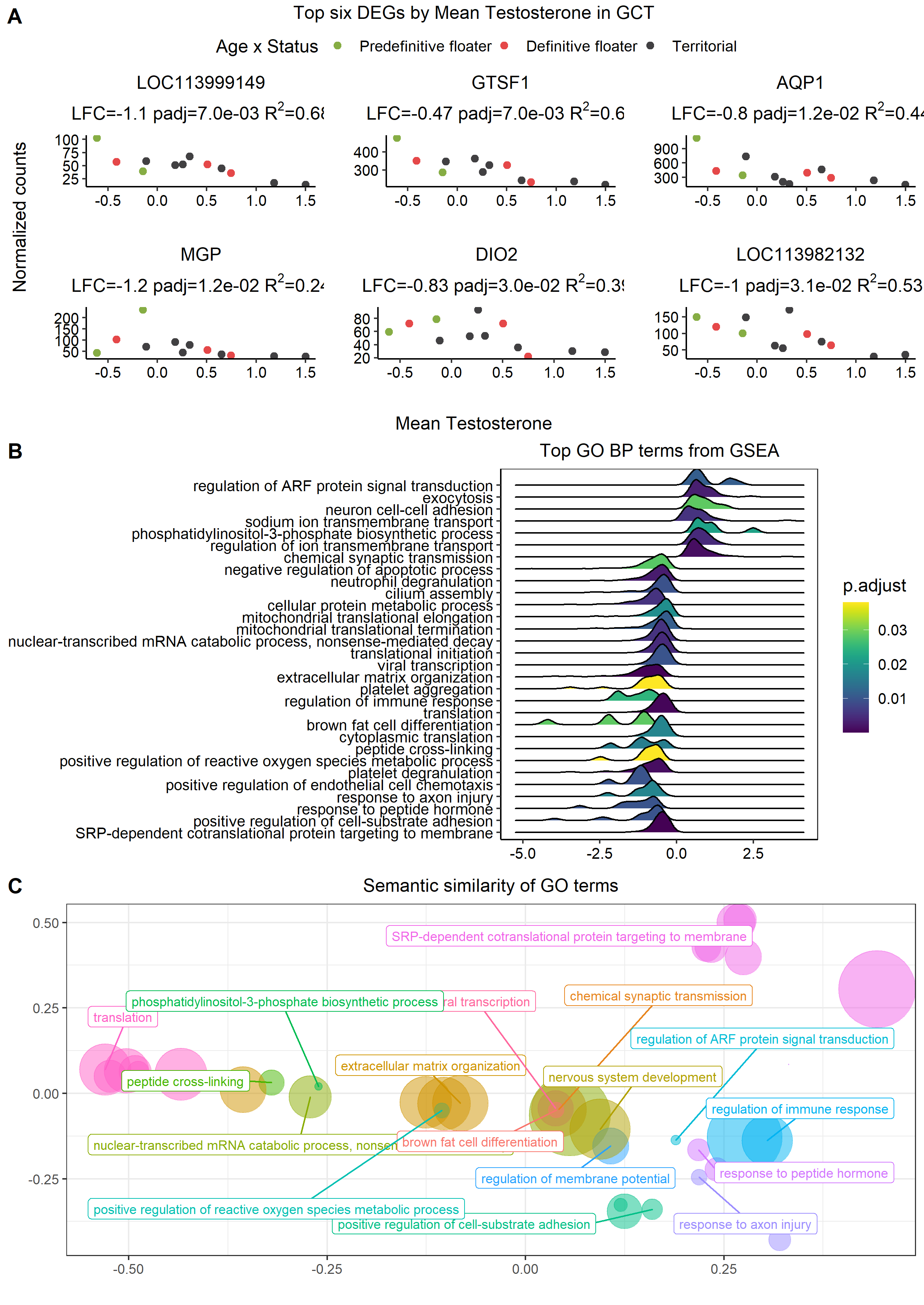

###### Figure S3.27: Mean testosterone phenotype associated DEGs in the midbrain central grey (GCt).

Explanations as in Figure S3.20.

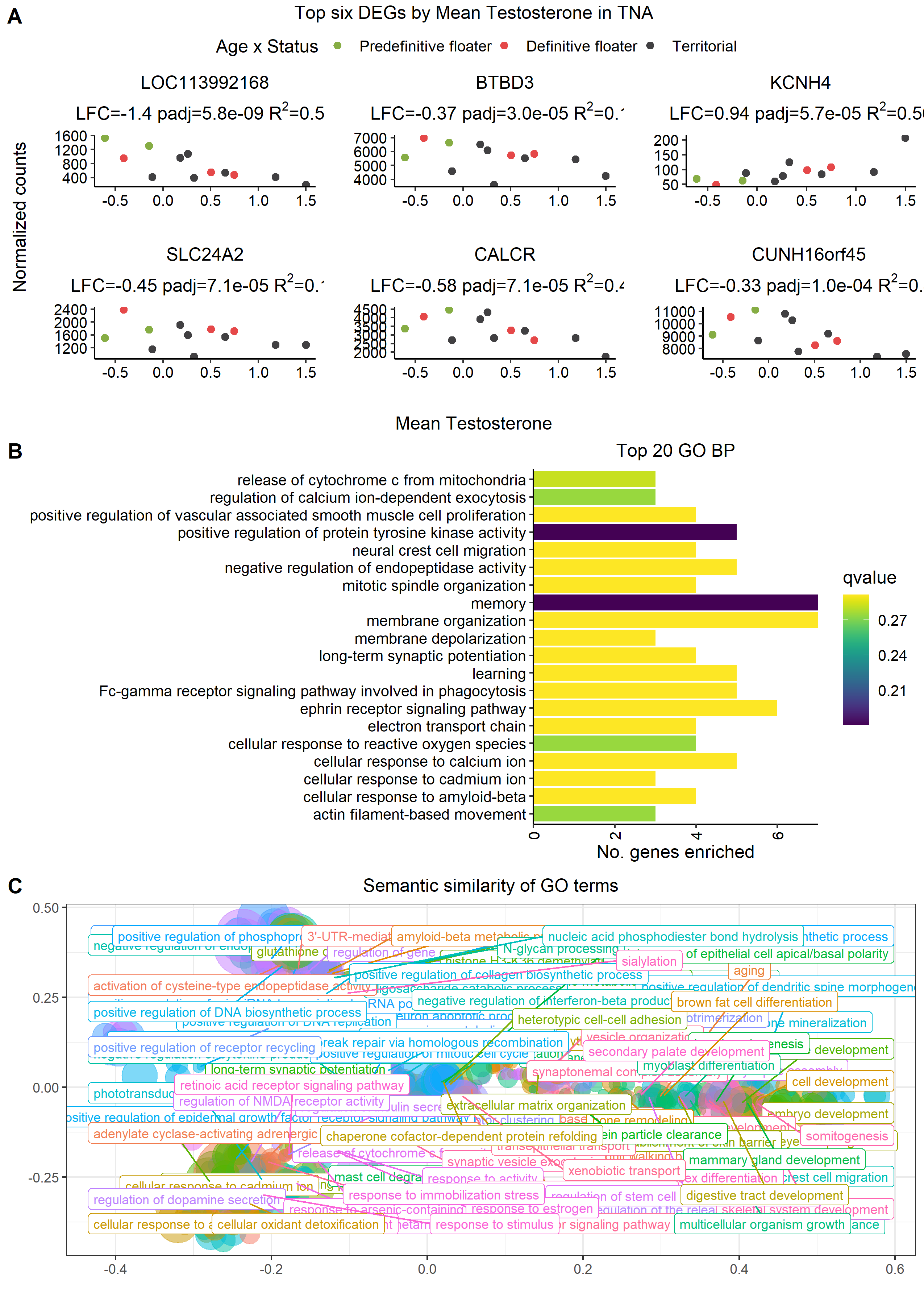

###### Figure S3.28: Mean testosterone phenotype associated DEGs in the nucleus taenia (TnA).

Explanations as in Figure S3.20.

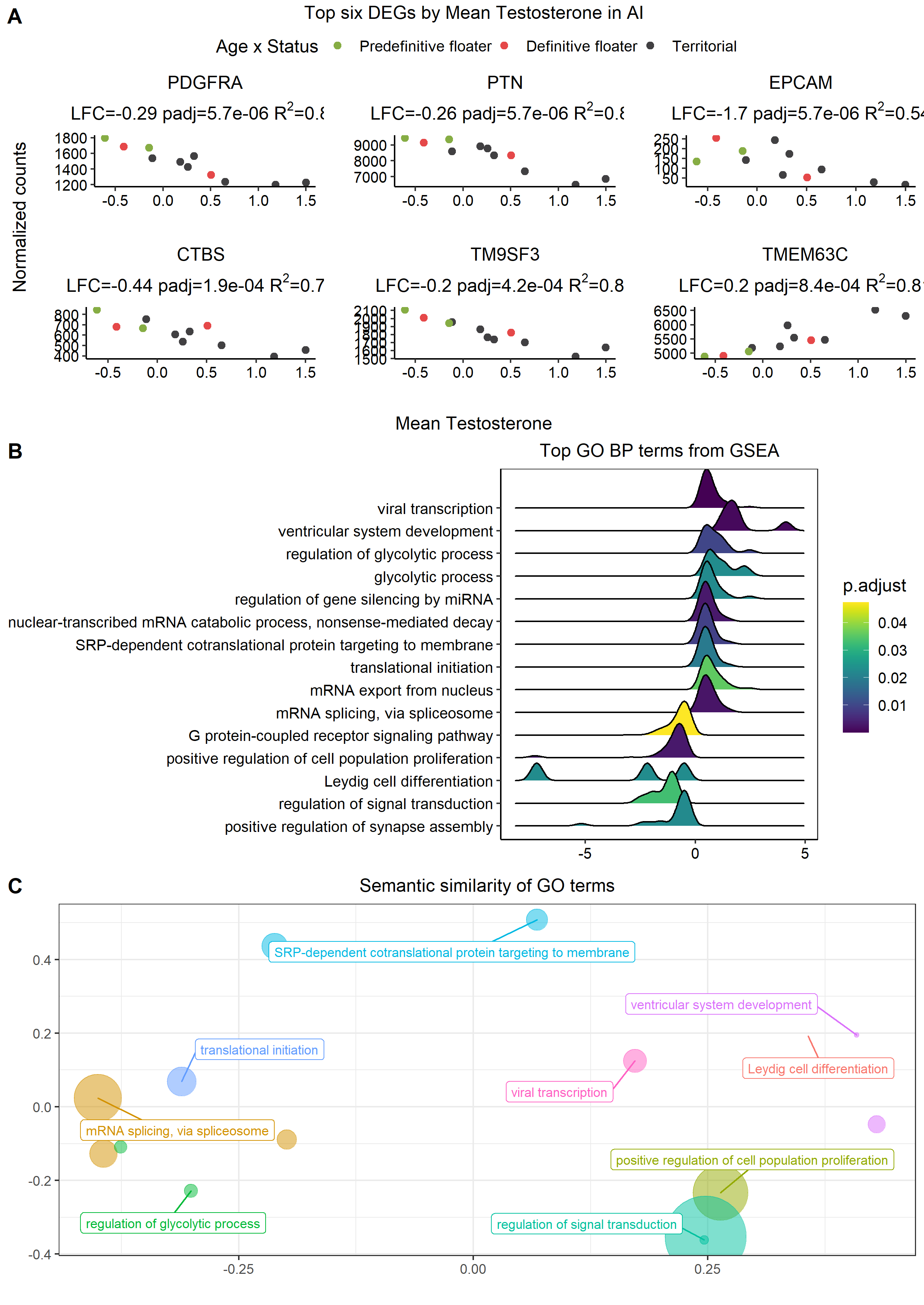

###### Figure S3.29: Mean testosterone phenotype associated DEGs in the arcopallium intermedium (AI).

Explanations as in Figure S3.20.

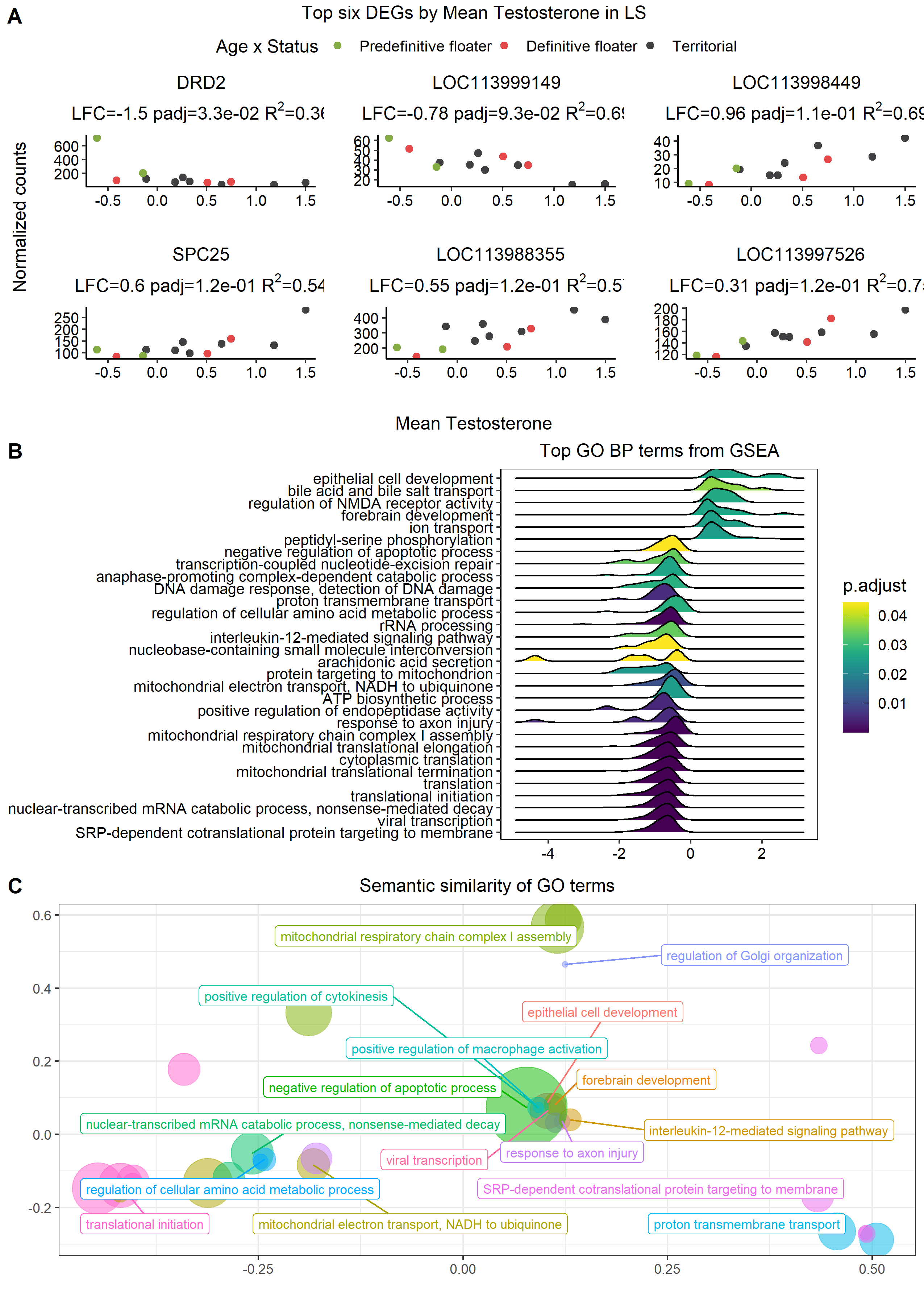

###### Figure S3.30: Mean testosterone phenotype associated DEGs in the lateral septum (LS).

Explanations as in Figure S3.20.

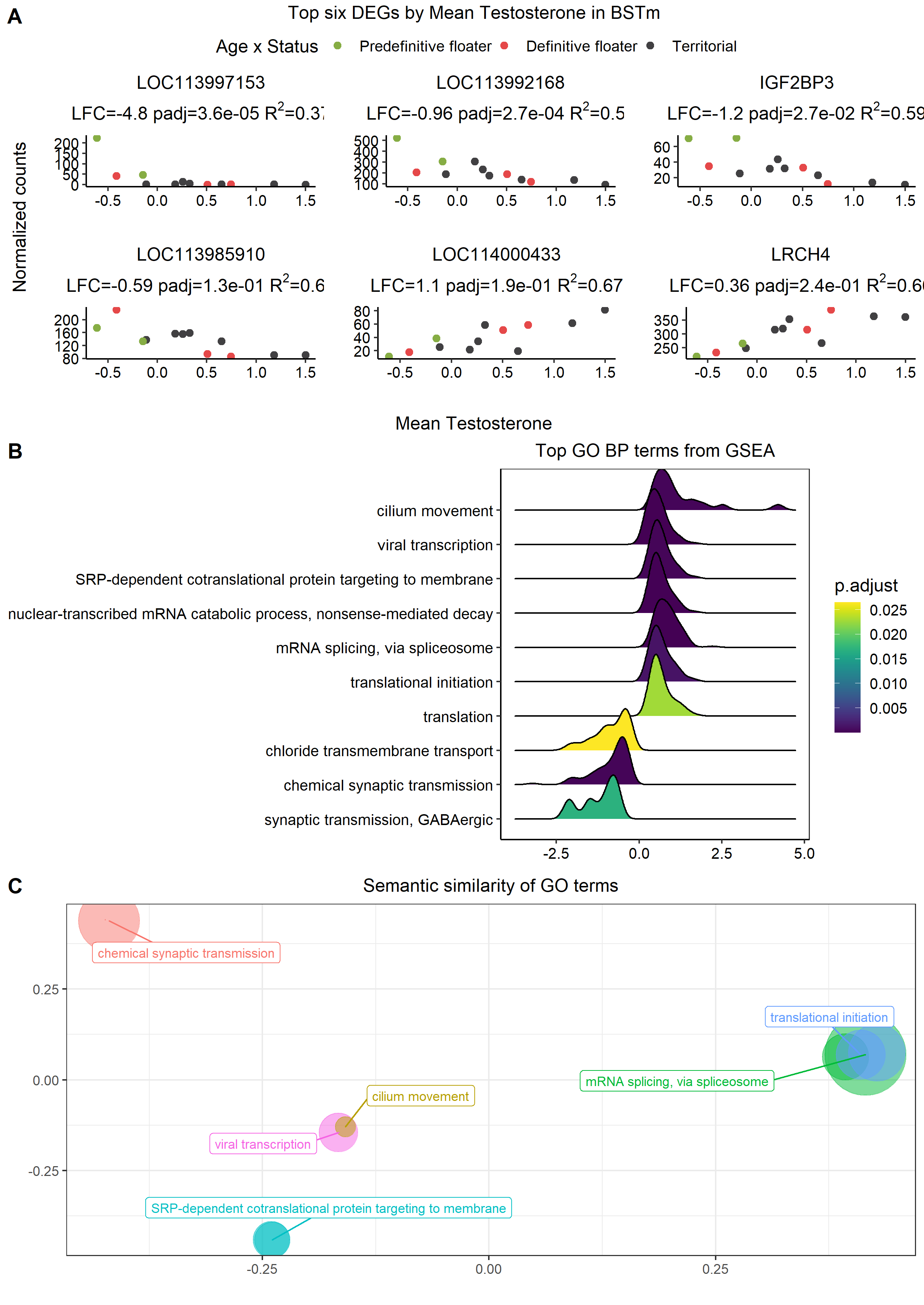

###### Figure S3.31: Mean testosterone phenotype associated DEGs in the bed nucleus of the stria terminalis (BSTm).

Explanations as in Figure S3.20.

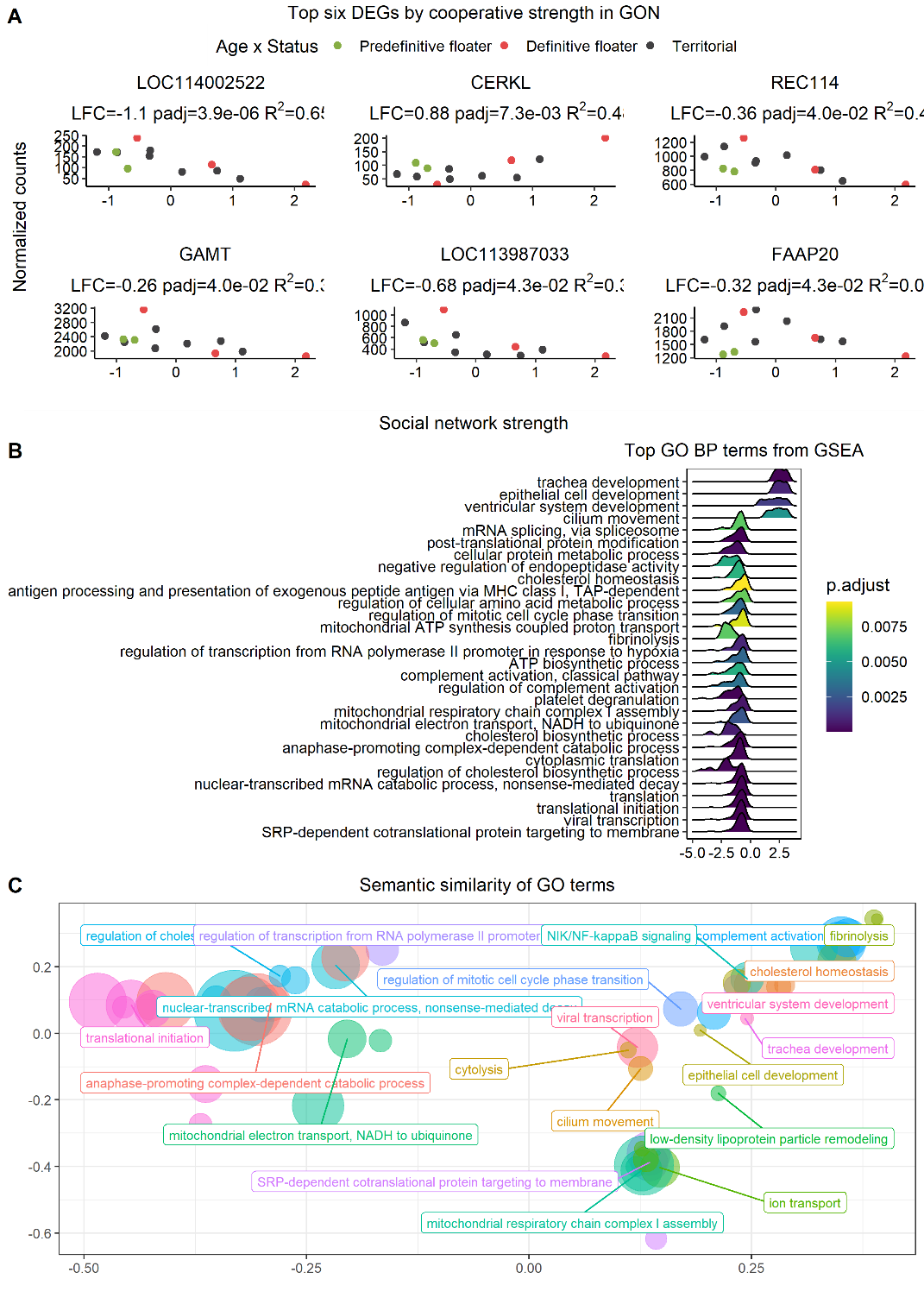

###### **Figure S3.32: Social network strength associated DEGs in the testes (GON).**

A) The six genes with the lowest p-values for difference in gene expression in individuals with high or low cooperative tendencies (social network strength). B) Ridge plot showing the density of GO terms along a directional, log-transformed of gene p-values, therefore genes with low p-values are on the left and right edges of the plot. C) Bubble plot of semantically clustered GO terms where colors indicate the clustered GO terms, with the parent term labelled. The three-dimensional distance between points represents the semantic distance between terms, and the bubble size indicates enrichment score.

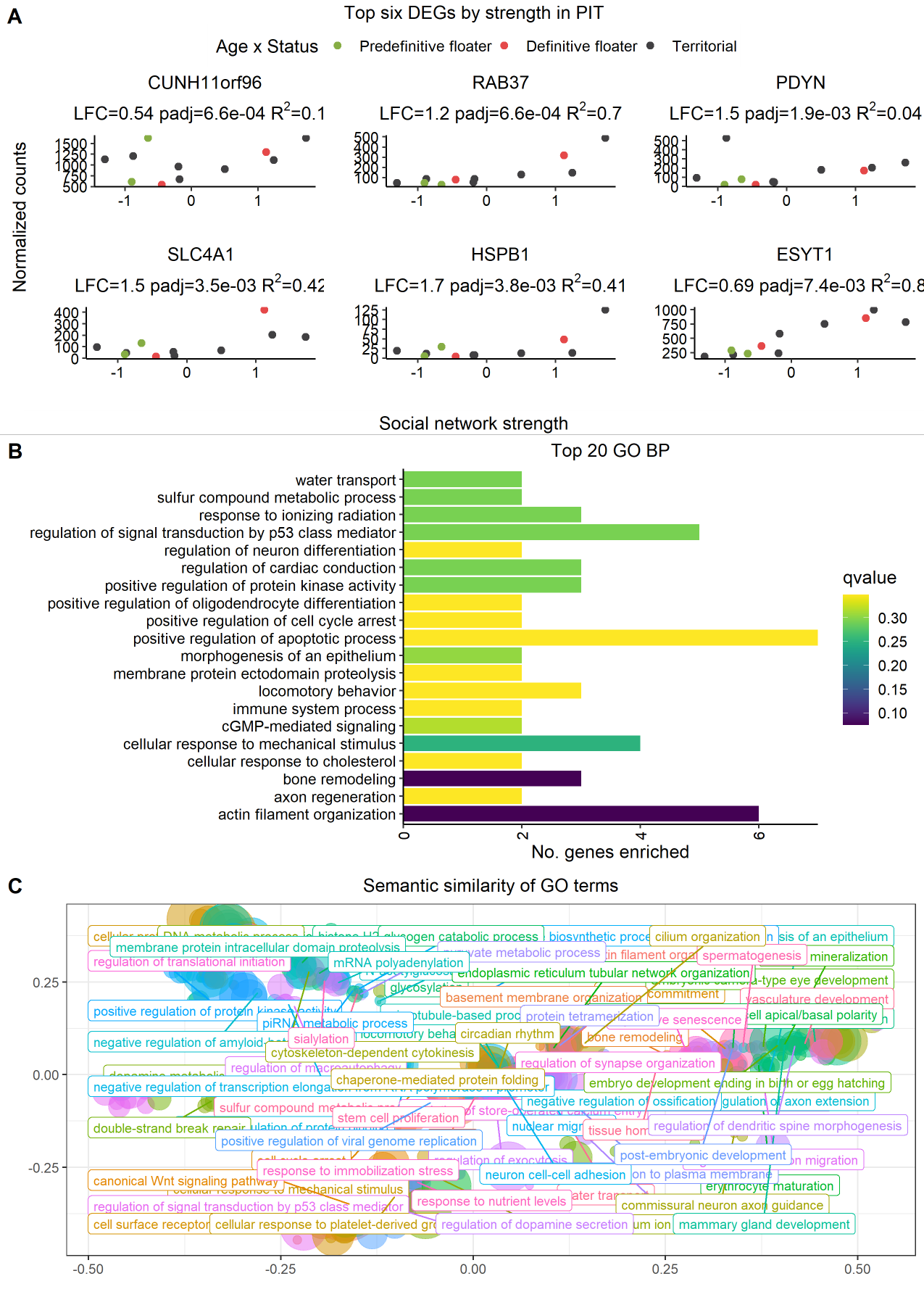

###### **Figure S3.33: Social network strength associated DEGs in the pituitary (PIT).**

Explanations as in Figure S3.32.

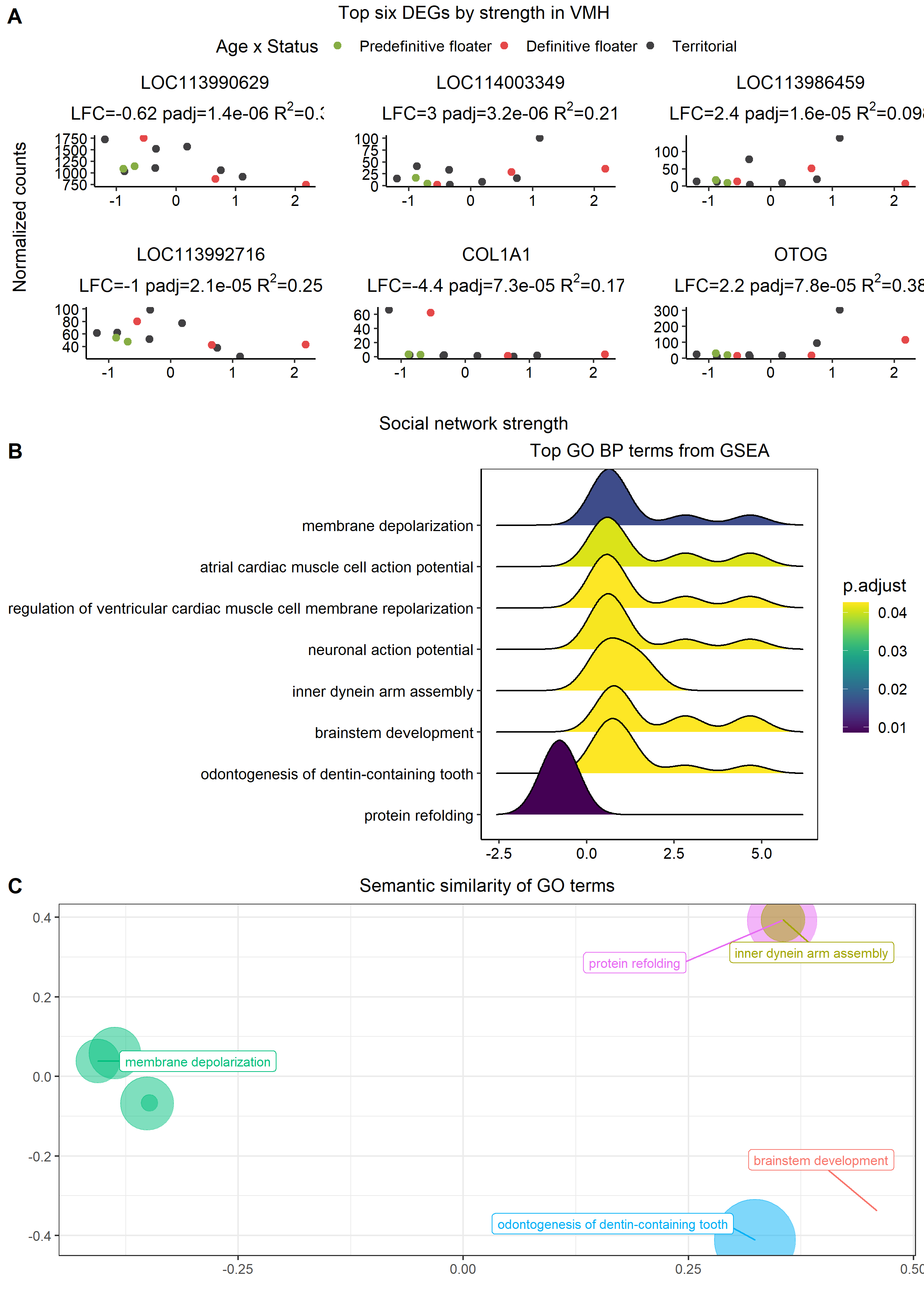

###### **Figure S3.34: Social network strength associated DEGs in the ventromedial hypothalamus (VMH).**

Explanations as in Figure S3.32.

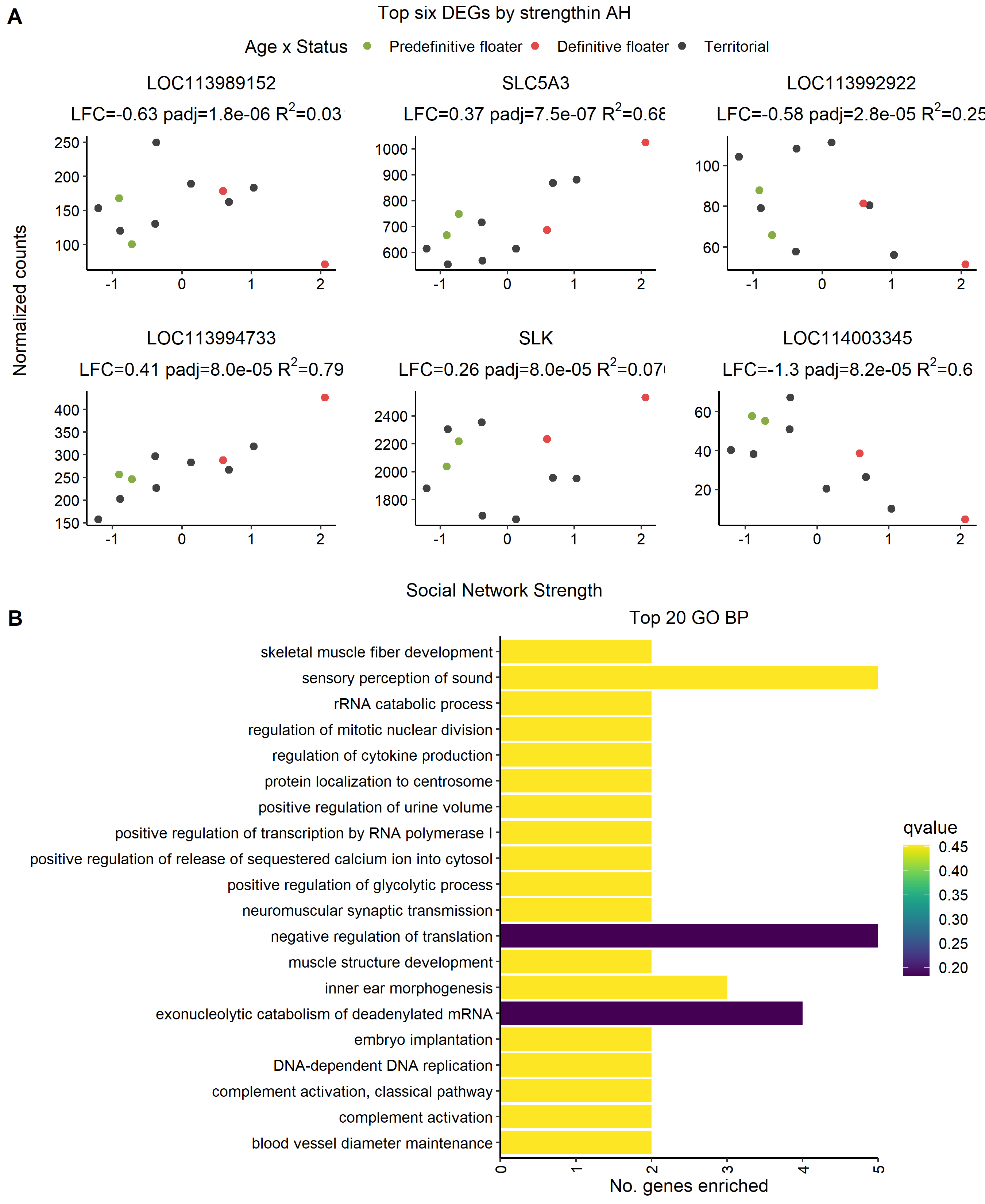

###### **Figure S3.35: Social network strength associated DEGs in the anterior hypothalamus (AH).**

Explanations as in Figure S3.32.

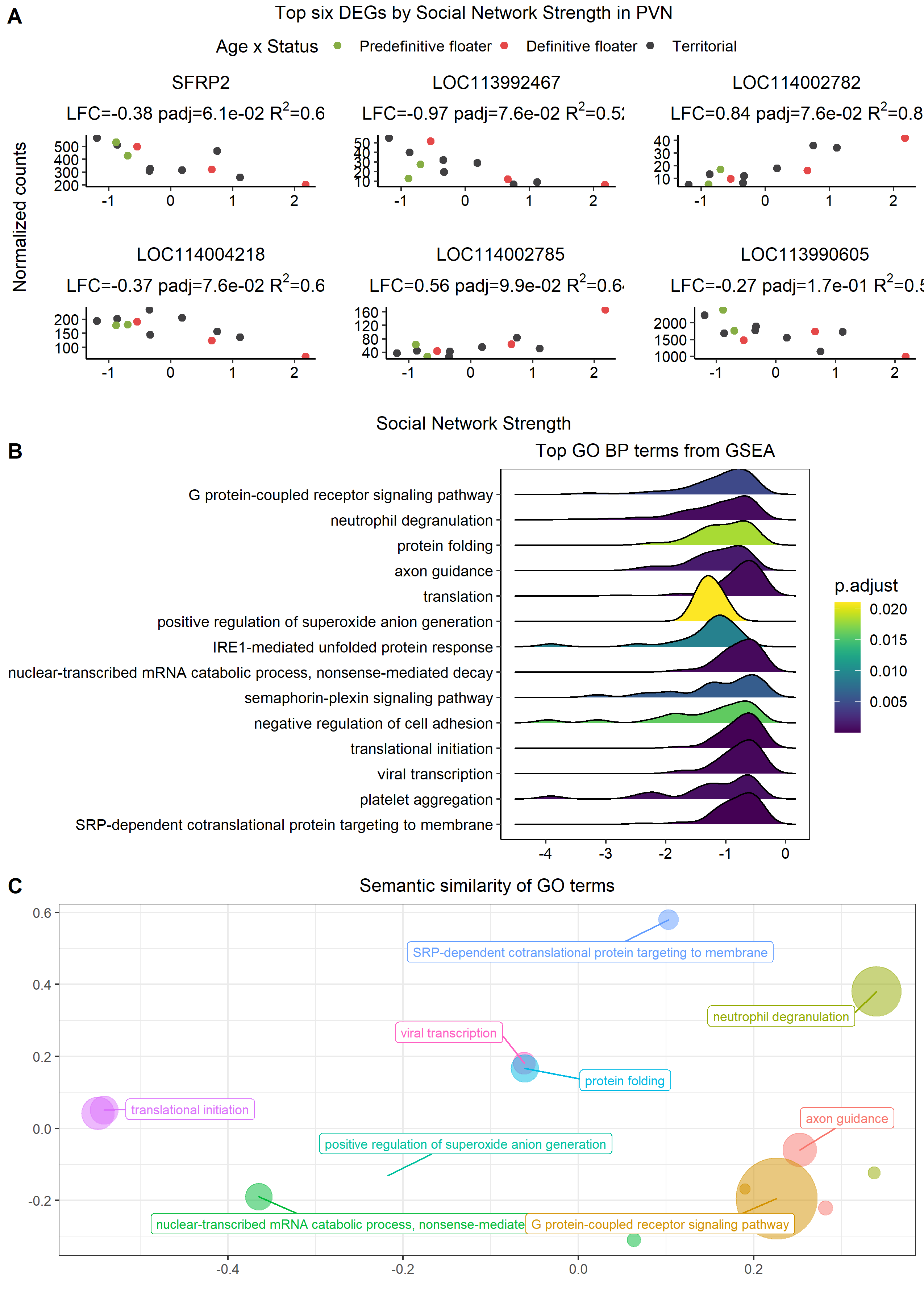

###### **Figure S3.36: Social network strength associated DEGs in the paraventricular nucleus of the hypothalamus (PVN).**

Explanations as in Figure S3.32.

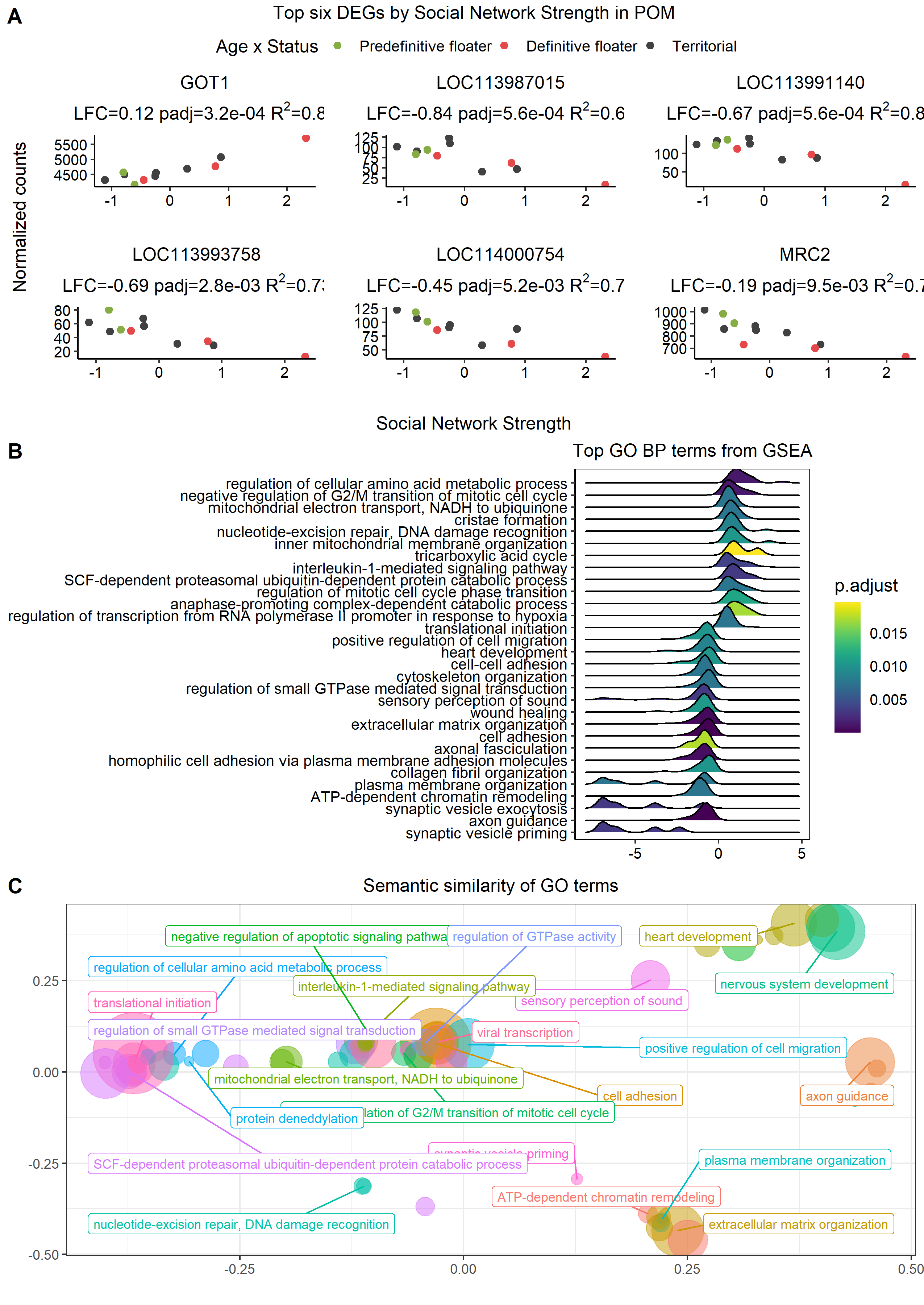

###### **Figure S3.37: Social network strength associated DEGs in the medial preoptic nucleus (POM).**

Explanations as in Figure S3.32.

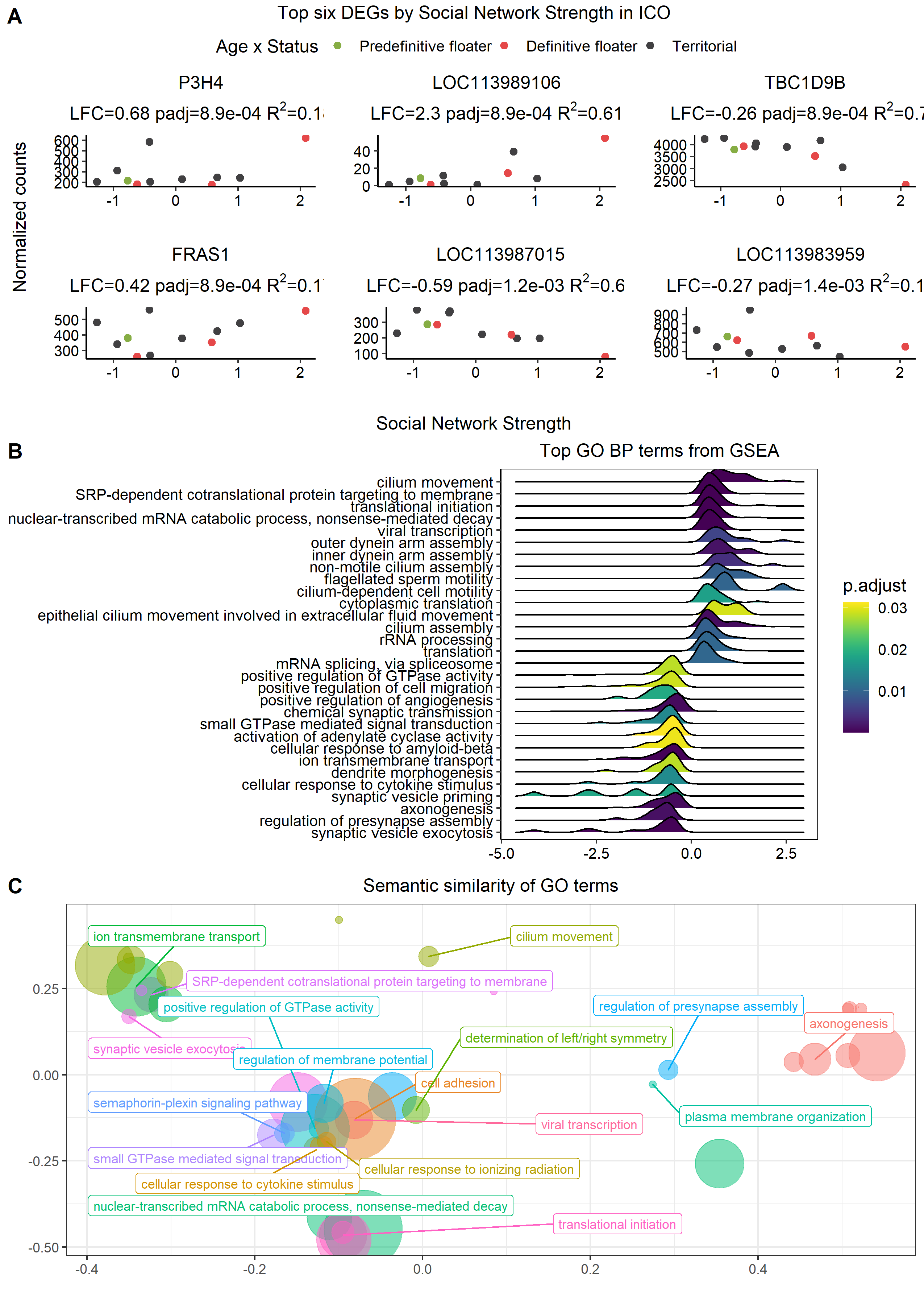

###### **Figure S3.38: Social network strength associated DEGs in the intercollicular nucleus (ICo).**

Explanations as in Figure S3.32.

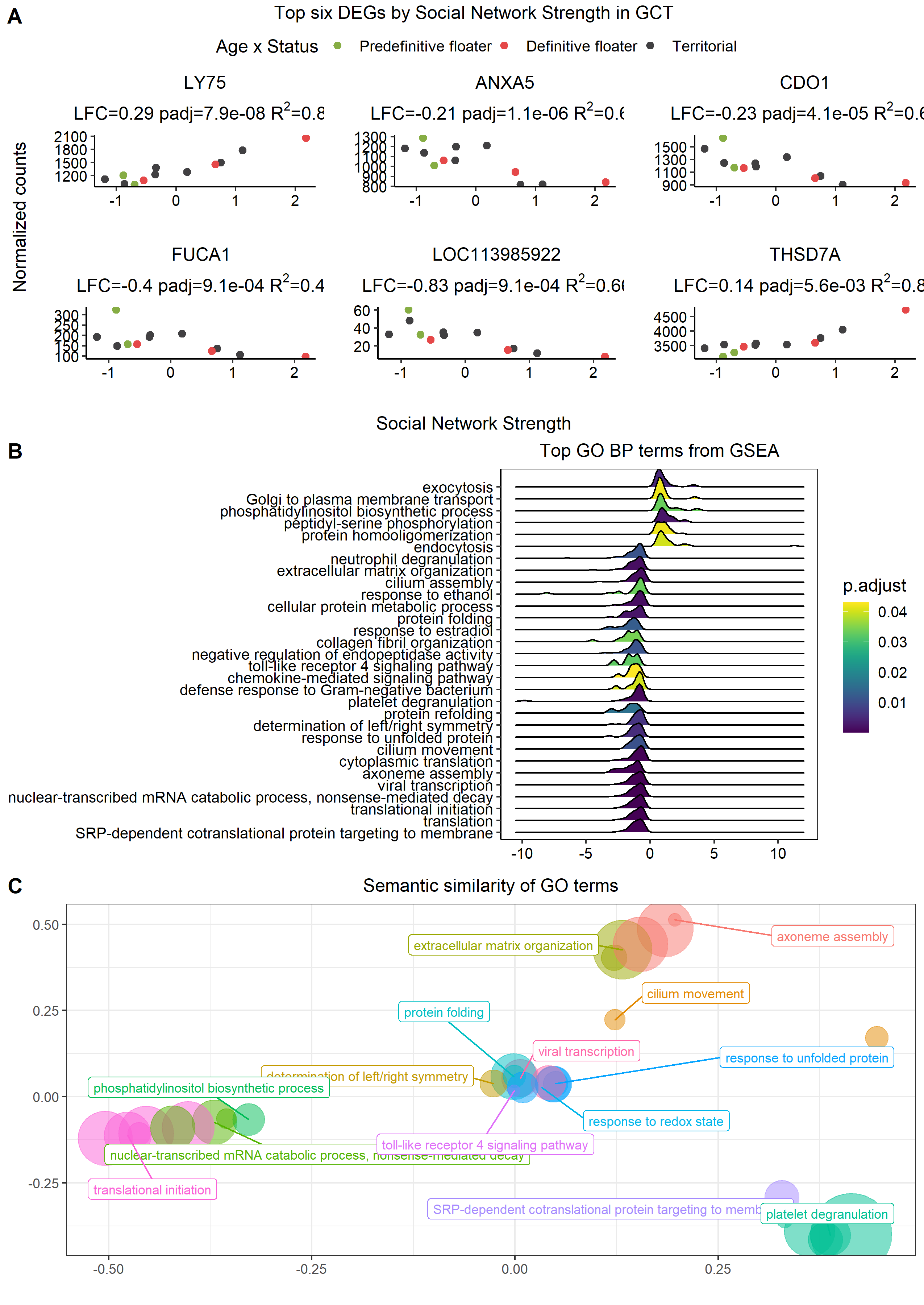

###### **Figure S3.39: Social network strength associated DEGs in the midbrain central grey (GCt).**

Explanations as in Figure S3.32.

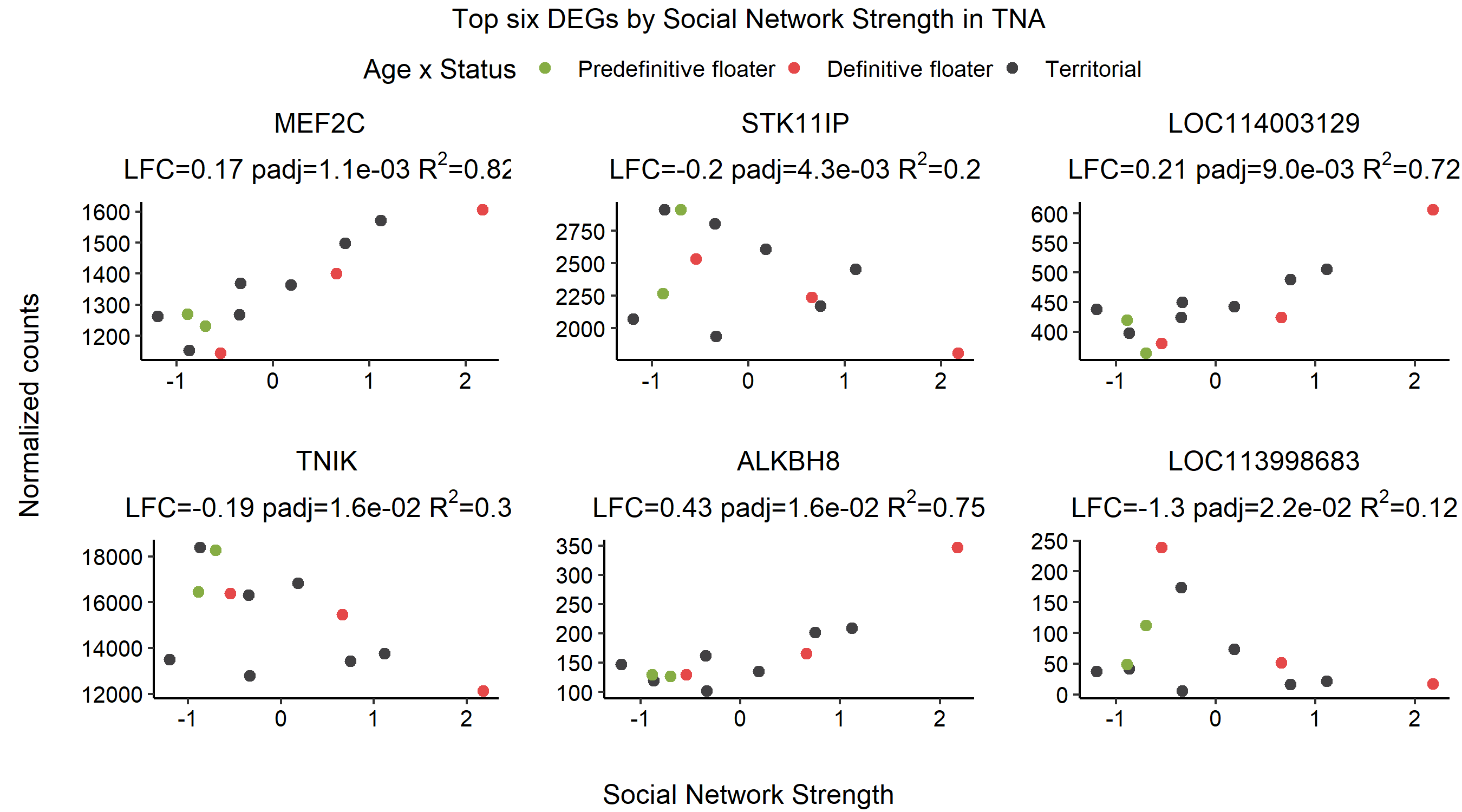

###### **Figure S3.40: Social network strength associated DEGs in the nucleus taenia (TnA).**

Explanations as in Figure S3.32. There were no enriched GO categories.

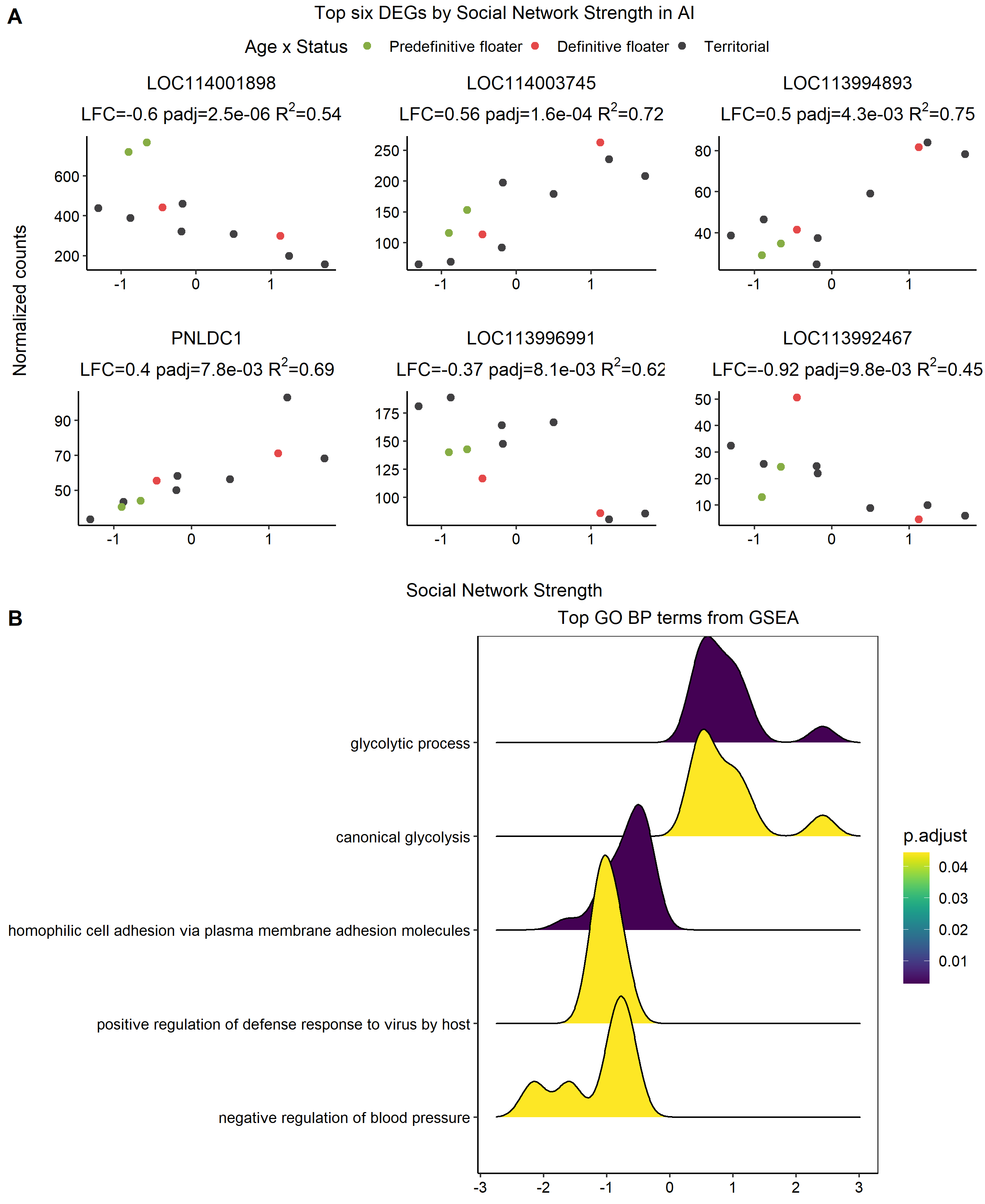

###### **Figure S3.41: Social network strength associated DEGs in the arcopallium intermedium (AI).**

Explanations as in Figure S3.32.

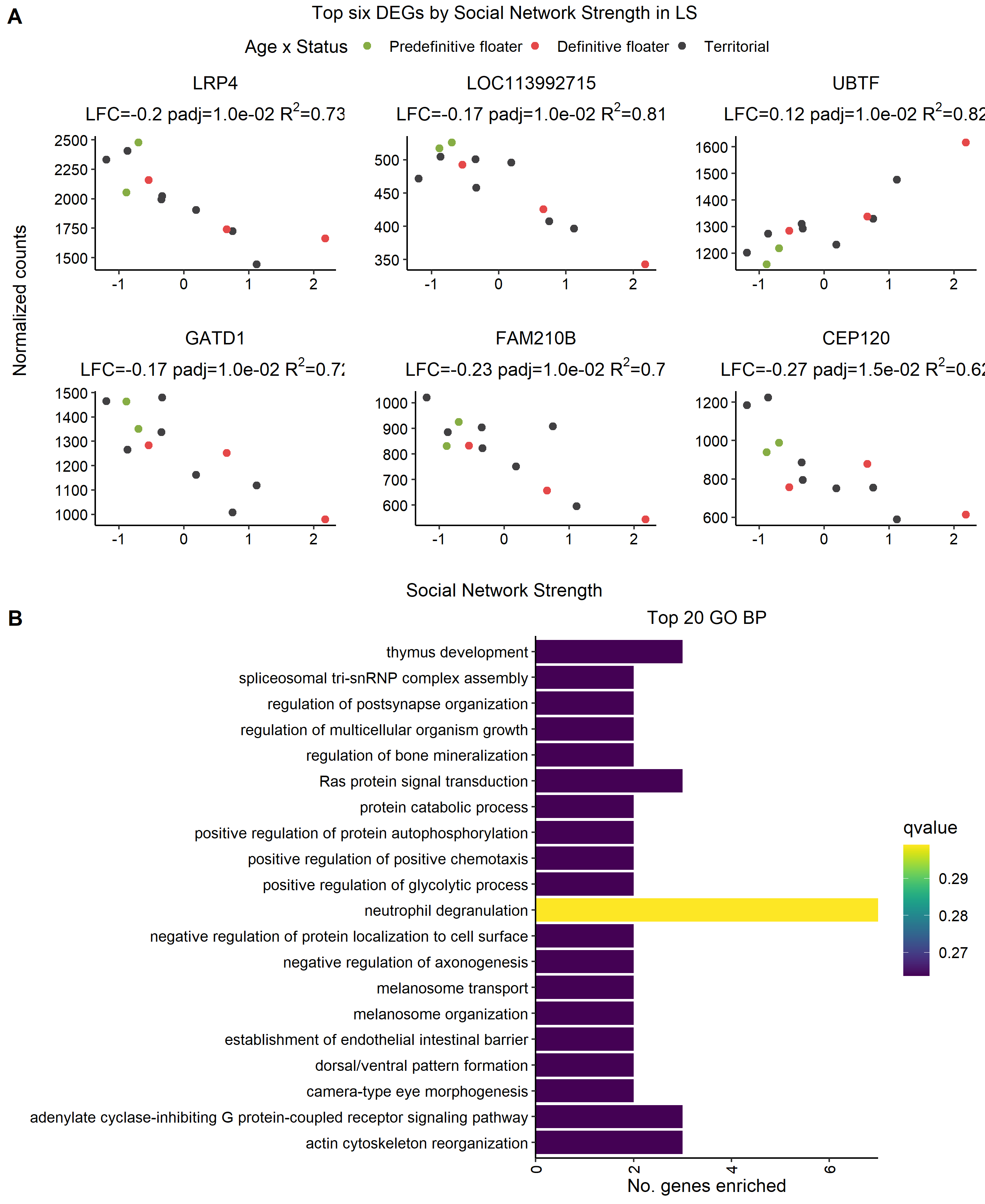

###### **Figure S3.42: Social network strength associated DEGs in the lateral septum (LS).**

Explanations as in Figure S3.32.

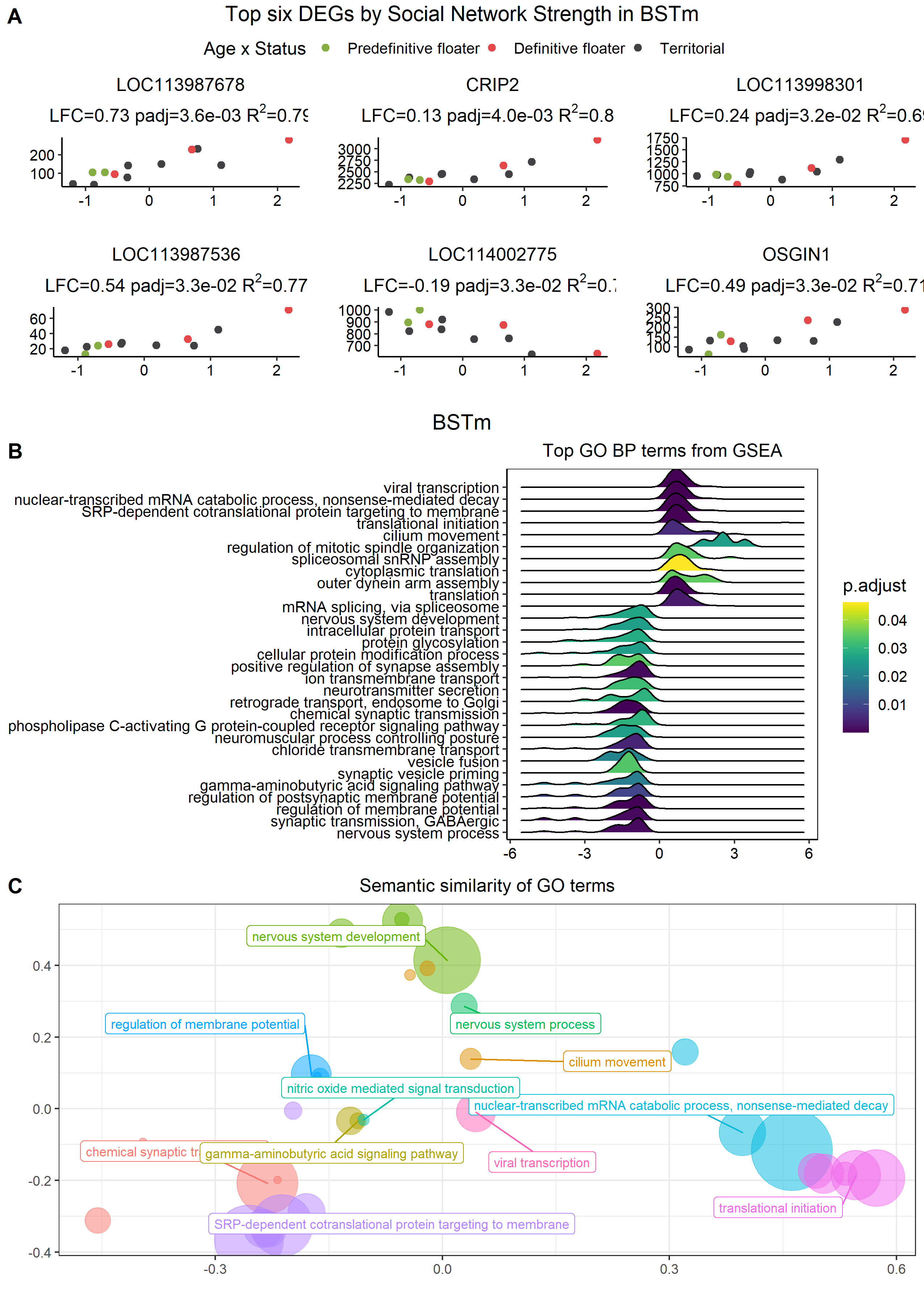

###### **Figure S3.43: Social network strength associated DEGs in the bed nucleus of the stria terminalis (BSTm).**

Explanations as in Figure S3.30.

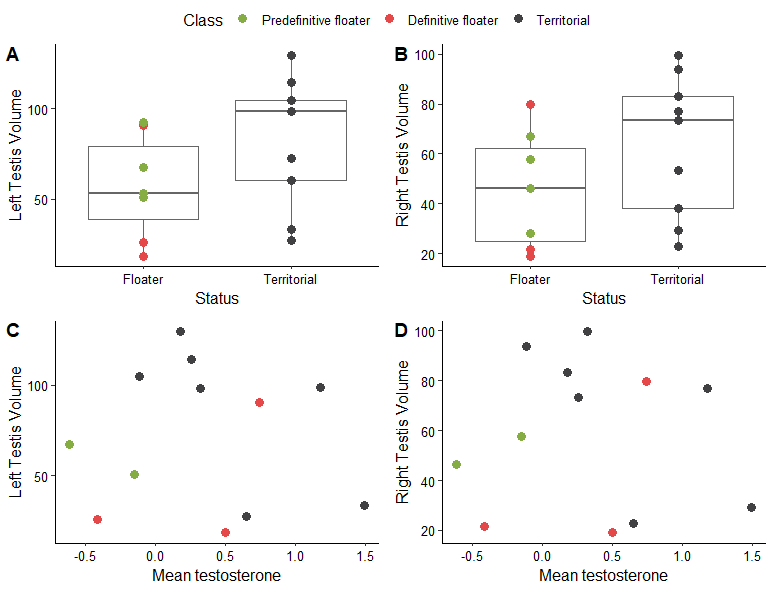

###### **Figure S3.44: Testes size in male wire-tailed manakins.**

Comparing testis size (volume mm^3^) against social status in the left A) and right B) testis for the total sample of 16 males, and the testis volume for left C) and right D) for males with repeated testosterone measures. Colors represent social status and plumage type.

###### Table 3.1 Results from linear regression on the interactive effect of mean T and status on testis size.

Where size is volume in mm^3^ and the sample size is 12 males with repeated testosterone measures.

| Testis | Variable | β | t | P (t) | F | P (F) |
| --- | --- | --- | --- | --- | --- | --- |
| Left | Status | 60.84 | 2.53 | 0.04* | 3.28 | 0.11 |
|  | Mean testosterone | 9.88 | 0.34 | 0.74 | 1.41 | 0.27 |
|  | Status * mean T | -53.42 | -1.422 | 0.19 | 2.02 | 1.19 |
| Right | Status | 42.66 | 3.77 | 0.05 | 2.31 | 0.17 |
|  | Mean testosterone | 14.32 | 2.27 | 0.54 | 0.94 | 0.36 |
|  | Status * mean T | -47.55 | -1.62 | 0.14 | 2.63 | 0.14 |
| Mean | Status | 51.75 | 2.50 | 0.04* | 3.04 | 0.12 |
|  | Mean testosterone | 12.10 | 0.49 | 0.64 | 1.27 | 0.29 |
|  | Status * mean T | -50.48 | -1.56 | 0.16 | 2.44 | 0.16 |

### 4. Candidate gene analysis

#### Methods

Using the results from differential expression analysis (Section 3) we also explored the expression of 18 candidate genes identified *a priori* for their likely involvement in androgen moderated social behavior in each tissue (Table S1.3). The candidate genes are broadly divided into sex hormone metabolism and sex hormone receptors (Figure S4.1), and neuropeptides and their receptors (Figure S4.2).

Additionally, we identified candidate genes with potential status-specific modulation by mean testosterone by using the general model from above with a status * mean T interaction in *DESeq2* (Love *et al.* 2014). All candidate genes with an uncorrected p-value <0.05 in any analysis were extracted and plotted for further visual inspection. We filtered this list of “significant” genes by removing relationships with highly influential observations, or for continuous predictors with R^2^ ≤ 0.2.

Some candidate genes were significantly correlated to more than one of our traits of interest, or were significant in the interaction model. As such, all these contenders for significant candidate genes were further were further validated using a stepdown-like procedure using AIC. We compared the AIC of the following models:

1. gene expression ~ [Batch] + [Year] + NULL
2. gene expression ~ [Batch] + [Year] + status
3. gene expression ~ [Batch] + [Year] + mean_T
4. gene expression ~ [Batch] + [Year] + status x mean T

If the gene was also associated with social network strength, then it was included in the larger model set described above. For consistency, we included Batch and Year effects when they were included in the original *DESeq2* models for that tissue. We accepted the model with the lowest AIC, only if ΔAIC≥2 compared to the null model, and from other models in the testing set. Accepted models are annotated with an “a”. If there were two models with similar AIC but both better than the null model, we coded this as indeterminate or “?”. We report both the raw p-value and the *DESeq2* adjusted p-value that is corrected for the number of genes expressed in the tissue. We also report our final list of candidate gene relationships after the above exclusion process.

#### Results

This section will include the raw plotted results for each candidate gene, and include the full set of AIC tables. The detailed Rmarkdown is at this [link](https://periperipatus.github.io/PIFI_brain_transcriptome/05_candidate_gene_analysis.html). For the final list of genes and relationships after the filtering process see Figure 2C & 3B.

Two candidate genes, PRLR and AVPR1A were putatively associated with mean testosterone phenotype in the testes (GON) (Figure S4.3). VIPR1 was associated with social network strength but was discarded owing to low R^2^. VIPR2 appeared to be related to mean testosterone in a status-specific manner (Figure S4.4A), but no model was a substantially better fit to the data than the null model (Figure S4.4B), therefore we conclude it is unlikely there is a status-specific relationship in the testes data.

Seven candidate genes were associated with interest variables in the pituitary (PIT) (Figure S4.5). However, AVP and OXT were all spurious relationships with unusual count distributions, and so we excluded these from the final list, and ESR1 ~ Status was excluded on the basis of a single influential observation. Note that the results for social network strength exclude individual PFT3, as it was highly influential on a large proportion of genes (see Supplement 3 methods, and [GitHub](https://periperipatus.github.io/PIFI_brain_transcriptome/02_DESeq2.html#541_Analysis)). PRL showed a strong correlation with (LFC=-0.4, p=0.0001,q=0.02, R^2^=0.56) and without this potential outlier (Figure S4.5, Figure 3C). AVPR1B showed a significant relationship before FDR correction when PFT3 was removed (Figure S4.5), but this relationship was not significant with PFT3 included (LFC=0.2, p=0.08, q=0.4, R^2^=0.098). We kept in the final list AVPR1B as all results relating to social network strength in the PIT reported exclude PFT3. VIPR2 was associated with both mean testosterone and social network strength. PFT3 had little influence on the social network strength model as the original model also showed a correlation between VIPR2 and social network strength (LFC=-0.4, p=0.001, q=0.06, R^2^=0.48). When comparing models that included PFT3, social network strength had the higher R^2^ and ΔAIC found this model to be the best fit. Therefore we excluded the VIPR2 ~ mean T result (result not shown). Further, AVPR1A showed status-specific relationship with mean testosterone phenotype in the DESeq2 interaction analysis (Figure S4.6A), and the model AIC was at the cut-off of ΔAIC (Figure S4.6B) and so was kept in the final list.

There were six candidate genes putatively associated with variables of interest in the ventromedial hypothalamus (VMH) (Figure S4.7). Again, like in the PIT, there appeared to be unusual count distributions and low R^2^ in GNRH1, OXT, and AVP, and these results were removed from the final list. AR showed a weak negative relationship with social network strength, but was excluded on the basis of its R^2^. ESR2 showed decreased expression in territorial males. AVPR2 showed a negative relationship with mean T, but the counts involved are low (<30). PGR, AVPR1A, and GNRH1 were identified as putatively status-specific in expression, but all three models were not a better fit for the data, nor were the visualizations compelling, so these results are only shown in the [GitHub](https://periperipatus.github.io/PIFI_brain_transcriptome/05_candidate_gene_analysis.html#41_Status_specific_gene_expression_in_the_VMH).

In the paraventricular nucleus of the hypothalamus (PVN), there were six genes associated with variables of interest (Figure S4.8). However, AVP again did not meet the minimum requirements for R^2^ and was excluded from the final list. ESR2 showed decreased expression in territorial males, while VIPR1 expression increased in territorial males. AVPR2 showed a weak positive relationship with social network strength, but also associated with low read counts (<40). Oxytocin (OXT) showed a weak negative correlation in association with social network strength. Although Androgen receptor (AR) showed weak positive correlation with mean T and strength, AIC preferred the interaction model identified by *DESeq2* (Figure S4.9). Therefore, we discarded the AR ~ mean T and strength relationships in favor of the interaction.

There were five candidate genes associated with social status and mean testosterone in the medial preoptic nucleus (POM) (Figure S4.10). Aromatase (CYP19A1) and GNRH1 showed weak positive relationships with mean testosterone. Progesterone receptor (PGR) showed increased expression in territorial males. The candidate gene VIP was also identified in the *DESeq2* interaction models (Figure S4.11A), but this model fit was not better than the null model (Figure S4.11B), and this was excluded from the final list. Both AR and PRLR were associated with both mean T and Status, suggesting a potential interaction between these variables (Figure S4.11C,E). We compared model fit among these models but found that the model including mean testosterone phenotype only had the lowest AIC, and was a substantially better fit than the null models (Figure S4.11D,F). Despite similar AIC (ΔAIC<2), we excluded the interaction model from further consideration as p-values for the interaction model were not significant (Figure S4.11F).

There were no candidate genes identified as related to our interest variables in the anterior hypothalamus (AH).

In the midbrain, there was strong evidence of the involvement of the VIP system in regulating cooperation. VIP showed a negative relationship with mean testosterone phenotype, while VIPR2 was negatively associated with social network strength in the dorsomedial intercollicular nucleus (ICo) (Figure S4.12). Further genes (AVPR1A, AVP, OXTR) were putatively regulated in a status-specific manner (Figure S4.13 top row). AVPR1A (Figure S4.13A) was excluded because it was not substantially better than the null model. OXTR had an unusual count distribution, with many individuals with very low counts (Figure S4.13C), and so was excluded on this basis.

The midbrain central grey (GCt) showed five putative differentially expressed genes (Figure S4.14) and may be a hotbed for status-specific regulation (Figure S4.15). AR ~ Strength was excluded on the basis of its low R^2^, and OXT was excluded on the basis of a highly influential observation. Low counts of CYP19A1 showed a negative relationship with mean T, while PGR and VIP were associated with multiple traits. AIC analysis strongly favoured the interaction model with VIP, which decreased in expression in floater males with high testosterone (Figure S4.15AB). A similar relationship was observed with PGR (Figure S4.15CD). ESR1 showed increasing expression with mean testosterone phenotype in territorial individuals (Figure S4.15EF), and ESR1 has been previously demonstrated to be an effective receptor even with low transcript counts in the white-throated sparrow nucleus taenia (Horton *et al.* 2014). Finally, PRLR showed a status-specific relationship similar to ESR1 (Figure S4.15GH).

The nucleus taenia (TnA) showed that five candidate genes were potentially associated with our traits of interest (Figure S4.16), AR, PGR, AVP, and AVPR1A were all excluded from downstream analysis owing to low R^2^ and influential observations. The interaction analysis in *DESeq2* identified four candidate genes with potential status-specific gene expression (Figure S4.17), but only PRLR and VIPR1 showed improvement over the null models.

There were three genes associated with interest variables in the arcopallium intermedium (AI) (Figure S4.18). VIP was associated with both social network strength and mean testosterone phenotype, model comparisons revealed that both were substantially better from the null model, and mean testosterone showed only a minor (ΔAIC<2) improvement in model performance (Figure S4.19CD) and so VIP was retained on lists for status and social network strength (Figure 2 & 3). The interaction analysis additionally suggested AVP, VIPR1, and AVPR1A as possible candidates for status-specific gene regulation in the AI. However, the AVP model was substantially worse than the null (ΔAIC>2) (Figure S4.19). VIPR1 and AVPR1 were excluded and not presented because the models did not improve on the null and visualization was not compelling, but can these results be seen on [GitHub](https://periperipatus.github.io/PIFI_brain_transcriptome/05_candidate_gene_analysis.html).

There were three genes that were putatively associated with our traits of interest in the lateral septum (LS) (Figure S4.20). However, after filtering the genes for outliers and low R^2^, only SRD5A2 remained a convincing candidate for significantly increased expression in territorial males. Interaction analysis revealed AR as a possible candidate for status-specific expression in the LS (Figure S4.21A), but model fit was not a substantial improvement on the null model (Figure S4.21B).

There were also three genes putatively associated with traits in the bed nucleus of the stria terminalis (BSTm), but ESR2 was filtered from downstream analysis owing to low R^2^ (Figure S4.22). Remaining is a plausible increase in expression of AR in territorial males (Figure 2F; Figure S4.22), and a decrease in PGR expression with highly cooperative males (Figure S4.22). Interaction analysis revealed both ESR1 and ESR2 as possible candidates for status-specific gene expression in the BSTm (Figure S4.23A,C). Only ESR2 was a plausible candidate for status-specific expression (Figure S4.23B,D).

These results reveal an interesting pattern suggesting certain nuclei have a density of sex steroid-related genes possibly related to status-specific regulation of cooperative behavior. While these results provide compelling initial evidence, more targeted studies are needed to ascertain their roles in modulation of male-male cooperative behavior.

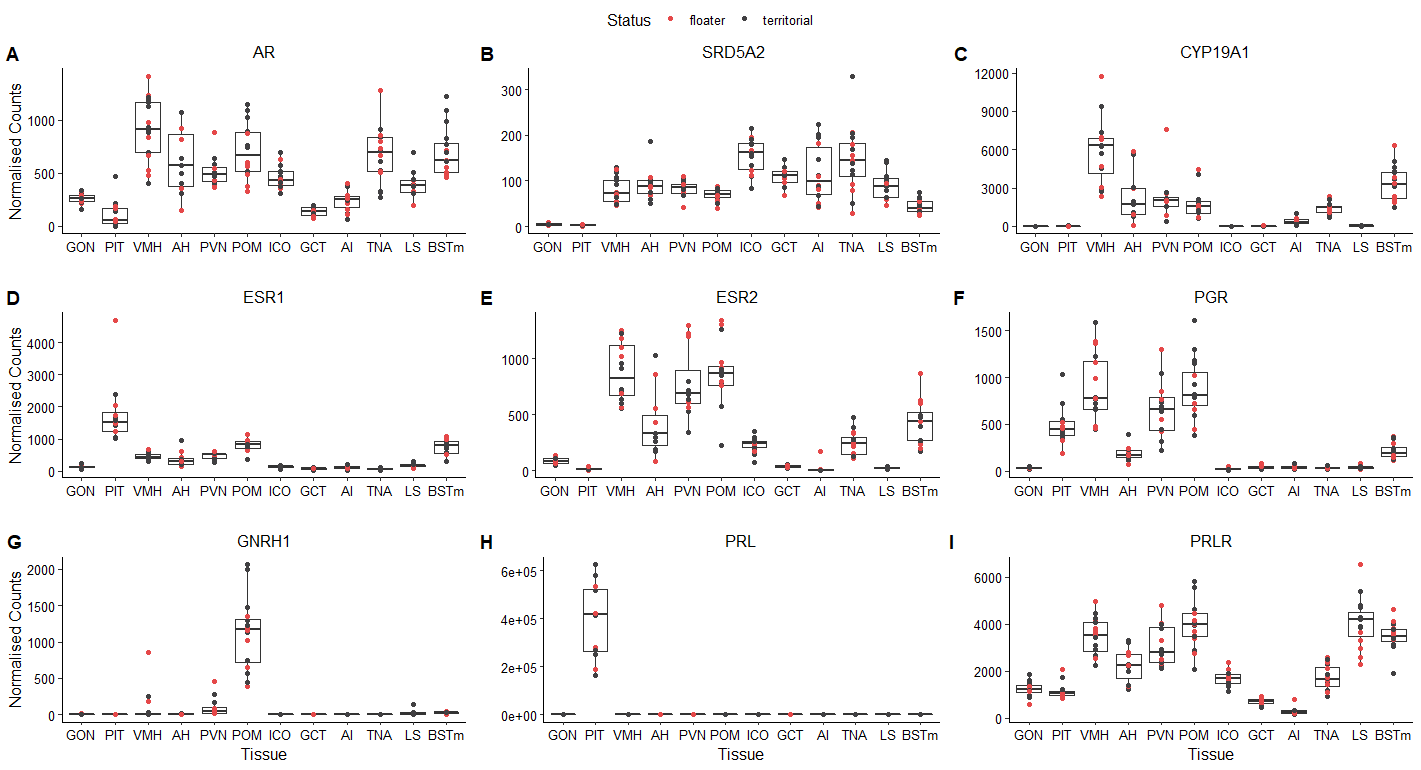

###### **Figure S4.1: Tissue-specific expression of sex steroid related candidate genes**.

Normalization method is the Median of Ratios from *DESeq2* (Love *et al.* 2014). Points are color coded according to social status for visualization only – the full results are presented later in this section.

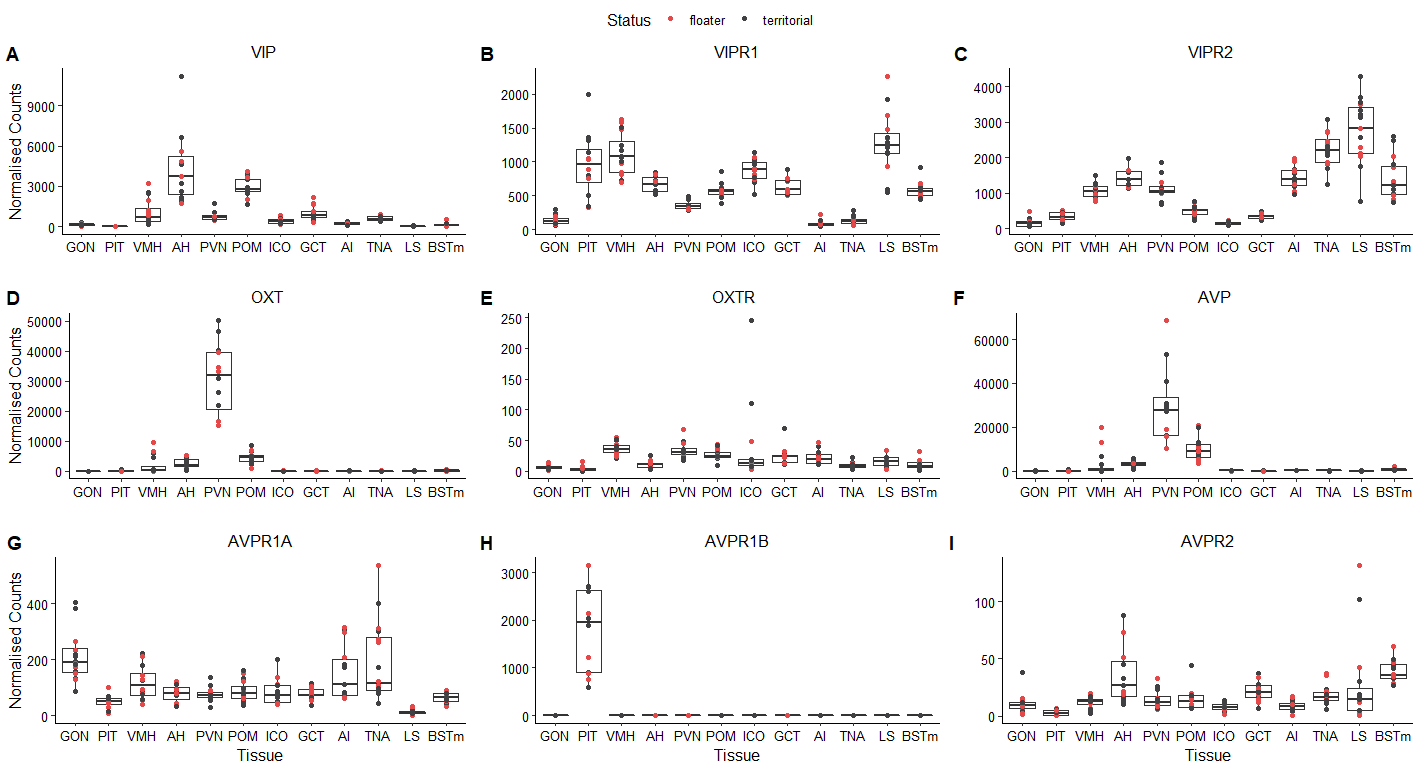

###### **Figure S4.2: Tissue-specific expression of candidate neuropeptides.**

Normalization method is the Median of Ratios from *DESeq2* (Love *et al.* 2014). Points are color coded according to social status for visualization only – the full results are presented later in this section

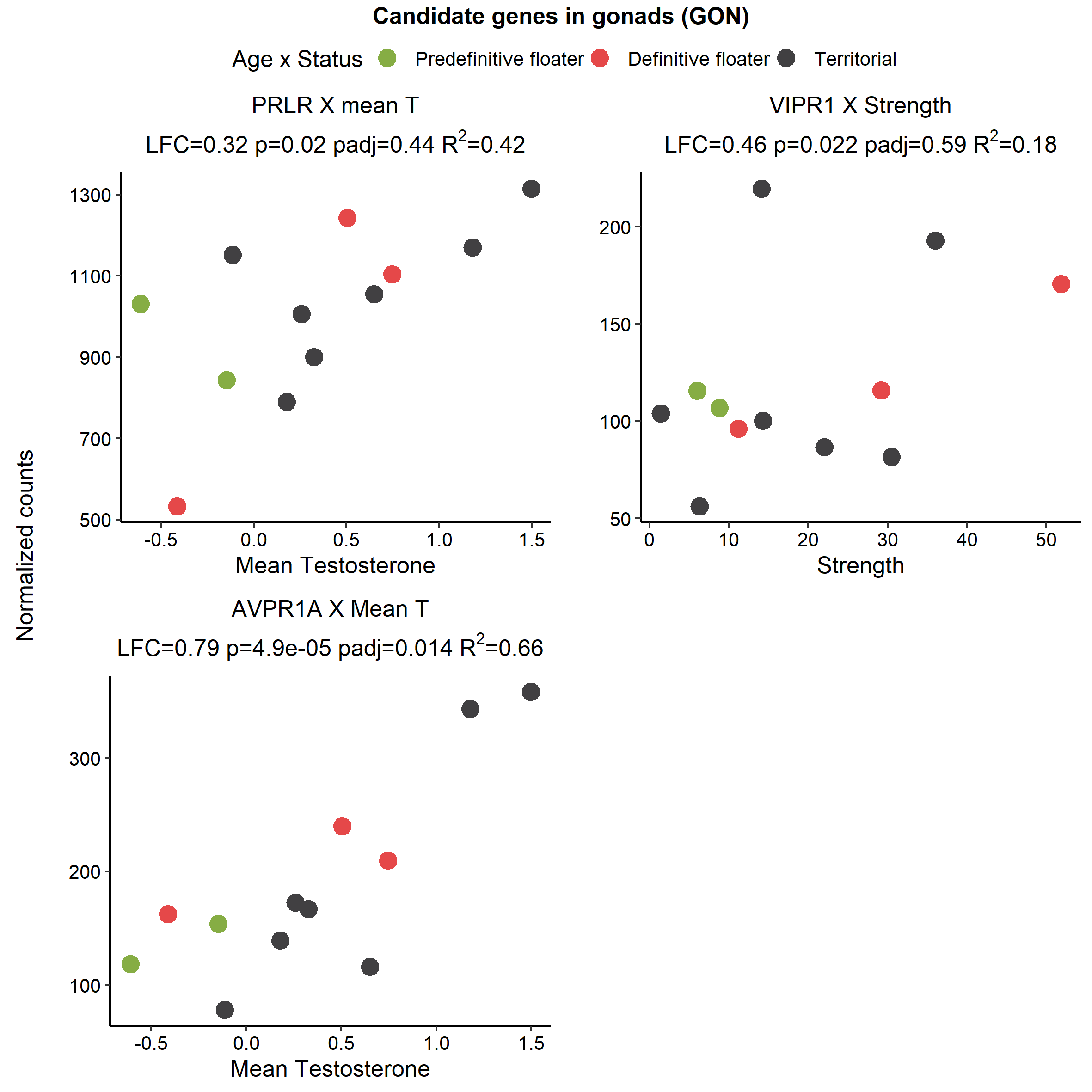

###### Figure S4.3: Candidate genes differentially expressed in the testes (GON).

Each panel represents a candidate gene by trait comparison relevant to that tissue, with uncorrected p-values <0.05. The particular trait the gene expression is related to is in the panel heading and the x-axis label. Points are color coded by social status and plumage state for illustration purposes only.

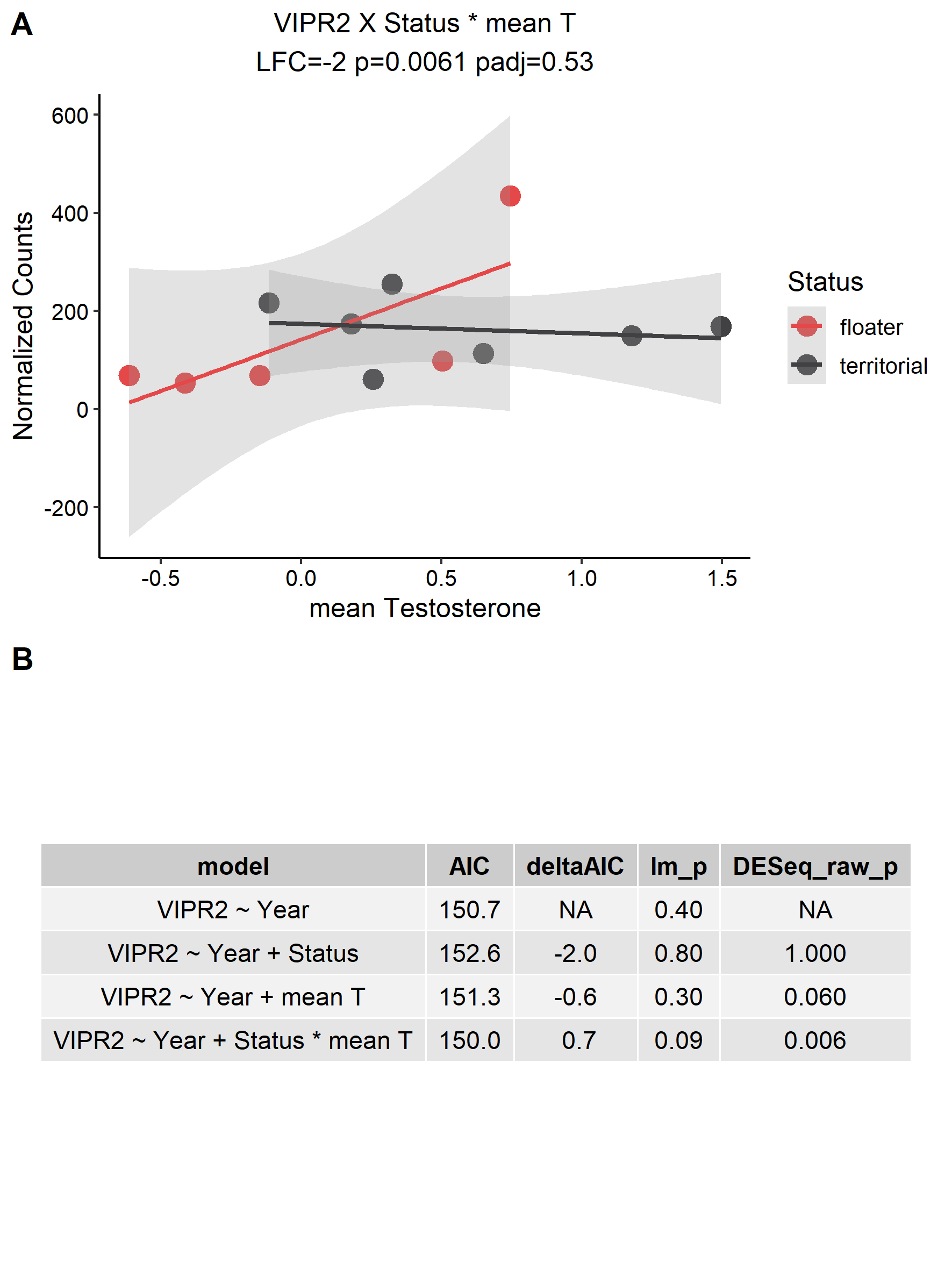

###### Figure S4.4: Possible interaction effects in the testes (GON).

DESeq2 analysis of the interaction between mean testosterone phenotype (mean T) and status revealed a putative interaction in VIPR2 in the testes A). This relationship was tested using AIC amongst a series of models and a null model (that incorporates any necessary nuisance variables), to determine whether this interaction relationship was any better than the simpler or null models B). Based on this result, we did not accept that there was a convincing relationship between VIPR2 expression and status/mean testosterone in the testes.

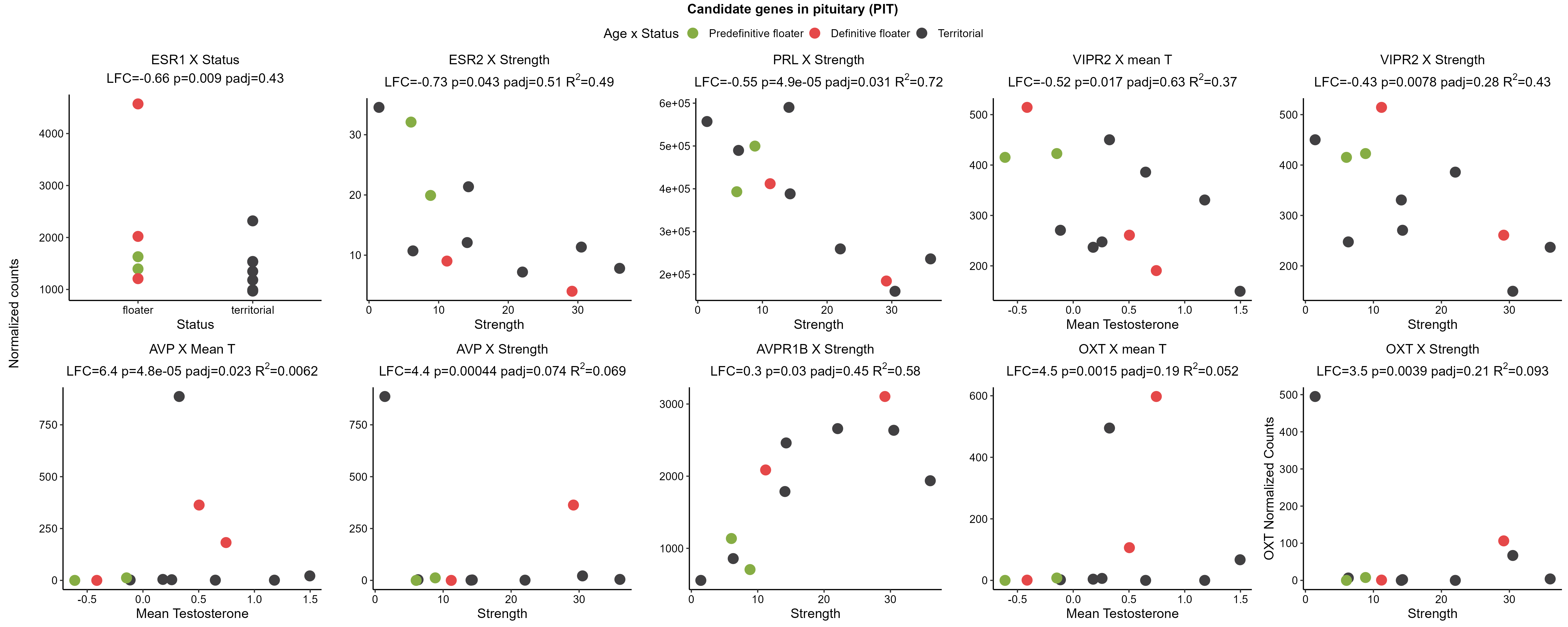

###### Figure S4.5: Candidate genes differentially expressed in the pituitary (PIT).

Explanations as in Figure S4.3. ESR1, AVP and OXT genes were disregarded from further analysis owing to satisfying both the “influential observation” and/or low R^2^ criteria. AIC analysis preferred VIPR2 ~ Strength and so the relationship with mean T was excluded from the final list. Note that PFT3 has been excluded from all models assessing social network strength (Supplement 3).

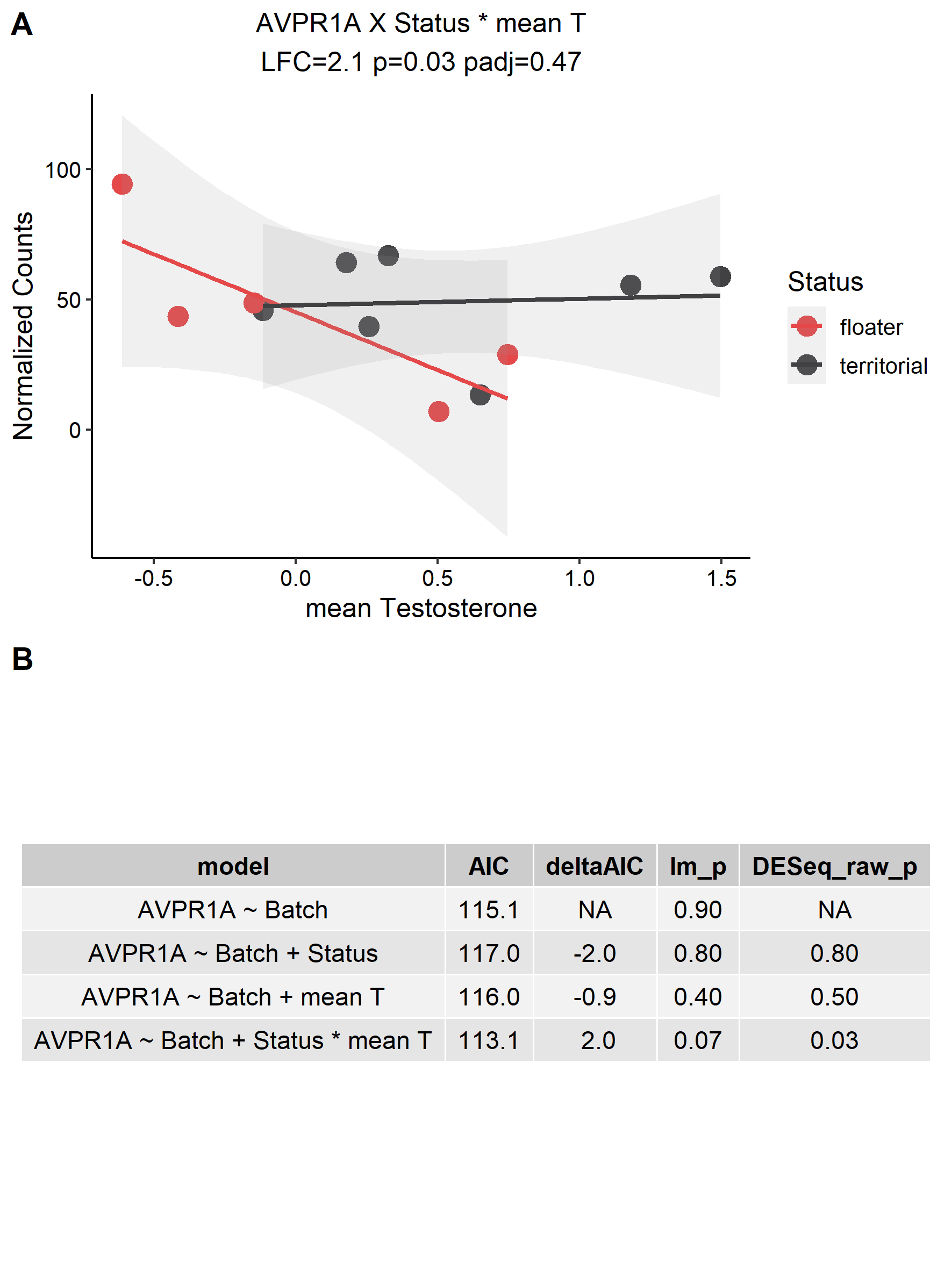

###### **Figure S4.6: Possible interaction effects in the pituitary (PIT).**

Explanations as in Figure S4.4. The interaction model was accepted.

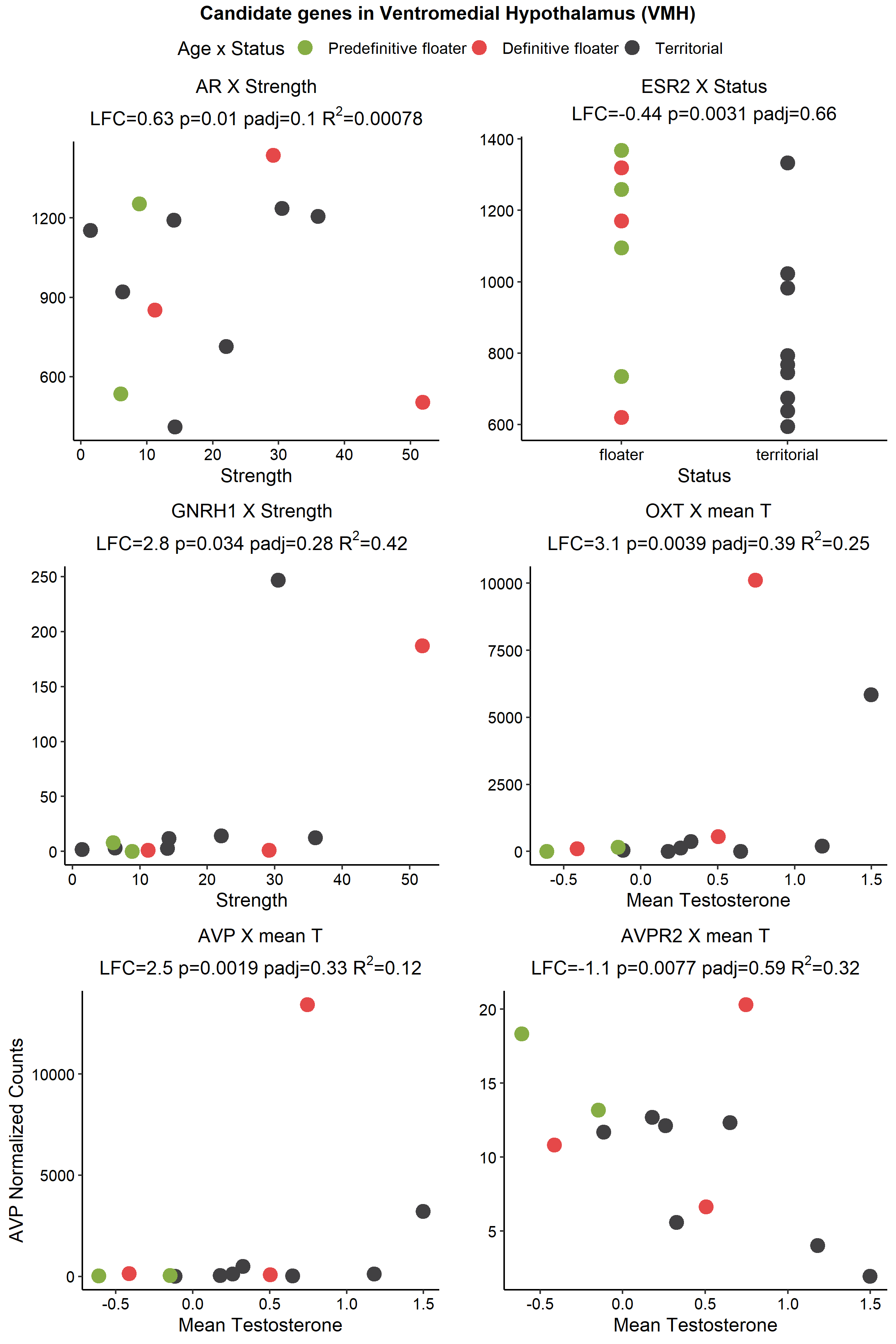

###### Figure S4.7: Candidate genes differentially expressed in the ventromedial hypothalamus (VMH).

Explanation as in Figure S4.3. GNRH1, OXT, and AVP were excluded from further consideration based on influential observations. AR was excluded on the basis of low R^2^.

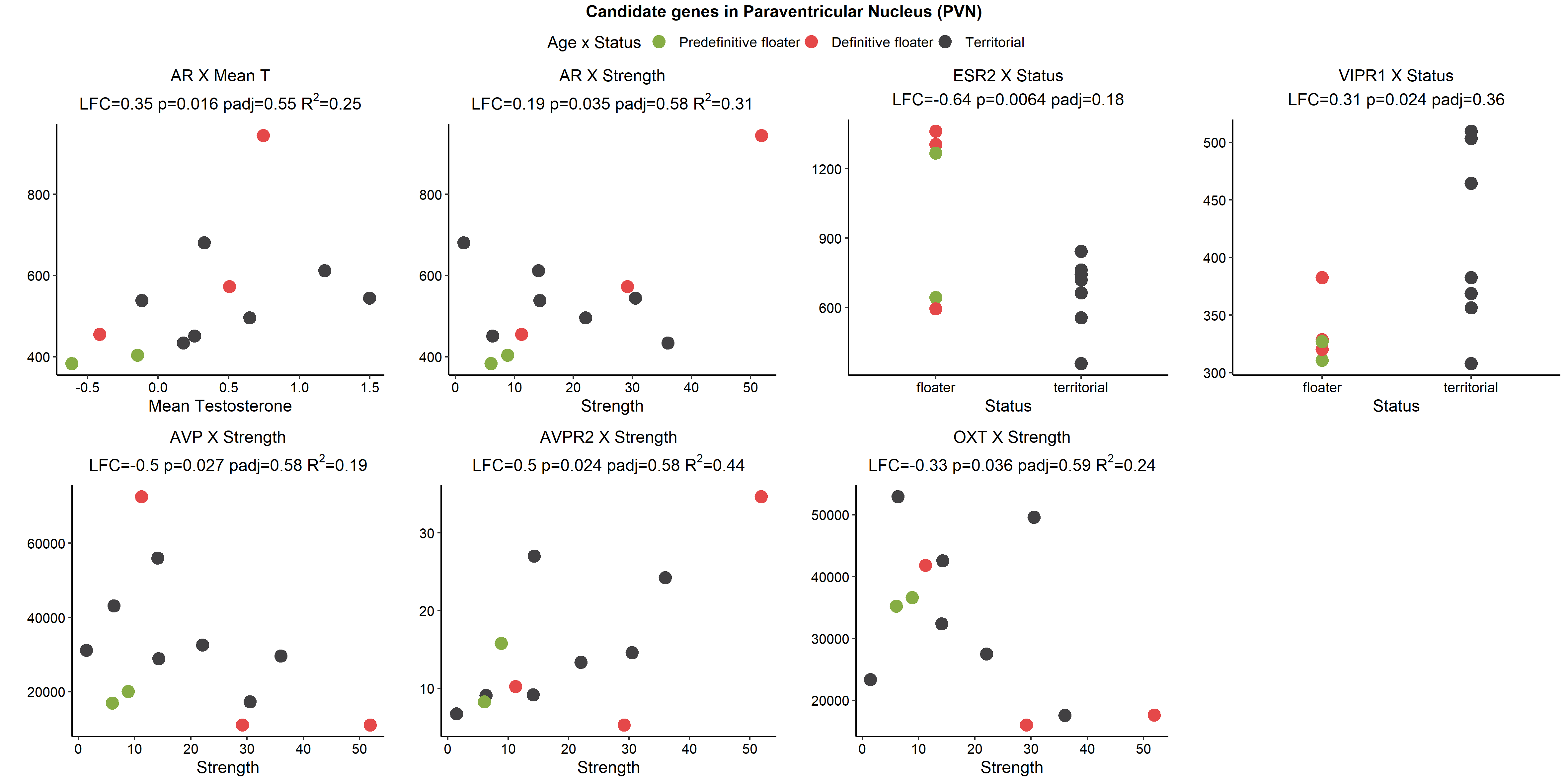

###### Figure S4.8: Candidate genes differentially expressed in the paraventricular nucleus hypothalamus (PVN).

Explanation as in Figure S4.13. AVP was filtered from further analysis owing to low R^2^. Explanations as in Figure S4.3.

###### Figure S4.9: Possible interaction effects in the paraventricular nucleus (PVN).

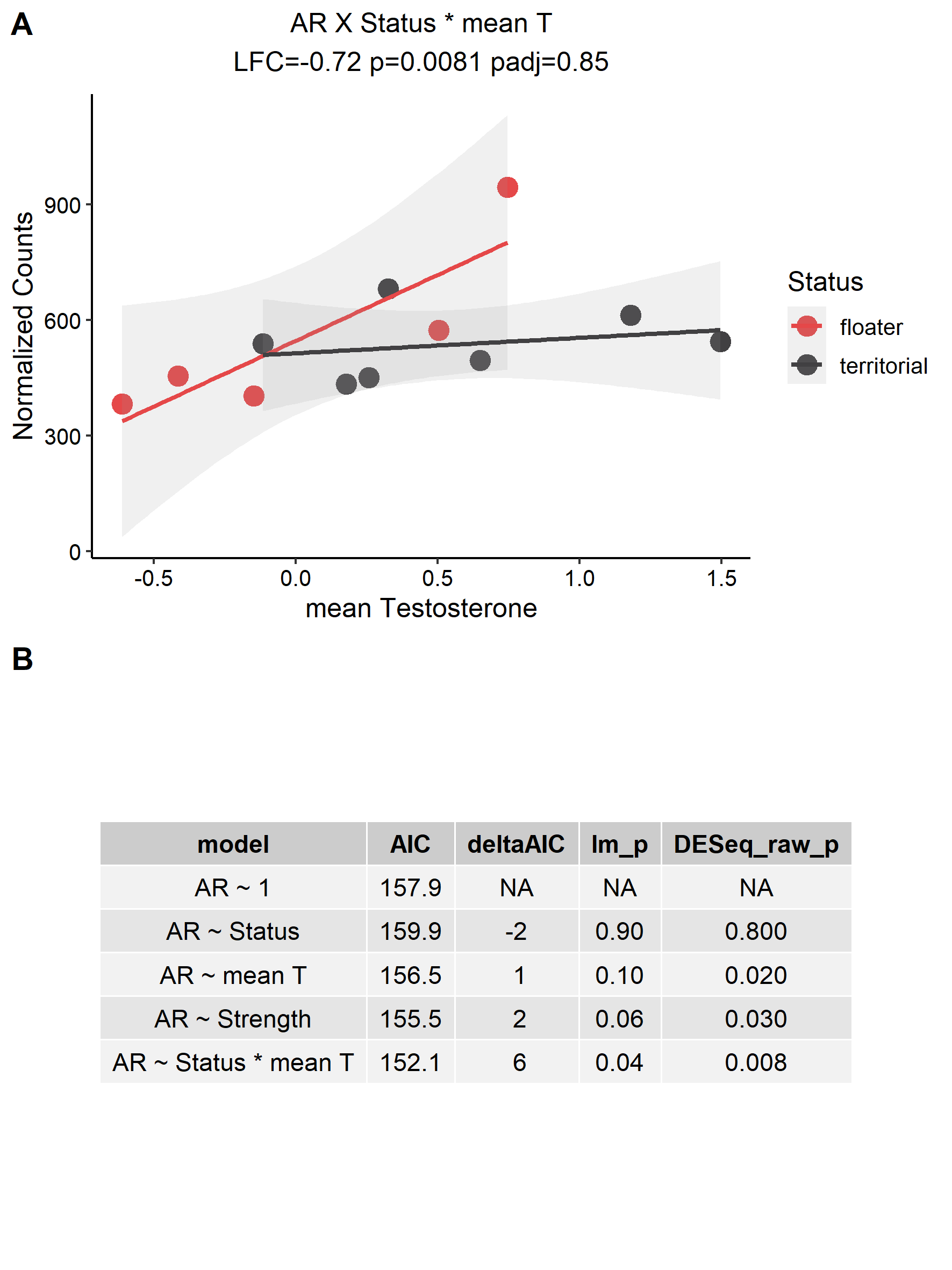

**a**

Explanation as in Figure S4.4. We accepted the interaction model.

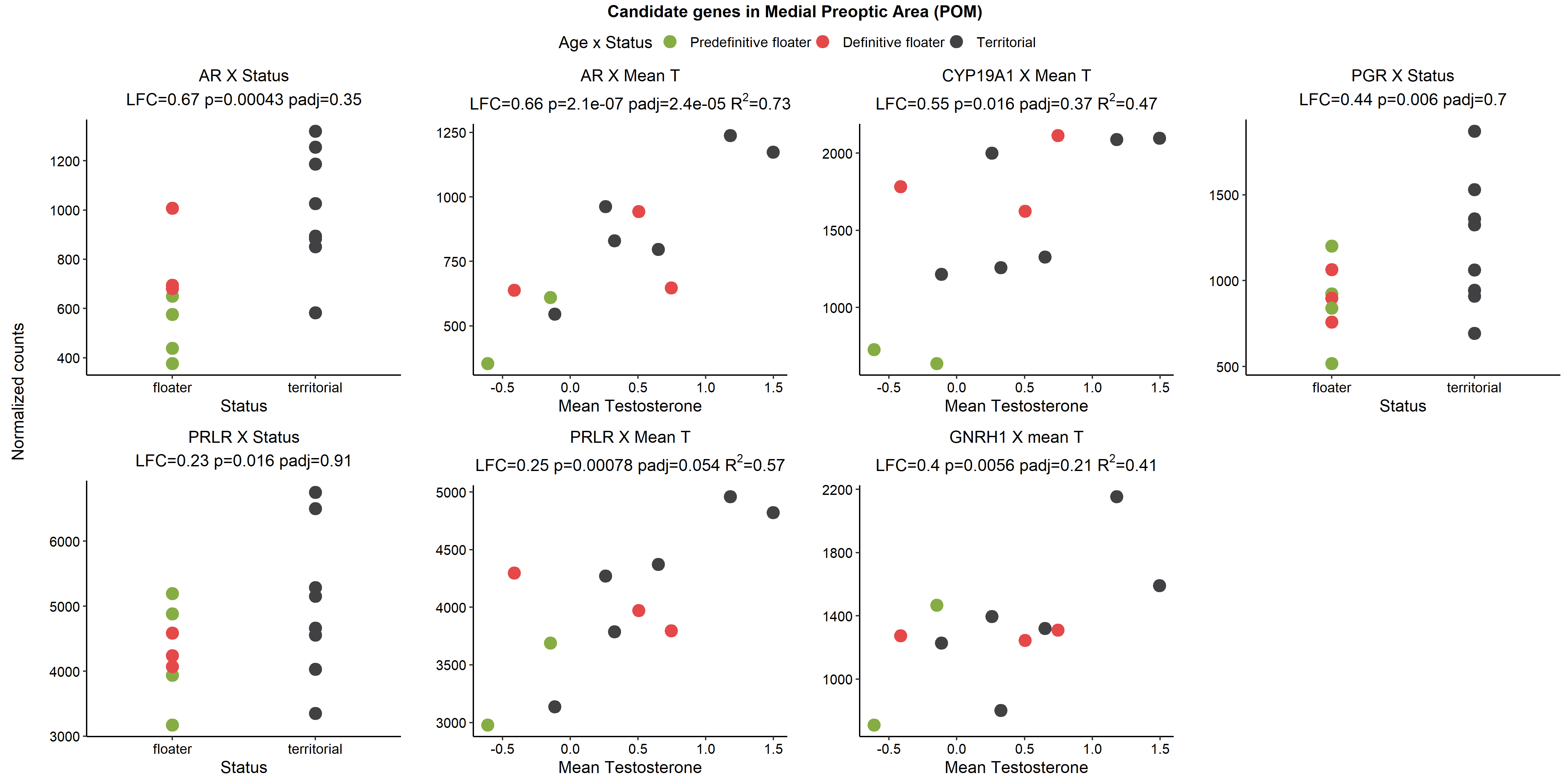

###### Figure S4.10: Candidate genes differentially expressed in the medial preoptic nucleus (POM).

Explanation as in Figure S4.3.

###### Figure S4.11: Possible interaction effects in the medial preoptic area (POM).

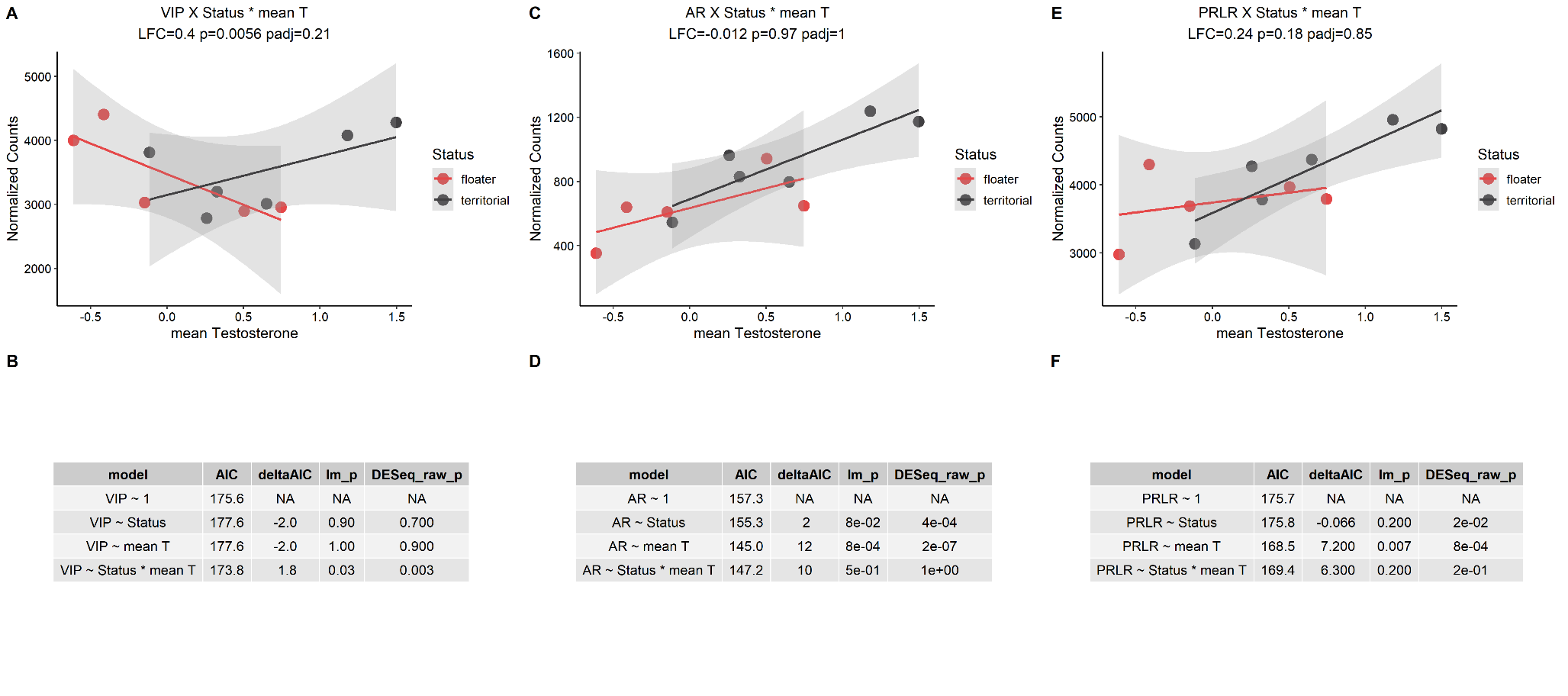

**a**

**a**

VIP was identified in the additional DESeq2 interaction analysis A), but this model was not substantially different from the null model B). AR identified as associated with Status and mean T in the individual models (Figure S4.10), and so an interaction effect was explored C). We accepted the AR ~ mean T relationship, as it’s ΔAIC relative to interaction model was 2.2 D). PRLR was identified as associated with Status and mean T in the individual models (Figure S4.10), and so interaction and model fits were explored E). We chose PRLR ~ mean T on the basis of lowest AIC and significant p-values F).

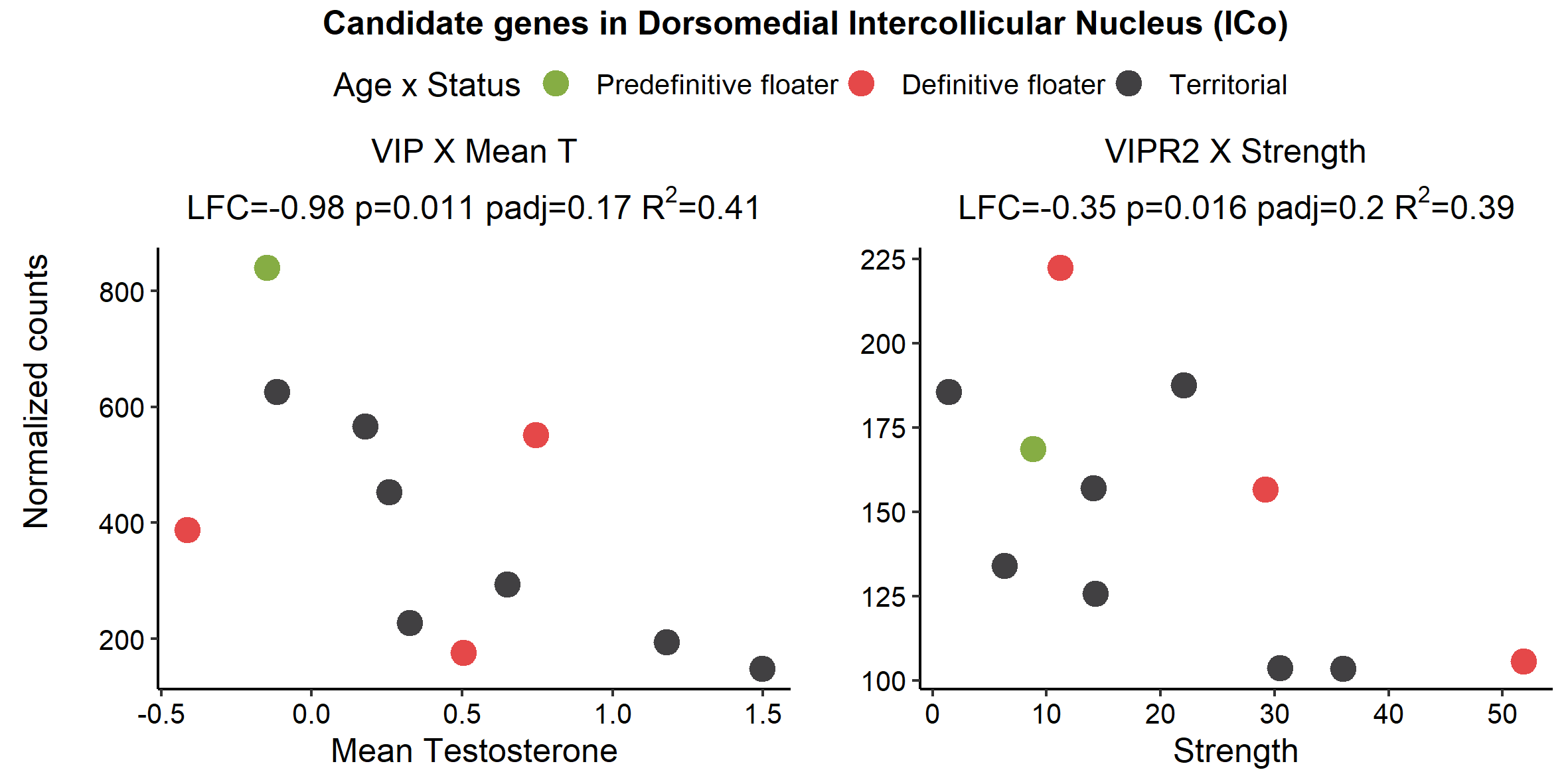

###### Figure S4.12: Candidate genes differentially expressed in the intercollicular nucleus (ICo).

Explanation as in Figure S4.3.

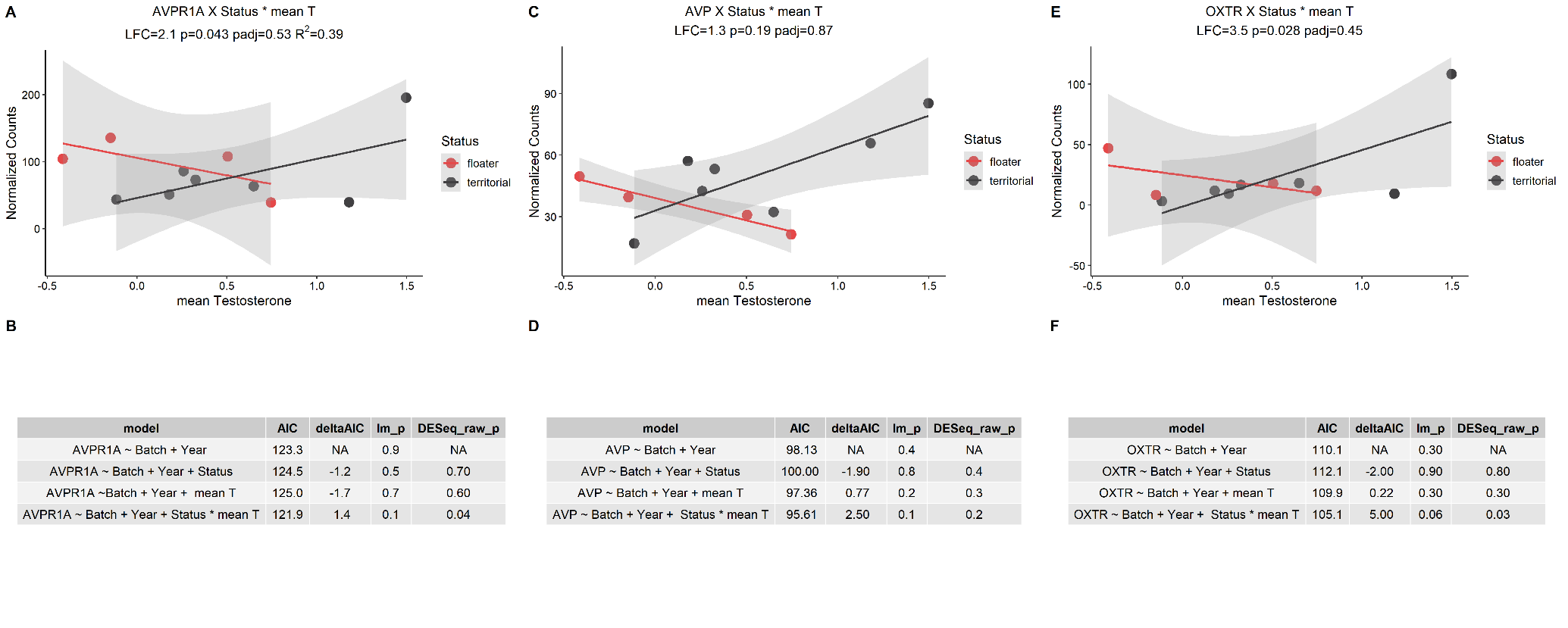

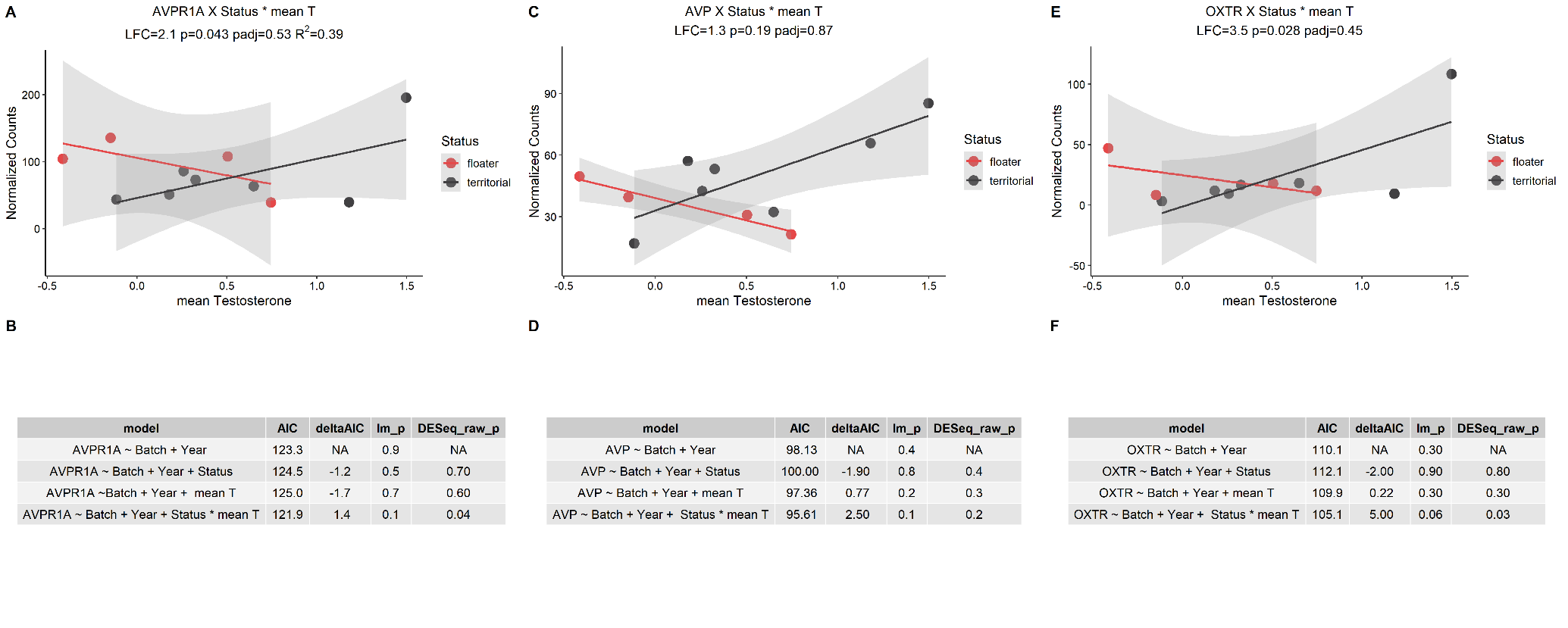

###### Figure S4.13: Possible interaction effects in the intercollicular nucleus (ICo).

Explanations as Figure S4.4. DESeq2 identified AVPR1A to have a possible interaction effect A), but the model’s AIC was not substantially better than the null and was thus discarded B). OXTR showed many individuals with very low counts and so was excluded on this basis C) despite support for the interaction model D).

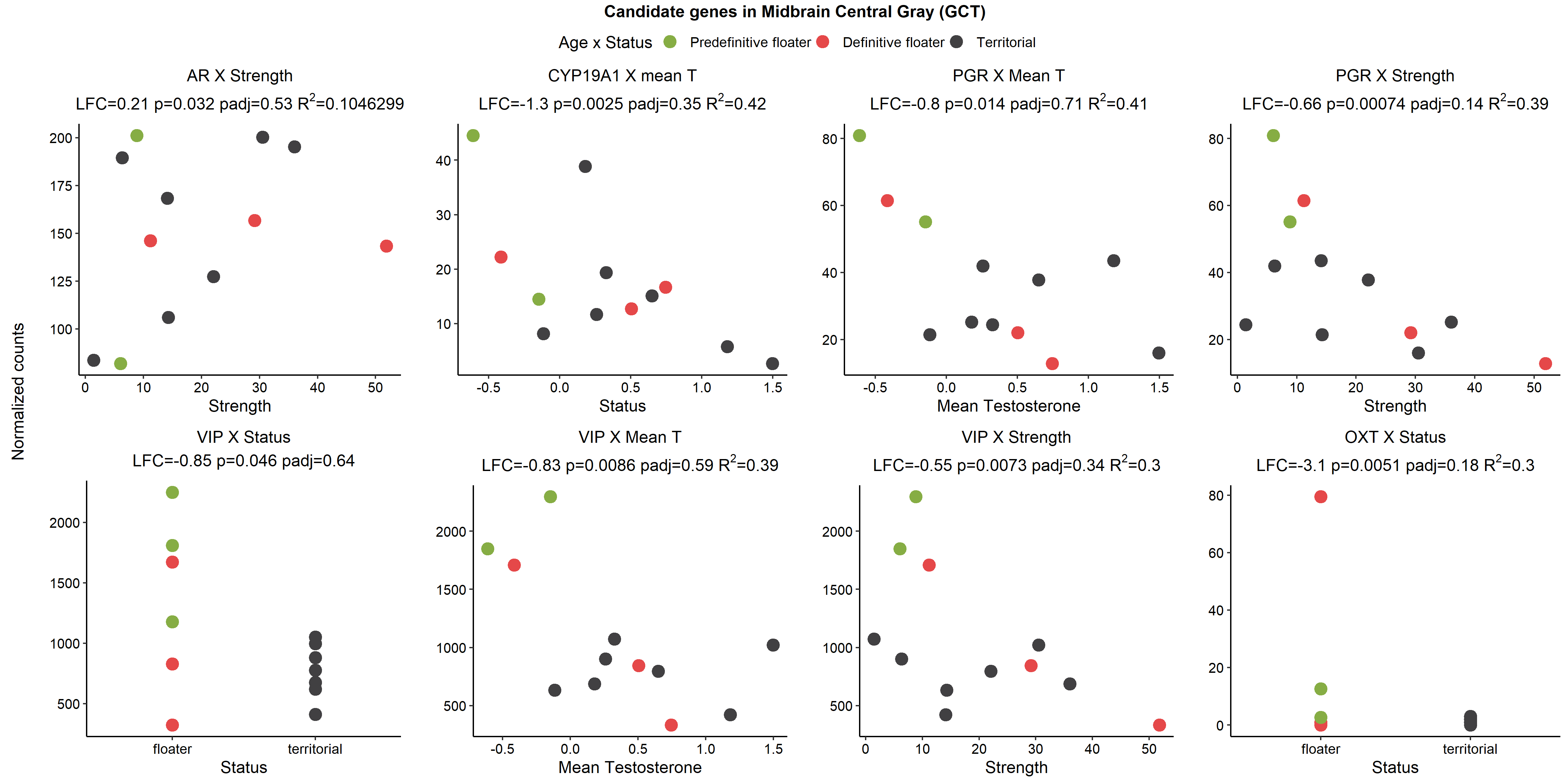

###### Figure S4.14: Candidate genes differentially expressed in the midbrain central grey (GCt).

Explanations as in Figure S4.3. AR was excluded on the basis of low R^2^, and OXT was excluded on the basis of an influential observation.

###### Figure S4.15: Possible interaction effects in the midbrain central grey (GCt).

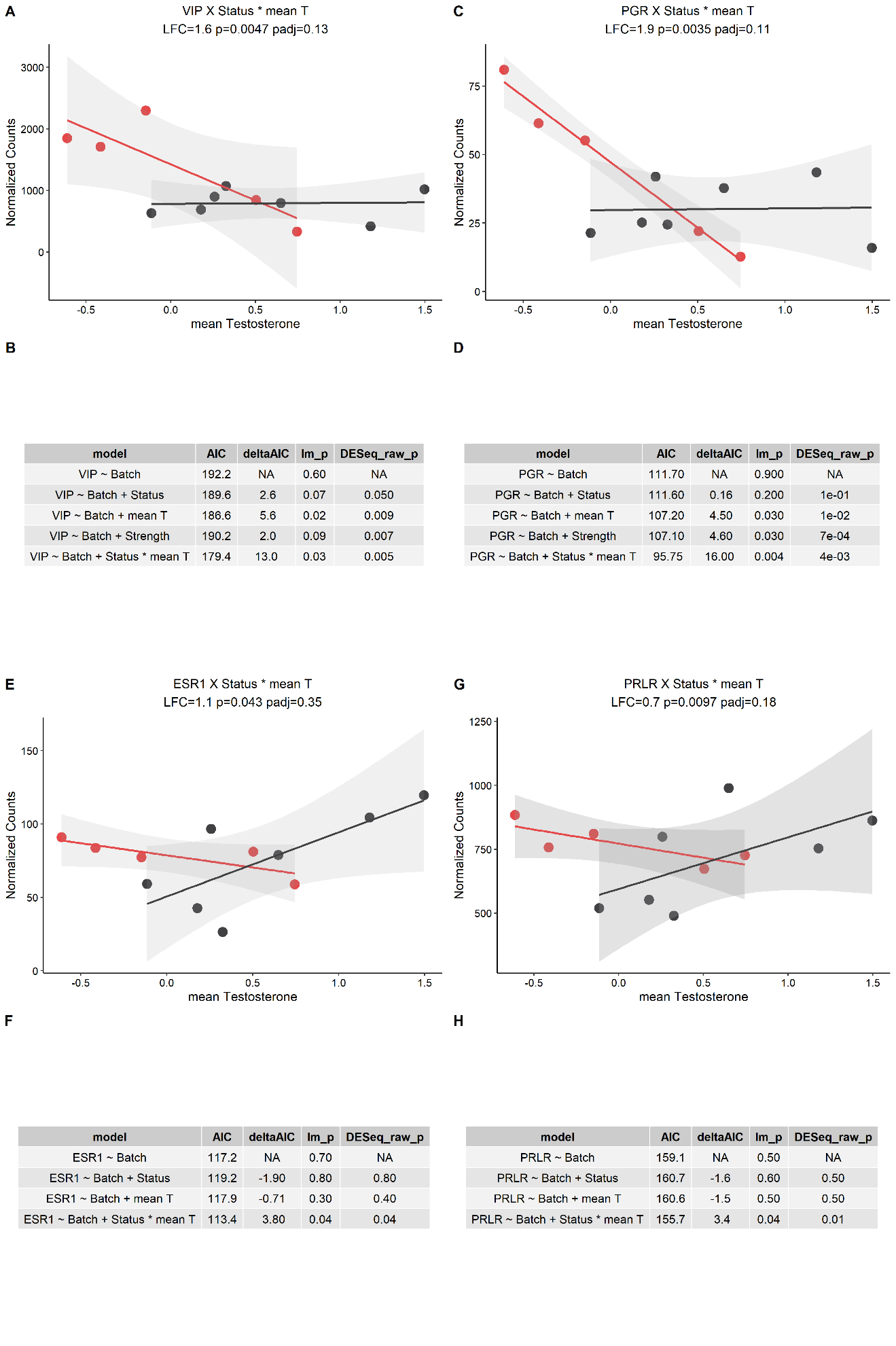

**a**

**a**

**a**

**a**

All four candidate genes were substantially better than the null and simpler models. Explanations as in Figure S4.4.

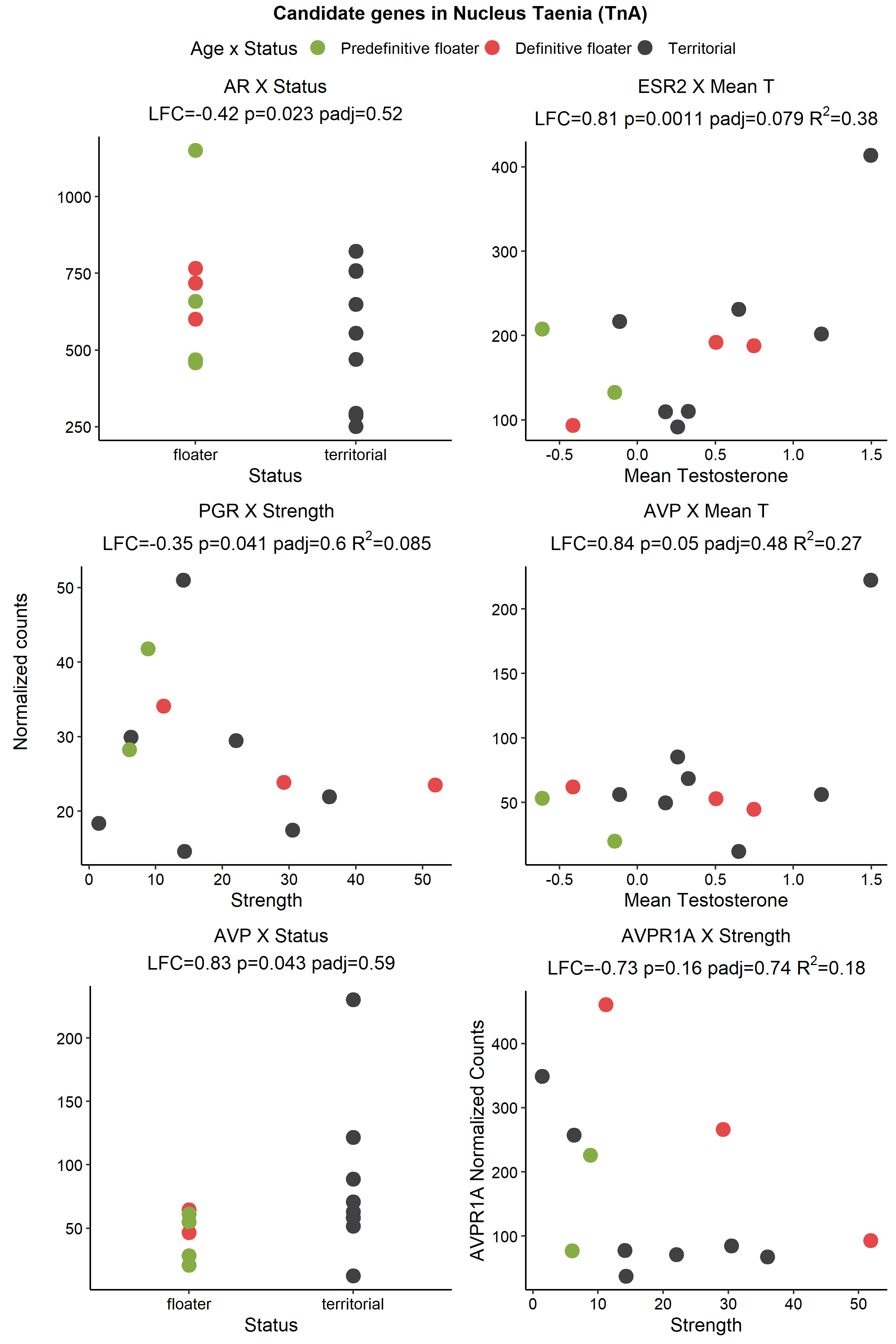

###### Figure S4.16: Candidate genes differentially expressed in the nucleus taenia (TnA).

AR, PGR, AVP, and AVPR1A were all excluded from further analysis owing to R^2^<0.3 and influential observations. Explanations as in Figure S4.3.

###### Figure S4.17: Possible interaction effects in the nucleus taenia (TnA).

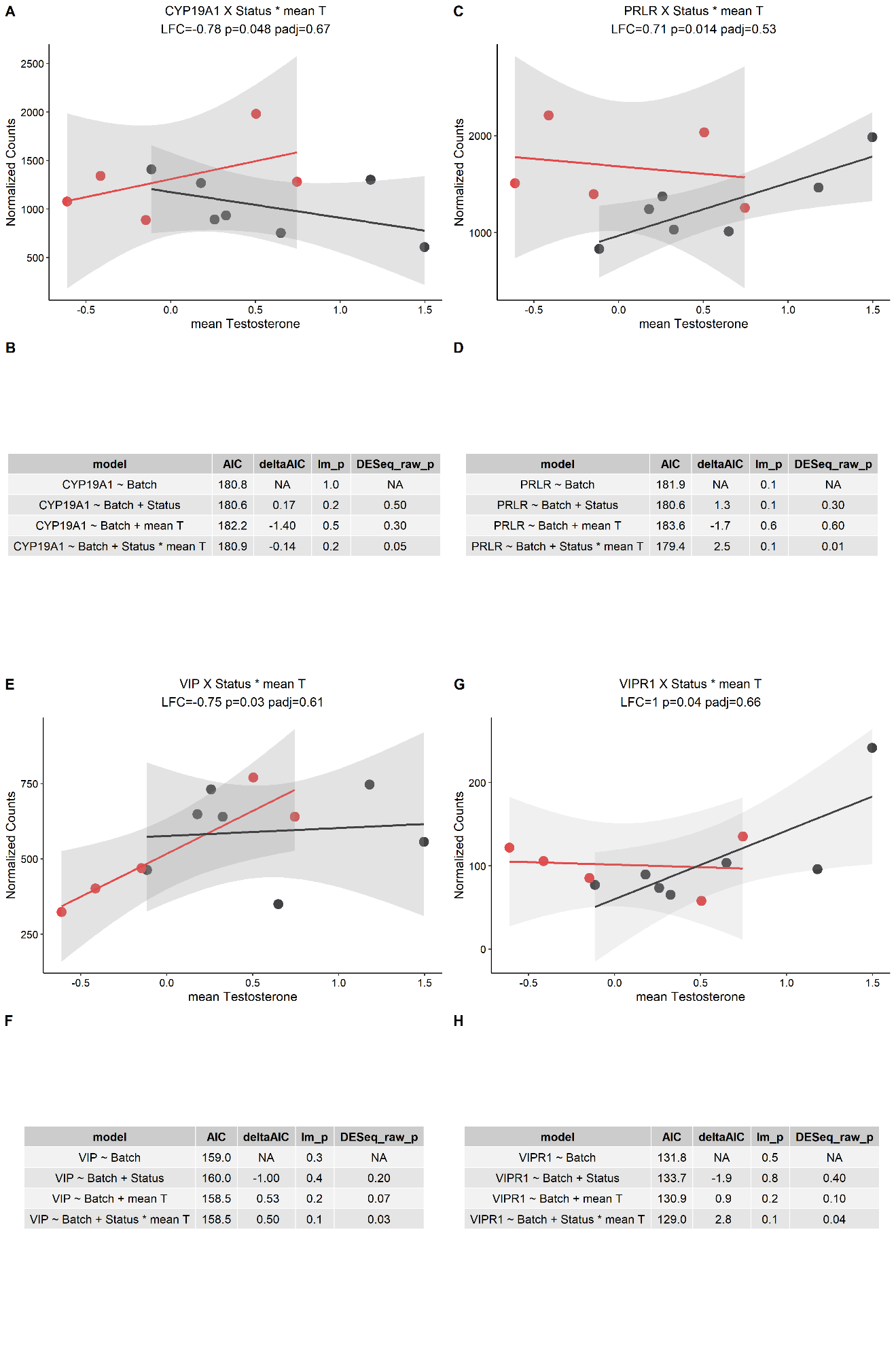

**a**

**a**

Only PRLR and VIPR1 showed substantial improvement over the null model. Explanations as Figure S4.4.

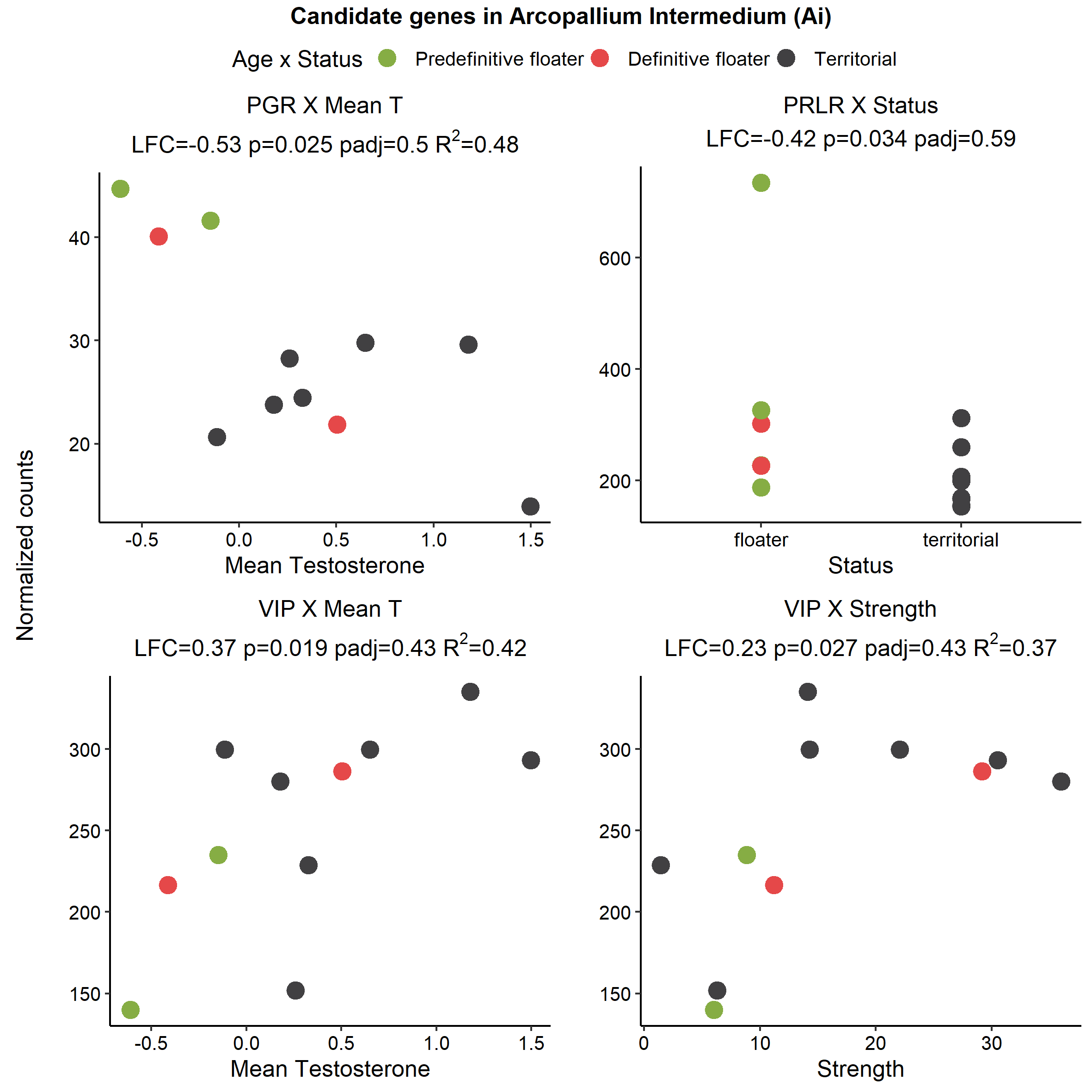

###### Figure S4.18: Candidate genes differentially expressed in the arcopallium intermedium (AI).

Explanations as in Figure S4.3.

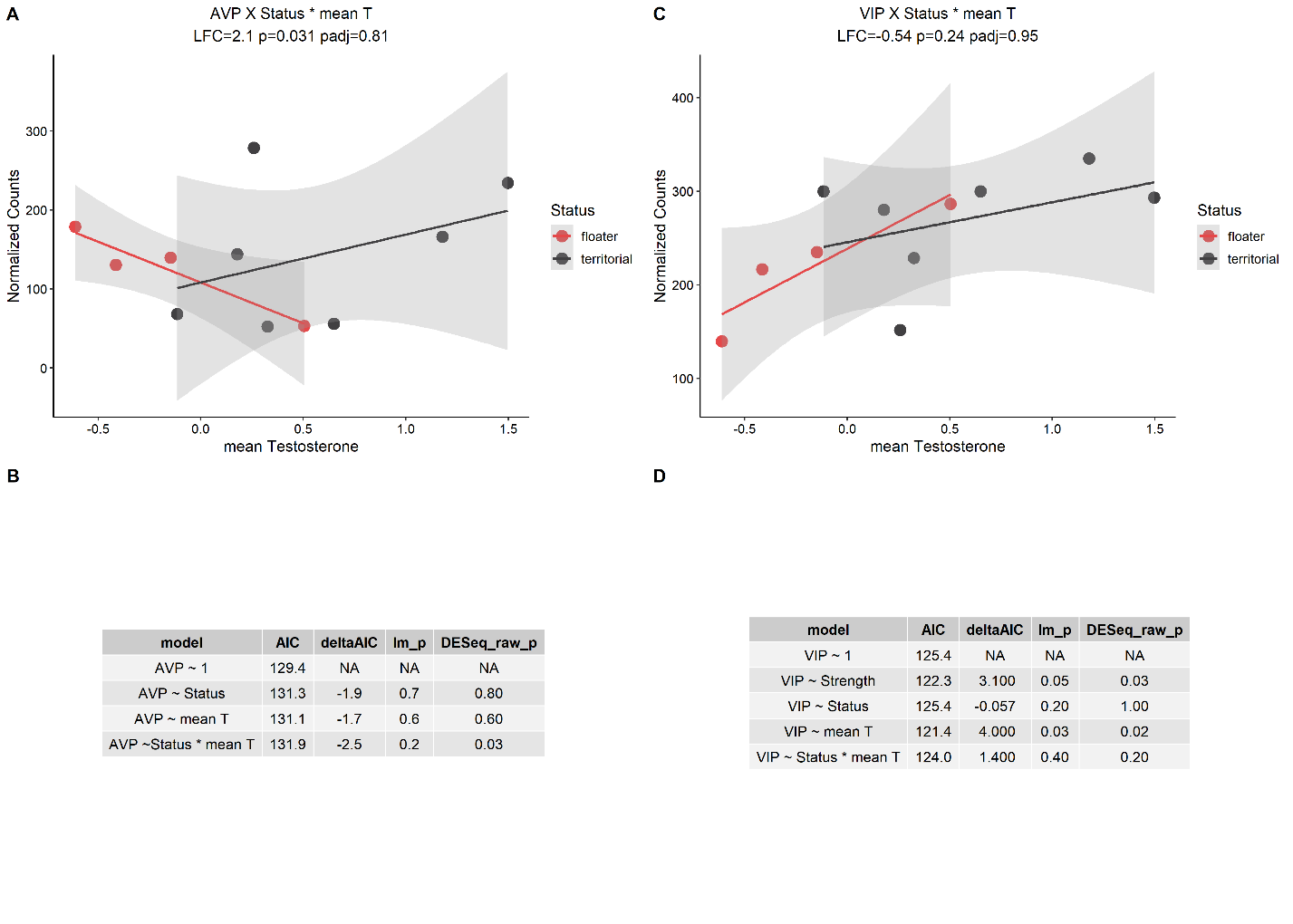

###### Figure S4.19: Possible interaction effects in the arcopallium intermedium (AI).

Explanations as Figure S4.4. AVP A) was excluded from the final list as no model improved from the null B). VIP was significantly associated with both Strength and mean T (Figure S4.18), and both were better than the null D), and were retained on both lists. VIPR1 and AVPR1A were not plotted here because they did not show a compelling visual relationship, as well as no model improvements. See the [GitHub](https://periperipatus.github.io/PIFI_brain_transcriptome/05_candidate_gene_analysis.html#11_Arcopallium_Intermedium_(Ai)) for more details.

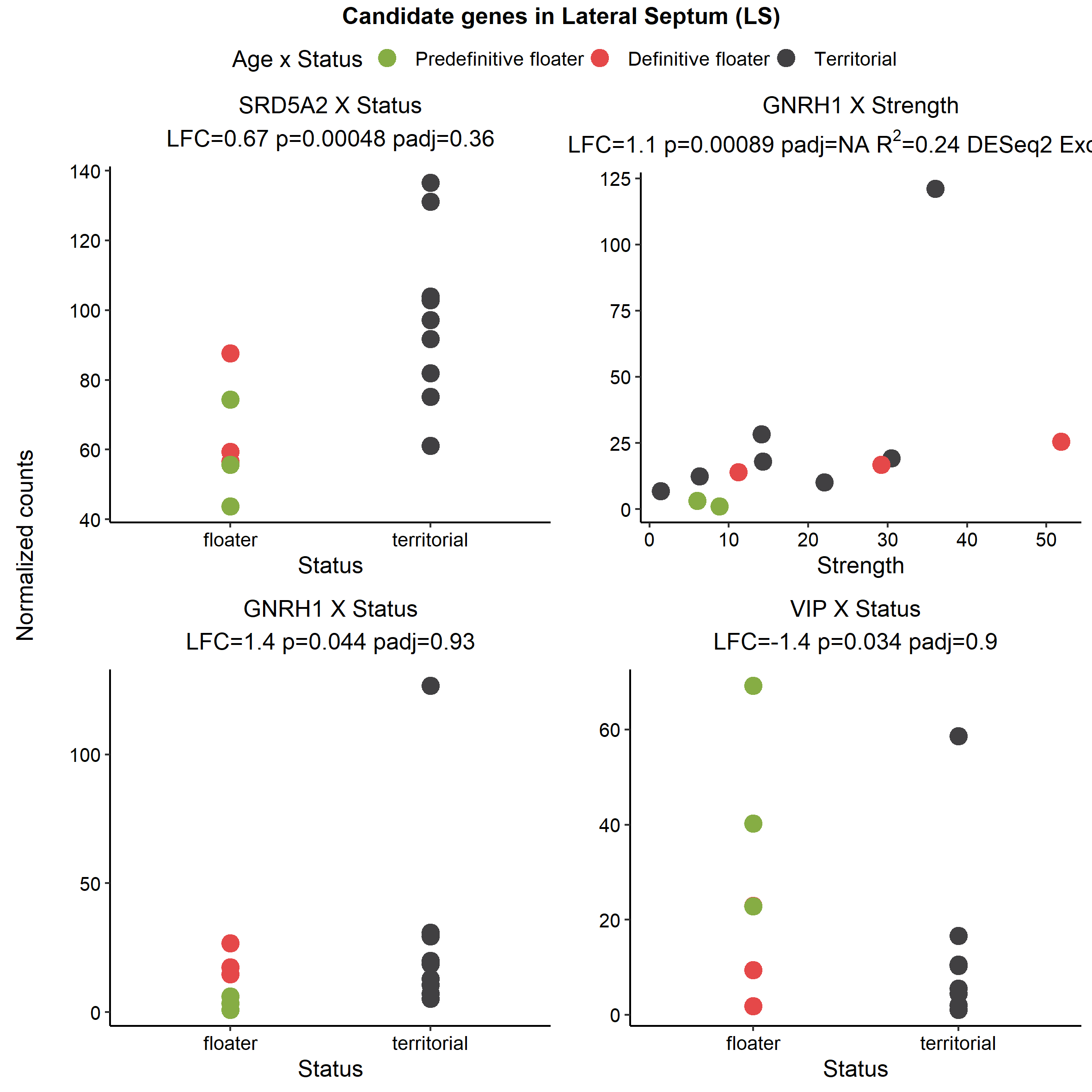

###### Figure S4.20: Candidate genes differentially expressed in the lateral septum (LS).

GNRH1 x Strength comparison was excluded from further analysis by DESeq2 based on outliers, and GNRH1 x Status and VIP were excluded by the authors for the same reasons. Explanations as in Figure S4.3.

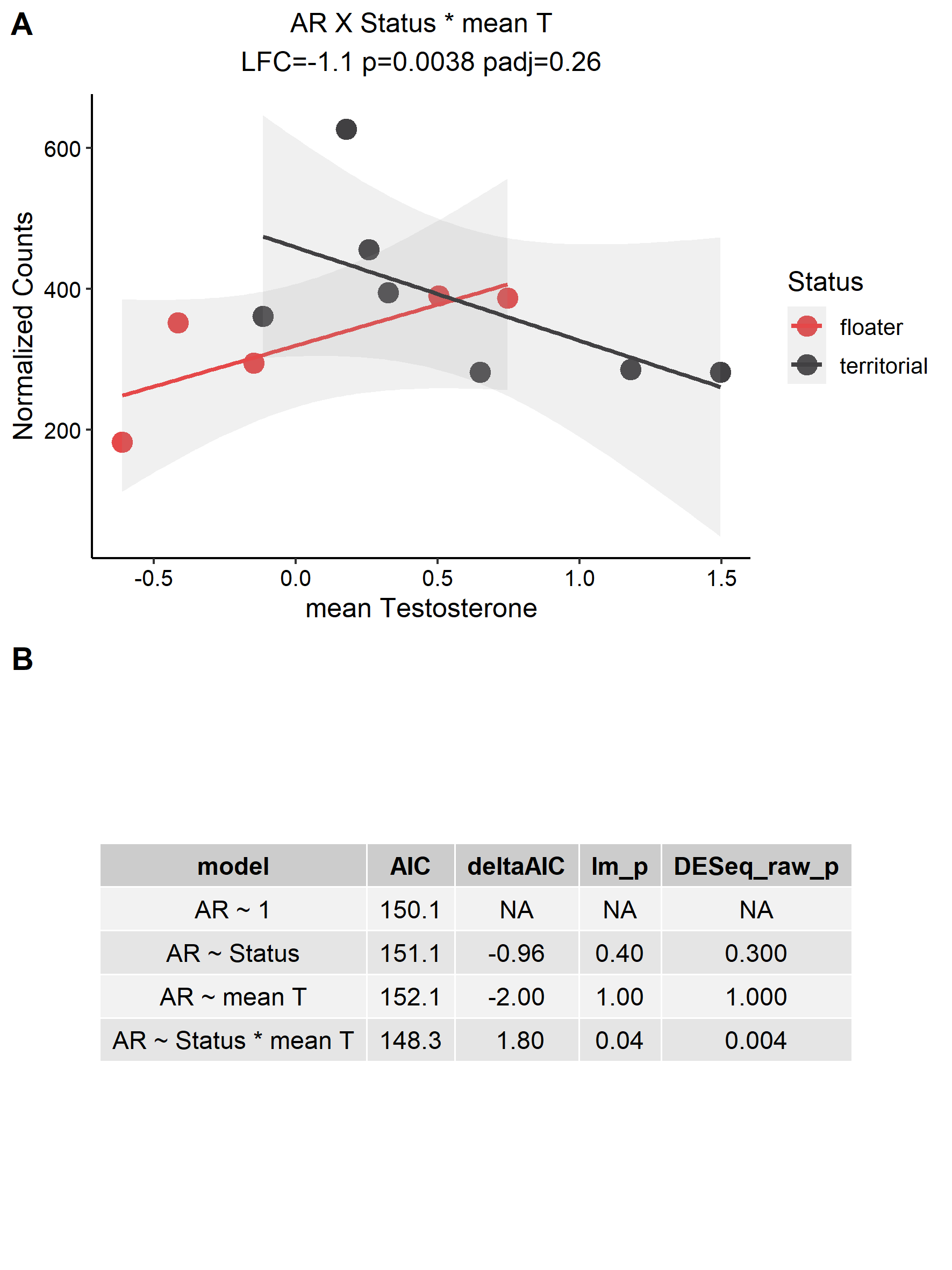

###### Figure S4.21: Possible interaction effects in the lateral septum (LS).

Explanations as Figure S4.4. AR was excluded from the list of genes with interaction effects, as it did not improve substantially on the null model B).

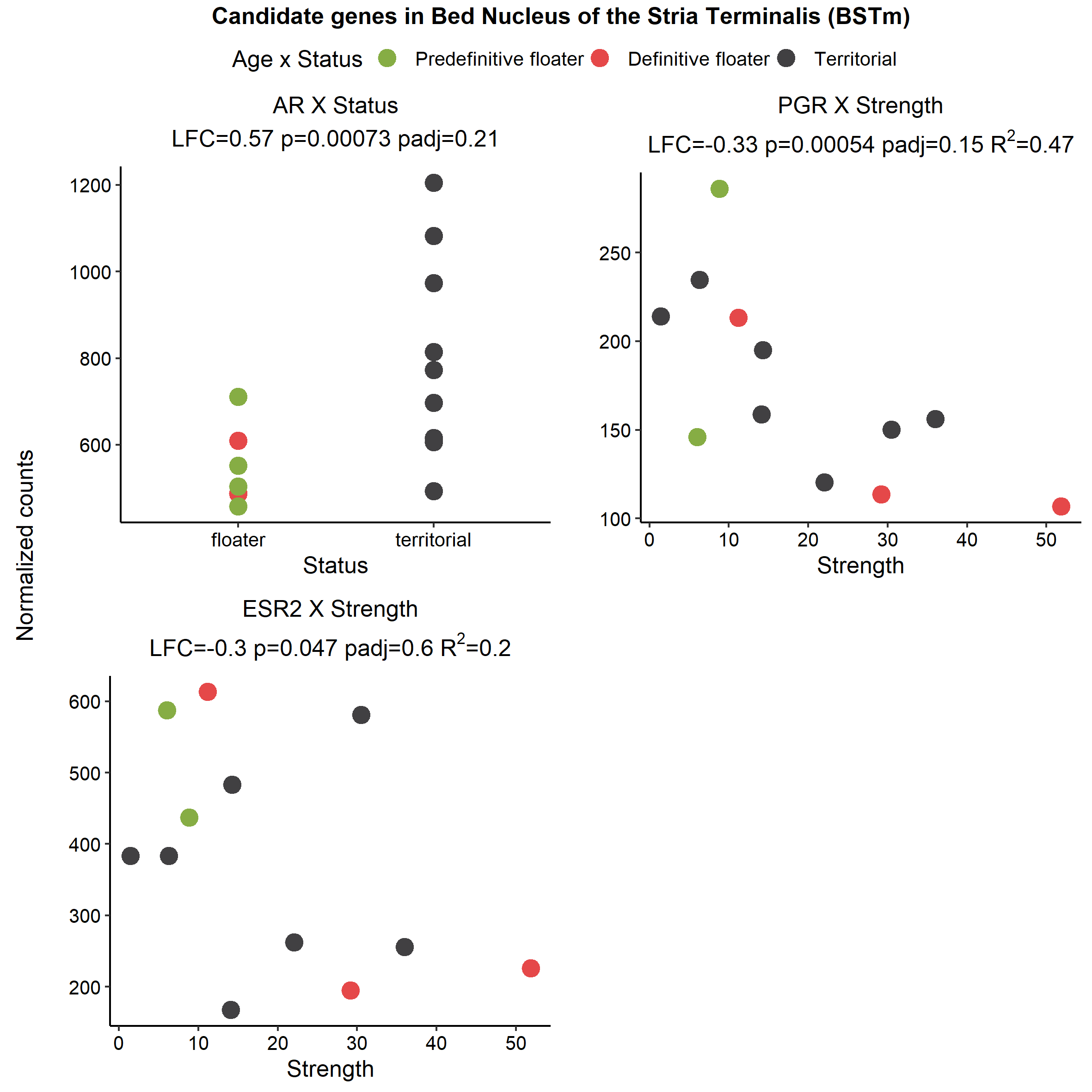

###### Figure S4.22: Candidate genes differentially expressed in the bed nucleus of the stria terminalis (BSTm).

ESR2 was excluded from further analysis due to low R^2^. Explanations as in Figure S4.3.

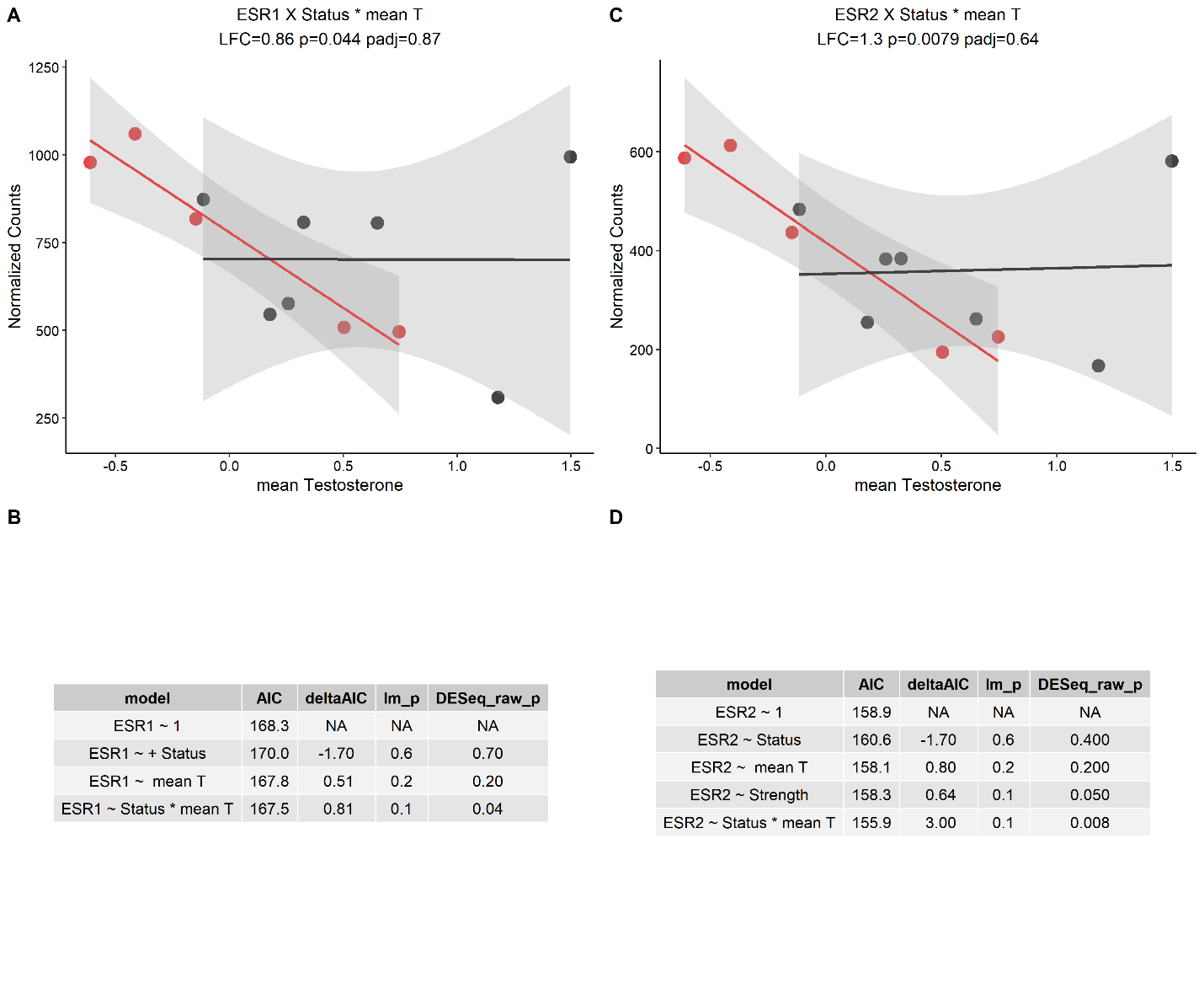

**a**

###### Figure S4.23: Possible interaction effects in the bed nucleus of the stria terminalis (BSTm).

Explanations as Figure S4.4. DESeq2 identified a possible interaction effect with ESR1 A) but this model was not better than the null B). ESR2 was associated with both social network strength and DESeq2 suggested an interaction effect C). Strength was discarded from ESR2 as it had a low R2 and also did not improve model fit from the interaction, and the interaction was improved from the null D).

### 5. Within Tissue WGCNA

To explore potential regulatory networks associated with our social network strength, status, and mean T, we conducted WGCNA analysis (Langfelder and Horvath 2008). Because the between-tissue signal is the strongest co-expression pattern (Horton *et al.* 2020), we created co-expression matrices for each individual tissue, after correcting the data for batch effects using *limma::RemoveBatchEffect*. Co-expression adjacency matrices were generated using Pearson correlation and signed network types, with soft-thresholding power (β) assigned empirically for each tissue (range 9-22). Modules were assigned using dynamic tree cutting, with a minimum module size of 30, and similar modules were merged on a similarity threshold of 0.3, meaning 70% of genes must be shared amongst two modules before merging. Module eigengenes were Pearson correlated to our traits of interest, with a correlation of p<0.05 considered compelling. Hub genes were determined based on Module Membership scores.

The full detail of these methods and results can be found at the [GitHub](https://periperipatus.github.io/PIFI_brain_transcriptome/index.html). These analyses served as an independent estimate of relationship between DEGs and traits of interest.

See figures S5.1-S5.14 for associations between co-expression modules and our proposed explanatory variables, and for selected nuclei relevant GO enrichment results (S5.7 and S5.11)

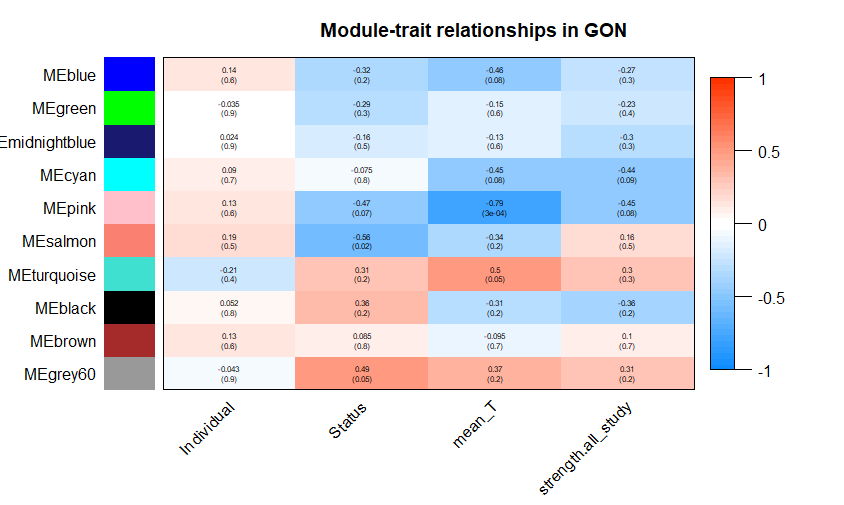

###### Figure S5.1: Correlation between gene co-expression modules and traits in the testes (GON).

Module eigengenes Pearson correlated with traits of interest. The y-axis indicates the module, denoted by a randomly assigned color. Red squares indicate a positive correlation coefficient between module eigengene and trait, and blue indicates a negative correlation. The correlation coefficient is indicated in the text, and the p-value is indicated in parentheses.

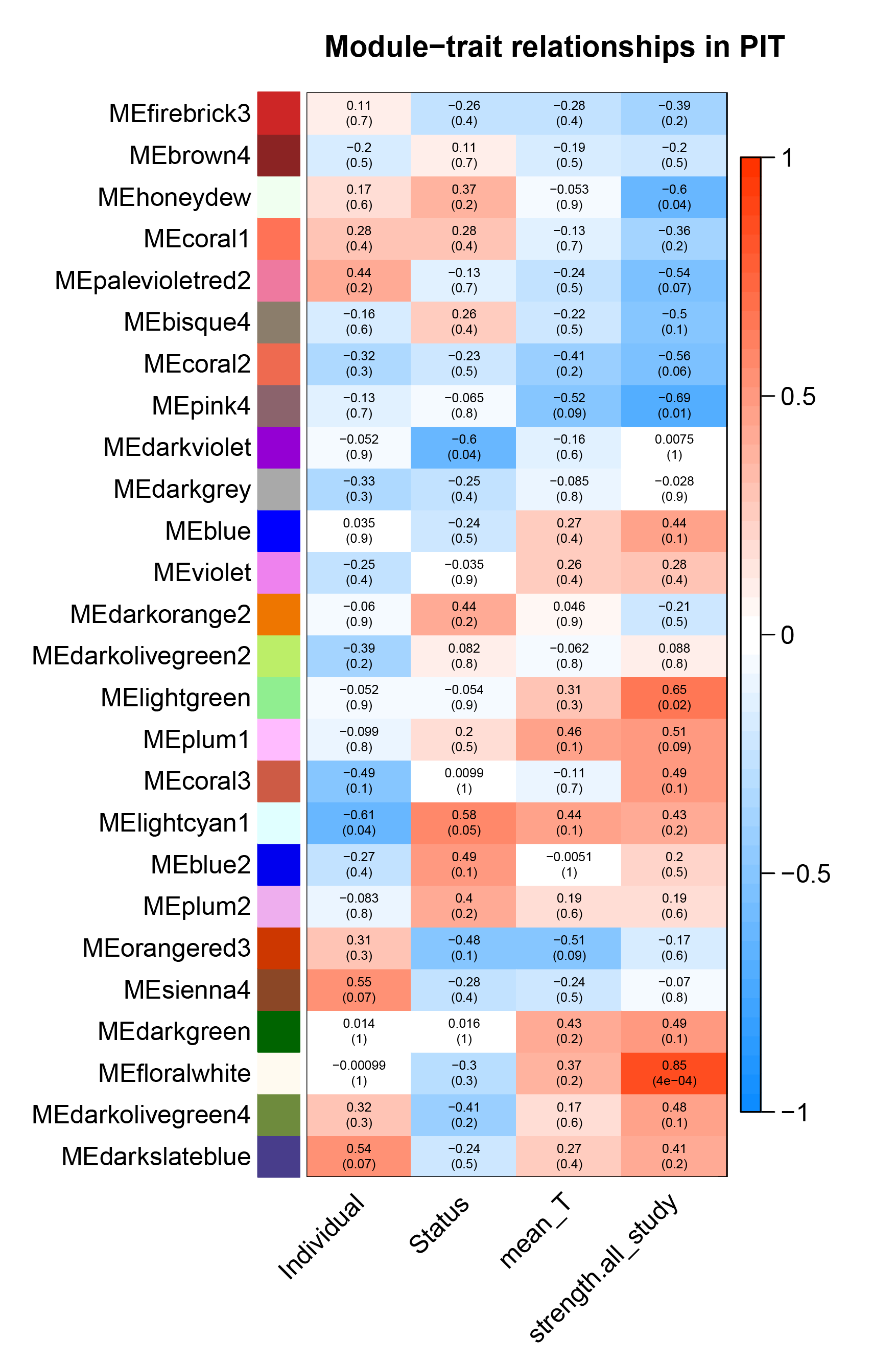

###### Figure S5.2: Correlation between gene co-expression modules and traits in the pituitary (PIT).

Explanation as in Figure S5.1.

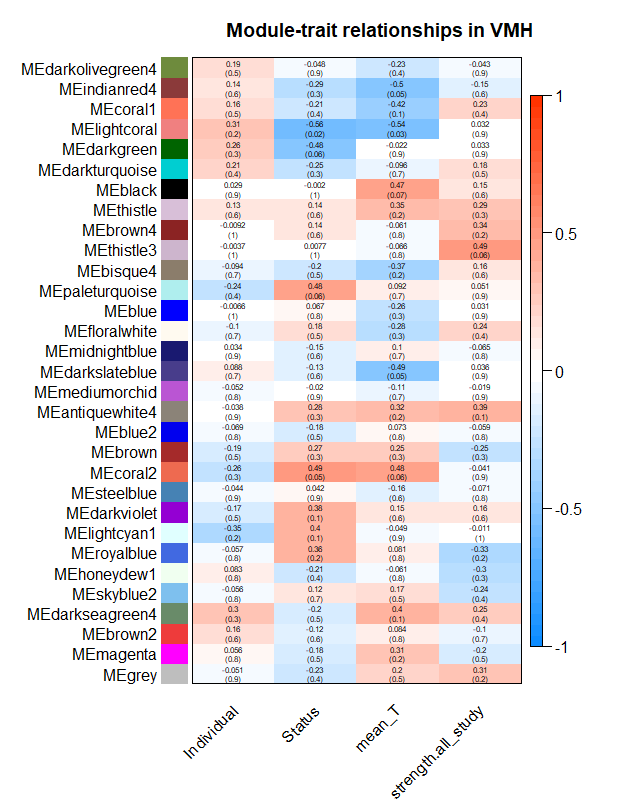

###### Figure S5.3: Correlation between gene co-expression modules and traits in the ventromedial hypothalamus (VMH).

Explanation as Figure S5.1.

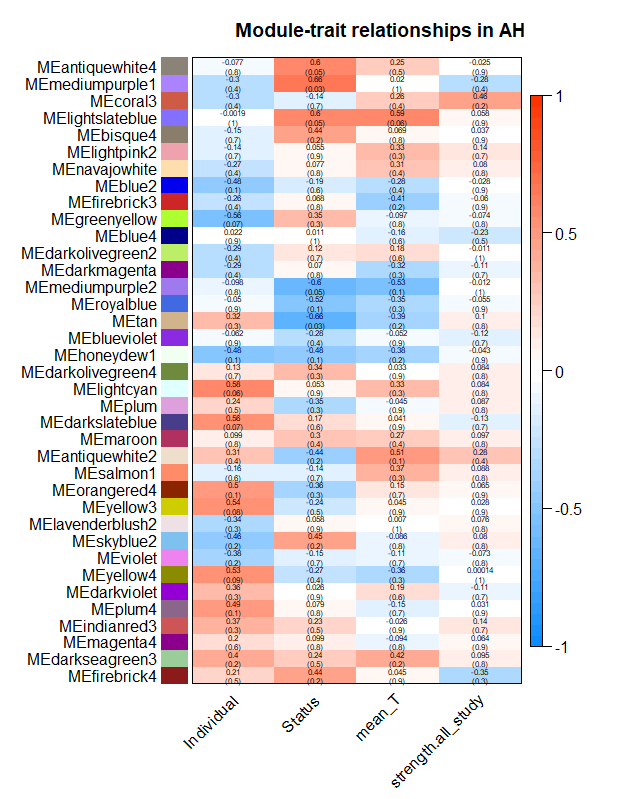

###### Figure S5.4: Correlation between gene co-expression modules and traits in the anterior hypothalamus (AH).

Explanations as Figure S5.1.

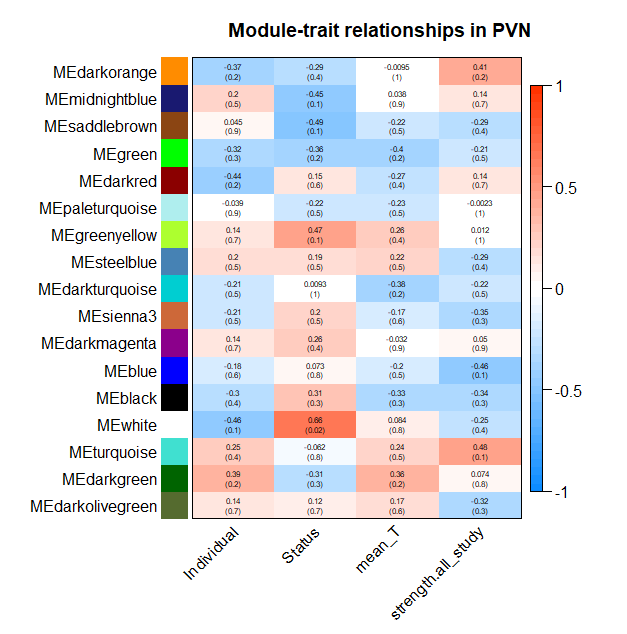

###### Figure S5.5: Correlation between gene co-expression modules and traits in the paraventricular nucleus (PVN).

Explanation as Figure S5.1.

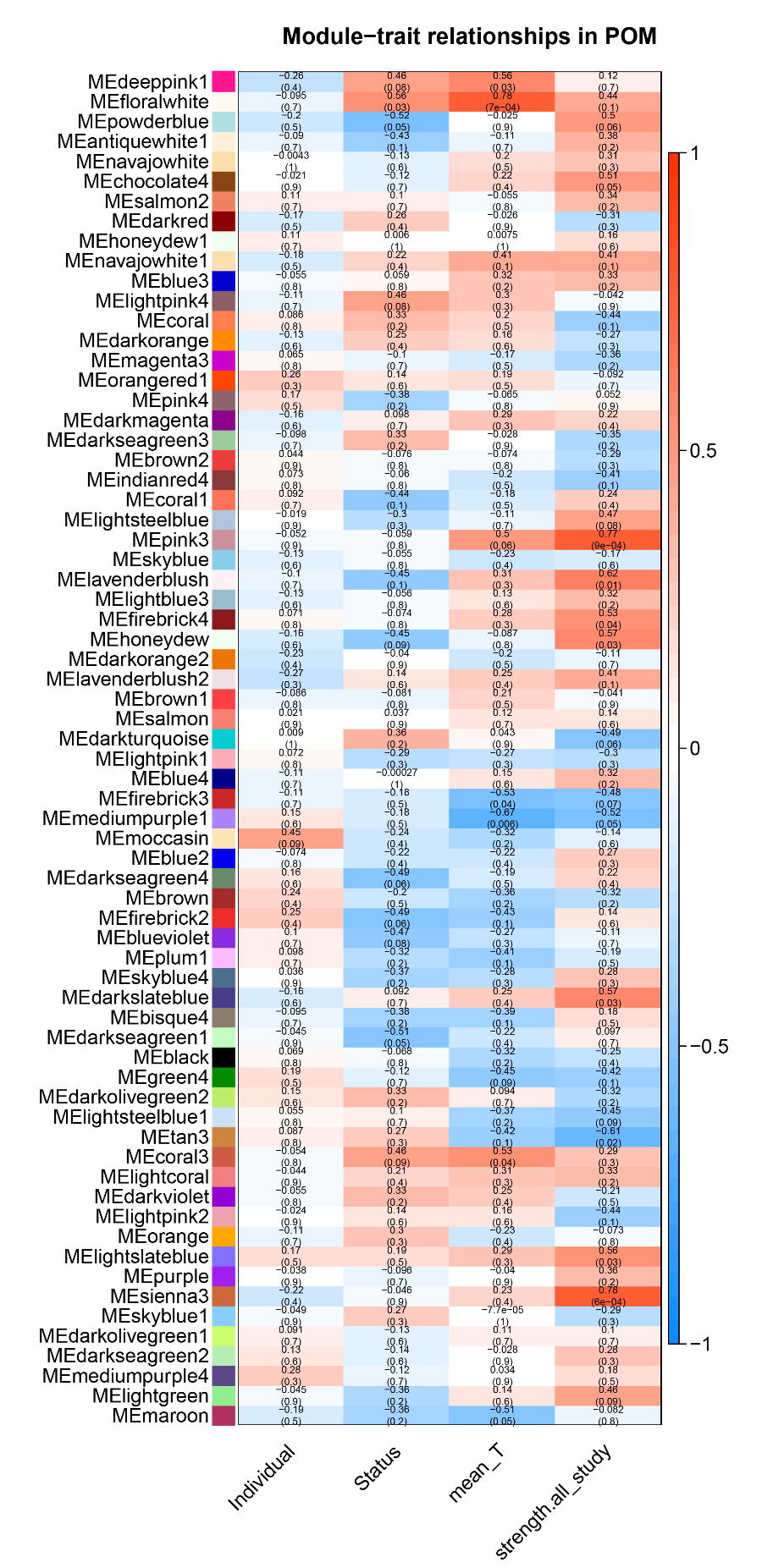

###### Figure S5.6: Correlation between gene co expression modules and traits in the medial preoptic nucleus (POM).

Explanation as Figure S5.1

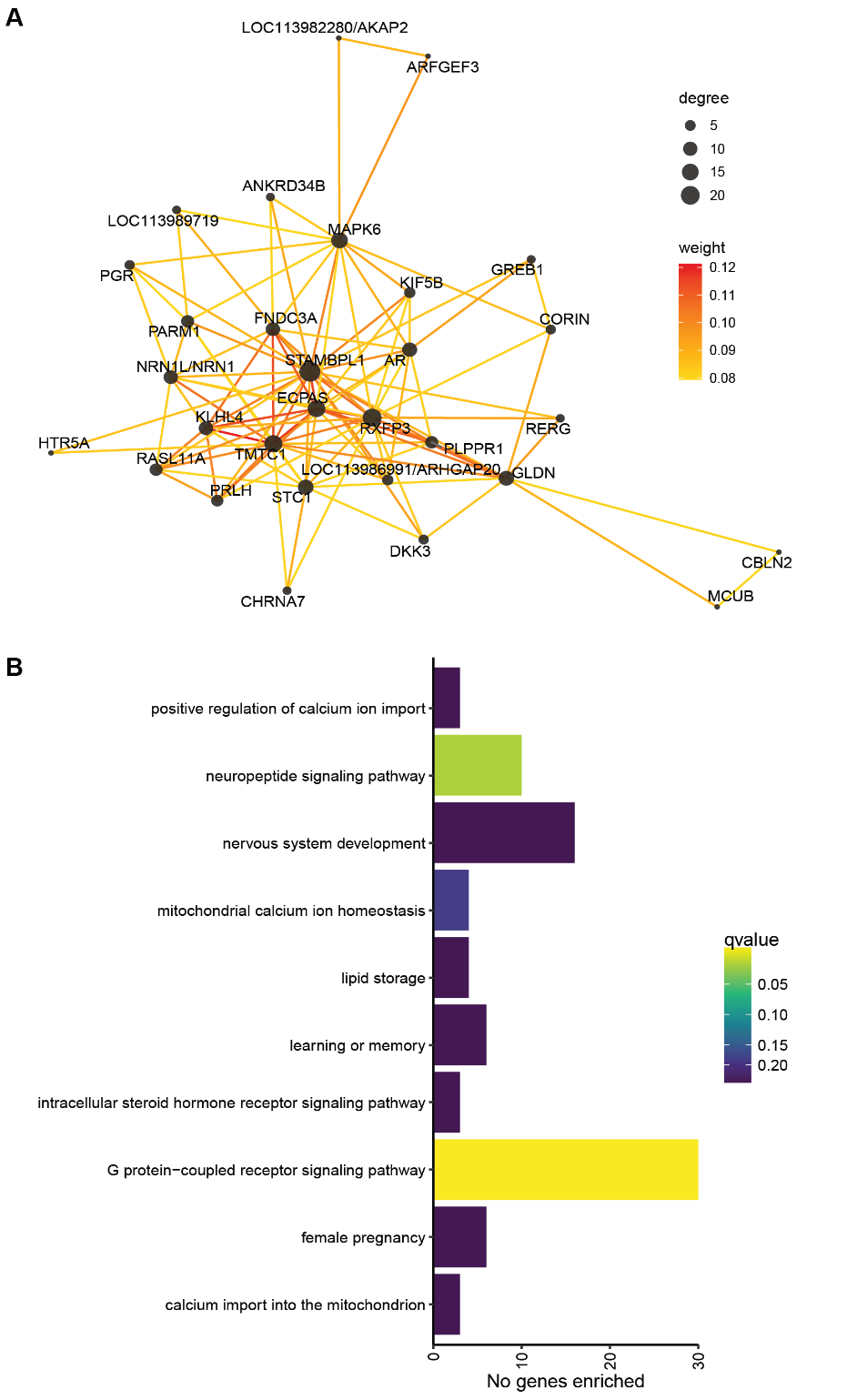

###### Figure S5.7: Network diagram of top 30 genes in POM’s ‘floralwhite’ module and GO enrichment.

A) Topological overlap network where size of the node represents number of connections (degree) and the edge color represents the topological overlap (roughly similarity of gene expression) between the genes. Notable hub genes are PRLH, HTR5A, PGR, AR, and GREB1, B) Biological Process GO enrichment for all 391 genes in the ‘floralwhite’ module. Where bar color represents the FDR adjusted p-value.

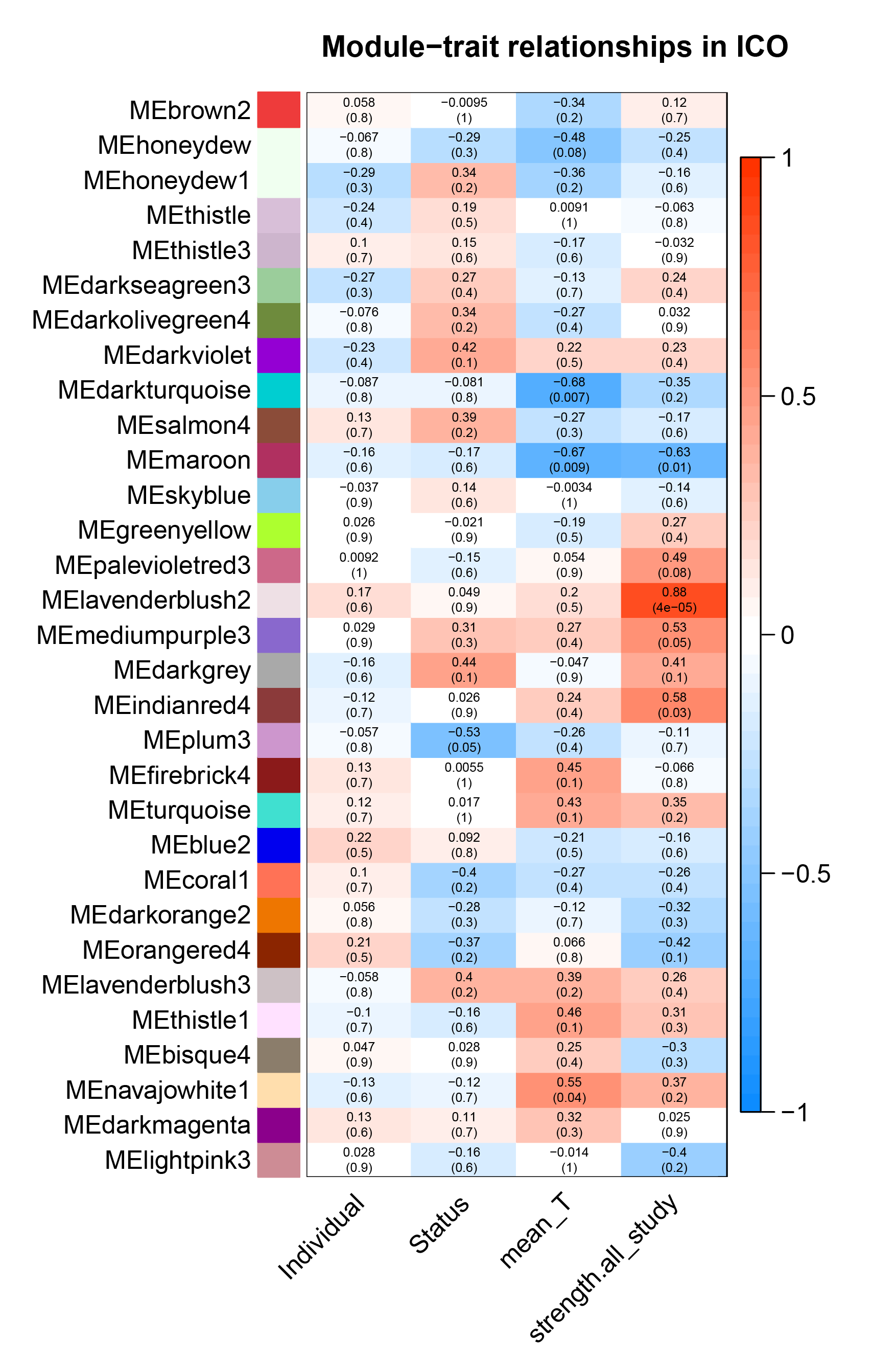

###### Figure S5.8: Correlation between gene co-expression modules and traits in the intercollicular nucleus (ICo).

Explanation as in Figure S5.1.

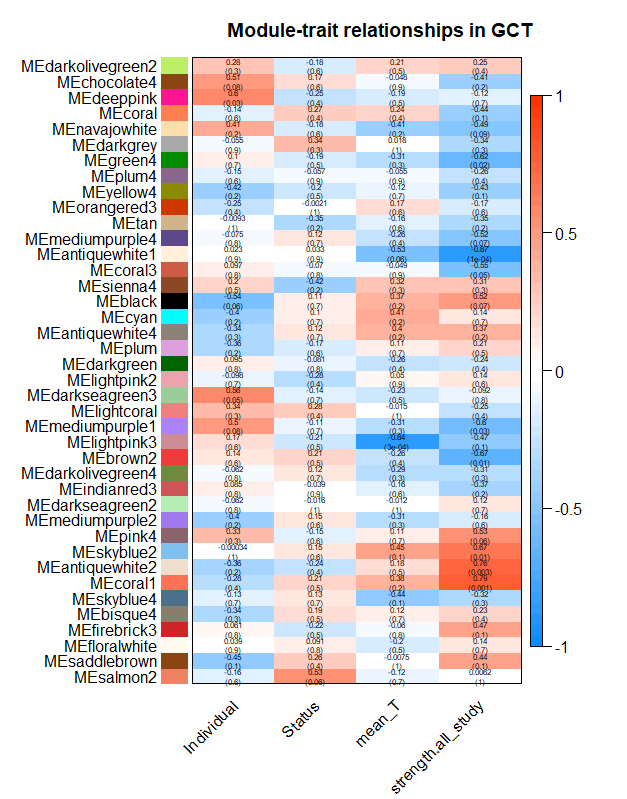

###### Figure S5.9: Correlation between gene co-expression modules and traits in the midbrain central grey (GCt).

Explanation as in Figure S5.1.

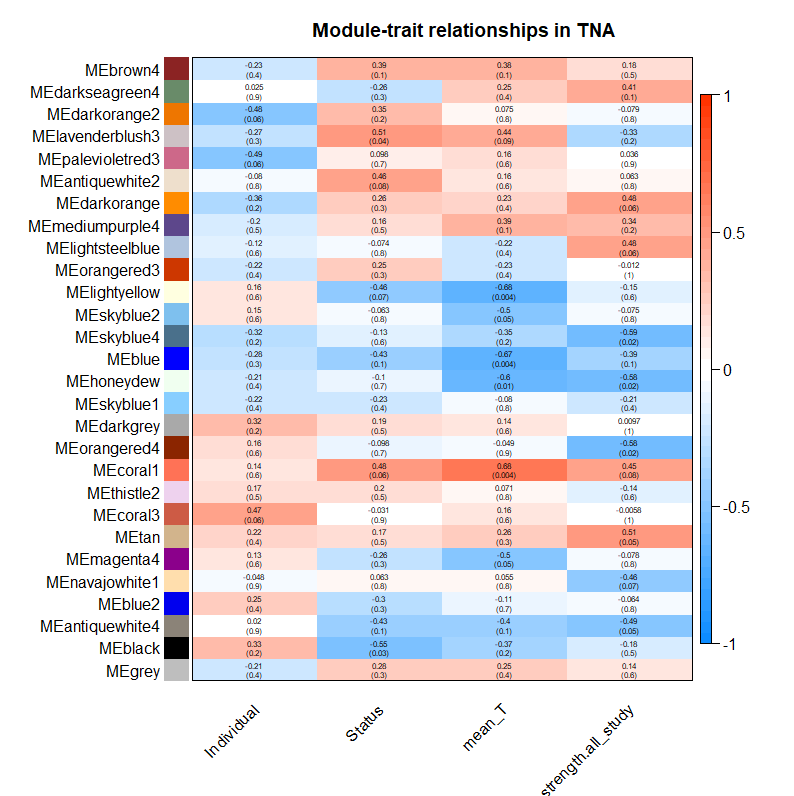

###### Figure S5.10: Correlation between gene co-expression modules and traits in the nucleus taenia (TnA).

Explanation as in Figure S5.1.

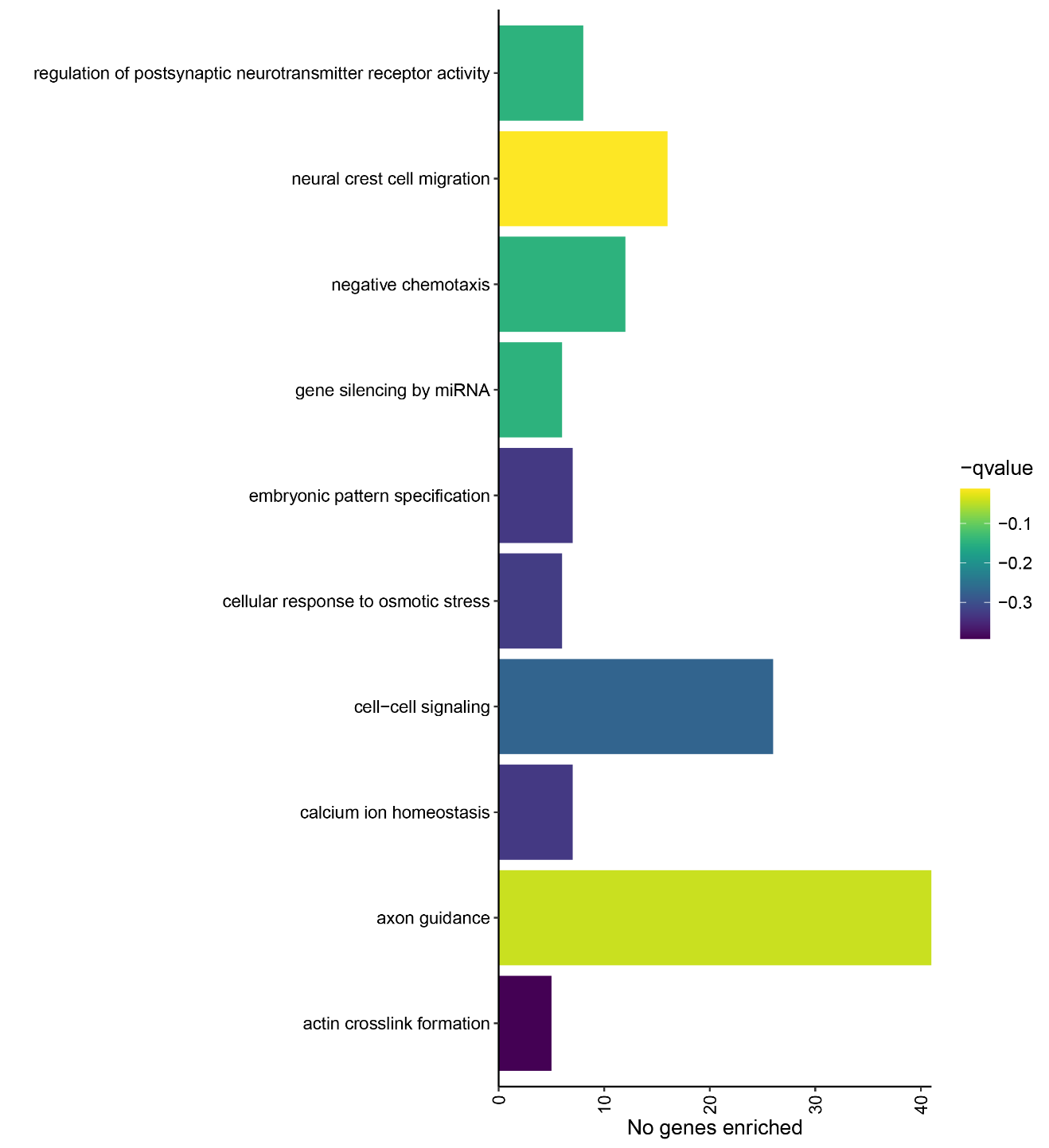

###### Figure S5.11: Biological Process GO enrichment in the TnA’s ‘coral1’ module.

Where bar color indicates the FDR corrected p-value (q-value).

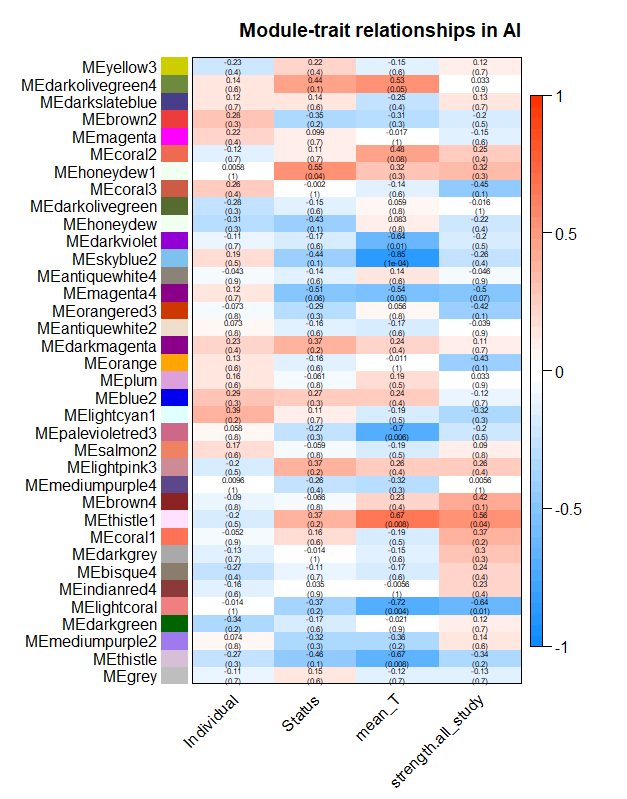

###### Figure S5.12: Correlation between gene co-expression modules and traits in the arcopallium intermedium (AI).

Explanation as Figure S5.1

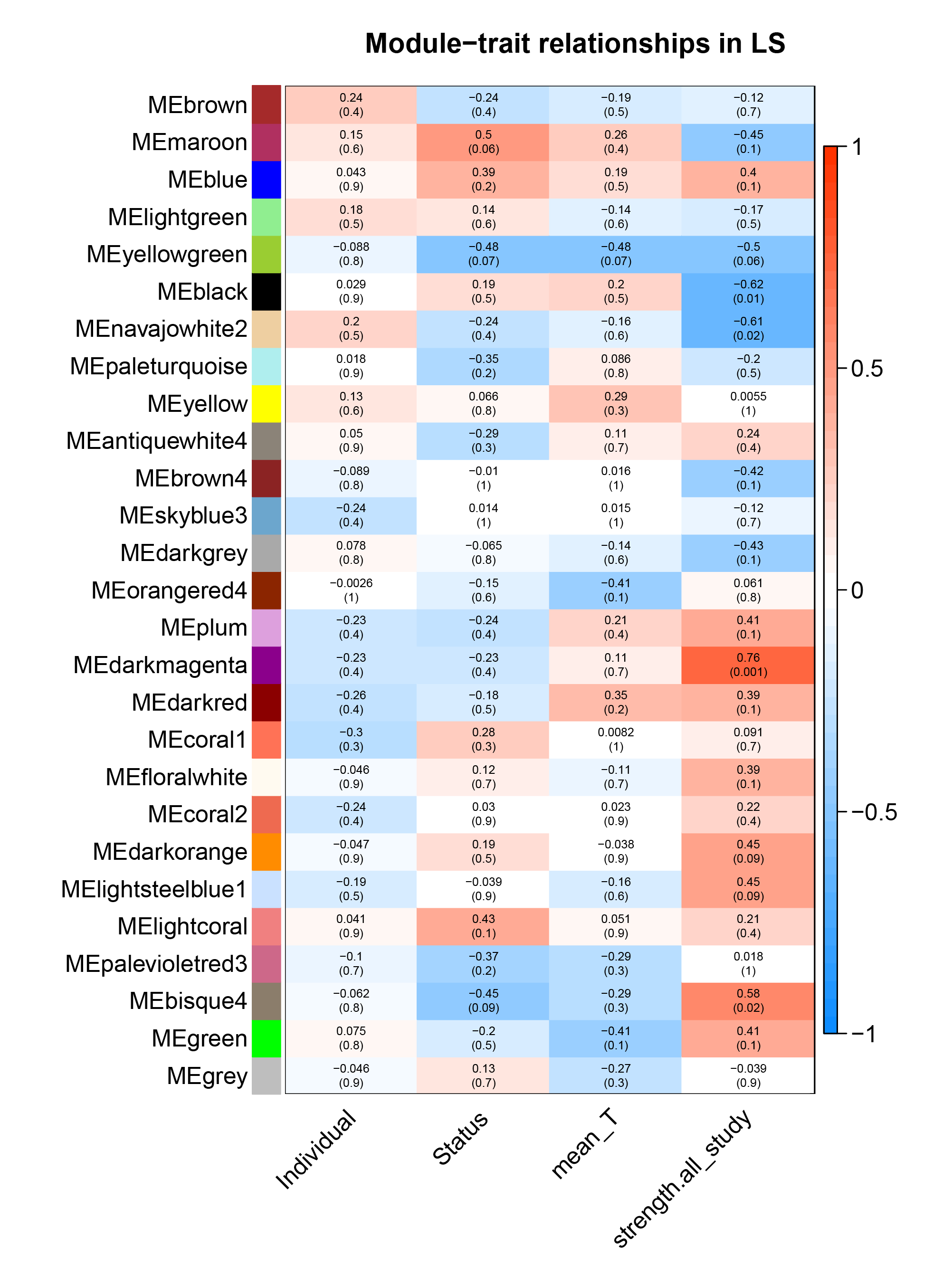

###### Figure S5.13: Correlation between gene co-expression modules and traits in the lateral septum (LS).

Explanation as in Figure S5.1

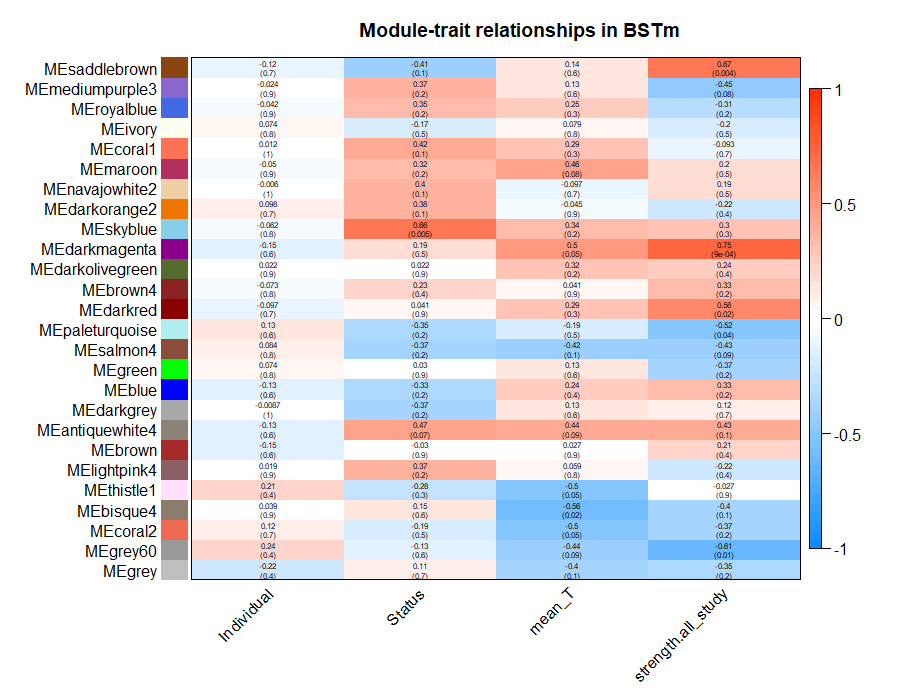

###### Figure S5.14: Correlation between gene co-expression modules and traits in the bed nucleus of the stria terminalis (BSTm).

Explanation as in Figure S5.1.

### 6. Overlap Analyses

#### Methods

There were two kinds of overlap analyses conducted for this work, one was aimed at describing overall patterns of genes with similar expression across the brain, and the other describing similarities between gene expression profiles in each pair of tissues.

To test whether there were global signatures of gene expression in the brain in relation to our variables of interest, we used uncorrected p-values from each single tissue differential expression analysis (Supplement 3). The gene lists were ranked by -log_10_ transforming the p-values and multiplying by the direction of expression. Genes that had a median p-value <0.05 (transformed value >|1.3|) were considered consistently differentially expressed. Gene Ontology enrichment for biological process was determined using custom annotations (Supplement 3) and foreground-background method using the *enricher()* function in the *clusterProfiler* v3.16.1 in R v4.0.2 (Yu *et al.* 2012; R Core Team 2020).

To test whether related brain tissues showed similar overall gene expression profiles in relation to our three interest variables we conducted Pearson correlation on the above log-transformed p-values between each tissue for each trait (social status, individual mean T, and social network strength). Results were plotted using R package *pheatmap* v1.0.12 (Kolde 2019). The clustering hierarchy was tested using 1,000 bootstrap resamples, and clusters with bootstrap probability >95 were indicated using a black square.

For clarity on where in the ordered gene list these similarities arose, for all tissue pairs we conducted Rank-Rank-Hypergeometric Overlap test with R package *RRHO2* (Cahill et al. 2018). This method compares the identity of genes in short sections of the ordered gene lists and tests significance using the hypergeometric distribution. The visual results of this package are not corrected for multiple comparisons. However, because we had a large number of pairwise comparisons, we also visualized the resultant plots scaled against the tissue comparison with the highest p-value. This would give a visual indication of which comparisons would be significant after a correction for multiple testing, but we did not know of any method that would not be too conservative given the number of comparisons (genes x genes + tissue x tissue).

#### Results

##### Social status and mean testosterone

When considering what genes have globally similar expression we found, as expected, that many of the same genes were associated with both status and testosterone, and many were also represented in the brain PCA analysis (Figure S2.1-2.3, PC5). There were 39 consistently upregulated genes (median p<0.05), and 21 consistently downregulated genes in territorial individuals, and 30 upregulated, 36 downregulated in individuals with high mean testosterone (see [GitHub](https://github.com/periperipatus/PIFI_brain_transcriptome/tree/main/DE_results) for gene list).

For example, there was significant GO enrichment for immune response genes in association with testosterone phenotype (Figure S6.2,Table S6.2), and many of these same genes were differentially regulated across social classes (e.g., C4A-like, C3, PLA3G4A, CLEC3B, CLEC2B-like). There were many paralogs of glutathione-S-transferases (GO:0006749, q=0.03) consistently differentially regulated (up LOC113990807, LOC113990865, down LOC113990605), and although these have diverse cellular functions, one of their notable roles is in testosterone and progesterone biosynthesis (GSTA3), and many genes involved in cholesterol biosynthesis and transport were consistently differentially expressed (LCAT, GGPS1, SC5D, CYPJ2J).

When only considering consistent regulation with respect to social status, there were no statistically significant GO enrichments after FDR correction, but many GO categories were represented in the consistently differentially expressed genes (Table S6.1). One of the notable categories was ‘regulation of cell migration’ (GO:0030334, p=0.03) containing two developmental genes, including ROBO4, which was consistently upregulated in territorial individuals (Figure 4). The top upregulated gene in territorial individuals is an immune related gene, a possible paralog of CLEC2B (LOC113988442, C-type lectin domain 2B-like). Glutamate receptor GRINA was consistently upregulated in territorial individuals (and high testosterone), suggesting enhanced post-synaptic excitation in this status class. Finally, social status showed consistent differential regulation of telomere maintenance pathways (POLD-1-like (LOC114000574), RAD5, CC4).

##### Social network strength (cooperative behavior)

There were 10 genes consistently upregulated (median p<0.05) and seven consistently downregulated across the brain in association with a male’s cooperative phenotype (see archived data for gene list. Amongst these there was no statistically significant patterns of GO enrichment after FDR correction (Table S6.3), but some interesting pathways were represented. For example, inflammatory response genes (GO:0006954, q=0.08: CDO1, CSF1R) were marginally enriched, and CDO1 is also involved in the stress response. A single gene (C2D3) represented mild enrichment for neural tube development (Table S6.3). Some genes were shared in regulatory pattern with results from testosterone phenotype, such as downregulation of a glutathione-s-transferase (LOC113990605 Figure S6.4B) and immune CLEC3B. The upregulated gene with the highest median p-value was SLC4A1AP (Kanadaptin) (Figure S6.4A), meaning individuals that high strength of cooperative behavior tended to universally express more SLC4A1AP. Little is known about this gene, but gene-wide association studies in humans have associated this gene with functions including serum testosterone and sex hormone binding globulins (Buniello *et al.* 2019).

##### Between tissue analyses

The analyses describing raw correlation between gene expression profiles between tissues is a blunt instrument. This approach does not provide information on the expression direction in which genes are similar, or rank in the gene list. The results from these analyses for testosterone and mean T are presented in Figure 3, with unscaled outputs presented in for the social status (Figure S6.1). The RRHO2 analysis provides information on where in the ordered gene list the similarities of expression are occurring. The full set of pairwise comparisons that are not scaled to the tissue comparison with the largest log_10_P-value are presented [here](https://github.com/periperipatus/PIFI_brain_transcriptome/tree/main/Overlap_results/RRHO2). The full set of scaled comparisons, which we refer to in the main text (Figure 3), is also viewable [here](https://github.com/periperipatus/PIFI_brain_transcriptome/tree/main/Overlap_results/RRHO2). The main text also focuses on the similarity between LS and BSTm, and the unscaled comparisons are provided in Figure S6.3. These show that there are some genes that have similar expression in relation to mean T and Status, but the strength results suggest that the middle of those gene lists are much more similar than any fragment in the other comparisons.

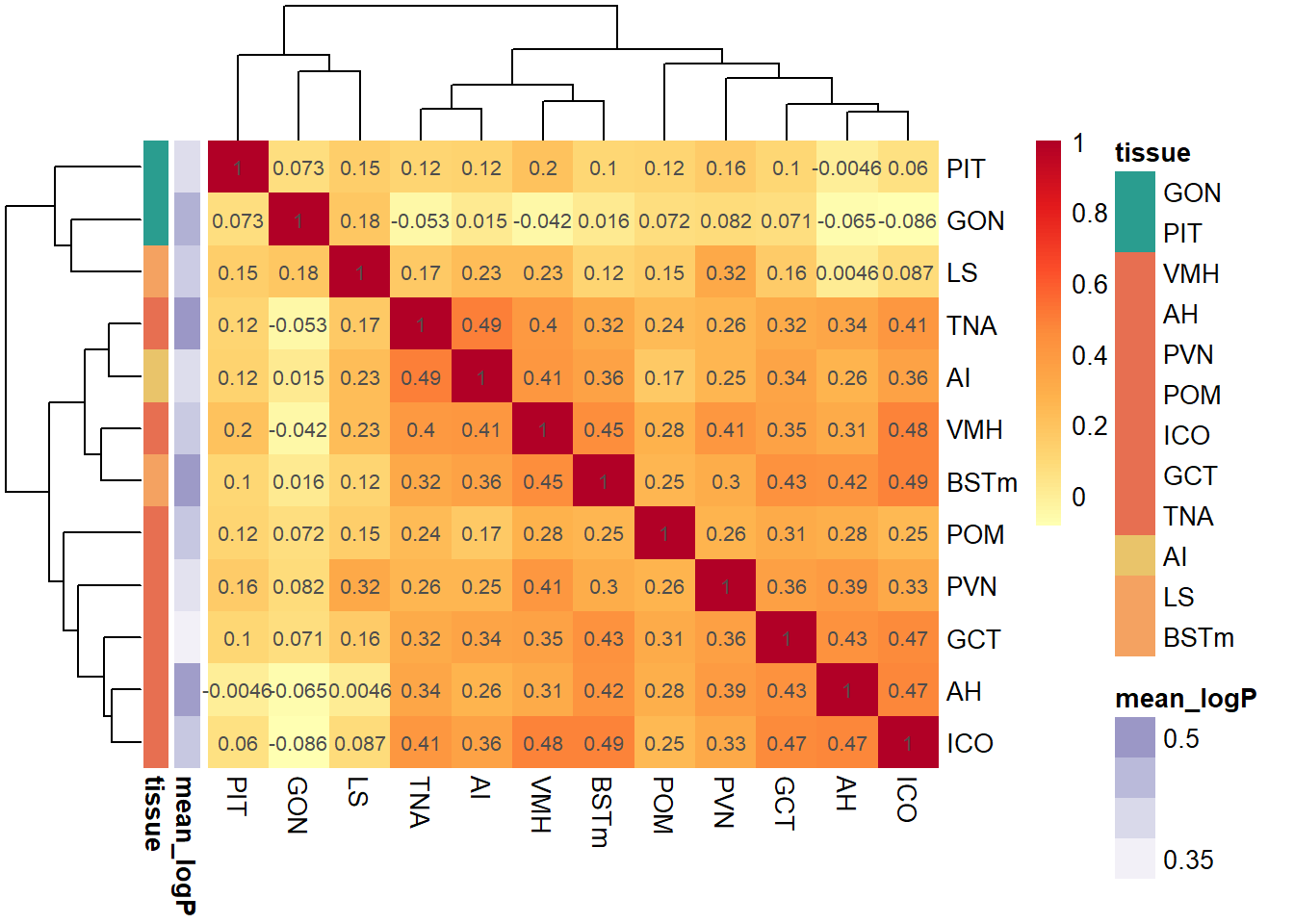

###### Figure S6.1: Similarity of gene expression profiles between tissues in association with social status.

Grid square color and number are the Pearson correlation coefficients of gene expression between tissues, correlating log10 transformed differential expression p-values (times the direction of differential expression). Black boxes outline significant tissue correlation clusters based on 1,000 bootstrap resamples of the correlation matrix.

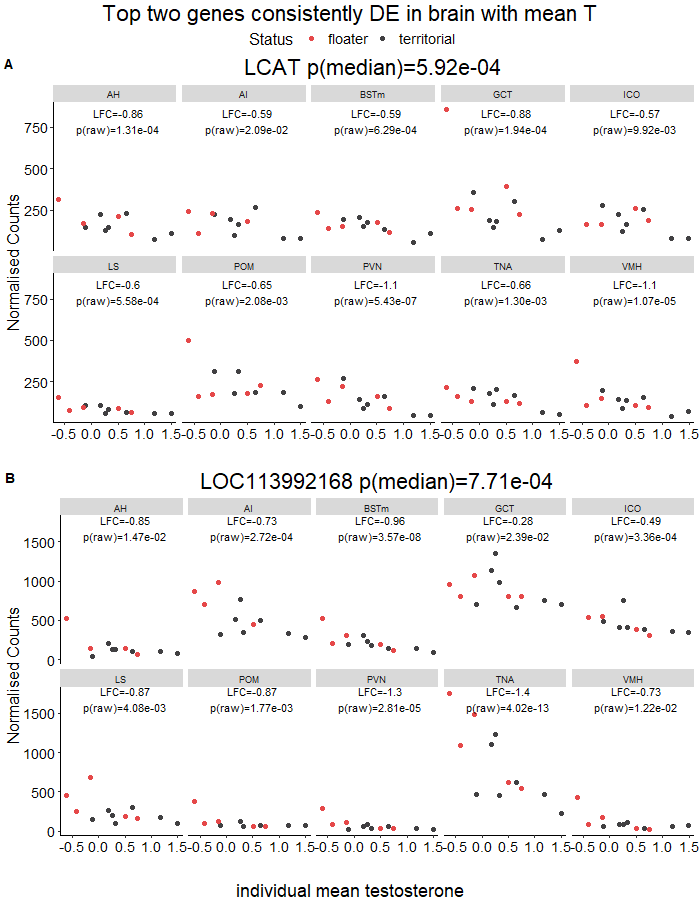

###### Figure S6.2: The top two genes that were consistently downregulated in individuals with high mean testosterone.

A) The strongest median p-value was associated with the gene LCAT involved in cholesterol transport. B) LOC113992168 (C4 complement-like) is an immune related gene that was consistently downregulated in individuals with high mean testosterone.

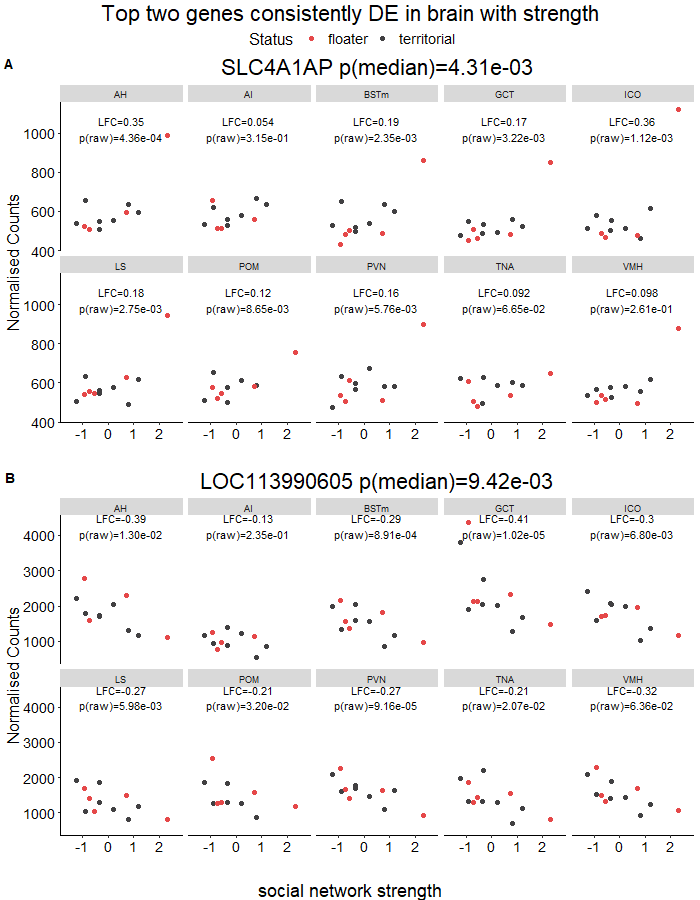

###### Figure S6.3: The top two genes that were consistently up- and downregulated in individuals with high strength of cooperative behavior.

A) The most consistently upregulated gene was SLC4A1AP. B) The most consistently downregulated gene was LOC113990605 (GSTA5-like).

###### Figure S6.4: Similarity of gene expression profiles between LS and BSTm for social status (left) testosterone (middle) and strength (right).

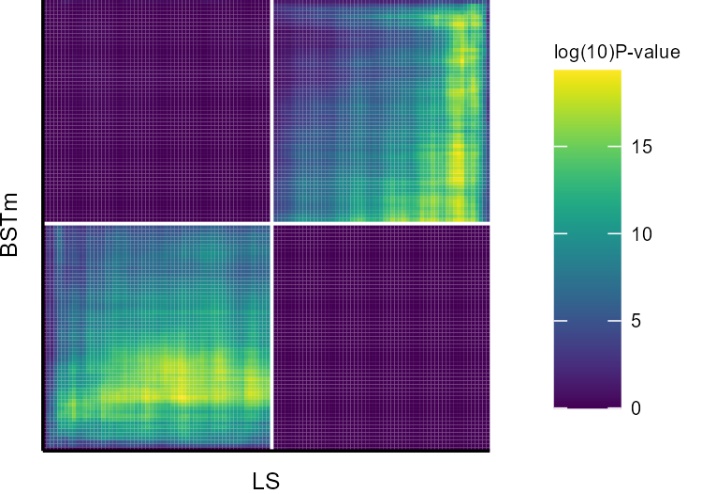

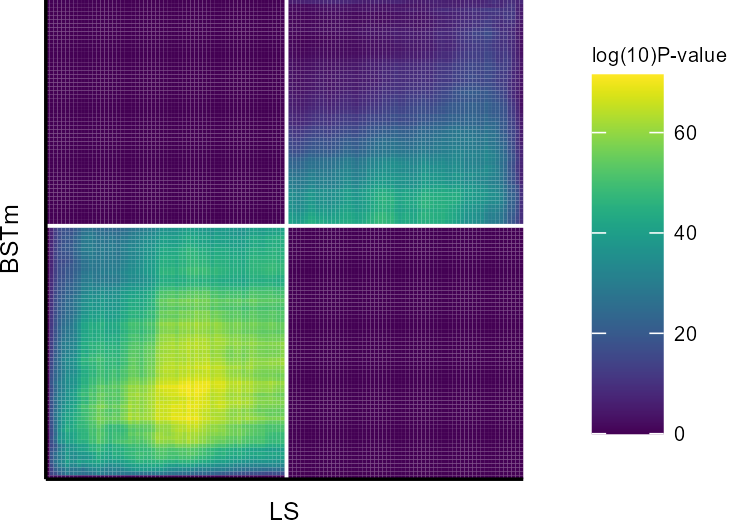

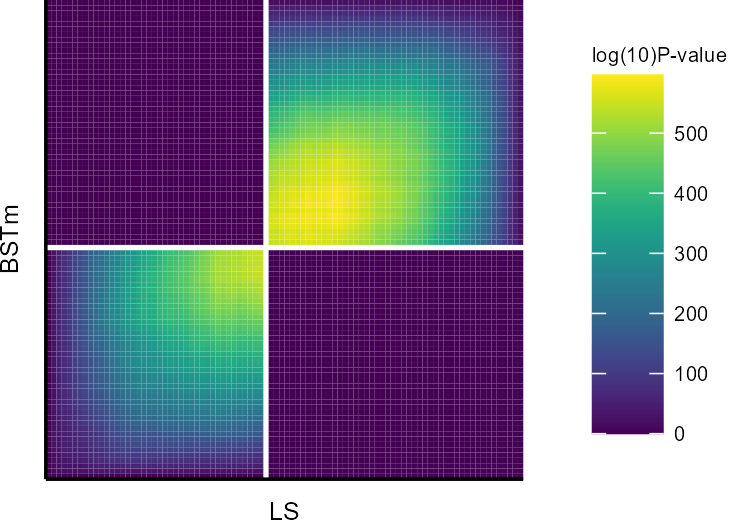

This method takes an ordered gene list (ranked p-values with sign indicating direction of expression) and breaks it up into fragments and compares gene identity among fragments. The white lines represent zero in the ordered gene list, and the axes are the direction of expression for each tissue. Colors represent the raw log_10_ p-value from each fragment’s identity comparison, so note the scale on each legend. The full suite of these RRHO analyses can be found on the [GitHub](https://github.com/periperipatus/PIFI_brain_transcriptome/tree/main/Overlap_results/RRHO2).

###### Table S6.1: The top 10 GO enrichment results for genes that have consistently low p-values across brain tissues in association with social status

The full gene list is deposited with the archived data.

| **ID** | **Description** | **p-value** | **q-value** | **Gene IDs** |
| --- | --- | --- | --- | --- |
| GO:0010951 | negative regulation of endopeptidase activity | 0.0016 | 0.23 | ITIH5/LOC113991485/LOC113992168 |
| GO:0043085 | positive regulation of catalytic activity | 0.0034 | 0.23 | FGFR4/SLC5A3 |
| GO:0045540 | regulation of cholesterol biosynthetic process | 0.008 | 0.23 | GGPS1/SC5D |
| GO:0009636 | response to toxic substance | 0.019 | 0.23 | LOC114001565/RAD51 |
| GO:0006936 | muscle contraction | 0.024 | 0.23 | GAMT/ITGA1 |
| GO:0030334 | regulation of cell migration | 0.028 | 0.23 | MEAK7/ROBO4 |
| GO:0000731 | DNA synthesis involved in DNA repair | 0.039 | 0.23 | LOC114000574 |
| GO:0007602 | phototransduction | 0.039 | 0.23 | PLEKHB1 |
| GO:0034356 | NAD biosynthesis via nicotinamide riboside salvage pathway | 0.039 | 0.23 | LOC113994416 |
| GO:0045651 | positive regulation of macrophage differentiation | 0.039 | 0.23 | IL34 |

###### Table S6.2: The top 10 GO enrichment results for genes that have consistently low p-values across brain tissues in association with mean testosterone.

The full gene list is deposited with the archived data.

| **ID** | **Description** | **p-value** | **q-value** | **Gene IDs** |
| --- | --- | --- | --- | --- |
| GO:0006958 | complement activation, classical pathway | 0.00005 | 0.007933 | C3/LOC113984271/LOC113992168 |
| GO:0006805 | xenobiotic metabolic process | 0.000069 | 0.007933 | LOC113990605/LOC113990807/ LOC113990865/LOC113991532 |
| GO:0006749 | glutathione metabolic process | 0.000368 | 0.028156 | LOC113990605/LOC113990807/ LOC113990865 |
| GO:0006469 | negative regulation of protein kinase activity | 0.002232 | 0.075498 | EPHA1/LOC114002160/TESC |
| GO:0006956 | complement activation | 0.002346 | 0.075498 | C3/LOC113992168 |
| GO:0033628 | regulation of cell adhesion mediated by integrin | 0.002346 | 0.075498 | LOC114002160/TESC |
| GO:0051604 | protein maturation | 0.002346 | 0.075498 | LOC114002160/TESC |
| GO:1901687 | glutathione derivative biosynthetic process | 0.002632 | 0.075498 | LOC113990807/LOC113990865 |
| GO:0007131 | reciprocal meiotic recombination | 0.004293 | 0.109468 | RAD51D/SYCE3 |
| GO:0030449 | regulation of complement activation | 0.005892 | 0.135201 | C3/LOC113992168 |

###### Table S6.3: The top 10 GO enrichment results for genes that have consistently low p-values across brain tissues in association with strength.

The full gene list is deposited with the archived data.

| **ID** | **Description** | **p-value** | **q-value** | **Gene IDs** |
| --- | --- | --- | --- | --- |
| GO:0006954 | inflammatory response | 0.007 | 0.08 | CDO1/CSF1R |
| GO:0070374 | positive regulation of ERK1 and ERK2 cascade | 0.008 | 0.08 | CSF1R/PRXL2C |
| GO:0006488 | dolichol-linked oligosaccharide biosynthetic process | 0.02 | 0.08 | ALG1 |
| GO:0000266 | mitochondrial fission | 0.02 | 0.08 | MTFR1 |
| GO:0006998 | nuclear envelope organization | 0.02 | 0.08 | SUN1 |
| GO:2000249 | regulation of actin cytoskeleton reorganization | 0.02 | 0.08 | CSF1R |
| GO:0008589 | regulation of smoothened signaling pathway | 0.02 | 0.08 | C2CD3 |
| GO:0071539 | protein localization to centrosome | 0.02 | 0.08 | C2CD3 |
| GO:0021915 | neural tube development | 0.03 | 0.08 | C2CD3 |
| GO:0030316 | osteoclast differentiation | 0.03 | 0.08 | CSF1R |
